# Efficient scar-free knock-ins of several kilobases by engineered CRISPR/Cas endonucleases

**DOI:** 10.1101/2023.11.24.568561

**Authors:** Tom Schreiber, Anja Prange, Petra Schäfer, Thomas Iwen, Ramona Grützner, Sylvestre Marillonnet, Alain Tissier

## Abstract

In plants and mammals, non-homologous end-joining is the dominant pathway to repair DNA double strand breaks, making it challenging to generate knock-in events. Using a transient assay in *Nicotiana benthamiana* we identified two groups of exonucleases, respectively from the Herpes Virus and from the bacteriophage T7 families, hat confer up to 38-fold increase in HDR frequencies when fused to Cas9. We achieved precise and scar-free insertion of several kilobases of DNA both in transient and stable transformation systems. In *Arabidopsis thaliana*, fusion of Cas9 to a Herpes Virus family exonuclease leads to 10-fold higher frequencies of knock-ins. Our results open perspectives for the routine production of knock-in and gene replacement events in plants.

**One-Sentence Summary:** Fusions of CRISPR endonucleases to specific 5′-exonucleases leads to significant increase in scar-free multikilobase knock-ins.

## Introduction

The discovery of CRISPR systems and their adaptation for gene editing purposes have significantly improved our capacity to introduce targeted modifications in the genome of living organisms. CRISPR/Cas endonucleases such as Cas9 or Cas12a introduce a double strand break in the DNA, which upon error-prone repair leads to indels leading to the production of knock-outs in coding sequences. There is however strong interest for the targeted insertion of sequences, for example to correct genetic defects or to introduce specific variants in DNA. The modification of individual bases or insertion of small and larger sequence stretches is possible by base editors, prime editing or CRISPR-associated transposons (Anzalone et al., 2020). However, these approaches do not permit the precise integration or replacement of larger DNA fragments of foreign sequences without footprint. Homology directed repair (HDR), a precise form of DSB repair, allows the scar-free integration or replacement of large DNA fragments but occurs at low frequency, typically a few percent or less of the transformation events, in plants and mammalian cells (Lieber, 2010; Wang et al., 2023). In these organisms, the repair of DSBs occurs primarily via non-homologous end joining (NHEJ). All repair pathways of DSBs via HDR require the processing of DSB ends to produce longer free 3’ ends (>25bp) (Lieber, 2010; Schmidt et al., 2019). CRISPR endonucleases like Cas9 or Cas12a generate blunt ends and 5’ sticky ends (of 7 nt), respectively (Shi et al., 2019; Stella et al., 2017). We reasoned that fusion of 5’-exonucleases to CRISPR endonucleases could lead to production of free 3′ends and thereby to increased rates of HDR.

Here we demonstrate that fusion of 5′-exonucleases from the Herpes Virus family to Cas9 and Cas12a are the most efficient at enhancing HDR, with frequencies up to 38-fold higher in a transient assay system and over 10-fold for stable knock-ins in Arabidopsis. The fusion proteins developed here represent promising tools for gene targeting in plants and possibly in other organisms.

## Results

### 5′-exonuclease::Cas9 fusions increase homology directed repair efficiency

To assess the efficiency of Cas9 endonucleases fusions for HDR, we developed a transient assay based on a transgenic *Nicotiana benthamiana* line that contains a defective tobacco mosaic virus (TMV) viral vector harboring a GFP gene (Fig. 1A and fig. S1, see Note S1). Expression of GFP only occurs if the defective endogenous copy of the RNA-dependent RNA polymerase (RdRP) is precisely repaired via HDR using an appropriate donor template. Repair takes place after introduction of a DSB in the attB site using Cas9 and one or several guide RNAs, and co-delivery of a T-DNA carrying the repair template. The presence of the viral movement protein in the reconstituted viral vector allows the TMV to spread locally from cell to cell, leading to amplification of the GFP signal from individual events and results in GFP spots that can be easily counted (Fig. 1B and C, fig. S2).

**Figure 1.**
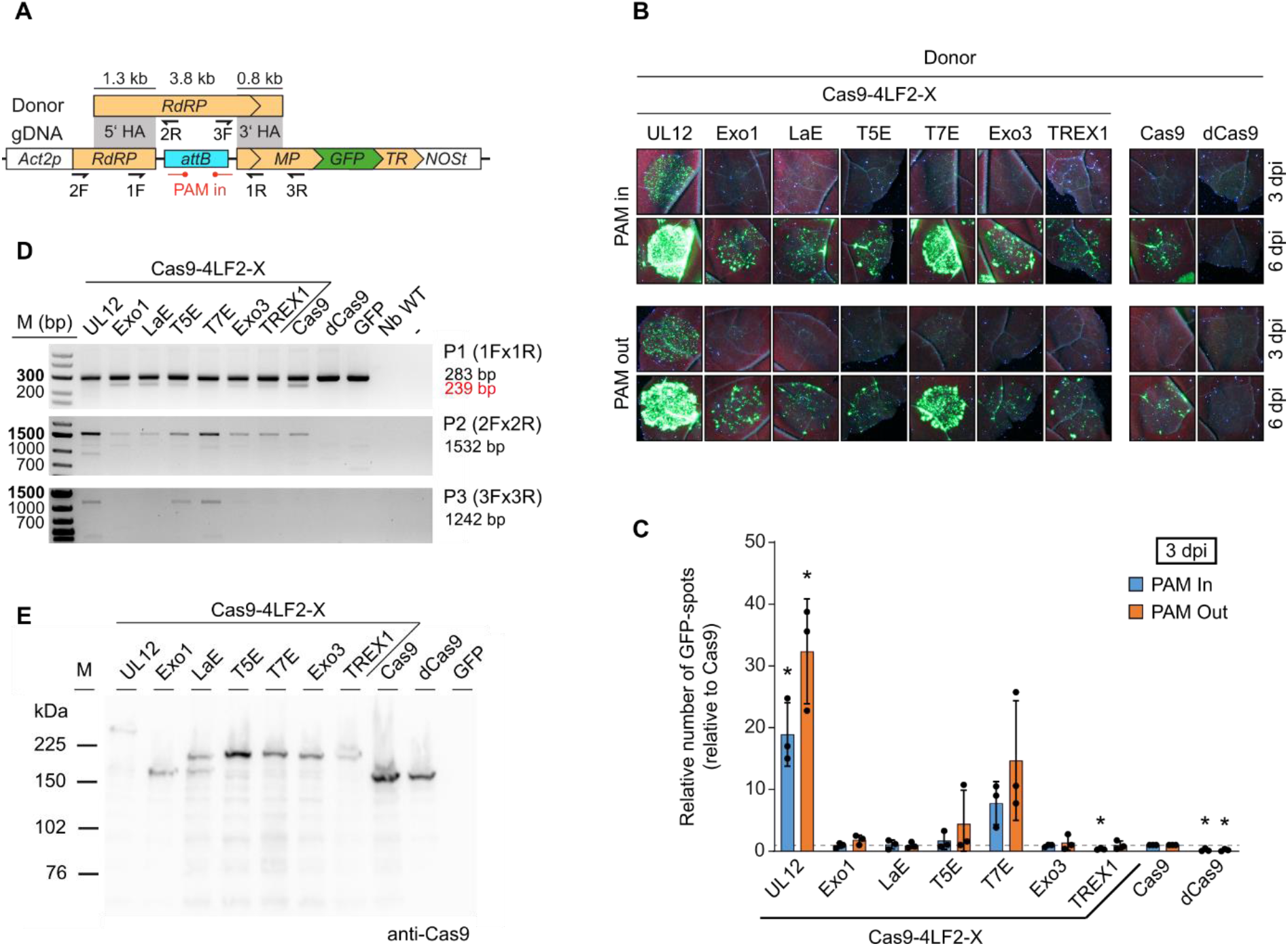
Fusion of UL12 and T7 5′ to 3′ exonucleases to Cas9 increases HDR efficiency. **(A)** Transgenic construct containing the defective TMV genome with a GFP reporter (bottom) and donor template used to repair the defective RdRP gene. **(B)** GFP signals of transgenic *Nicotiana benthamiana* leaves three and six days after inoculation (dpi) of individual exonuclease-Cas9 fusions together with sgRNAs and the TMV repair donor. Two sgRNAs are used, either in the PAM-In or PAM-Out orientation. Deactivated Cas9 (dCas9) serves as negative control. **(C)** Quantification of HDR efficiency by GFP spot count relative to Cas9. Plotted is the fold change of GFP spots from three infiltrated leaf areas, each from a distinct plant, with standard deviation relative to Cas9. Significance was evaluated with Student′s t-test; * p-value ≤ 0.05; ** p-value ≤ 0.01; *** p-value ≤ 0.001. **(D)** PCR-based genotyping of repair events. Primer pairs P2 (2F + 2R) and P3 (3F + 3R) amplify the upstream and downstream junction, respectively. Primer pair P1 (1F + 1R) is used for on-target amplification and monitors the deletion between the two cut sites, an event that is visible as a second smaller band (239 bp). Note that the fusion of UL12 or T7 exonucleases to Cas9 leads to a strong reduction in the intensity of the lower band. Expected fragment sizes are given on the right panel. **(E)** Detection of Cas9 and Cas9 fusion proteins from leaf extracts by Western blot using Cas9-specific antibody.

We first tested the fusion of Cas9 to 5’-exonucleases from bacteriophages T5, T7 and λ, to UL12 exonuclease from Herpes Simplex Virus 1 (HSV1), and to Arabidopsis thaliana Exo1. In addition, we also tested Exo III from Escherichia coli and human TREX1, which are 3’-exonucleases. C-terminal fusions to Cas9 showed that UL12 and T7-exonuclease (T7E) strongly increase the number of GFP spots compared to unmodified Cas9. UL12 leads to increases ranging from 15- to 38-fold with GFP spots visible already 3 days post inoculation (dpi, Fig. 1B; fig. S2D). Fusion of Cas9 to T7E leads to a 4 to 26-fold improvement over Cas9. By contrast, other 5’-exonucleases such as Arabidopsis Exo1, λ-exonuclease (LaExo), T5E and 3’-exonucleases (Exo III and TREX) did not increase frequencies of GFP spots (Fig. 1B). We observed similar results with the N-terminal fusions (fig. S2). Genotyping by PCR and sequencing of genomic DNA isolated from infiltrated leaf discs confirms the precise repair of the TMV RdRP gene (Fig. 1D, fig. S3). Using a Cas9-specific antibody, we show by Western blot that all fusion proteins except Cas9::Exo1 are correctly expressed (Fig. 1E).

We next checked the ability of a subset of the exonuclease-Cas9 fusions (UL12, T7E and T5E) to increase knock-in frequencies at two endogenous loci in *N. benthamiana* that are constitutively expressed (Phosphoglycerate kinase family protein (NbPGK) and Tetratricopeptide repeat transcription factor (NbTPR)) by introducing a translational fusion to the GUS reporter. For these two loci we see a significant increase of knock-in frequencies induced by exonuclease fusions compared to Cas9 (Fig. 2, fig. S4 and S5). In conclusion, fusion of UL12 or T7E to Cas9 significantly increases gene targeting efficiency in *N. benthamiana*.

**Figure 2.**
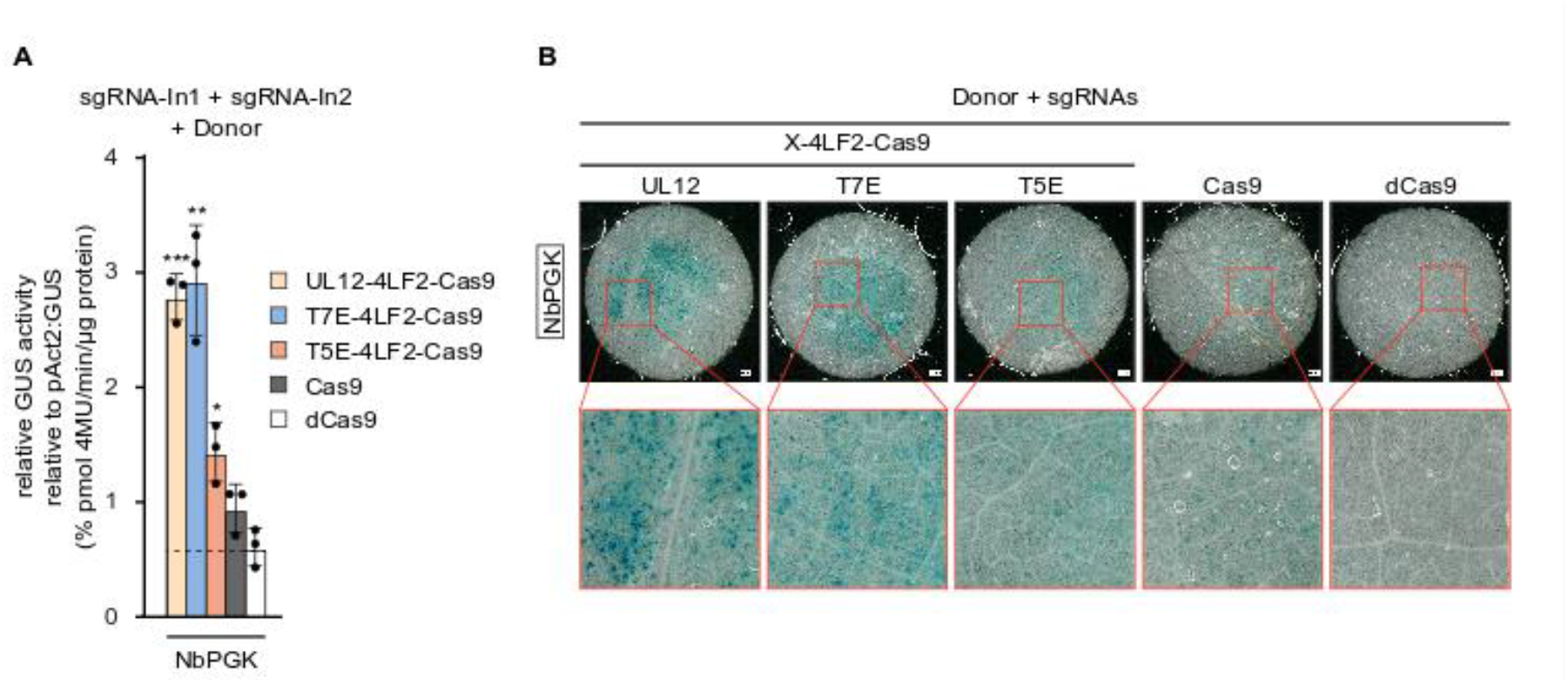
Exonuclease-fused Cas9 used for in frame GUS integration into a Nb endogene. **(A)** An in-frame intronized GUS sequence was used as donor with corresponding homology arms for the targeted insertion downstream of either a gene encoding Phosphoglycerate kinase family protein (*NbPGK*). Quantitative GUS assay from leaf extracts harvested 3 dpi from individual inoculated leaf areas. Values are relative to pAct2-driven GUS. Significance was evaluated with Student′s t-test; * p-value ≤ 0.05; ** p-value ≤ 0.01; *** p-value ≤ 0.001. **(B)** GUS stained leaf discs, harvested 3 dpi from corresponding inoculation spots. Blue spots indicate cells with gene targeting events.

**Figure 3.**
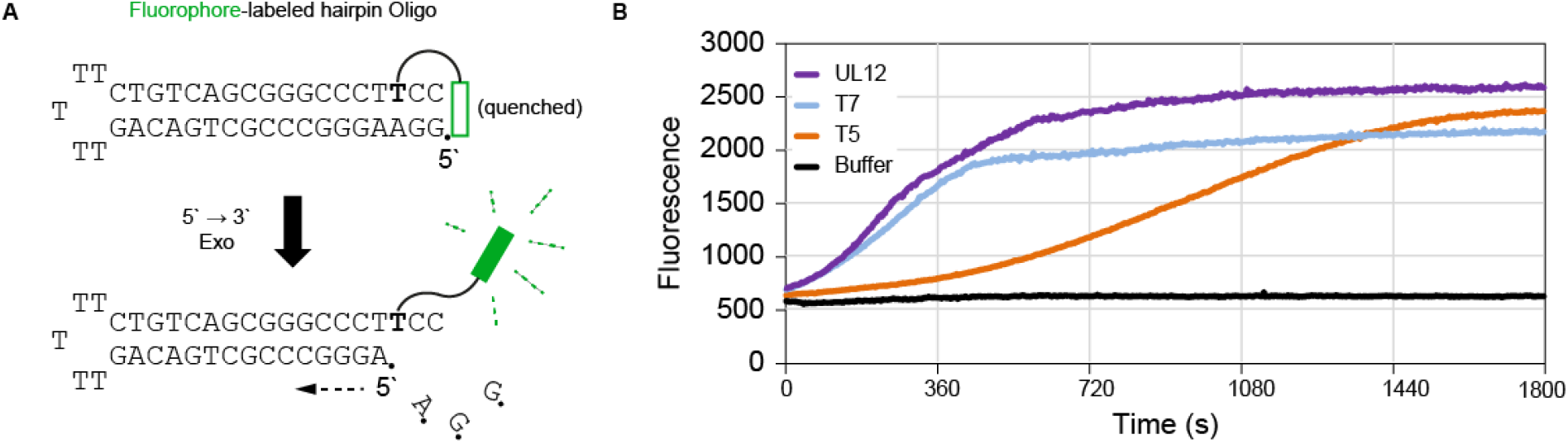
UL12 and T7E possess high exonuclease activity on blunt ended DNA substrates. **(A)** Schematic description of the fluorogenic in vitro assay based on fluorophore-labeled hairpin oligonucleotides. The fluorophore is quenched by stacking to the G-C base pair at the end of the hairpin. Exonucleolytic degradation of the oligo releases the fluorophore from quenching and results in fluorescent signals. **(B)** Fluorescence intensity plotted over time (30 minutes) after recombinant exonucleases were added to the reaction.

### Linkers, length of homology arms and co-expression with single-strand DNA binding proteins

Tests with different linkers for the fusions showed that the 4xLF2 is the best while the position of the fusion, either as N- or C-terminal did not have a significant impact (fig. S6). We then tested homology arms (HA) of different lengths (100, 250, 500 bp) and found that 250 bp was sufficient and gave the highest HDR frequencies (fig. S7). We did not observe significant differences when only one sgRNA instead of two were used (Fig. S8). We also found that paired single-stranded nicks generated by Cas9 nickases fused to UL12 lead to increased HDR albeit with reduced efficiency compared to the Cas9 fusions (fig. S9, see Note S2).

The N-terminal region of UL12 (amino acids 1-126) was shown to interact with the Mre11-Rad50-Nbs1 (MRN) complex, which is responsible for DSB sensing and repair initiation (Balasubramanian et al., 2010). Fusion of this UL12 MRN-recruitment domain (N1-126) to Cas9 was also shown to slightly enhance HDR efficiency in human cells (Reuven et al., 2019). In order to dissect the relative contribution of the MRN recruitment domain or of the exonuclease activity of UL12 to increased HDR efficiency we analyzed UL12 variants truncated at the N-terminus or catalytically dead fused to Cas9 (fig. S10A and B). This showed that exonuclease activity of UL12 is essential for increased HDR efficiency with a minor contribution of the MRN recruiting domain. Since additional proteins, in particular single-strand DNA binding (SSB) proteins or proteins with DNA strand exchange activity (single strand annealing proteins - SSAPs), play important roles in HDR, we tested whether their co-expression would further increase gene targeting frequencies in our transient assay. None of the proteins tested (SSAPs: RAD51, RAD52a/b, ICP8; SSB: E. coli SSB) have an effect (fig. S10B, C and F). Interestingly, ICP8, the SSAP of HSV which reconstitutes a two-component recombination system together with UL12 (Reuven et al., 2003), reduces gene targeting frequencies triggered by UL12::Cas9. Furthermore, expression of ICP8 leads to reduced general expression rates and tissue collapse especially at later time points, an observation that is independent of nuclear and cytoplasmic localization (fig. S11).

### Exonucleases suited for direct endonuclease fusion

We next investigated why not all 5’-exonucleases lead to increased frequencies of HDR. We compared the exonuclease activity of recombinant UL12 (fig. S12), T7E, T5E and LaExo using a fluorogenic in vitro assay based on a hairpin oligonucleotide with blunt ends (Nikiforov, 2014) (fig. S13A and B). In this assay, UL12 was the fastest with T7E and LaExo slightly slower, whereas T5E was much slower. These results are in good agreement with the relative efficiencies of these exonucleases in the TMV-based transient assay, except for LaExo. However, LaExo acts as a trimer (Kovall and Matthews, 1997), which might explain the absence of effect in our HDR assay when a LaExo monomer is fused to Cas9. Co-expression of free LaExo only slightly improves HDR efficiency but does not reach the levels achieved by UL12::Cas9 fusions (fig. S13C). We next found that when fused to Cas12a, which generates short 5’-sticky ends, UL12 still outperforms T7E and T5E (Fig. 4A and B; fig. S14), despite the preference of T5E for 5’-overhangs (Garforth and Sayers, 1997). To gain further insights into the processing of Cas9- or Cas12a-induced DSBs by exonucleases (UL12-, T7E- and T5E-), we performed amplicon sequencing of the TMV-locus using two sgRNAs in either PAM-in or PAM-out orientation (Fig. 4C and D). Cas9 and Cas12a alone mainly induced deletions between the two cleavage sites, with minor additional DNA end processing according to their cleavage patterns. Exonuclease fusions increase the overall number of larger deletions with an average processing length of approximately 40-60 nt on both PAM proximal and PAM-distal sides for all tested exonucleases. Compared to UL12 and T7E, Cas9-fused T5E shows a lower overall number of larger deletions, confirming its lower affinity for blunt ends (Fig. 4C).

**Figure 4.**
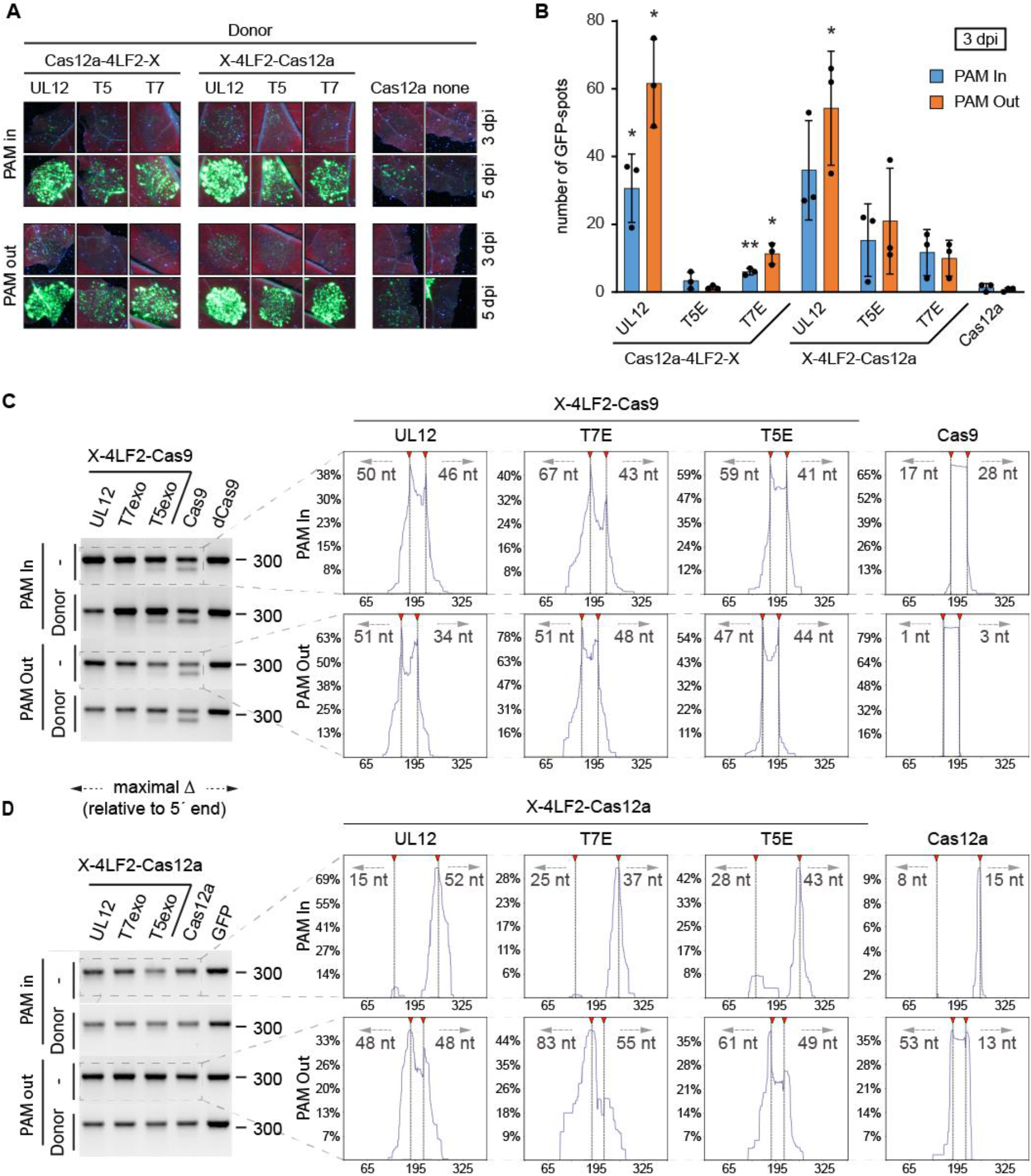
Fusion of 5′ to 3′ exonucleases to CRISPR endonucleases increases the number of larger deletions. **(A)** GFP signals of transgenic *Nicotiana benthamiana* leaves three and five days post inoculation (dpi) of individual exonuclease-Cas12a combinations together with corresponding crRNAs (PAM-In; PAM-out) and the TMV repair donor. **(B)** Quantification of HDR efficiency by GFP spot count relative to Cas12a. Plotted is the average GFP spot number and standard deviation of three individual plants. Significance was evaluated with Student′s t-test; * p-value ≤ 0.05; ** p-value ≤ 0.01; *** p-value ≤ 0.001. **(C and D)** Amplicon sequencing of TMV on target-spanning PCR products (primer pair P1) amplified from isolated genomic DNA after expression of the indicated exonuclease-endonuclease fusion proteins. Gel pictures and corresponding amplicon reads are shown on the left and right panel, respectively. DNA DSB sites are indicated by red triangles (dashed lines) and the length of the largest deletion are given in base pairs counted from the 5’ end of the lesion. Note the weak performance of Cas12a crRNA in1, probably due to the non-canonical CTTC PAM (LbCas12a, fig. S1).

### Homologs of UL12 and T7E exhibit improved HDR performance

UL12 and T7E have 7% identity on the amino acid level and belong to distinct clades of exonucleases (fig. S15). Because both HSV1 and bacteriophage T7 are members of clades of viruses with a large number of variants, orthologs of these exonucleases from these clades could perform better than UL12 or T7E. We selected several orthologs of UL12 and T7E from related viral genomes and generated fusions with Cas9. From the exonucleases tested, the UL12 ortholog from Papiine alpha Herpes Virus 2 (PapE) (Black et al., 2014) and T7 homolog of bacteriophage IME15 (Huang et al., 2012) fused to Cas9 had even higher frequencies of HDR compared to UL12 and T7, respectively (fig. S16 and S17).

### 5′-Exonuclease-Cas9 fusion leads to increased gene targeting in Arabidopsis

We used the optimized exonuclease::Cas9 fusions to repair a disrupted Arabidopsis *THIAMIN C* (*AtTHIC*, At2G29630) gene, which is essential for vitamin B1 biosynthesis (Raschke et al., 2007). The *thiC* mutant harbors a T-DNA insertion in the third intron that leads to a complete loss of function and arrest of growth at the cotyledon stage (Kong et al., 2008). Arabidopsis *thiC* mutant plants can be rescued by external supply of thiamine, and thus be propagated in the homozygous state. Flowering *thiC* mutants were transformed by A. tumefaciens using the floral dip method with three distinct gene targeting constructs harboring either Cas9, PapE::Cas9 or ME15E::Cas9, as well as the donor module and three sgRNAs (Fig. 5A-C). The *thiC* locus was targeted at the junction between genomic DNA and the T-DNA insertion with sgRNA-ThiC (Fig 5C). The other sgRNAs, sgR-gl1 and sgR-D1, target the Arabidopsis GLABRA 1 (GL1) gene, whose mutation leads to trichome-free leaves, a phenotype readily detected by macroscopic inspection of the plants, and the donor module for donor release, respectively. Successfully edited T1 plants where the donor was inserted correctly at the *thiC* locus should survive without external supply of thiamine and in addition possess trichome-free leaves, if in parallel editing at the GL1 locus took place (Fig. 5B, right panel). Strikingly, the PapE::Cas9 fusion increased gene targeting efficiency by 10-fold compared to WT Cas9 (Table 1). By contrast, the ME15::Cas9 fusion only showed a two-fold increase in gene targeting. Sequencing of the junctions showed that events with a perfect repair by the 5’HA and a repair by NHEJ on the 3’ side with complete integration of the THIC transcription unit still resulted in a functional *THIC* gene (fig. S18). With the PapE::Cas9 fusions almost half (12/28) of the plants having a functional *THIC* gene had perfect repair on both sides. A SNP between the donor and the target locus located121 bp from the cut site in the 5’HA allowed us to determine that in 60% of the HDR events, the sequence came from the donor. Thus, in a significant fraction (40%) of successful repair events the conversion track is less than 121 bp, consistent with the data from the transient assay that 250 bp HAs are sufficient to generate high efficiency HDR. Notably, 2 plants out of the 12 with a perfect repair generated by PapE::Cas9 fusions do not contain the T-DNA used to transform the plants (Table 1). This indicates that the HDR events occurred before the T-DNA could integrate in the genome and underscores the efficiency of the PapE::Cas9 fusions.

**Figure 5.**
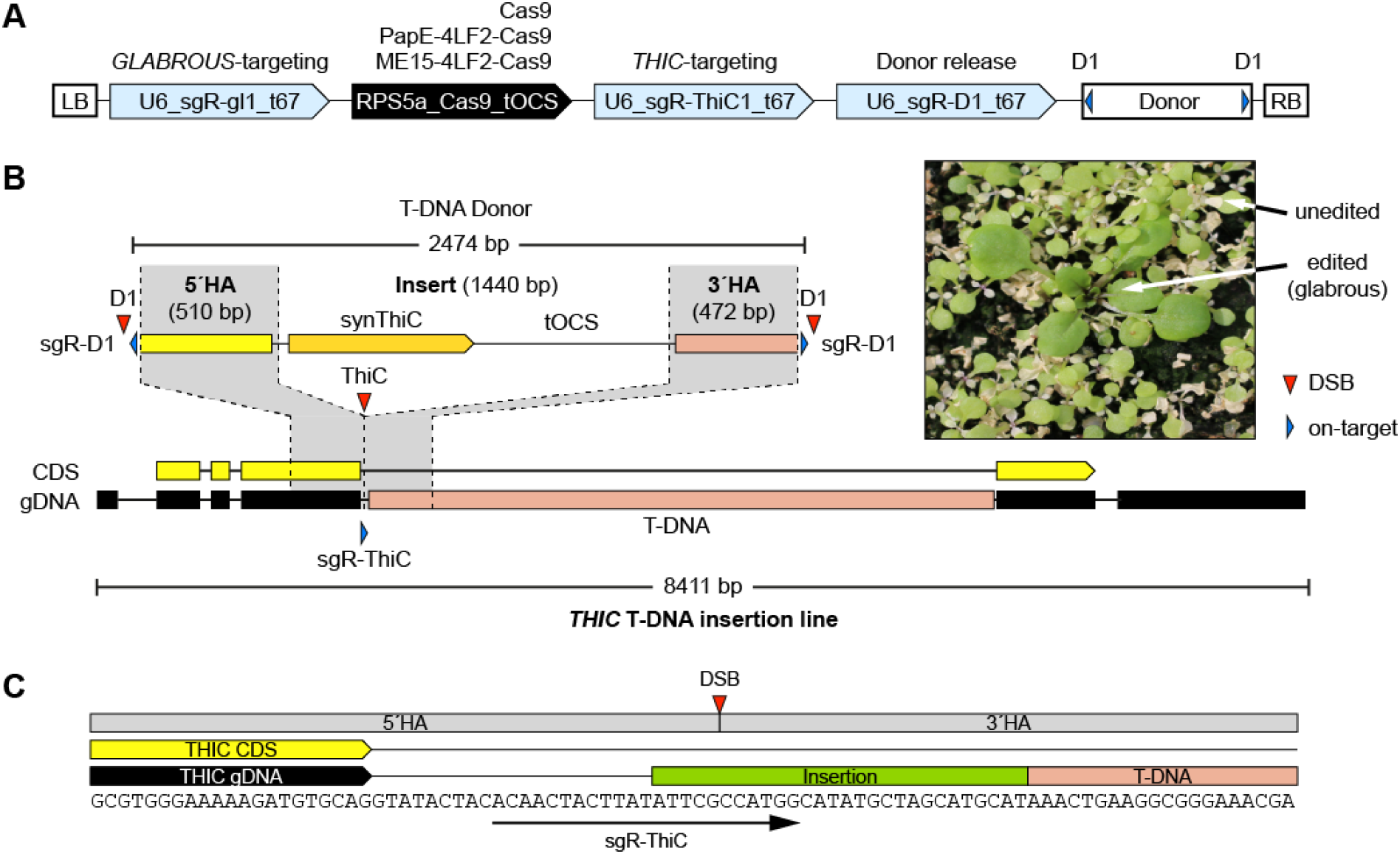
Construct design for a HDR assay at the *THIC* locus in Arabidopsis. **(A)** Schematic overview of constructs transformed into Arabidopsis. All constructs contain sgRNAs for targeting the genomic loci *GLABROUS 1* (sgR-gl1) and *THIC* (sgR-ThiC1) as well as a sgRNA for the donor release (sgR-D1). Endonuclease Cas9 was compared to PapE- and ME15-fused Cas9. **(B)** Schematic overview of the *thiC* locus (THIC T-DNA insertion line) and the donor design. Precise donor integration leads to restoration of the THIC coding sequence (CDS) with a synthetic sequence with a different codon composition compared to the native sequence to avoid unwanted recombination events, followed by the tOCS terminator. 5′ and 3′ homology arms (HA) of 510 bp and 472 bp were used, respectively. The donor sequence was flanked by sgR-D1 target sites in a PAM out orientation. **(C)** Close view of the *thic-2* locus. Target sequence of sgR-ThiC1 is indicated by an arrow (arrow head indicated PAM). 5′and 3′ HAs start directly at the Cas9 cleavage site (indicated by the red triangle).

**Table 1.** Comparison of gene targeting at the thiC locus with PapE::Cas9, ME15::Cas9 and Cas9. Results from three independent transformations. For each transformation the three constructs were transformed in parallel. Three pots with five plants each were transformed.

|  | 5'-HA<br>HDR | FC vs<br>Cas9 | 3'-HA<br>HDR | FC vs<br>Cas9 | % of BsaI<br>site repaired | Absence of the T-DNA in HDR events |
| --- | --- | --- | --- | --- | --- | --- |
| Cas9 | 3 | 1 | 1 | 1 | 33 % | 0 |
| ME15::Cas9 | 6 | 2 | 2 | 2 | 83 % | 0 |
| PapE::Cas9 | 28 | 9.3 | 12 | 12 | 57% | 2 |

## Discussion

The targeted and scar-free integration of several kb of DNA holds much potential in plant breeding and for therapeutic applications. In plants, beneficial alleles, for example from related wild species, could be directly pasted in at the homologous locus in elite cultivars or breeding lines, thereby shortening development times considerably and avoiding linkage drag (Voss-Fels et al., 2017). Although CRISPR/based tools for targeted repair, such as base editors and prime-editing, have been developed in recent years, these are limited to single or less than 100 bases (Anzalone et al., 2022; Zheng et al., 2023). Naturally occurring RNA-guided transposons can be harnessed for the targeted integration of larger payloads, but cannot be used to exchange sequences, generate a footprint and have low efficiencies in human cells (Lampe et al., 2023). In plants, base editors and prime editing have been successfully applied for stable integration of substitutions, short insertions and deletions, but to our knowledge efficient targeted and scar-free integration of several kb has not been achieved so far (Gupta et al., 2023; Lin et al., 2021; Veillet et al., 2019; Zong et al., 2022). The endonuclease::exonuclease fusion system described here allows the targeted integration of large DNA fragments in a seamless fashion and at frequencies that are at least one order of magnitude higher than with Cas9 alone.

The rationale for fusing 5′-exonucleases to CRISPR endonucleases is to rapidly generate longer 3′ ssDNA overhangs, which are known to be substrates for HDR, and thereby prevent processing via the NHEJ machinery. In vivo, competition between the 5′-exonuclease fused to Cas endonuclease and endogenous repair factors for free DNA ends after cleavage is likely to occur. Therefore, one would expect that the time required for the fused exonuclease to start processing the DSBs generated by Cas9 should play a critical role in the outcome of the repair. Various aspects can influence the time to process as well as the exonuclease activity, such as general positioning relative to the endonuclease, affinity of the exonuclease for DSB-ends, accessibility of the DNA substrate, mono- or oligomeric state of the exonuclease or exonuclease processivity. We find that the 5′-exonucleases that are the most active in the in vitro assay, i.e. UL12 and T7, are also the ones that perform best in the HDR assay in *N. benthamiana* (Fig. 1 Fig. 3 and Fig. 4, fig. S13B), suggesting that high affinity for DSBs is crucial. One exception is the λ exonuclease, which has high in vitro activity but is trimeric (Kovall and Matthews, 1997), a feature that is likely to be a disadvantage in the context of a fusion with Cas9. Because UL12 and T7 are monomeric (Banks et al., 1983; Takahashi et al., 2014), we propose that monomeric 5′-exonucleases with high affinity for DSBs are best suited to increase HDR when fused to endonucleases. The fact that relatives of UL12 and T7 exonucleases, respectively PapE and ME15, exhibit further enhanced HDR activity indicates that there is potential for further improvement given the large virus families to which these exonucleases belong. Amplicon sequencing of lesions produced by Cas9 fused to 5′-exonucleases in the absence of the donor revealed that the depth of 5′ end resection is similar for UL12, T7E and T5E with 34-67 nt in both PAM distal and PAM proximal direction for Cas9 fusions and 37-83 nt for Cas12a fusions (Fig. 4C and D). With Cas9 fusions there is a clear difference between T5E and T7E/UL12 in the deletion pattern, with proportionally fewer smaller deletions with T5E than with T7E or UL12, consistent with the results from in vitro assays and confirming that T5E has lower affinity for blunt end DSBs also in vivo. The Cas12a fusions display similar deletion patterns for all exonucleases tested, consistent with the improved HDR activity of the T5E fusions with Cas12a. A reason for the superior performance of UL12-type over T7E and T5E, regardless of the endonuclease used, might be its ability to interact with endogenous DNA repair factors. In human cells, UL12 is essential for HSV replication, which heavily depends on recombination (Schumacher et al., 2012). The physical interaction between UL12 and the MRN complex could also contribute to increased HDR activity in plants, since MRN is evolutionary conserved (Balasubramanian et al., 2010). The effect of the MRN recruiting domain (1-126 aa) from UL12 on HDR efficiency could be an indication of this interaction (fig. S6). Whether and how UL12 interacts with plant endogenous DNA repair pathways needs further investigation. In human cells, expression of UL12 or ICP8 alone leads to increased single strand annealing and HR activity (Schumacher et al., 2012; Valledor et al., 2018). By contrast, upon, co-expression of UL12::Cas9 with ICP8 we observed a decrease of HDR activity compared to UL12::Cas9 alone (fig. S10). Reasons for this difference are not known but could be due to the partial toxicity observed when ICP8 is expressed (fig. S11).

The improvement we observe in the frequencies of stable knock-ins in Arabidopsis by 5′-exonuclease fusions over Cas9 alone is substantial, around 10-fold, but not as strong as in the transient assay in N. benthamiana (20 to 40 fold). One reason for this could be the high copy number of T-DNAs that are delivered individually to plant cells in the transient assays, providing both higher levels of T-DNA donor and of Cas9 expression. Regardless of potential improvements, the 5’-exonuclease::Cas endonuclease fusions presented here make it possible to generate knock-ins or gene replacement events for which no selection is available routinely and will find use both for fundamental research as well as applications in crop plants.

## Materials and Methods

### Bacterial, plant growth conditions and Arabidopsis transformation

*Escherichia coli* strain DH10B and *Agrobacterium tumefaciens* strain GV3101::pMP90 (Koncz and Schell, 1986) were grown in lysogeny broth/medium (LB medium [Duchefa Biochemie]: 10 g/l tryptone, 10 g/l sodium chloride, 5 g/l yeast extract) with selective antibiotics at 37°C and 28°C, respectively. *Nicotiana benthamiana* plants were grown in a phytochamber (day and night temperatures of 21°C and 18°C, respectively) with 16-h light and 50% to 60% humidity. Arabidopsis plants were grown in the greenhouse, and in case of the *thic* T-DNA insertion lines, were supplied with thiamine by watering with a 300 mM thiamine solution. Arabidopsis plants were transformed using Agrobacterium GV3101::pMP90 strains by the floral dip method (Clough and Bent, 1998).

### Construct design and vectors

All constructs were generated via Golden-Gate cloning (GG-cloning) with the syntax of the modular cloning system (Engler et al., 2014; Weber et al., 2011). All sequences and cloning procedures can be found in the supporting information. Exonucleases were codon optimized for *Nicotiana benthamia* (Nb) and synthesized via Geneart Regensburg (Thermo) as level -1 modules. Corresponding Exonuclease level -1 modules were combined with different linker level -1 modules to obtain linker-fused exonuclease level 0 modules. Linker-fused exonuclease level 0 modules were combined with other level 0 modules (promoter, endonuclease (Cas9 or Cas12a) and terminator) to assemble transcriptional units (level 1). For SpCas9 and ttLbCas12a the intronized version have been used (Grutzner et al., 2021; Schindele et al., 2023). Cas9 single guide RNAs (sgRNAs) were designed according to Grützner et al. (Grutzner et al., 2021), whereas the flip extension sgRNA scaffold and the Arabidopsis U6-26 t67 terminator were used as template for PCR-based sgRNA amplification (Castel et al., 2019; Chen et al., 2013; Dang et al., 2015). Purified sgRNA-t67 PCR products were assembled with pU6 Level 0 modules to obtain a transcriptional unit for sgRNA expression. Cas12a crRNAs (directed repeat (DR)) followed by the 20 nt spacer sequence were amplified via primer extension and cloned into Level 0 cloning vector pAGT6272. Corresponding DR-Spacer Level 0 modules were combined with SlU6 promoter- and t67 terminator-modules, to generate level 1 transcriptional units. Proper Cas9 or Cas12a target sequences were identified with the CRISPOR online tool (Concordet and Haeussler, 2018).

### Agrobacterium mediated transient expression

Agrobacterium GV3101::pMP90 strains with corresponding T-DNA vectors were grown on LB agar plates containing rifampicin, gentamycin and the vector-specific antibiotics for one to two days at 28 °C. Grown cells were resuspended in Agrobacterium infiltration media (AIM; 10 mM MES; 10 mM MgCl_2_; 150 mM acetosyringone) with an optical density of OD_600_ = 0.2 and mixed at equal concentration if multiple Agrobacterium strain were combined. Agrobacterium suspensions were infiltrated in the abaxial leaf side of 5 to 6 week-old Nb plants using a needleless syringe.

### TMV assays

A T-DNA containing the modified TMV genome was transformed into Nb and resulting transformants were screened for HR events after attB-site specific DSBs triggered by Cas9 or TALENs in the presense of the corresponding repair donor. Nb primary transformant nbi775 showed highest activity and was used for all further TMV assays. Leaves of three individual plants were inoculated with mixtures of different Agrobacterium strains (Nuclease (Cas9 or Cas12a), sgRNA, Donor) at varying positions. All constructs of one set of comparison were inoculated into the same leaf. GFP signals were visible three dpi and pictures were taken under UV-light at three and five or six dpi. For quantification, identical leaf areas of individual spots from one picture were digitally cut out and were used for GFP spot count. GFP spot numbers were normalized for comparison within individual plants.

### Western Blot

Three days after transient expression of exonuclease-fused Cas9 derivatives, one leaf disc (diameter 0.9 cm) from corresponding inoculated spots of three individual plants were pooled and disrupted using the TissueLyser II (Qiagen) for 30 sec at 30 Hz. Plant material was resuspended in 50 µl 1xPBS (137 mM NaCl, 2.7 mM KCl, 10 mM Na_2_HPO_4_, 1.8 mM KH_2_PO_4_) + 50 µl 2x Laemmli and boiled (95°C) for 10 minutes. The resulting extract was cleared by centrifugation (maximum speed, 8 minutes) and the supernatant was separated on an 8% SDS-PAGE. Proteins were blotted on nitrocellulose membrane using the Power Blotter-Semi-dry Transfer System (Thermo Fisher Scientific Inc.) according to manufacturer’s instructions. Antibody detection of the proteins was performed using the iBind Automated Western System (Thermo Fisher Scientific Inc.) according to manufacturer’s instructions. Cas9-specific antibody (Santa Cruz, 7A9-3A3; 1:150 dilution) and anti-mouse antibody (Sante Cruz, m-IgGk BP-HRP; 1:1000 dilution) were used.

### Isolation of genomic Nb DNA

Genomic DNA (gDNA) was isolated from *Nicotiana benthamiana* leaves using the cetyltrimethylammonium bromide (CTAB) method as described below. Two to three days after Agrobacterium mediated transient expression, two leaf discs (0.9 cm diameter) were harvested and immediately transferred into liquid nitrogen. Material was disrupted using the TissueLyser II (Qiagen) for 30 sec at 30 Hz and dissolved in 300 μl CTAB buffer (100mM Tris-HCL pH8, 1.5 M NaCl, 20 mM EDTA pH8, 2% C-TAB, 0.5% Na_2_SO_3_, 1% PEG 6000). After incubation for 20-40 minutes at 65°C, 200 μl chloroform was added followed by centrifugation (10 min, 18000 g). The aqueous phase was mixed with 200 μl isopropanol, followed by centrifugation (10 min, 18000 g). The resulting pellet was washed with 1 ml 70% EtOH and centrifuged again (5 min, 18000 g). The final pellet was dried for 10 minutes at 65 °C and dissolved in 200-300 μl of water. Concentration was determined using a NanoDrop spectrophotometer.

### Qualitative and quantitative GUS assay

Qualitative GUS staining was performed as described in Schornack et al. (2005)(Schornack et al., 2005) with minor changes. Briefly, the incubation of harvested leaf discs with GUS staining solution (10 mM sodium phosphate, pH 7, 10 mM EDTA, 0.1% Triton X-100, 0.1% 5-bromo-4-chloro-3-indolyl-β-D-glucuronide, 1 mM potassium ferricyanide, and 1 mM potassium ferrocyanide) at 37°C was reduced to 6 hours. Qualitative GUS assay was performed as described by Kay et al. (2007)(Kay et al., 2007) with minor changes. Two or three days after Agrobacterium mediated transient expression in three individual *Nicotiana benthamiana* plants, two leaf discs (diameter 0.9 cm) were harvested and pooled per plant. Error bars indicate the standard deviation of three biological replicates (plants). GUS-assays were repeated two to three times with similar results.

### UL12 Protein expression and purification

The construct for recombinant expression of His-tagged UL12 (pAGT8232) was transformed into E.coli BL21 DE3 cells and selected on Lysogeny broth (LB) agar plates with 50 µg/ml Kanamycin. A single colony was picked and inoculated into 20 ml liquid LB-media with 50 µg/ml kanamycin followed by growth over night at 37 °C. The pre-culture was used to inoculate 400 mL LB (+Kan) culture. At a cell density of 0.8 OD600, UL12 expression was induced by 1mM isopropyl ß-D-1-thiogalactopyranoside (IPTG) for 22h at 20 °C. Cells were separated into 9x 50 ml falcons and harvested by centrifugation at 4000 g for 20 min at 4°C. Pellets were resuspended in 20 ml lysis buffer (20 mM Tris-HLC pH 7.9, 500 mM NaCl, 10% glycerol; +DNase, + Lysozyme + proteinase inhibitor) and lysed by three freeze and thawing cycles. The soluble fraction was separated from the debris by centrifugation at 4000g for 40 minutes at 4°C and the supernatant was purified using 1 ml Ni-NTA columns (Macherey-Nagel). His-tagged UL12 was eluted using lysis buffer with increasing concentration of Imidazole (100 mM, 200 mM and 300 mM). Fractions were analyzed for UL12 presence by western blotting using an anti-6xHis antibody and elution fractions from 200 and 300 mM imidazole were combined and the buffer exchange to storage buffer (20 mM Tris-HCL pH7.9, 500mM NaCl, 10% glycerol) using PD-10 desalting columns (GE Healthcare). Protein concentration was calculated using NanoDrop using the UL12 protein extinction coefficient. Recombinant UL12 was concentrated using 30 kDa Amicon filters (Merk) to a final concentration of 24.4 µg/ml. Proteins were aliquoted and stored at -80°C.

### Fluorophore assay

A hairpin oligonucleotide with internally labelled Oregon green 488 was synthesized by Eurofins Genomics (labeled nucleotide underlined; 5′-GGAAGGGCCCGCTGACAGTTTTTCTGTCAGCGGGCCCT**<u>T</u>**CC-3′;(Nikiforov, 2014)). Fluorescence assays were performed using the TECAN Spark plate reader and blank 96-well plates. Emission wavelength (Emission = 520 nm; Excitation = 495 nm (width 10 nm each); Z-position = 18173; Gain 50) was monitored over 30 minutes at 28 °C. Reaction was performed in 100µl 1xNEB4 buffer (50 mM potassium acetate, 20 mM Tris-acetate, 10 mM magnesium acetate, 1 mM DTT, pH 7.9) + 5 mM fresh DTT, 200 nM labelled oligonucleotide, 5U/100µl exonuclease enzyme (T5E (New England Biolabs M0663S) and T7E (New England Biolabs M0263S), □□exonucleases (T7E and T5E) and purified UL12 we applied UL12 with the same molarity as T7E.

## Data availability

Sequences off all constructs and used primers are given in the supplemental information.

## Acknowledgement

The work presented here was funded by grant number 031B0548 in the frame of the program “Crop plants of the future” from the *Bundesministerium für Bildung und Forschung* to AT. The TMV-GFP reporter N. benthamiana line was kindly provided by Icon Genetics. We thank Thomas Lahaye for critical reading of the manuscript and helpful suggestions. We also thank the IPB gardeners and staff for excellent support.

## Author contributions

Conceptualization: AT, TS

Methodology: TS, AP, PS, TI, RG

Investigation: AP, PS, TI, RG, AT, TS

Visualization: TS, AT

Funding acquisition: AT

Project administration: AT

Supervision: SM, TS, AT

Writing – original draft: TS

Writing – review & editing: AT, SM, TS

## Supplemental Information

Supplementary Text

Supplemental Figures S1-S18

Supporting Information

