## Supplemental Figures and Information for "Efficient scar-free knock-ins of several kilobases by engineered CRISPR/Cas endonucleases"

**Supplemental Information**

**This section includes:**

Figs. S1 to S18

Supplementary Text (Note S1-S2)

Supporting Information

**Supplemental Figure 1**

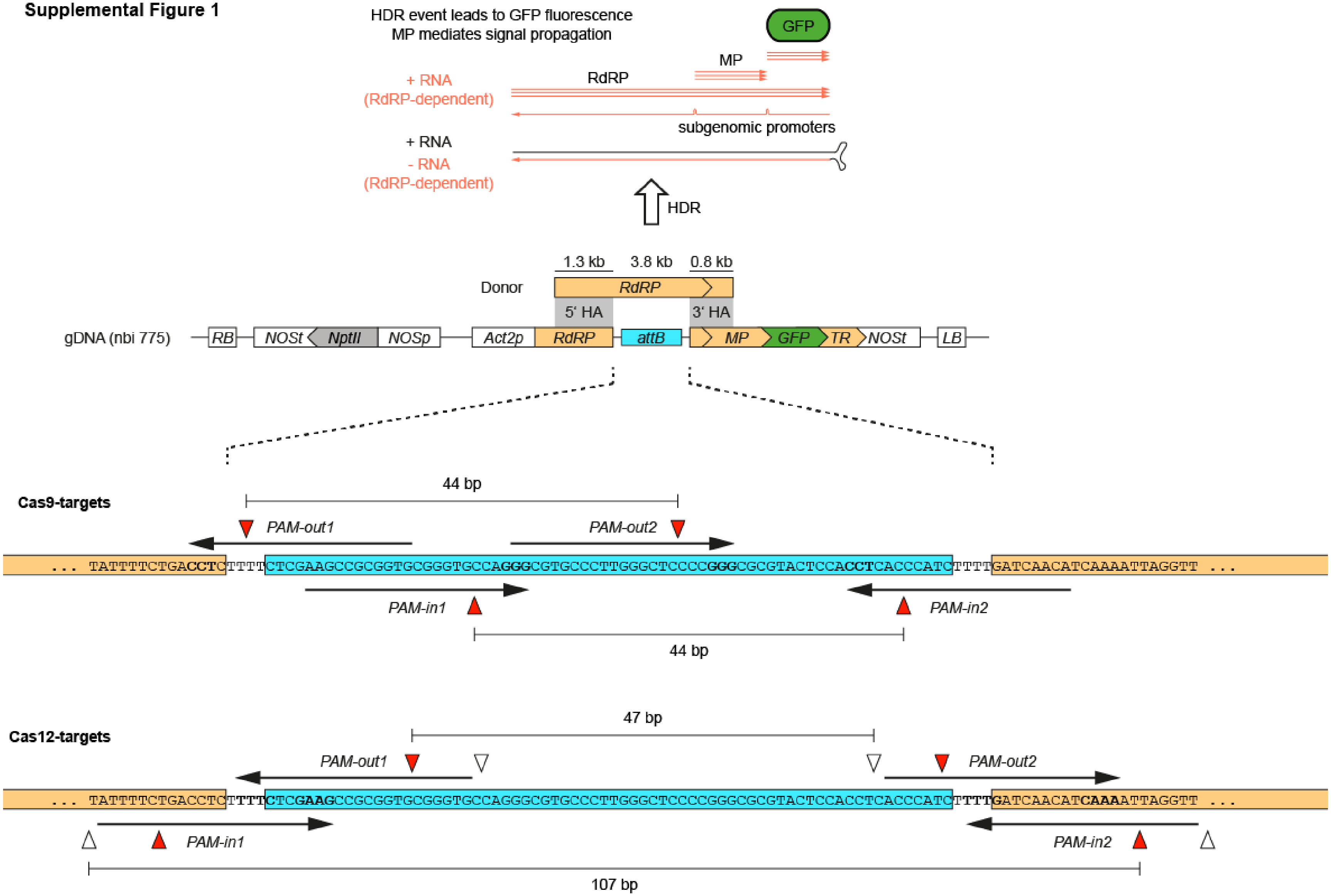


7

**Supplementary Fig. 1. A *Nicotiana benthamiana tobacco mosaic virus* (TMV) reporter line.** The TMV from the transgenic locus is unable to replicate, because a 3.8 kb fragment of the RNA-dependent RNA polymerase (RdRP) coding sequence has been deleted and replaced by an attB site. The attB site was initially placed in this construct to be used for recombinase mediated recombination, but is not used here for this purpose. Here the only role of this sequence is to serve as target site for cleavage by guide RNAs. The viral vector construct also contains GFP in place of the native TMV viral coat protein. A functional viral vector can be reconstituted from the inactive one by induction of DSBs in the attB, followed by homology-directed repair with a donor template delivered by agrobacterium. After repair, the viral vector is first transcribed from the ACT2 promoter, leading to production of a positive strand (+) RNA. This RNA is translated to produce RDRP, but also serve as a template for synthesis of a (-) RNA starting from the 3’ terminal region (TR) using the translated RDRP protein. The (-) RNA then serves as a template for synthesis of additional (+) RNA molecules, as well as for synthesis of sub-genomic RNAs transcribed from sub-genomic promoters.  The sub-genomic RNAs serve for translation of the movement protein (MP) and GFP. This explains why GFP cannot be expressed from the transgenic locus before repair, even though its entire coding sequence is present, as subgenomic RNAs are required for GFP expression. Expression of MP and GFP allows tracking of viral cell to cell movement (GFP signal) from single cell gene targeting events. The Cas9 and Cas12a target sites are given in the lower panel. For staggered DNA cleavage by Cas12a, nicking of the targeted and non-targeted strand is indicated by white and red rectangles, respectively.

**Supplemental Figure 2**

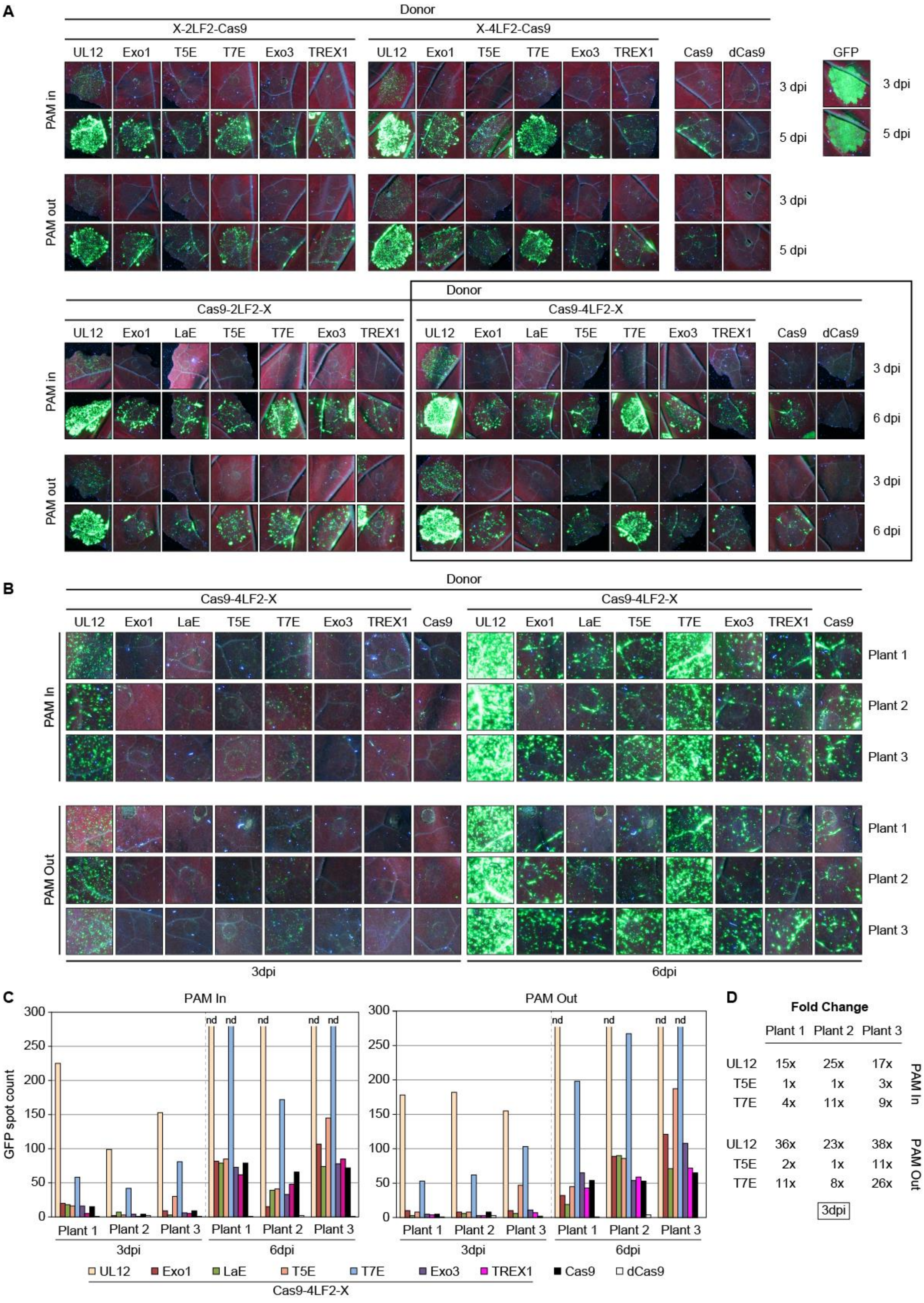

**Supplementary Fig. 2. Quantification of gene targeting events using the TMV reporter line.** **(A)** Different exonuclease-endonuclease combinations were inoculated on one leaf of three individual plants. Positions of the infiltrations for each construct were scrambled and differ between plants. Pictures of inoculated leaves taken three to five or six days post inoculation (dpi). Pictures of individual infiltrated areas are shown **(B)** Equal areas within the inoculated spots (from one picture) were sampled and used for GFP spot count. The areas used for GFP counting are shown. **(C and D)** Absolute numbers of GFP-spot count and fold change relative to Cas9 for three individual plants are given on the left and right panel, respectively.

**Supplemental Figure 3**

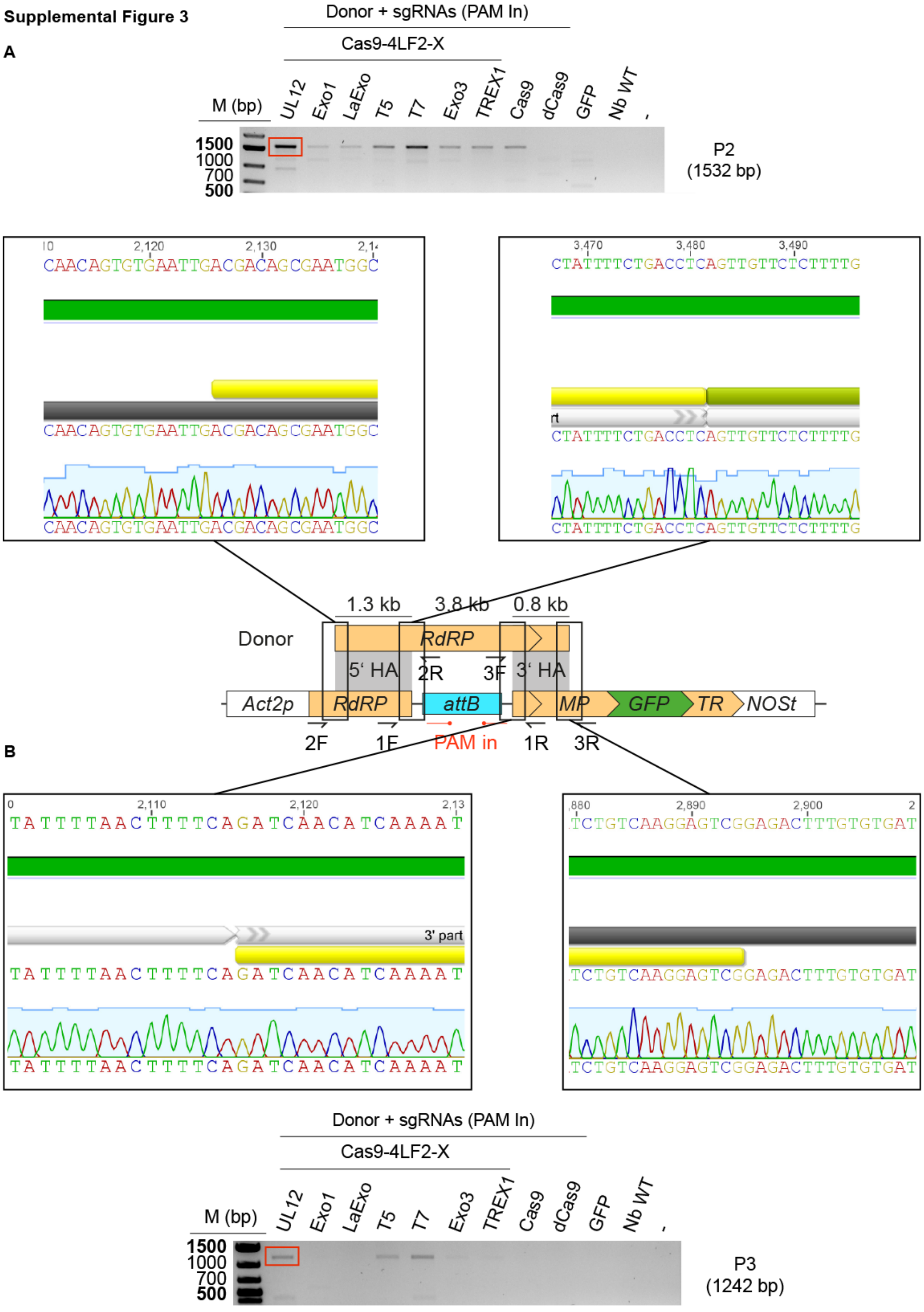


34

**Supplementary Fig. 3. Sequence confirmation of amplified 5´ and 3´ junctions from the TMV locus.** PCR fragments belong to Figure 1C. PCR fragments were subcloned and individual clones were sequenced. Borders of the 5´ **(A)** and 3´ **(B)** homology arms (HAs) are covered within the shown chromatograms.

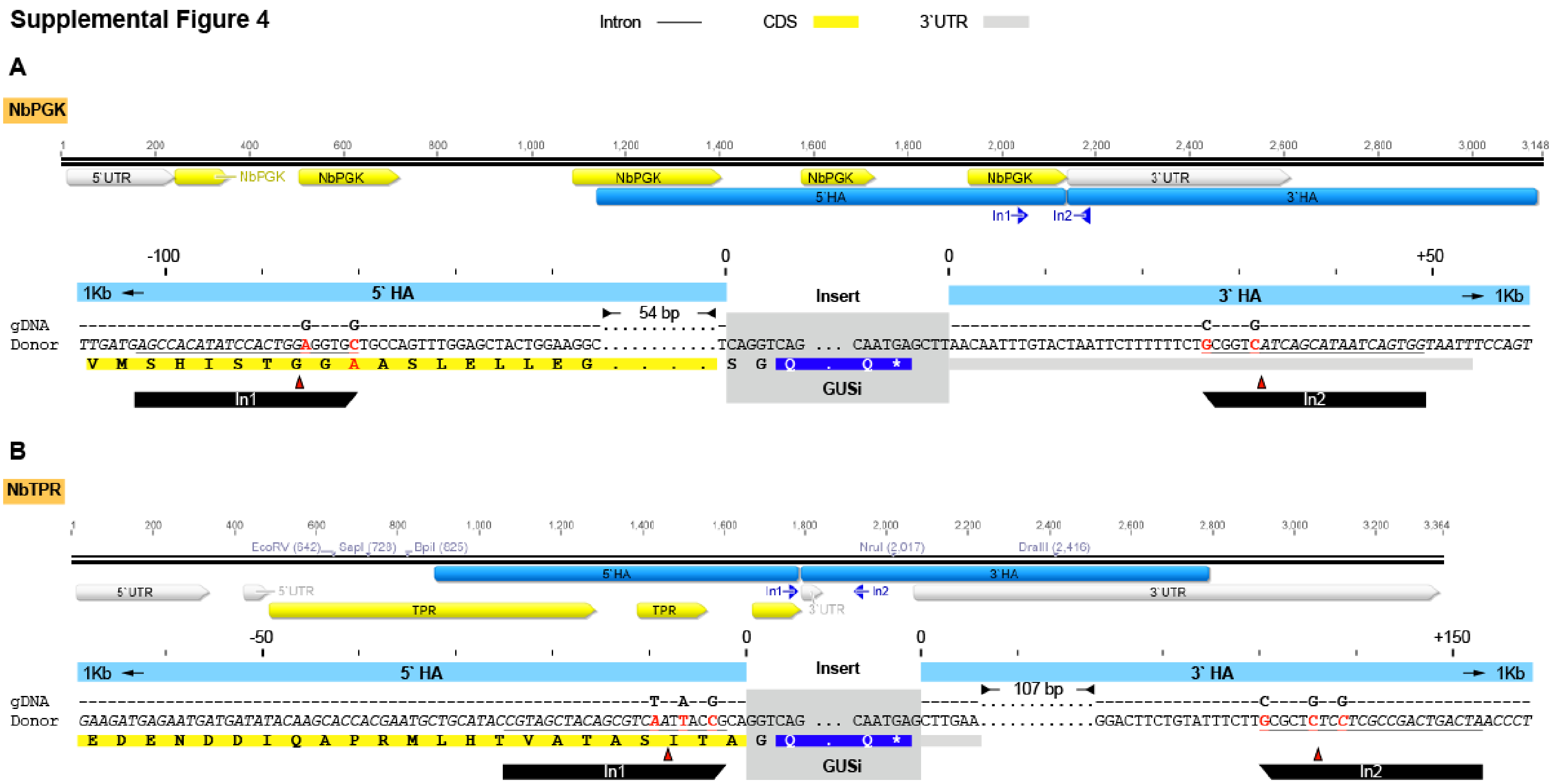


**Supplementary Fig. 4. Target design for in-frame GUS integration downstream of *Nicotiana benthamiana* endogenes *NbPGK* (A) and *NbTPR*. (B)** Exons and UTRs are given in yellow and grey, respectively. Homology arms (HA) of 1 kb each are indicated in blue. Target-specific sgRNAs were applied in a PAM-in orientation (dual sgRNAs; In1 and In2). Cas9 target sequences are underlined in the close-up view. Mutations in the donor to prevent cleavage after successful integration are indicated with red letters. Mutations are either at wobble positions to keep the amino acid sequence or based on C-C mismatches (lowest effect on the DNA double helix, pairing donor-gDNA) in UTR regions.

Supplemental Figure 5

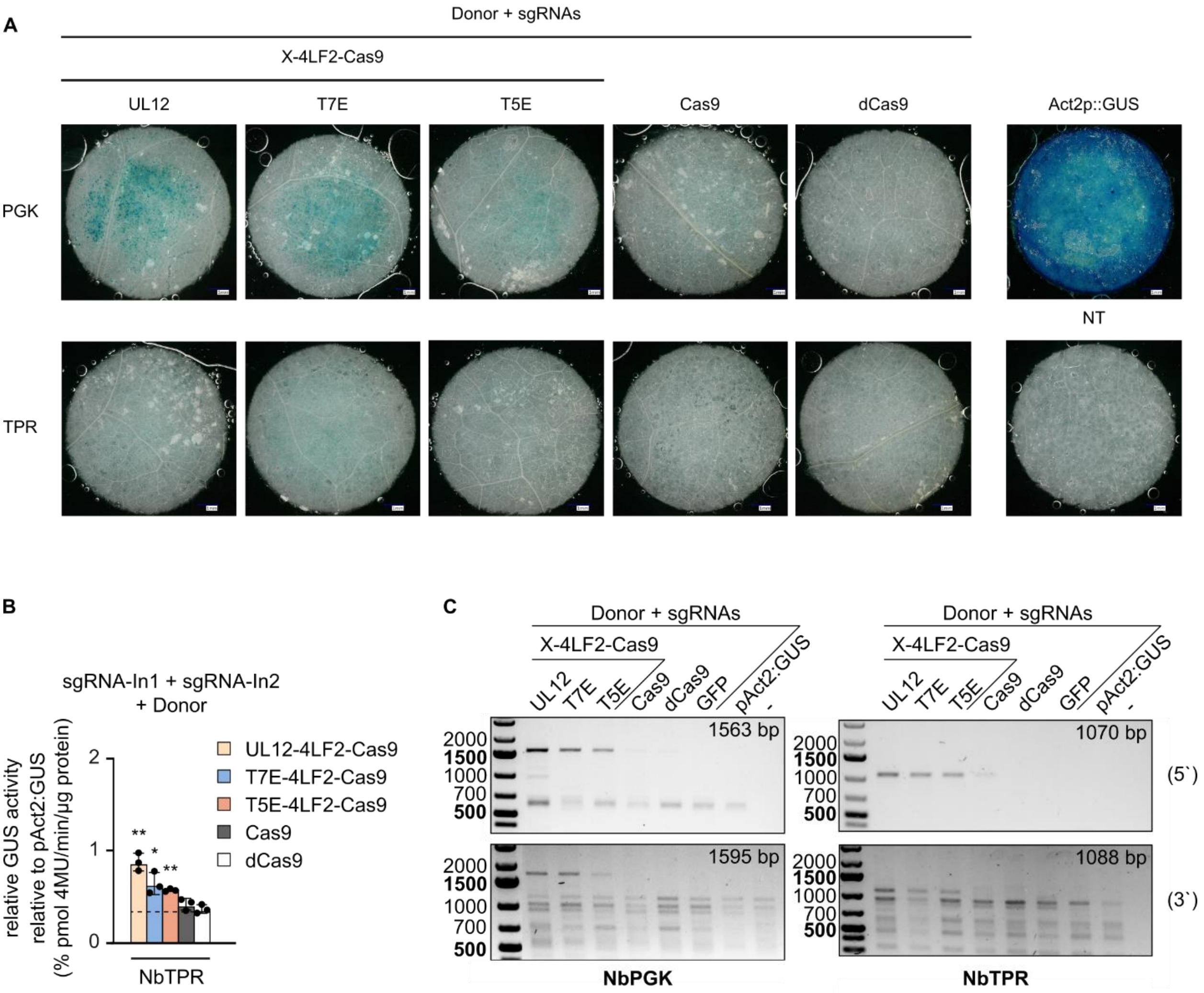


**Supplementary Fig. 5. GUS-staining of transformed leaves reveals increased gene targeting frequency for UL12- and T7E-fused Cas9 constructs on endogenous targets of N. benthamiana. (A)** GUS-stained leaf discs 3 dpi of targeting constructs for NbPGK and NbTPR. GUS positive spots indicates successful in frame integration of the GUS reporter. Actin2 promoter-driven GUS (Act2p::GUS) and non-treated leaf areas (NT) serve as positive and negative control, respectively. **(B)** Quantitative GUS assay from leaf extracts harvested 3 dpi from individual inoculated leaf areas. Values are relative to pAct2-driven GUS. Significance was evaluated with Student´s t-test; * p-value ≤ 0.05; ** p-value ≤ 0.01; *** p-value ≤ 0.001. **(C)** PCR-based genotyping of repair events. Amplification of upstream and downstream junctions are shown on the upper and lower panel respectively. Expected fragment sizes are given in the right top corner.

**Supplemental Figure 6**

**A**

2xLF2 = GS(GGGGS)$_6$

4xLF2 = GS(GGGGS)$_{12}$

T16 = GSGSETPGTSESATPES (16 aa of XTEN linker)

T40 = GSGSETPGTSESATPESGPGSEPATSGSETPGTSESATPES (40 aa of XTEN linker)

T144 = GSGTSTEPSEGSAPGTSESATPESGPGSEPATSGSETPGTSESATPESGPGSEPATSGSETPGTSESATPESGPGTSTEPSEGSAPGTSESATPESGPGSPAGSPTSTEEGSPAGSPTSTEEGSPAGSPTSTEEGTSESATPESGP (144 aa of XTEN linker)

**B**

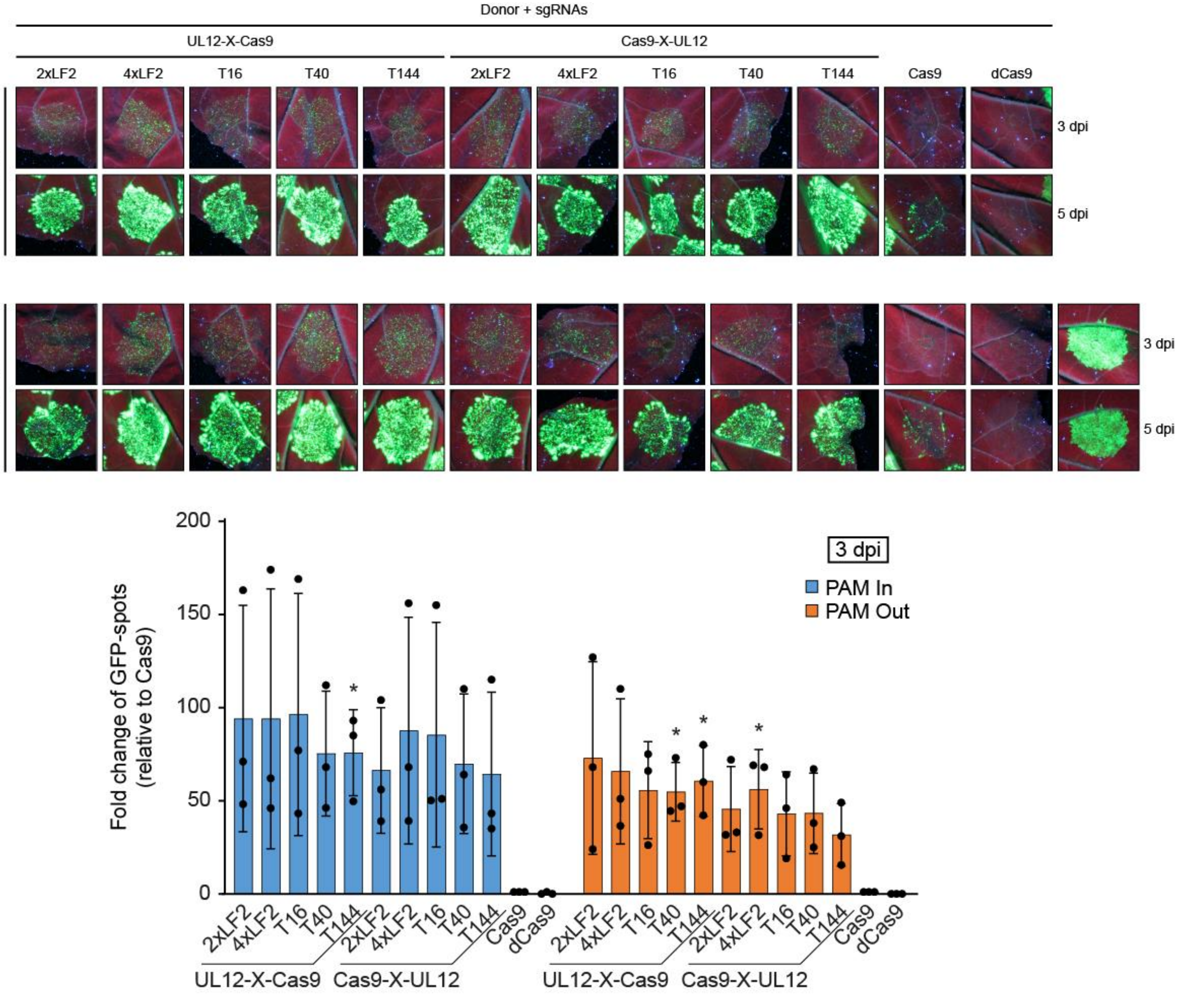


**C**

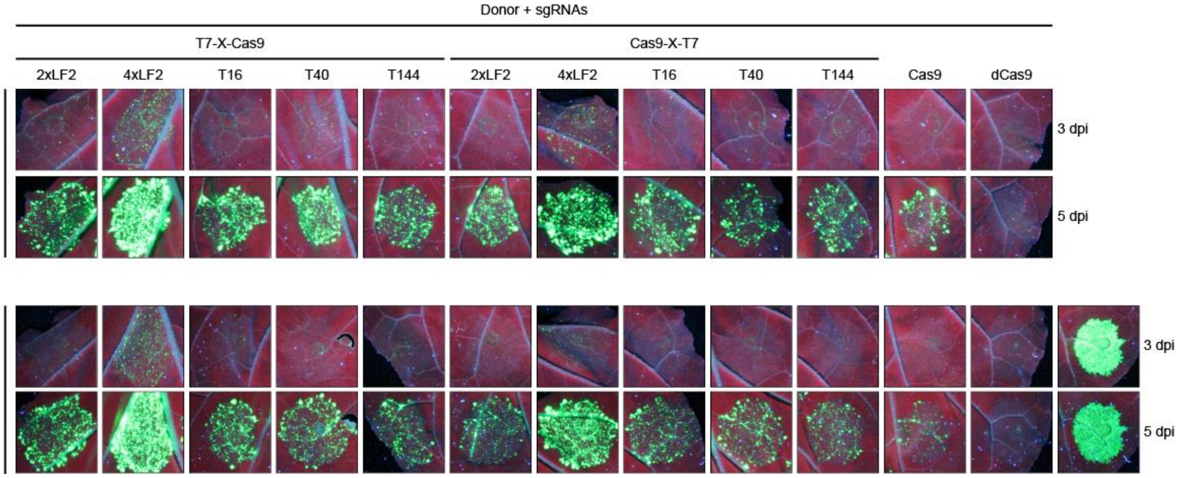

**Supplementary Fig. 6. Exonuclease Cas9 fusions work better with longer linkers. (A)** Amino acid sequence of linkers connecting UL12 or T7E to the N- or C-terminus of Cas9. **(B and C)** Analysis of gene targeting efficiency of different exonuclease Cas9 fusions using TMV reporter assay. **(B)** UL12 shows no linker preference. Relative number of GFP spots normalized to Cas9. Significance relative to Cas9 was evaluated with Student´s t-test; * p-value ≤ 0.05; ** p-value ≤ 0.01; *** p-value ≤ 0.001. , **(C)** T7E Cas9 fusion proteins showed a preference for the linker 4xLF2

#### Supplemental Figure 7

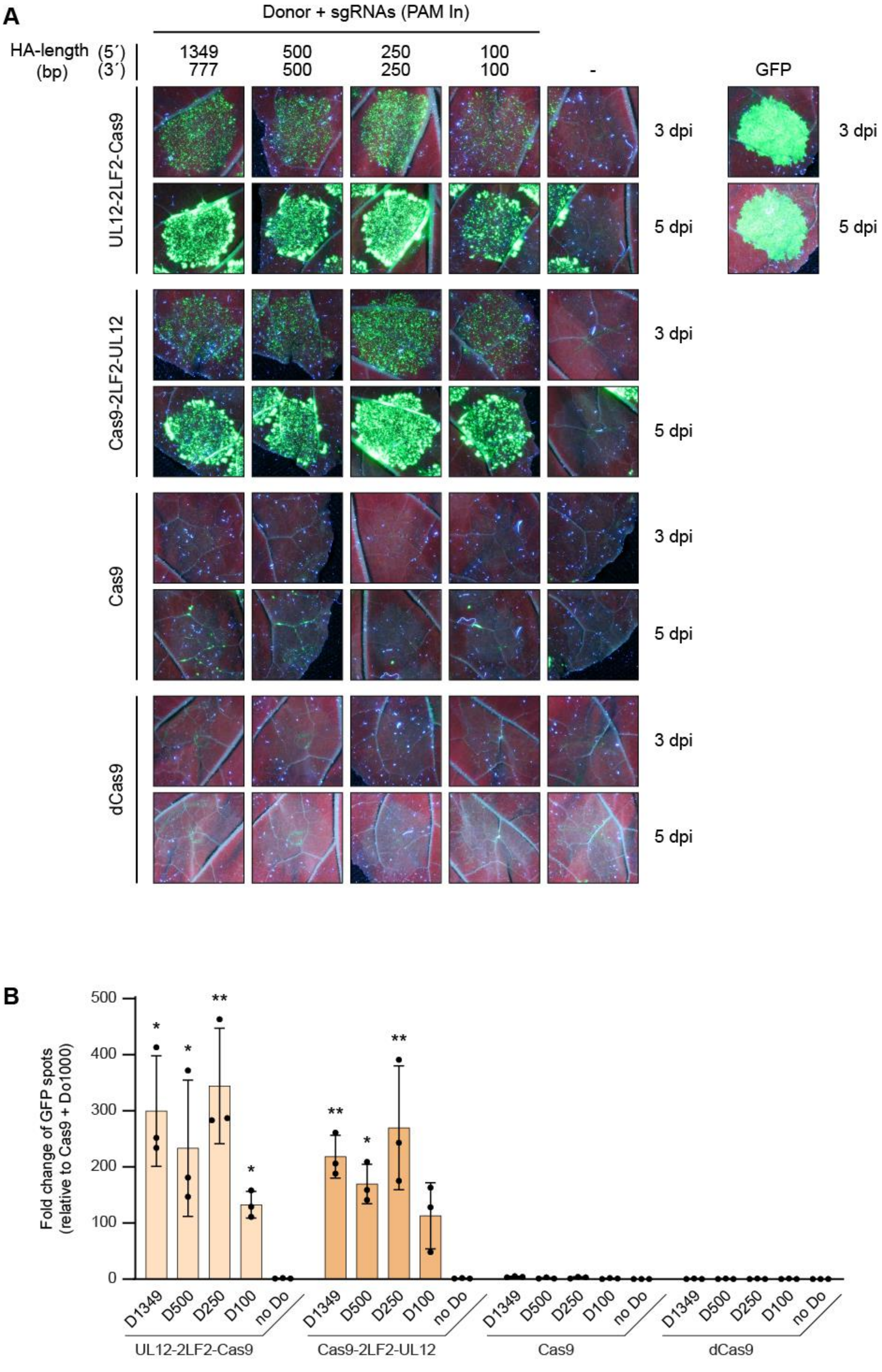


68

**Supplementary Fig. 7. Homology arms can be reduced to 250 bp without affecting gene targeting efficiency.** Several donors differing in the length of the 5´ and 3´ homology arms (HA) were tested using the TMV reporter assay (A and B). A slight increase of gene targeting efficiency for donor constructs possessing 250 bp was observed. HA length of 100 bp reduces gene targeting efficiency. (B) Fold change of GFP spots relative to Cas9. Significance relative to Cas9 was evaluated with Student´s t-test; * p-value ≤ 0.05; ** p-value ≤ 0.01; *** p-value ≤ 0.001.

#### Supplemental Figure 8

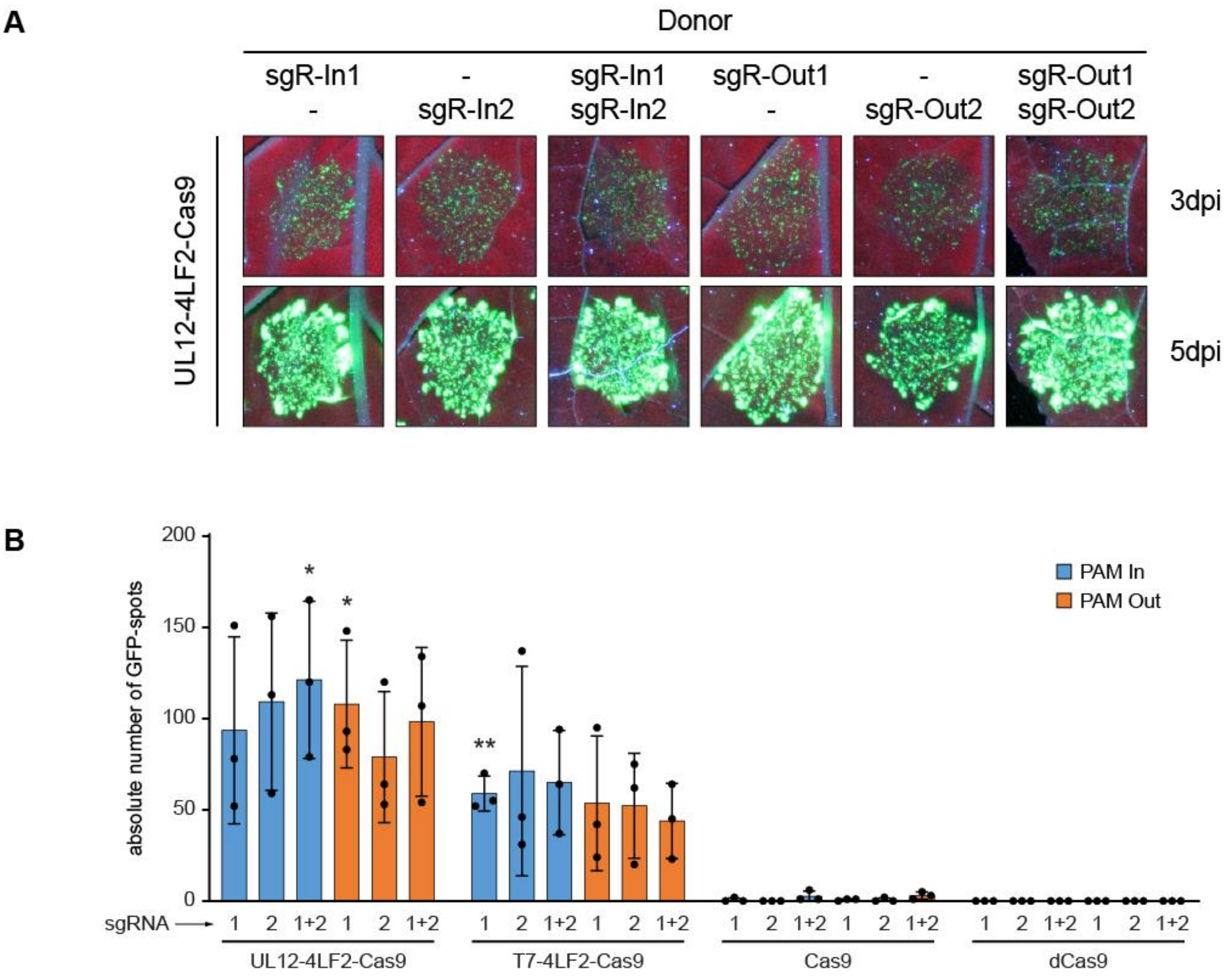


**Supplementary Fig. 8. Exonuclease fused Cas9 can be applied with a single sgRNA. (A)** Gene targeting efficiency of UL12-4LF2-Cas9 with all possible sgRNA combinations using the TMV reporter. **(B)** Graph shows absolute number of GFP spots, because no GFP spot could be observed with Cas9 in some cases. Significance relative to Cas9 was evaluated with Student´s t-test; * p-value ≤ 0.05; ** p-value ≤ 0.01; *** p-value ≤ 0.001. Individual sgRNAs can be used without negatively impacting gene targeting efficiency.

**Supplemental Figure 9**

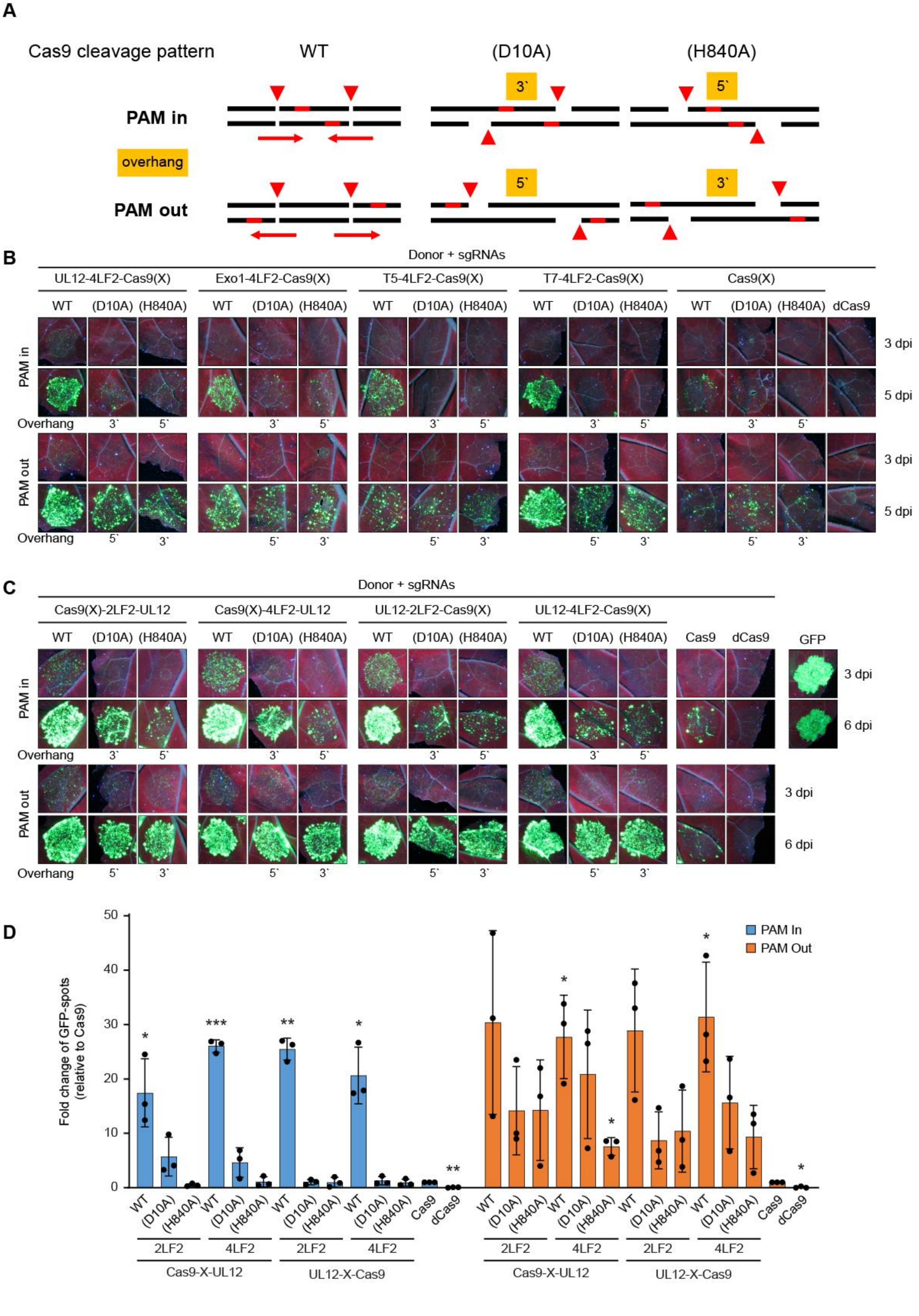

**Supplementary Fig. 9. Exonuclease fused Cas9 can be applied as dual nickase. (A)** Schematic overview of Cas9 and nCas9 cleavage patterns in dependency of dual sgRNA orientation (PAM in or PAM out). PAM is indicated as red line. DNA cleavage is indicated with a red triangle. **(B)** Gene targeting efficiency of different exonuclease-Cas9 nickase fusion proteins using two sgRNAs. SgRNAs in the PAM out configuration led to higher gene targeting efficiency. Combination of exonucleases with Cas9 (WT) led to the highest gene targeting efficiency in all cases. **(C)** Similar results were obtained with different UL12-fused Cas9 derivatives. Fusion of UL12 to Cas9(D10A) nickase with linker 4LF2 showed slightly increased activity in the PAM-out orientation. C-terminal fusion of UL12 to Cas9(D10A) by any linker is preferred in the PAM-in orientation. **(D)** Fold change of GFP spots relative to Cas9 counted from C. Relative number of GFP spots normalized to Cas9 with the same sgRNA setup. Significance relative to Cas9 was evaluated with Student´s t-test; * p-value $\leq 0.05$; ** p-value $\leq 0.01$; *** p-value $\leq 0.001$.

**Supplemental Figure 10**

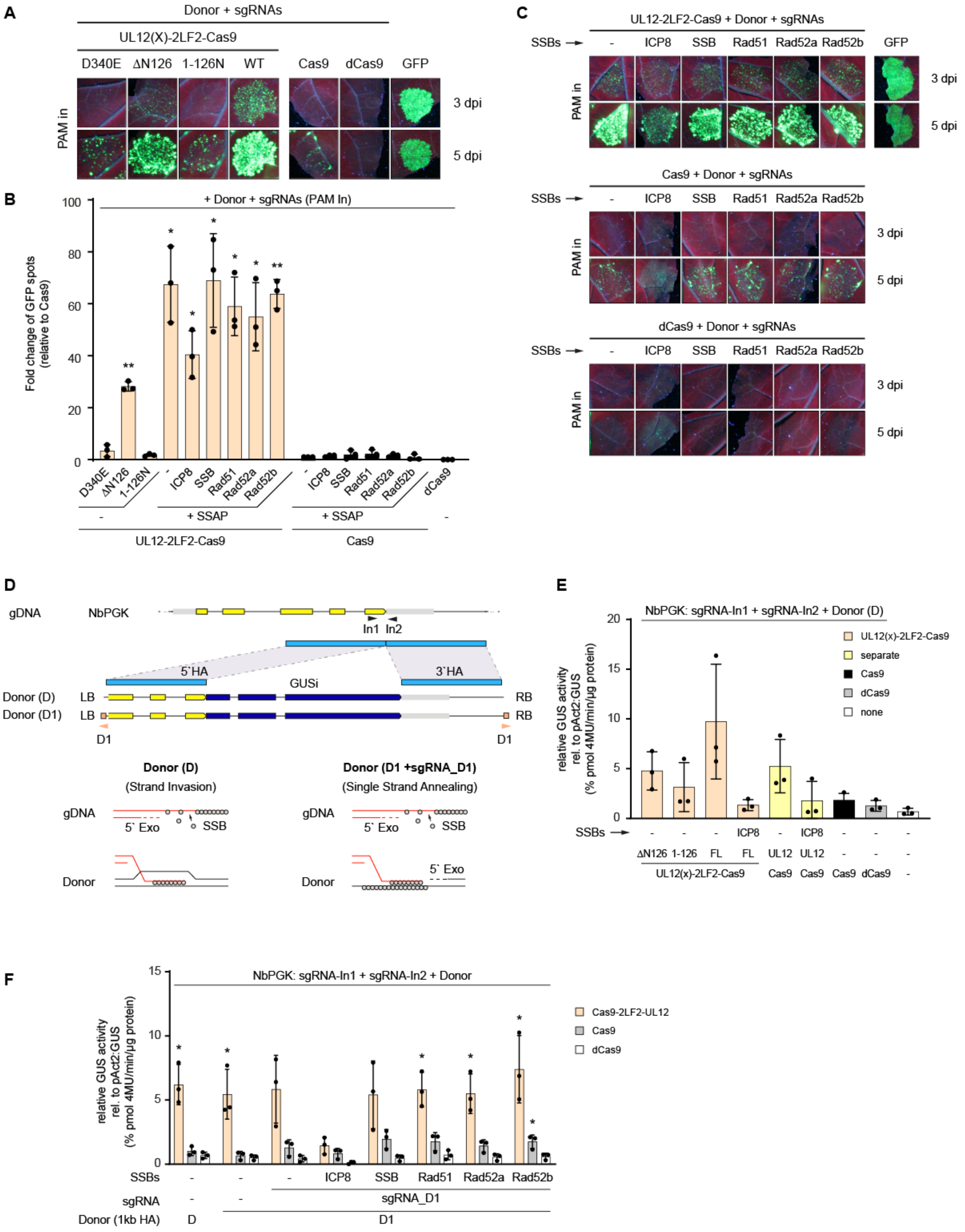

**Supplementary Fig. 10. Combination of active UL12-Cas9 with single strand annealing proteins (SSAPs). (A and B)** Effect of UL12 mutations on gene targeting frequency using the TMV reporter line. Exonuclease-dead UL12 (D340E) lost the ability to increase gene targeting frequency similar to the solei MRN-recruiting domain (1-126N) if fused to Cas9. Deletion of the MRN-recruiting domain (ΔN126) showed reduced gene targeting frequency. Fold change of GFP spots is given in B. Relative number of GFP spots normalized to Cas9 without SSAP. Significance relative to Cas9 was evaluated with Student´s t-test; * p-value ≤ 0.05; ** p-value ≤ 0.01; *** p-value ≤ 0.001. **(B and C)** Co expression of ICP8 negatively effectsgene targeting frequency. Similar results were obtained with in-frame GUS integration at the Nicotiana benthamiana locus NbPGK (D to F). **(D)** Rad51 and Rad52 possess strand invasion and single strand annealing (SSA) activity, respectively. To allow SSA we additionally cleave the donor at flanking D1 target sites. UL12 Exonuclease resects the 5´ ends leaving patches of ssDNA (3´ overhang), allowing association of Rad52 and SSA of the target gDNA and donor DNA. No significant changes on gene targeting frequency could be observed using the in-frame GUS integration downstream of NbPGK. **(E)** Analysis of gene targeting frequencies of Cas9-fused UL12 deletion derivatives and comparison between free and Cas9-fused UL12 using the in-frame GUS integration at the NbPGK locus. Direct fusion of UL12 to Cas9 leads to higher gene targeting frequency compared to separate expression of UL12 and Cas9. **(F)** Analysis of the effect of co-expression of SSAPs on gene targeting frequencies using the in-frame GUS integration downstream of NbPGK. Similar to the TMV assay a negative impact of ICP8 co expression on gene targeting frequencies could be observed.

#### Supplemental Figure 11

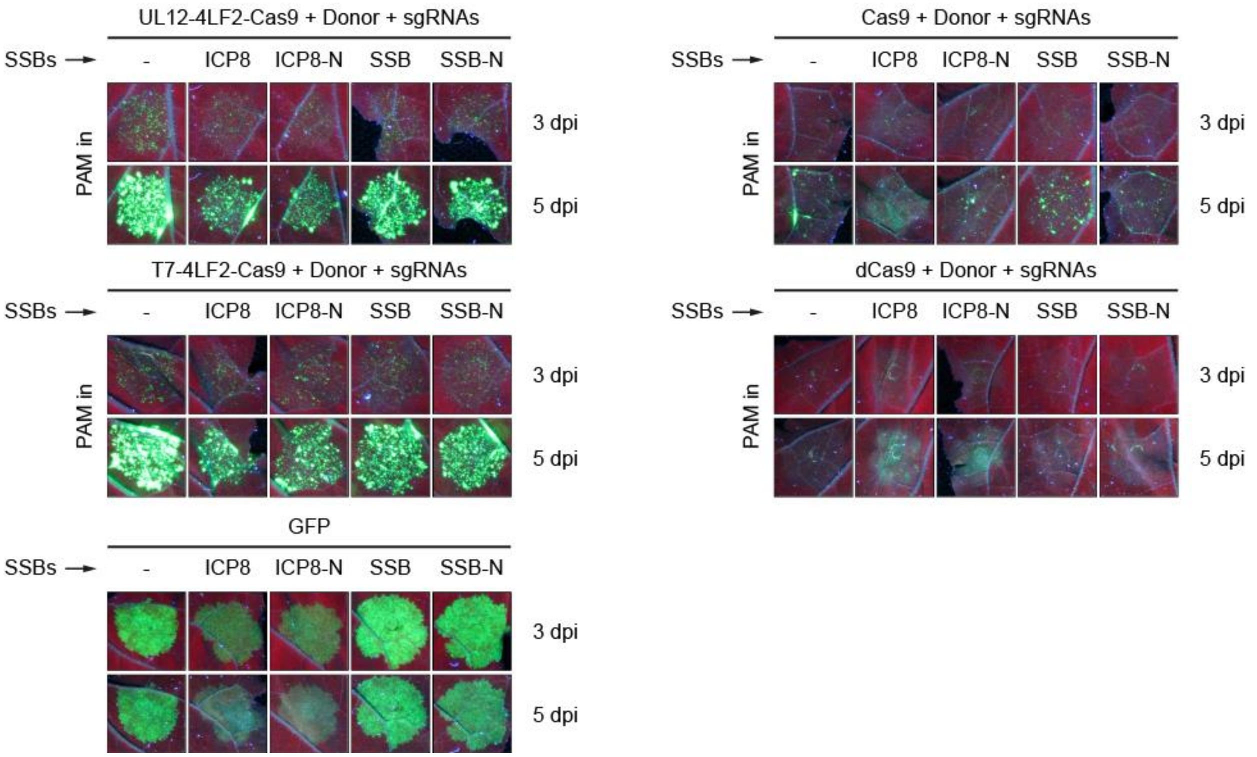


**Supplementary Fig. 11. Expression of ICP8 leads to cytotoxic effects. (A)** Transient expression of HSV ICP8 in *Nicotiana benthamiana* leaves triggers cytotoxic effects five days post inoculation. TMV reporter assay with different exonuclease-Cas9 combinations. Co-expression of ICP8 or ICP8 with NLS (ICP8-N) reduces expression of GFP and triggers cell death at later time points. E.coli SSB and SSB-N serve as control.

**Supplemental Figure 12**

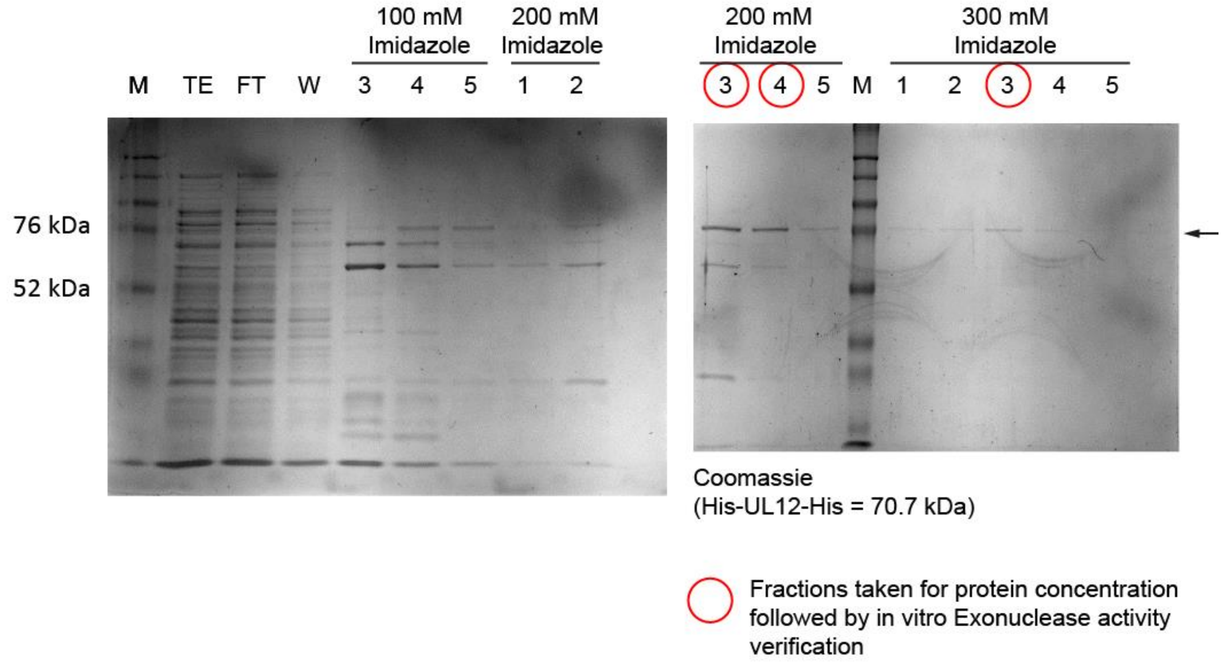


**Supplementary Fig. 12. Purification of recombinant his-tagged UL12 from E.coli.** Coomassie stained gel loaded with different fractions from the purification procedure. Elution fractions 3 and 4 (200 mM Imidazole) and 3 (300 mM Imidazole) were collected and combined for protein concentration. Molecular weight of His-UL12-His is 70.7 kDa.

**Supplemental Figure 13**

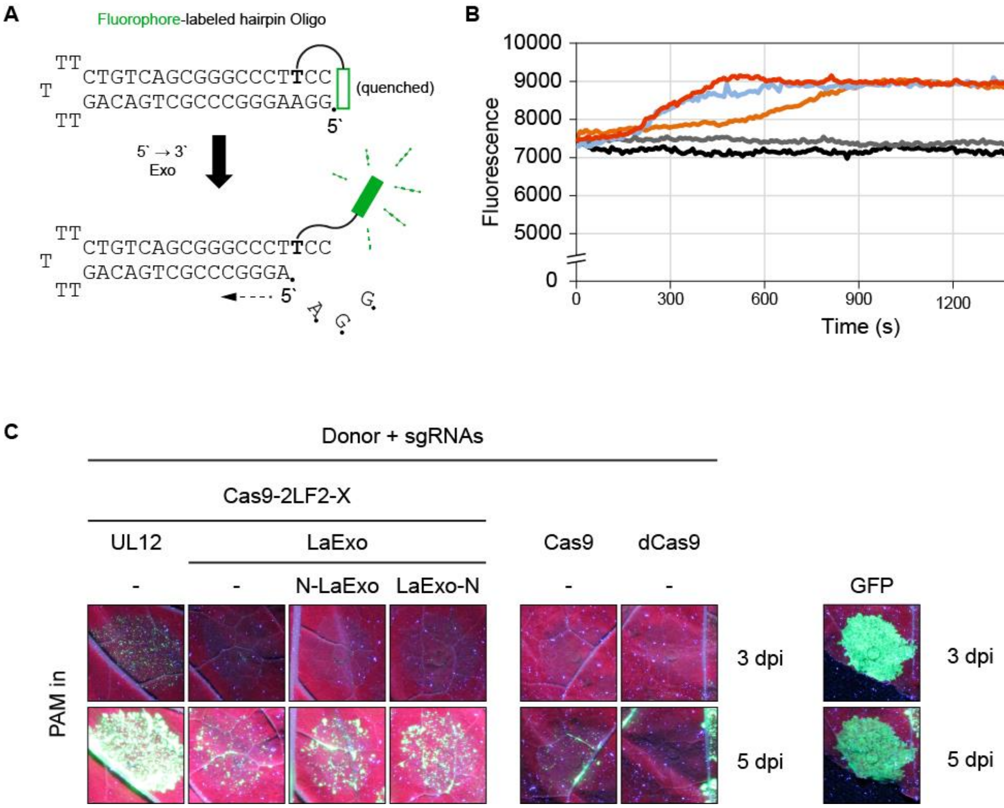


**Supplementary Fig. 13. Cas9-combined Lambda exonuclease does not increase gene targeting efficiency.** **(A)** Principle of the assay based on a hairpin oligonucleotide that carries a fluorophore, which is quenched by the terminal C:G base pair. Upon 5'-exonuclease activity the quenching is released, triggering fluorescence emission. **(B)** Fluorescence intensity plotted over time (30 minutes) after recombinant exonucleases were added to the reaction. LaExo resects blunt end substrates as efficiently as T7E. T5E possesses reduced affinity on blunt end DNA substrates. **(C)** Co-expression of WT or NLS-fused LaExo does not significantly increase gene targeting efficiency of LaExo-fused Cas9. UL12-fused Cas9 outperforms LaExo-fused Cas9 constructs.

#### Supplemental Figure 14

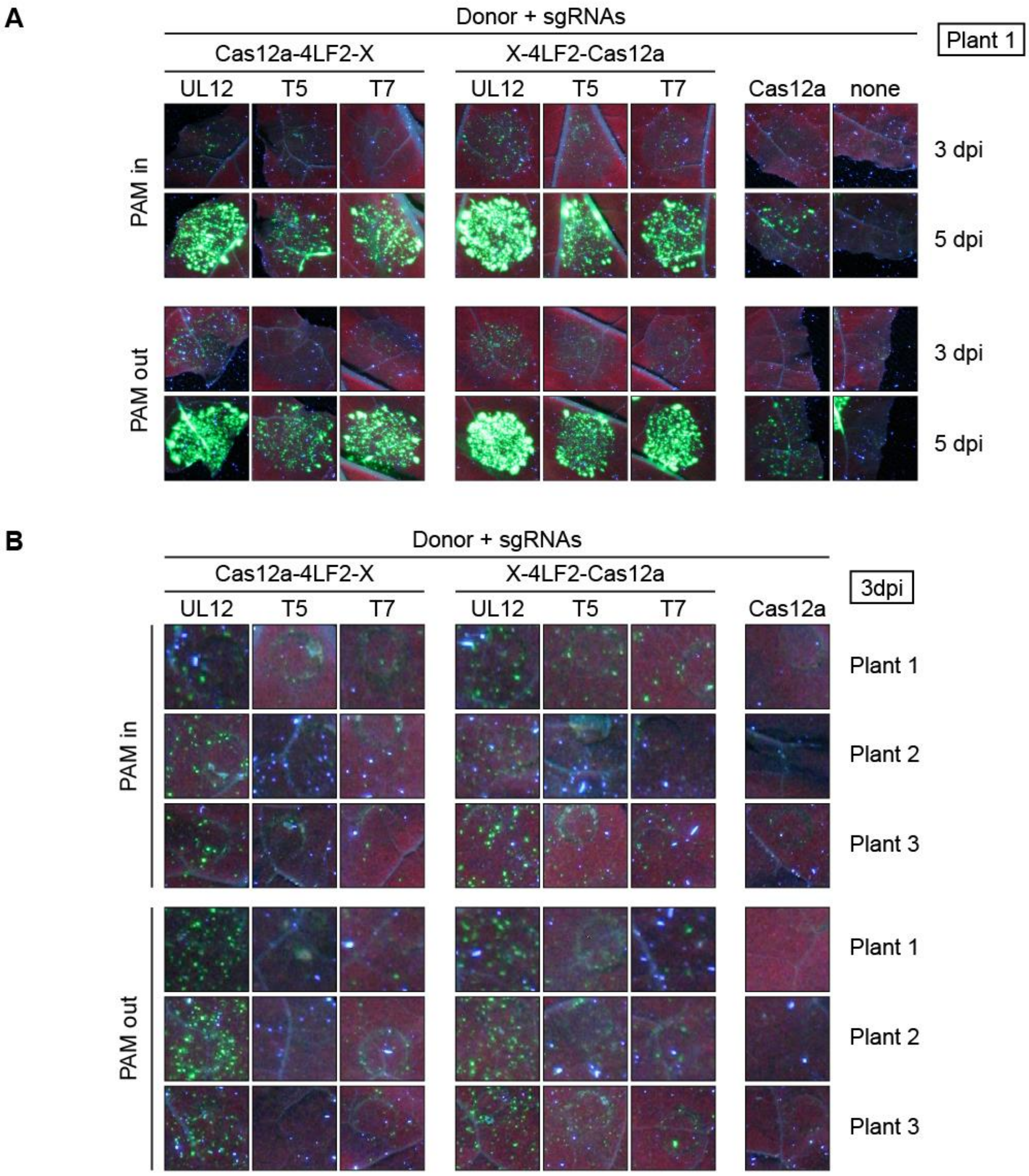


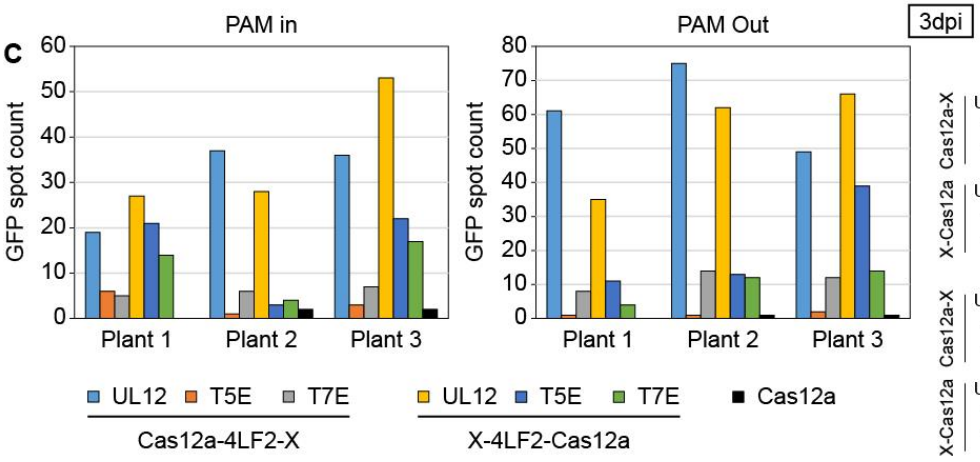


**Fold Change to Cas12a**

| | | Plant 1 | Plant 2 | Plant 3 | |
|---|---|---|---|---|---|
| Cas12a-X | UL12 | - | 19x | 18x | PAM In |
| | T5E | - | 1x | 2x | |
| | T7E | - | 3x | 4x | |
| X-Cas12a | UL12 | - | 14x | 27x | |
| | T5E | - | 2x | 11x | |
| | T7E | - | 2x | 9x | |
| Cas12a-X | UL12 | - | 75x | 49x | PAM Out |
| | T5E | - | 1x | 2x | |
| | T7E | - | 14x | 12x | |
| X-Cas12a | UL12 | - | 62x | 66x | |
| | T5E | - | 13x | 39x | |
| | T7E | - | 12x | 14x | |

**Supplementary Fig. 14. Fusion of UL12 to Cas12a increases gene targeting efficiency. (A)** Analysis of gene targeting efficiency after transient expression of indicated Cas12a variants in the transgenic Nb TMV reporter line (Plant1). **(B)** Areas of the same size from the infiltrated leaf discs taken for GFP spot count. **(C)** Summary of absolute GFP spot counts for individual constructs. Fusion of UL12 outperforms T5E and T7E. For exonucleases fused to the N-terminus of Cas12a, T5E showed increased gene targeting efficiency over T7E.

**Supplemental Figure 15**

**A**

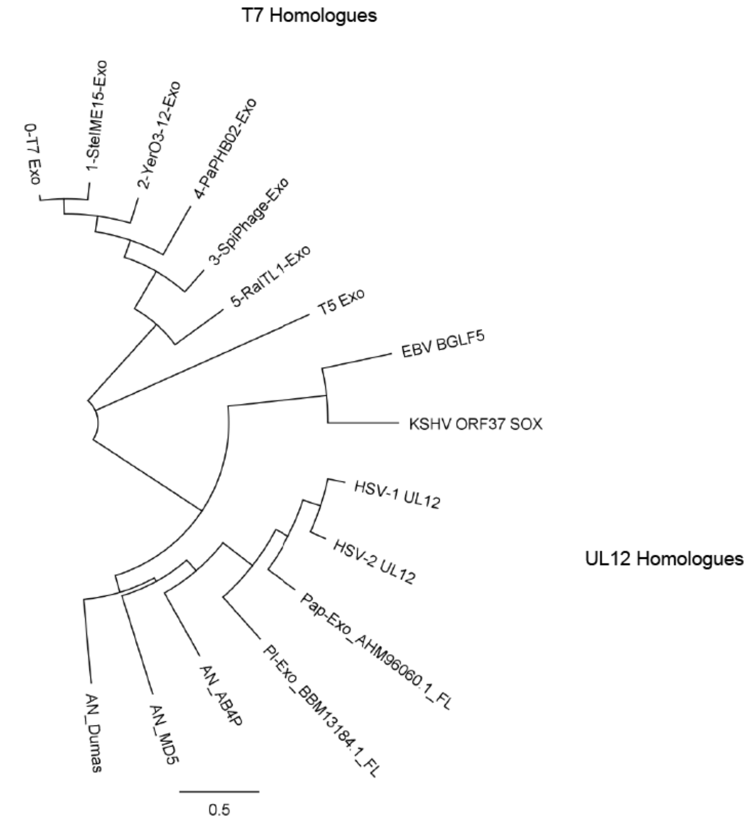


**B**

| Homologues | | HSV-1 UL12 | HSV-2 UL12 | Pap-Exo_AH... | PI-Exo_BBM1... | AN_AB4P | AN_MD5 | AN_Dumas | EBV BGLF5 | KSHV ORF37... | 0-T7 Exo | 1-SteIME15-... | 2-YerO3-12-... | 3-SpiPhage-... | 4-PaPHB02-... | 5-RaITL1-Exo | T5 Exo |
|---|---|---|---|---|---|---|---|---|---|---|---|---|---|---|---|---|---|
| UL12 | HSV-1 UL12 | | 79.9% | 61.2% | 50.6% | 36.0% | 27.2% | 26.1% | 16.4% | 16.3% | 7.0% | 6.7% | 7.0% | 8.0% | 5.9% | 7.7% | 5.8% |
| | HSV-2 UL12 | 79.9% | | 62.3% | 49.9% | 36.5% | 26.4% | 26.1% | 15.9% | 15.5% | 7.0% | 6.7% | 7.0% | 7.8% | 6.2% | 8.2% | 5.8% |
| | Pap-Exo_AHM96060.1_FL | 61.2% | 62.3% | | 46.4% | 36.6% | 27.8% | 29.2% | 16.6% | 16.5% | 7.1% | 7.0% | 7.0% | 8.1% | 6.5% | 8.6% | 5.3% |
| | PI-Exo_BBM13184.1_FL | 50.6% | 49.9% | 46.4% | | 31.4% | 25.0% | 26.6% | 16.3% | 14.0% | 6.9% | 6.6% | 7.4% | 7.2% | 6.1% | 8.6% | 5.7% |
| | AN_AB4P | 36.0% | 36.5% | 36.6% | 31.4% | | 30.4% | 32.9% | 16.8% | 16.1% | 6.3% | 5.7% | 6.2% | 7.2% | 6.4% | 7.0% | 6.5% |
| | AN_MD5 | 27.2% | 26.4% | 27.8% | 25.0% | 30.4% | | 28.7% | 17.0% | 15.5% | 7.9% | 7.3% | 7.8% | 7.6% | 6.8% | 6.9% | 6.1% |
| | AN_Dumas | 26.1% | 26.1% | 29.2% | 26.6% | 32.9% | 28.7% | | 17.9% | 16.7% | 7.3% | 7.0% | 6.6% | 7.8% | 7.0% | 8.2% | 6.6% |
| | EBV BGLF5 | 16.4% | 15.9% | 16.6% | 16.3% | 16.8% | 17.0% | 17.9% | | 41.7% | 5.5% | 4.9% | 6.3% | 7.4% | 5.7% | 6.6% | 4.7% |
| | KSHV ORF37 SOX | 16.3% | 15.5% | 16.5% | 14.0% | 16.1% | 15.5% | 16.7% | 41.7% | | 6.3% | 7.0% | 5.5% | 6.9% | 5.5% | 6.8% | 7.7% |
| T7 | 0-T7 Exo | 7.0% | 7.0% | 7.1% | 6.9% | 6.3% | 7.9% | 7.3% | 5.5% | 6.3% | | 87.3% | 72.8% | 65.4% | 54.5% | 40.7% | 9.1% |
| | 1-SteIME15-Exo | 6.7% | 6.7% | 7.0% | 6.6% | 5.7% | 7.3% | 7.0% | 4.9% | 7.0% | 87.3% | | 71.3% | 66.1% | 54.5% | 38.8% | 8.4% |
| | 2-YerO3-12-Exo | 7.0% | 7.0% | 7.0% | 7.4% | 6.2% | 7.8% | 6.6% | 6.3% | 5.5% | 72.8% | 71.3% | | 56.3% | 54.0% | 38.3% | 8.7% |
| | 3-SpiPhage-Exo | 8.0% | 7.8% | 8.1% | 7.2% | 7.2% | 7.6% | 7.8% | 7.4% | 6.9% | 65.4% | 66.1% | 56.3% | | 49.7% | 39.4% | 8.8% |
| | 4-PaPHB02-Exo | 5.9% | 6.2% | 6.5% | 6.1% | 6.4% | 6.8% | 7.0% | 5.7% | 5.5% | 54.5% | 54.5% | 54.0% | 49.7% | | 37.2% | 10.7% |
| | 5-RaITL1-Exo | 7.7% | 8.2% | 8.6% | 8.6% | 7.0% | 6.9% | 8.2% | 6.6% | 6.8% | 40.7% | 38.8% | 38.3% | 39.4% | 37.2% | | 10.7% |
| | T5 Exo | 5.8% | 5.8% | 5.3% | 5.7% | 6.5% | 6.1% | 6.6% | 4.7% | 7.7% | 9.1% | 8.4% | 8.7% | 8.8% | 10.7% | 10.7% | |

**Supplementary Fig. 15. Sequence analysis of T7E- and UL12-homologues. (A)** Phylogenetic tree showing the sequence-based distances between T7E, UL12 and their tested homologues. **(B)** Table of amino acid sequence identity between individual exonucleases.

#### Supplemental Figure 16

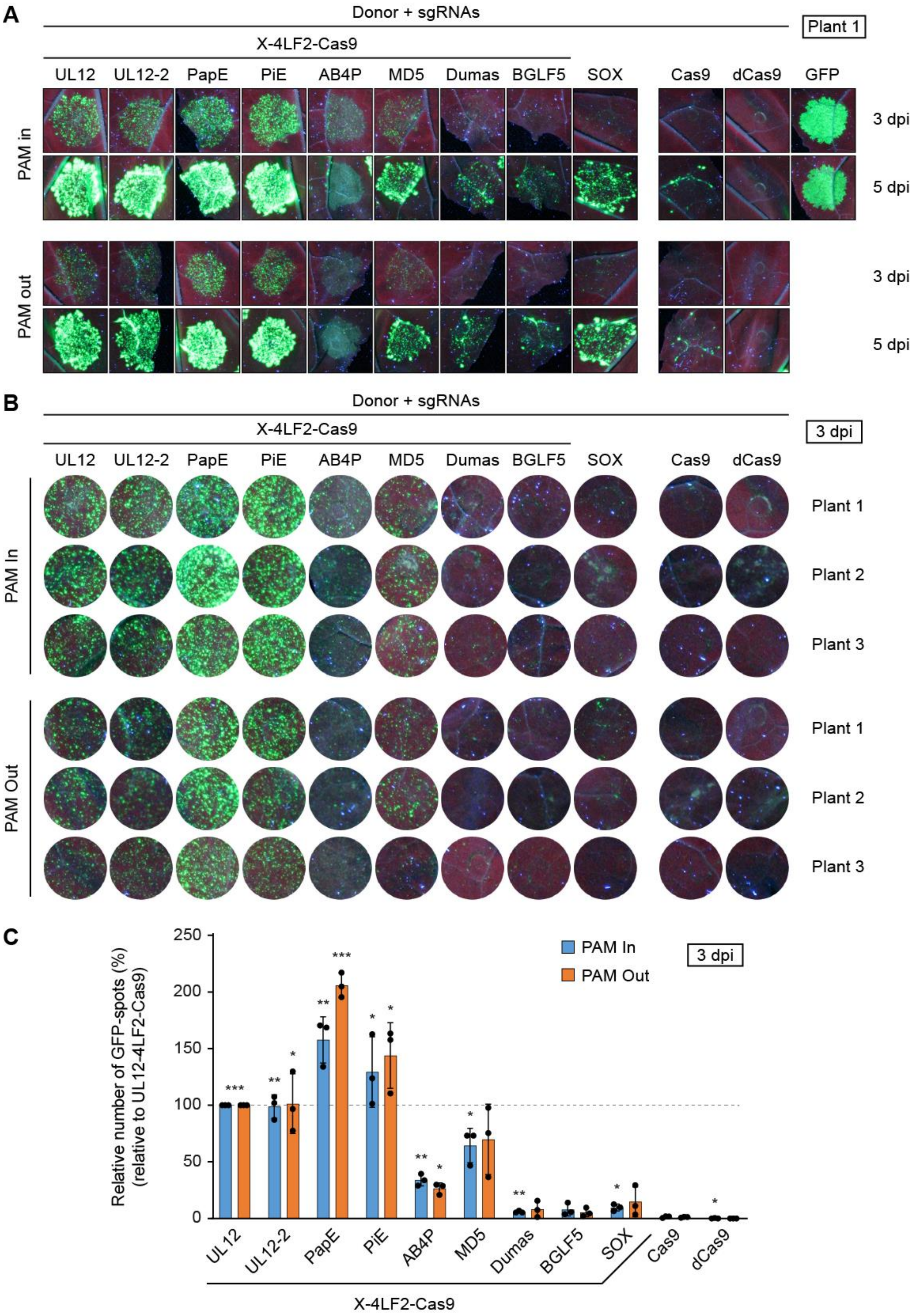


163

**Supplementary Fig. 16. Comparative analysis of Cas9-fused UL12 homologues. (A)** Analysis of gene targeting efficiency after transient expression of indicated Cas9 variants in the transgenic Nb TMV reporter line (Plant 1). **(B)** Areas of the same size from the infiltrated leaf discs taken for GFP spot count. **(C)** Summary of relative GFP spot counts for individual constructs. Number of GFP-spots are relative to UL12-fused Cas9 for which the GFP-spot number was set to 100%. Among the tested candidates PapE led to the highest number of GFP spots.

#### Supplemental Figure 17

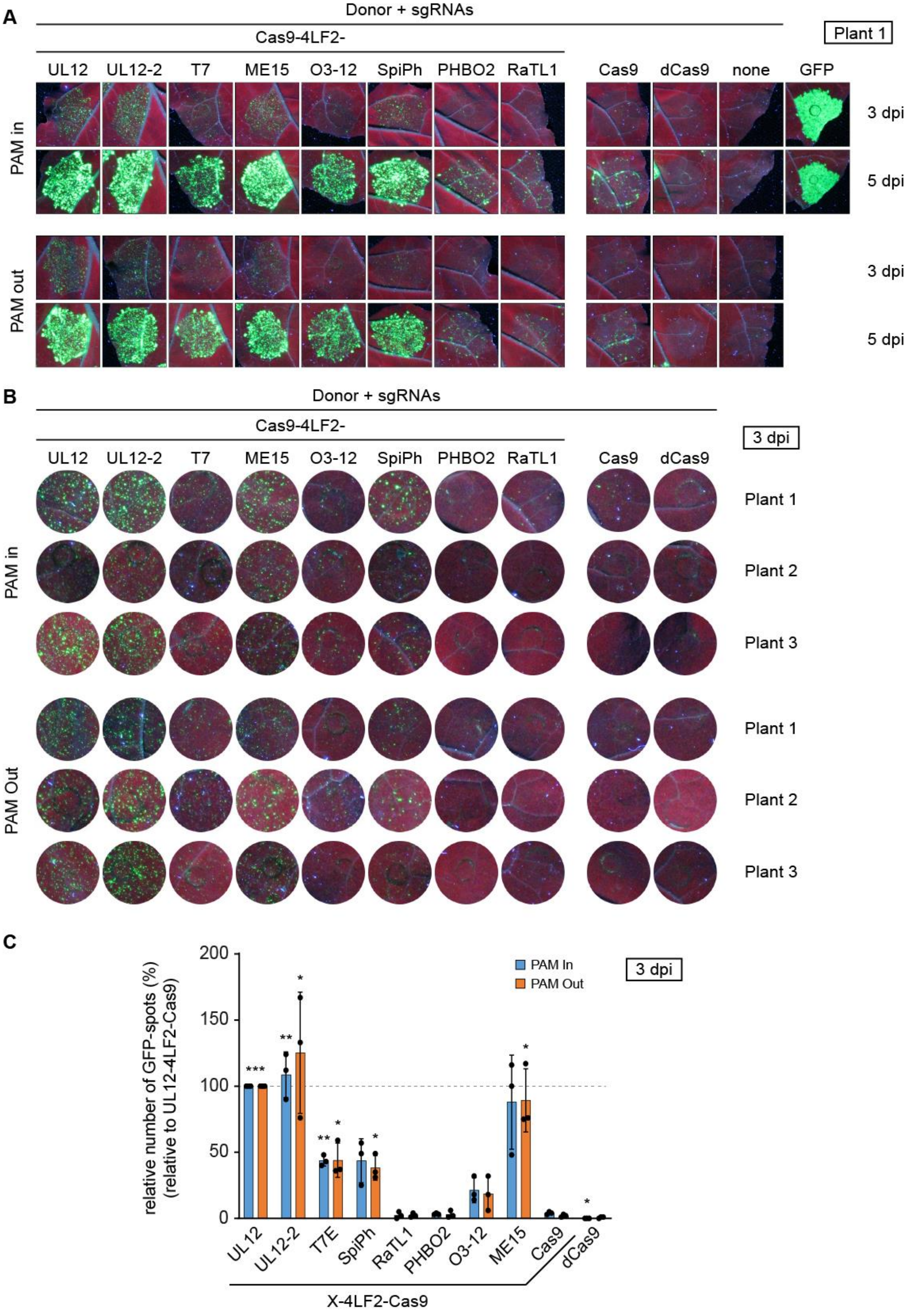

**Supplementary Fig. 17. Comparative analysis of Cas9-fused T7E homologues. (A)** Analysis of gene targeting efficiency after transient expression of indicated Cas9 variants in the transgenic Nb TMV reporter line (Plant1). **(B)** Areas of the same size from the infiltrated leaf discs taken for GFP spot count. **(C)** Summary of relative GFP spot counts for individual constructs. Number of GFP-spots are relative to UL12-fused Cas9 for which the GFP-spot number was set to 100%. Among the tested T7E homologues ME15 led to the highest number of GFP spots.

Supplemental Figure 18

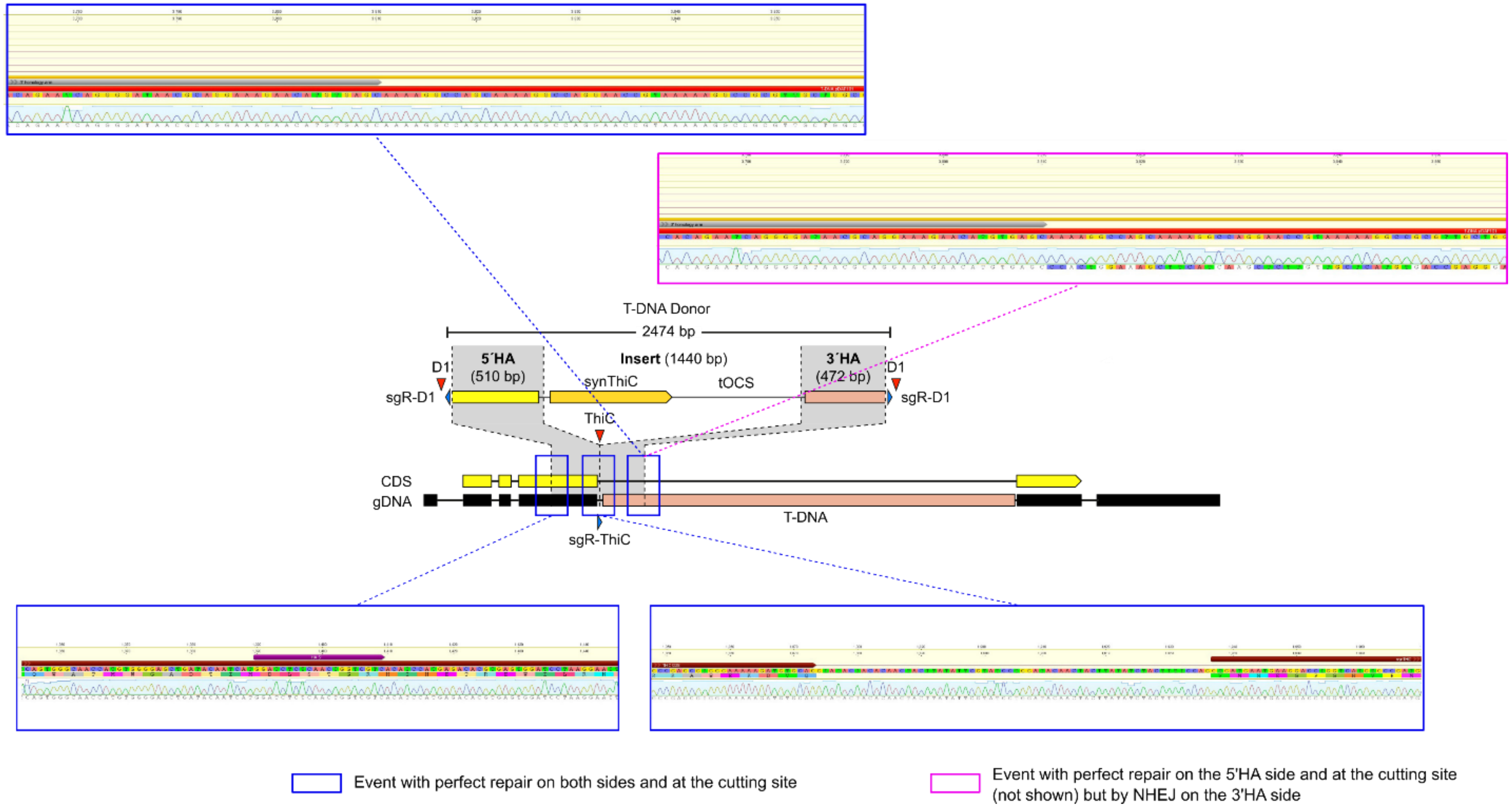


**Supplementary Fig. 18. Sequences of knock-ins at the *thic* locus.**

The scheme showing the structure of the locus and of the donor fragment is shown in the middle. Sequences framed in blue correspond to a successful knock-in event where both sides and the junction at the Cas9 cutting site correspond to the provided donor. The sequence framed in pink comes from an event where repair on the 3’HA side occurred via NHEJ as can be seen from the mismatched sequence beyond the homology arm. The expected sequence of the locus after perfect knock-in is shown above, the actual sequence chromatograms from the events is shown below.

**Supplementary Text**

<u>Note S1 - Transgenic Nicotiana benthamiana TMV-reporter line (Nbi775)</u>

To identify gene targeting events (precise repair) in *Nicotiana benthamiana*, we employed a transgenic reporter line containing a defective tobacco mosaic virus (TMV)-based reporter on the transgene. TMV is a (+)-RNA virus that encodes for three open reading frames (ORFs) on the (+) RNA, encoding for the RNA-dependent RNA polymerase (RdRP), the movement protein (MP), and the coat protein (CP). Upon injection into plant cells, the (+)-RNA undergoes translation by the host translational machinery, yet only results in the production of the initial ORF, the RdRP. Conversely, the expression of the MP and CP relies on a (-)-RNA intermediate, which acts as a template for producing MP- and CP-specific (+) RNAs from subgenomic promoters. The (-) RNA intermediate is created by the RdRP, which recognises a t-RNA-like structure located at the 3´ end of the genomic (+) RNA. In conclusion, neither the movement protein (MP) nor the coat protein (CP) are expressed in the absence of the RNA-dependent RNA polymerase (RdRP). The tobacco mosaic virus (TMV)-reporter has undergone two modifications. Firstly, the coat protein has been replaced by green fluorescent protein (GFP), rendering the TMV non-infectious and enabling the monitoring of viral replication through GFP fluorescence. Secondly, the RdRP has been disrupted by a 3.8 Kb deletion, replaced instead by a short 76 bp attB site. The attB site at the deletion endpoint, originally included in the construct for site-specific recombination, has no function in this context but rather acts as a target sequence for Cas9 or Cas12a cleavage. The defective TMV-reporter in the transgene is expressed constitutively by pAct2, but replication (and thus GFP expression) is reliant on precise repair of the RdRP. After inducing a DNA double strand break at this specific location, repairing the damage through homology-directed repair (HDR) using a T-DNA donor, which incorporates the missing RdRP fragment and flanking sequences that are homologous to both sides of the deletion endpoint, leads to the reconstruction of a functional TMV replicase. Consequently, there is GFP expression from subgenomic promoters (fig. S1). The presence of the MP enables the spread of Replication to surrounding cells, and facilitates the macroscopic monitoring of HDR events originating from a single cell. This can be achieved by directly observing the leaf under UV light. By quantifying the emerged GFP spots, gene targeting efficiencies can be determined.

Note S2 - Optimization of the Exo-Cas9 constructs

The *Nicotiana benthamiana* (Nb) TMV reporter line and the in-frame GUS-integration at the NbPGK locus in WT Nb allows the direct comparison of numerous constructs with a fast readout. Our assumption is that designs that perform well in transient assays are likely to perform similarly well in stable transformation approaches. We have utilized the TMV reporter to compare different linkers for fusion of UL12 or T7E to the N- or C-terminus of Cas9 (Fig S6). We examined the effectiveness of five versatile linkers, including two GGGGS-motif linkers (2xLF2 - $GS(GGGGS)_6$; 4LF2 - $GS(GGGGS)_{12}$), and three linkers derived from the XTEN linker (T16; T40; T144). For fusions of UL12 to Cas9, we were unable to find significant differences between the respective constructs. However, T7E favored linker 4LF2, regardless of whether it was fused to the N- or to the C-terminus of Cas9 (Fig. S6).

Molecular analyzes of the Cas9 DNA-binding mechanism have shown that there are strong interactions between the Cas9 protein and the target DNA at the PAM proximal site and that only the PAM-distal non-targeted strand is freely accessible. The dual sgRNA strategy was developed based on this idea. Accordingly, PAM in oriented targets should have a higher editing rate, since the 3' is directly accessible and the fragment to be replaced is shielded from repair at the ends by a Cas9 molecule. However, it appears that the activity is not affected by the orientation of the sgRNAs targeting the TMV-locus (PAM-In vs. PAM-Out; Fig. 1). To lower the risk of off-target cleavage, it would be beneficial if a single sgRNA could achieve comparable HDR rates. A direct comparison demonstrated that cleavage with a single sgRNA was as effective as using two sgRNAs (Fig. S8).

The impact of varying lengths of homology arms (HA) in the Donor T-DNA was also assessed (Fig. S7). TMV donor T-DNAs were created with HA lengths of 500bp, 250bp and 100bp on both the 5´ and 3´ sides, and compared to a donor with 1348bp and 777bp HA lengths on the 5´ and 3´ side, respectively. It was found that 250bp HAs resulted in the highest HDR efficiency, whereas 100bp HAs gave the lowest frequency of HDR (Fig. S7). Normally, a 1 kb homology arm length is employed for HDR donors, which means that the average length of the donor can be shortened by 1.5 kb.

When Cas9 is used as a nickase (D10A – no cleavage of the non-targeted strand; H840A – no cleavage of the targeted strand), the dual guide RNA approach can enhance specificity. Through nickases, a DNA double-stranded break only occurs when two close nicks are introduced in opposite strands (two on-targets). By using sgRNAs that target the TMV locus alongside Cas9 nickases, DNA double-stranded breaks with 44 base pair overhangs could be generated (see Fig. S1 and Fig. S9). Whether a 3' or 5' overhang is determined by the specific nickase used (Fig.S9A). In general, Cas9 nickases are more effective when applied in the PAM out orientation, as the interaction of Cas9 with the PAM-proximal DNA in the PAM in orientation might shield the dsDNA between the spatially close nicks from melting into a DNA double-strand break. The polarity bias for PAM out targets using Cas9 nickases was confirmed, but no effect of the DNA lesion (blunt end, 5' vs. 3' overhang) on gene targeting frequencies was detected (Fig.S9B). When fused to UL12 or T7E, the WT Cas9 resulted in the highest HDR frequencies upon introducing blunt end lesions (or two DNA DSBs) (Fig.S9B). If UL12 is fused to the C-terminus of Cas9, we observed higher HDR rates for Cas9(D10A) over Cas9(H840A) if applied in PAM in orientation (Fig.S9C and D).

The direct fusion of UL12 with Cas9 provides direct access to the DNA substrate and prevents UL12's mutagenic activity at unrelated DSB-sites. However, direct fusion could also negatively affect UL12 activity. We compared the Cas9-UL12 fusion with separately expressed Cas9 and UL12 at the NbPGK locus in WT Nb, using the in-frame GUS knock-in reporter (Fig.S10E). Measured GUS activities demonstrated that separate expression of UL12 and Cas9 increases gene targeting frequencies compared to just Cas9, although not to the same extent as the directly-fused UL12. To minimize the risk of off-target cleavage, we suggest using the Cas9-fused variant.

### Supporting Information

### Supplementary Table 1: Constructs used in this study

| | | Assembled from Level -1 and Level 0 Modules | | | | | |
|---|---|---|---|---|---|---|---|
| **Name** | **Construct (Level 1)** | **Promoter (Level 0)** | **Enhancer (Level 0)** | **Exo-Linker (Level -1)** | **Endonuclease /GOI (Level 0)** | **Linker-Exo (Level -1)** | **Terminator (Level 0)** |
| UL12-2xLF2-Cas9 | **pAGT6484** | pICH45089 | pICH41402 | pAGT6499, pAGT6467, pAGT6464 | pAGT6471 | - | pAGT5439 |
| Exo1-2xLF2-Cas9 | **pAGT7337** | pICH45089 | pICH41402 | pAGT7286, pAGT7288, pAGT6464 | pAGT6471 | - | pAGT5439 |
| T5E-2xLF2-Cas9 | **pAGT7339** | pICH45089 | pICH41402 | pAGT7323, pAGT6464 | pAGT6471 | - | pAGT5439 |
| T7E-2xLF2-Cas9 | **pAGT7340** | pICH45089 | pICH41402 | pAGT7324, pAGT6464 | pAGT6471 | - | pAGT5439 |
| Exo3-2xLF2-Cas9 | **pAGT7341** | pICH45089 | pICH41402 | pAGT7325, pAGT6464 | pAGT6471 | - | pAGT5439 |
| TREX1-2xLF2-Cas9 | **pAGT7342** | pICH45089 | pICH41402 | pAGT7326, pAGT6464 | pAGT6471 | - | pAGT5439 |
| UL12-4xLF2-Cas9 | **pAGT6487** | pICH45089 | pICH41402 | pAGT6499, pAGT6467, pAGT6463 | pAGT6471 | - | pAGT5439 |
| Exo1-4xLF2-Cas9 | **pAGT7343** | pICH45089 | pICH41402 | pAGT7286, pAGT7288, pAGT6463 | pAGT6471 | - | pAGT5439 |
| T5E-4xLF2-Cas9 | **pAGT7345** | pICH45089 | pICH41402 | pAGT7323, pAGT6463 | pAGT6471 | - | pAGT5439 |
| T7E-4xLF2-Cas9 | **pAGT7346** | pICH45089 | pICH41402 | pAGT7324, pAGT6463 | pAGT6471 | - | pAGT5439 |
| Exo3-4xLF2-Cas9 | **pAGT7347** | pICH45089 | pICH41402 | pAGT7325, pAGT6463 | pAGT6471 | - | pAGT5439 |
| TREX1-4xLF2-Cas9 | **pAGT7348** | pICH45089 | pICH41402 | pAGT7326, pAGT6463 | pAGT6471 | - | pAGT5439 |
| Cas9-2xLF2-UL12 | **pAGT6478** | pICH45089 | pICH41402 | - | pAGT6468 | pAGT6462, pAGT6499, pAGT6500 | pICH41432 |
| Cas9-2xLF2-Exo1 | **pAGT7310** | pICH45089 | pICH41402 | - | pAGT6468 | pAGT6462, pAGT7286, pAGT7287 | pICH41432 |
| Cas9-2xLF2-LaExo | **pAGT7311** | pICH45089 | pICH41402 | - | pAGT6468 | pAGT6462, pAGT7295 | pICH41432 |
| Cas9-2xLF2-T5 | **pAGT7312** | pICH45089 | pICH41402 | - | pAGT6468 | pAGT6462, pAGT7296 | pICH41432 |
| Cas9-2xLF2-T7 | **pAGT7313** | pICH45089 | pICH41402 | - | pAGT6468 | pAGT6462, pAGT7297 | pICH41432 |
| Cas9-2xLF2-Exo3 | **pAGT7314** | pICH45089 | pICH41402 | - | pAGT6468 | pAGT6462, pAGT7298 | pICH41432 |
| Cas9-2xLF2-TREX1 | **pAGT7315** | pICH45089 | pICH41402 | - | pAGT6468 | pAGT6462, pAGT7299 | pICH41432 |
| Cas9-4xLF2-UL12 | **pAGT6478** | pICH45089 | pICH41402 | - | pAGT6468 | pAGT6461, pAGT6499, pAGT6500 | pICH41432 |
| Cas9-4xLF2-Exo1 | **pAGT7310** | pICH45089 | pICH41402 | - | pAGT6468 | pAGT6461, pAGT7286, pAGT7287 | pICH41432 |
| Cas9-4xLF2-LaExo | **pAGT7311** | pICH45089 | pICH41402 | - | pAGT6468 | pAGT6461, pAGT7295 | pICH41432 |
| Cas9-4xLF2-T5 | **pAGT7312** | pICH45089 | pICH41402 | - | pAGT6468 | pAGT6461, pAGT7296 | pICH41432 |
| Cas9-4xLF2-T7 | **pAGT7313** | pICH45089 | pICH41402 | - | pAGT6468 | pAGT6461, pAGT7297 | pICH41432 |
| Cas9-4xLF2-Exo3 | **pAGT7314** | pICH45089 | pICH41402 | - | pAGT6468 | pAGT6461, pAGT7298 | pICH41432 |

| | | | | | | | |
|---|---|---|---|---|---|---|---|
| Cas9-4xLF2-TREX1 | **pAGT7315** | pICH45089 | pICH41402 | - | pAGT6468 | pAGT6461, pAGT7299 | pICH41432 |
| Cas9 | **pAGT5997** | pICH45089 | pICH41402 | - | pAGM47523 | - | pICH41432 |
| Cas9(D10A) | **pAGT8260** | pICH45089 | pICH41402 | - | pAGT6469 | - | pAGT5439 |
| Cas9(H840A) | **pAGT8261** | pICH45089 | pICH41402 | - | pAGT6470 | - | pAGT5439 |
| dCas9 | **pAGT7934** | pICH45089 | pICH41402 | - | pAGT6395 | - | pICH41432 |
| UL12(D340E)-2xLF2-Cas9 | **pAGT8114** | pICH45089 | pICH41402 | pAGT6499, pAGT8112, pAGT6464 | pAGT6471 | - | pAGT5439 |
| UL12(ΔN126)-2xLF2-Cas9 | **pAGT7642** | pICH45089 | pICH41402 | pAGT7289, pAGT6467, pAGT6464 | pAGT6471 | - | pAGT5439 |
| UL12(N1-126)-2xLF2-Cas9 | **pAGT7643** | pICH45089 | pICH41402 | pAGT7290, pAGT6464 | pAGT6471 | - | pAGT5439 |
| ICP8 | **pAGT7357** | pICH41388 | pICH41402 | - | pAGT7355 | - | pAGT5439 |
| SSB | **pAGT7356** | pICH41388 | pICH41402 | - | pAGT7354 | - | pAGT5439 |
| Rad51 | **pAGT6576** | pICH41388 | pICH41402 | - | pAGT6575 | - | pICH41432 |
| Rad52a | **pAGT7885** | pICH41388 | pICH41402 | - | pAGT7883 | - | pAGT5439 |
| Rad52b | **pAGT7886** | pICH41388 | pICH41402 | - | pAGT7884 | - | pAGT5439 |
| UL12 | **pAGT7407** | pICH41388 | pICH41402 | - | pAGT7406 | - | pAGT5439 |
| ICP8-NLS | **pAGT8090** | pICH41388 | pICH41402 | - | pAGT7355 | pAGT8089 | pICH41432 |
| SSB-NLS | **pAGT8091** | pICH41388 | pICH41402 | - | pAGT7354 | pAGT8089 | pICH41432 |
| UL12-T16-Cas9 | **pAGT8026** | pICH45089 | pICH41402 | pAGT6499, pAGT6467, pAGT6941 | pAGT6471 | - | pAGT5439 |
| UL12-T40-Cas9 | **pAGT8027** | pICH45089 | pICH41402 | pAGT6499, pAGT6467, pAGT6973 | pAGT6471 | - | pAGT5439 |
| UL12-T144-Cas9 | **pAGT8028** | pICH45089 | pICH41402 | pAGT6499, pAGT6467, pAGT6963 | pAGT6471 | - | pAGT5439 |
| Cas9-T16-UL12 | **pAGT8029** | pICH45089 | pICH41402 | - | pAGT6468 | pAGT8017, pAGT6499, pAGT6500 | pICH41432 |
| Cas9-T40-UL12 | **pAGT8030** | pICH45089 | pICH41402 | - | pAGT6468 | pAGT8018, pAGT6499, pAGT6500 | pICH41432 |
| Cas9-T144-UL12 | **pAGT8031** | pICH45089 | pICH41402 | - | pAGT6468 | pAGT8019, pAGT6499, pAGT6500 | pICH41432 |
| T7-T16-Cas9 | **pAGT8456** | pICH45089 | pICH41402 | pAGT7324, pAGT6941 | pAGT6471 | - | pAGT5439 |
| T7-T40-Cas9 | **pAGT8457** | pICH45089 | pICH41402 | pAGT7324, pAGT6973 | pAGT6471 | - | pAGT5439 |
| T7-T144-Cas9 | **pAGT8458** | pICH45089 | pICH41402 | pAGT7324, pAGT6963 | pAGT6471 | - | pAGT5439 |
| Cas9-T16-T7 | **pAGT8450** | pICH45089 | pICH41402 | - | pAGT6468 | pAGT8017, pAGT7297 | pICH41432 |
| Cas9-T40-T7 | **pAGT8451** | pICH45089 | pICH41402 | - | pAGT6468 | pAGT8018, pAGT7297 | pICH41432 |
| Cas9-T144-T7 | **pAGT8452** | pICH45089 | pICH41402 | - | pAGT6468 | pAGT8019, pAGT7297 | pICH41432 |
| UL12-4xLF2-Cas9(D10A) | **pAGT6488** | pICH45089 | pICH41402 | pAGT6499, pAGT6467, pAGT6463 | pAGT6472 | - | pAGT5439 |
| UL12-4xLF2-Cas9(H840A) | **pAGT6488** | pICH45089 | pICH41402 | pAGT6499, pAGT6467, pAGT6463 | pAGT6473 | - | pAGT5439 |
| Exo1-4xLF2-Cas9(D10A) | **pAGT8152** | pICH45089 | pICH41402 | pAGT7286, pAGT7288, pAGT6463 | pAGT6472 | - | pAGT5439 |
| Exo1-4xLF2-Cas9(H840A) | **pAGT8155** | pICH45089 | pICH41402 | pAGT7286, pAGT7288, pAGT6463 | pAGT6473 | - | pAGT5439 |
| T5E-4xLF2-Cas9(D10A) | **pAGT8153** | pICH45089 | pICH41402 | pAGT7323, pAGT6463 | pAGT6472 | - | pAGT5439 |
| T5E-4xLF2-Cas9(H840A) | **pAGT8156** | pICH45089 | pICH41402 | pAGT7323, pAGT6463 | pAGT6473 | - | pAGT5439 |

| | | | | | | | |
|---|---|---|---|---|---|---|---|
| T7E-4xLF2-Cas9(D10A) | **pAGT8154** | pICH45089 | pICH41402 | pAGT7324, pAGT6463 | pAGT6472 | - | pAGT5439 |
| T7E-4xLF2-Cas9(H840A) | **pAGT8157** | pICH45089 | pICH41402 | pAGT7324, pAGT6463 | pAGT6473 | - | pAGT5439 |
| UL12-2xLF2-Cas9(D10A) | **pAGT6484** | pICH45089 | pICH41402 | pAGT6499, pAGT6467, pAGT6464 | pAGT6472 | - | pAGT5439 |
| UL12-2xLF2-Cas9(H840A) | **pAGT6484** | pICH45089 | pICH41402 | pAGT6499, pAGT6467, pAGT6464 | pAGT6473 | - | pAGT5439 |
| Cas9(D10A)-2xLF2-UL12 | **pAGT6478** | pICH45089 | pICH41402 | - | pAGT6469 | pAGT6462, pAGT6499, pAGT6500 | pICH41432 |
| Cas9(H840A)-2xLF2-UL12 | **pAGT6478** | pICH45089 | pICH41402 | - | pAGT6470 | pAGT6462, pAGT6499, pAGT6500 | pICH41432 |
| Cas9(D10A)-4xLF2-UL12 | **pAGT6478** | pICH45089 | pICH41402 | - | pAGT6469 | pAGT6461, pAGT6499, pAGT6500 | pICH41432 |
| Cas9(H840A)-4xLF2-UL12 | **pAGT6478** | pICH45089 | pICH41402 | - | pAGT6470 | pAGT6461, pAGT6499, pAGT6500 | pICH41432 |
| 6xHis-Thrombin-UL12-6xHis (in pAGT8225) | **pAGT8232** | - | - | pAGT8226 | pAGT8228 | - | - |
| N-LaExo | **pAGT8115** | pICH41388 | pICH41402 | - | pAGT8081 | pAGT6462, pAGT7295 | pAGT5439 |
| LaExo-N | **pAGT8116** | pICH41388 | pICH41402 | pAGT7322, pAGT6464 | pAGT8080 | - | pAGT5439 |
| Cas12ai-4xLF2-UL12 | **pAGT8165** | pICH45089 | pICH41402 | - | pAGT8163 | pAGT6461, pAGT6499, pAGT6500 | pICH41432 |
| Cas12ai-4xLF2-T5 | **pAGT8168** | pICH45089 | pICH41402 | - | pAGT8163 | pAGT6461, pAGT7296 | pICH41432 |
| Cas12ai-4xLF2-T7 | **pAGT8169** | pICH45089 | pICH41402 | - | pAGT8163 | pAGT6461, pAGT7297 | pICH41432 |
| UL12-4xLF2-Cas12ai | **pAGT8175** | pICH45089 | pICH41402 | pAGT6499, pAGT6467, pAGT6463 | pAGT8173 | - | pAGT5439 |
| T5-4xLF2-Cas12ai | **pAGT8178** | pICH45089 | pICH41402 | pAGT7323, pAGT6463 | pAGT8173 | - | pAGT5439 |
| T7-4xLF2-Cas12ai | **pAGT8179** | pICH45089 | pICH41402 | pAGT7324, pAGT6463 | pAGT8173 | - | pAGT5439 |
| Cas12ai | **pAGT8186** | pICH45089 | pICH41402 | - | pAGT8163 | - | pAGT5439 |
| dCas12ai | **pAGT9121** | pICH45089 | pICH41402 | - | pAGT8162 | - | pAGT5439 |
| UL12-2-4xLF2-Cas9 | **pAGT7861** | pICH45089 | pICH41402 | pAGT7846, pAGT7847, pAGT6463 | pAGT6471 | - | pAGT5439 |
| PapE-4xLF2-Cas9 | **pAGT9054** | pICH45089 | pICH41402 | pAGT9046, pAGT6463 | pAGT6471 | - | pAGT5439 |
| PiE-4xLF2-Cas9 | **pAGT9055** | pICH45089 | pICH41402 | pAGT9047, pAGT6463 | pAGT6471 | - | pAGT5439 |
| AB4P-4xLF2-Cas9 | **pAGT8111** | pICH45089 | pICH41402 | pAGT8098, pAGT8099, pAGT6463 | pAGT6471 | - | pAGT5439 |
| MD5-4xLF2-Cas9 | **pAGT8110** | pICH45089 | pICH41402 | pAGT8096, pAGT8097, pAGT6463 | pAGT6471 | - | pAGT5439 |
| Dumas-4xLF2-Cas9 | **pAGT8109** | pICH45089 | pICH41402 | pAGT8094, pAGT8095, pAGT6463 | pAGT6471 | - | pAGT5439 |
| BGLF5-4xLF2-Cas9 | **pAGT7862** | pICH45089 | pICH41402 | pAGT7848, pAGT7849, pAGT6463 | pAGT6471 | - | pAGT5439 |
| SOX-4xLF2-Cas9 | **pAGT7863** | pICH45089 | pICH41402 | pAGT7850, pAGT7851, pAGT6463 | pAGT6471 | - | pAGT5439 |
| Cas9-4xLF2-ME15 | **pAGT8750** | pICH45089 | pICH41402 | - | pAGT6468 | pAGT6461, pAGT8740 | pICH41432 |
| Cas9-4xLF2-O3-12 | **pAGT8751** | pICH45089 | pICH41402 | - | pAGT6468 | pAGT6461, pAGT8741 | pICH41432 |

| | | | | | | | |
|---|---|---|---|---|---|---|---|
| Cas9-4xLF2-SpiPh | **pAGT8752** | pICH45089 | pICH41402 | - | pAGT6468 | pAGT6461, pAGT8742 | pICH41432 |
| Cas9-4xLF2-PHBO2 | **pAGT8753** | pICH45089 | pICH41402 | - | pAGT6468 | pAGT6461, pAGT8743 | pICH41432 |
| Cas9-4xLF2-RaTL1 | **pAGT8754** | pICH45089 | pICH41402 | - | pAGT6468 | pAGT6461, pAGT8744 | pICH41432 |
| At Cas9 | **pAGM51323** | pJOG603 | - | - | pAGM47523 | - | pICH49344 |
| At PapE-4xLF2-Cas9 | **pAGT9569** | pJOG603 | - | pAGT9046, pAGT6463 | pAGT6471 | - | pAGT8149 |
| At ME15-4xLF2-Cas9 | **pAGT9570** | pJOG603 | - | pAGT9278, pAGT6463 | pAGT6471 | - | pAGT8149 |

**Level -1 Modules (frame underlined)**

**pAGT6461 (Linker: TTCG_4xLF2_AATG)**
GG-overhang_4xLF2_GG-overhang
TTCGGGTGGAGGTGGTTCTGGTGGAGGTGGATCAGGAGGAGGAGGTTCGGGTGGAGGTGGTTCTGGTGGAGGTGGAT
CAGGAGGAGGAGGATCAGGTGGAGGTGGTTCTGGTGGAGGTGGATCAGGAGGAGGAGGTTCAGGTGGAGGTGGTTCT
GGTGGAGGTGGATCAGGAGGAGGAGGATCAATG

**pAGT6461 (Linker: TTCG_2xLF2_AATG)**
GG-overhang_2xLF2_GG-overhang
TTCGGGTGGAGGTGGTTCTGGTGGAGGTGGATCAGGAGGAGGAGGTTCGGGTGGAGGTGGTTCTGGTGGAGGTGGAT
CAGGAGGAGGAGGATCAATG

**pAGT8017 (Linker: TTCG_XTEN16_AATG)**
GG-overhang_XTEN16_GG-overhang
TTCGGGTTCAGAAACCCCTGGTACTAGCGAGTCAGCTACACCAGAGTCAATG

**pAGT8018 (Linker: TTCG_XTEN40_AATG)**
GG-overhang_XTEN40_GG-overhang
TTCGGGTTCAGAAACCCCTGGTACTAGCGAGTCAGCTACACCAGAGTCTGGACCAGGATCTGAACCTGCTACTAGTG
GATCTGAGACACCTGGAACTAGTGAGAGTGCAACTCCTGAGTCAATG

**pAGT8019 (Linker: TTCG_XTEN144_AATG)**
GG-overhang_XTEN40_GG-overhang
TTCGGGTACTTCTACTGAGCCTTCTGAAGGTTCTGCTCCTGGAACTTCTGAGTCTGCTACTCCTGAATCTGGTCCTG
GTTCTGAGCCAGCTACTTCTGGTTCAGAAACCCCTGGTACTAGCGAGTCAGCTACACCAGAGTCTGGACCAGGATCT
GAACCTGCTACTAGTGGATCTGAGACACCTGGAACTAGTGAGAGTGCAACTCCTGAGTCAGGACCTGGTACTTCAAC
AGAACCTAGTGAGGGTAGTGCTCCAGGCACTAGTGAATCTGCAACTCCAGAAAGTGGACCTGGATCTCCTGCTGGTT
CTCCTACTTCTACAGAAGAGGGTAGTCCTGCTGGAAGCCCTACATCTACTGAAGAAGGTTCTCCAGCTGGCAGTCCA
ACTTCAACTGAAGAGGGAACTTCAGAGAGCGCTACACCTGAAAGTGGTCCAATG

**pAGT6463 (Linker: TTCG_4xLF2_AGGT)**
GG-overhang_4xLF2_GG-overhang
TTCGGGTGGAGGTGGTTCTGGTGGAGGTGGATCAGGAGGAGGAGGTTCGGGTGGAGGTGGTTCTGGTGGAGGTGGAT
CAGGAGGAGGAGGATCAGGTGGAGGTGGTTCTGGTGGAGGTGGATCAGGAGGAGGAGGTTCAGGTGGAGGTGGTTCT
GGTGGAGGTGGATCAGGAGGAGGAGGATCAGGT

**pAGT6464 (Linker: TTCG_2xLF2_AGGT)**
GG-overhang_2xLF2_GG-overhang
TTCGGGTGGAGGTGGTTCTGGTGGAGGTGGATCAGGAGGAGGAGGTTCGGGTGGAGGTGGTTCTGGTGGAGGTGGAT
CAGGAGGAGGAGGATCAGGT

**pAGT6941 (Linker: TTCG_XTEN16_AGGT)**
GG-overhang_XTEN16_GG-overhang
TTCGGGTTCAGAAACCCCTGGTACTAGCGAGTCAGCTACACCAGAGTCAGGT

**pAGT6973 (Linker: TTCG_XTEN40_AGGT)**
GG-overhang_XTEN40_GG-overhang
TTCGGGTTCAGAAACCCCTGGTACTAGCGAGTCAGCTACACCAGAGTCTGGACCAGGATCTGAACCTGCTACTAGTG
GATCTGAGACACCTGGAACTAGTGAGAGTGCAACTCCTGAGTCAGGT

**pAGT6963 (Linker: TTCG_XTEN144_AGGT)**
GG-overhang_XTEN144_GG-overhang
TTCGGGTACTTCTACTGAGCCTTCTGAAGGTTCTGCTCCTGGAACTTCTGAGTCTGCTACTCCTGAATCTGGTCCTG
GTTCTGAGCCAGCTACTTCTGGTTCAGAAACCCCTGGTACTAGCGAGTCAGCTACACCAGAGTCTGGACCAGGATCT
GAACCTGCTACTAGTGGATCTGAGACACCTGGAACTAGTGAGAGTGCAACTCCTGAGTCAGGACCTGGTACTTCAAC
AGAACCTAGTGAGGGTAGTGCTCCAGGCACTAGTGAATCTGCAACTCCAGAAAGTGGACCTGGATCTCCTGCTGGTT
CTCCTACTTCTACAGAAGAGGGTAGTCCTGCTGGAAGCCCTACATCTACTGAAGAAGGTTCTCCAGCTGGCAGTCCA
ACTTCAACTGAAGAGGGAACTTCAGAGAGCGCTACACCTGAAAGTGGTCCAGGT

**pAGT6499 (UL12 module A: AATG_UL12-A_CACC)**
GG-overhang_UL12-A_GG-overhang
AATGGAATCTACTGTGGGTCCTGCTTGTCCTCCTGGTAGGACTGTTACTAAGAGGCCTTGGGCTCTTGCTGAGGATA
CTCCTAGAGGTCCTGACTCTCCACCAAAGAGGCCTAGACCTAACTCTCTTCCTCTGACTACTACCTTCAGGCCTTTG
CCACCTCCTCCACAAACTACCTCTGCTGTGGATCCTTCTTCTCACTCTCCTGTGAATCCTCCAAGGGATCAGCATGC
TACTGATACCGCTGATGAGAAGCCTAGAGCTGCTTCTCCTGCTCTGTCTGATGCTTCTGGTCCTCCTACTCCTGATA
TCCCTCTTTCTCCTGGTGGTACTCATGCTAGGGATCCTGATGCAGATCCTGATAGCCCTGATCTGGACTCTATGTGG
TCTGCTTCTGTGATCCCTAACGCTCTGCCTTCTCACATTCTGGCTGAGACTTTCGAGAGGCACCTTAGGGGTTTGCT
TAGAGGTGTTAGGGCTCCTCTTGCTATTGGTCCTCTTTGGGCTAGACTGGACTACCTTTGCTCTCTTGCTGTGGTGC
TTGAAGAGGCTGGTATGGTGGATAGAGGTCTTGGTAGACACCTTTGGAGGCTTACTAGAAGAGGTCCTCCAGCTGCT
GCTGATGCTGTTGCTCCTAGACCTCTTATGGGATTCTACGAGGCTGCTACTCAGAACCAGGCTGATTGTCAACTTTG
GGCTCTGCTTAGAAGGGGTCTTACTACCGCTTCTACTCTTAGATGGGGTCCTCAGGGACCTTGCTTTTCTCCTCAAT
GGCTGAAACACAACGCTAGCCTTAGGCCTGATGTGCAGTCATCTGCTGTGATGTTCGGTAGGGTTAACGAGCCTACC
GCTCGGTCTTTGCTTTTCAGGTATTGCGTTGGCAGGGCTGATGATGGTGGTGAAGCTGGTGCTGATACCAGGCGGTT
TATTTTCCAC

**pAGT7289 (UL12(ΔN126) module A: AATG_UL12(ΔN126)-A_CACC)**
GG-overhang_UL12(ΔN126)-A_GG-overhang
AATGTGGTCTGCTTCTGTGATCCCTAACGCTCTGCCTTCTCACATTCTGGCTGAGACTTTCGAGAGGCACCTTAGGG
GTTTGCTTAGAGGTGTTAGGGCTCCTCTTGCTATTGGTCCTCTTTGGGCTAGACTGGACTACCTTTGCTCTCTTGCT
GTGGTGCTTGAAGAGGCTGGTATGGTGGATAGAGGTCTTGGTAGACACCTTTGGAGGCTTACTAGAAGAGGTCCTCC
AGCTGCTGCTGATGCTGTTGCTCCTAGACCTCTTATGGGATTCTACGAGGCTGCTACTCAGAACCAGGCTGATTGTC
AACTTTGGGCTCTGCTTAGAAGGGGTCTTACTACCGCTTCTACTCTTAGATGGGGTCCTCAGGGACCTTGCTTTTCT
CCTCAATGGCTGAAACACAACGCTAGCCTTAGGCCTGATGTGCAGTCATCTGCTGTGATGTTCGGTAGGGTTAACGA
GCCTACCGCTCGGTCTTTGCTTTTCAGGTATTGCGTTGGCAGGGCTGATGATGGTGGTGAAGCTGGTGCTGATACCA
GGCGGTTTATTTTCCAC

**pAGT6500 (UL12 module B1: CACC_UL12-B1-stop_GCTT)**
GG-overhang_UL12-B1_GG-overhang
CCACGAGCCATCTGATCTGGCCGAAGAGAATGTTCATACCTGCGGTGTGCTTATGGATGGTCACACTGGAATGGTGG
GCGCTTCTCTTGATATTCTTGTGTGCCCTAGGGACATCCACGGTTACCTTGCTCCAGTTCCTAAGACTCCTCTGGCC
TTTTACGAGGTTAAGTGCAGGGCTAAGTACGCTTTCGATCCTATGGACCCTTCTGACCCTACTGCTTCTGCTTACGA
GGATCTGATGGCTCATAGAAGCCCTGAGGCTTTCAGGGCTTTCATCCGGTCTATTCCTAAGCCGAGCGTGAGATACT
TTGCTCCTGGAAGAGTTCCTGGTCCTGAGGAAGCTCTTGTTACTCAAGATCAGGCTTGGTCTGAGGCTCATGCTTCA
GGTGAGAAGAGAAGATGCTCAGCTGCTGATAGGGCACTCGTTGAGCTTAATTCTGGCGTGGTGTCTGAGGTGTTGCT
TTTCGGTGCTCCTGATCTCGGTAGGCACACTATTTCTCCAGTGAGCTGGTCCTCTGGTGATCTTGTTAGAAGGGAAC
CCGTGTTCGCTAATCCTAGGCACCCTAACTTCAAGCAGATTCTGGTGCAGGGTTACGTGCTGGATTCTCACTTTCCA
GATTGCCCTCCACATCCTCACCTTGTGACTTTCATTGGTCGGCATAGGACCTCAGCTGAAGAGGGTGTTACTTTCAG
GCTTGAGGATGGTGCTGGTGCTCTTGGTGCTGCTGGTCCTTCTAAGGCTTCTATTCTTCCTAACCAGGCCGTGCCTA

TCGCTCTTATTATCACCCCTGTGAGGATCGACCCCGAGATCTATAAGGCTATCCAGAGGTCATCTCGGCTGGCTTTC
GATGATACTTTGGCTGAGCTTTGGGCCTCTAGATCTCCTGGTCCAGGTCCTGCTGCTGCAGAAACTACTTCTTCTTC
ACCTACCACCGGCAGGTCATCTAGATAGGCTT

**pAGT6467 (UL12 module B1: CACC_UL12-B2- no stop_TTCG)**

GG-overhang_UL12-B2_GG-overhang

CCACGAGCCATCTGATCTGGCCGAAGAGAATGTTCATACCTGCGGTGTGCTTATGGATGGTCACACTGGAATGGTGG
GCGCTTCTCTTGATATTCTTGTGTGCCCTAGGGACATCCACGGTTACCTTGCTCCAGTTCCTAAGACTCCTCTGGCC
TTTTACGAGGTTAAGTGCAGGGCTAAGTACGCTTTCGATCCTATGGACCCTTCTGACCCTACTGCTTCTGCTTACGA
GGATCTGATGGCTCATAGAAGCCCTGAGGCTTTCAGGGCTTTCATCCGGTCTATTCCTAAGCCGAGCGTGAGATACT
TTGCTCCTGGAAGAGTTCCTGGTCCTGAGGAAGCTCTTGTTACTCAAGATCAGGCTTGGTCTGAGGCTCATGCTTCA
GGTGAGAAGAGAAGATGCTCAGCTGCTGATAGGGCACTCGTTGAGCTTAATTCTGGCGTGGTGTCTGAGGTGTTGCT
TTTCGGTGCTCCTGATCTCGGTAGGCACACTATTTCTCCAGTGAGCTGGTCCTCTGGTGATCTTGTTAGAAGGGAAC
CCGTGTTCGCTAATCCTAGGCACCCTAACTTCAAGCAGATTCTGGTGCAGGGTTACGTGCTGGATTCTCACTTTCCA
GATTGCCCTCCACATCCTCACCTTGTGACTTTCATTGGTCGGCATAGGACCTCAGCTGAAGAGGGTGTTACTTTCAG
GCTTGAGGATGGTGCTGGTGCTCTTGGTGCTGCTGGTCCTTCTAAGGCTTCTATTCTTCCTAACCAGGCCGTGCCTA
TCGCTCTTATTATCACCCCTGTGAGGATCGACCCCGAGATCTATAAGGCTATCCAGAGGTCATCTCGGCTGGCTTTC
GATGATACTTTGGCTGAGCTTTGGGCCTCTAGATCTCCTGGTCCAGGTCCTGCTGCTGCAGAAACTACTTCTTCTTC
ACCTACCACCGGCAGGTCATCTAGAGGTTCG

**pAGT8112 (dUL12 module B1: CACC_dUL12-B2- no stop_TTCG)**

GG-overhang_dUL12-B2_GG-overhang

CCACGAGCCATCTGATCTGGCCGAAGAGAATGTTCATACCTGCGGTGTGCTTATGGATGGTCACACTGGAATGGTGG
CTGCTGCTCTTGATATTCTTGTGTGCCCTAGGGACATCCACGGTTACCTTGCTCCAGTTCCTAAGACTCCTCTGGCC
TTTTACGAGGTTAAGTGCAGGGCTAAGTACGCTTTCGATCCTATGGACCCTTCTGACCCTACTGCTTCTGCTTACGA
GGATCTGATGGCTCATAGAAGCCCTGAGGCTTTCAGGGCTTTCATCCGGTCTATTCCTAAGCCGAGCGTGAGATACT
TTGCTCCTGGAAGAGTTCCTGGTCCTGAGGAAGCTCTTGTTACTCAAGATCAGGCTTGGTCTGAGGCTCATGCTTCA
GGTGAGAAGAGAAGATGCTCAGCTGCTGATAGGGCACTCGTTGAGCTTAATTCTGGCGTGGTGTCTGAGGTGTTGCT
TTTCGGTGCTCCTGATCTCGGTAGGCACACTATTTCTCCAGTGAGCTGGTCCTCTGGTGATCTTGTTAGAAGGGAAC
CCGTGTTCGCTAATCCTAGGCACCCTAACTTCAAGCAGATTCTGGTGCAGGGTTACGTGCTGGATTCTCACTTTCCA
GATTGCCCTCCACATCCTCACCTTGTGACTTTCATTGGTCGGCATAGGACCTCAGCTGAAGAGGGTGTTACTTTCAG
GCTTGAGGATGGTGCTGGTGCTCTTGGTGCTGCTGGTCCTTCTAAGGCTTCTATTCTTCCTAACCAGGCCGTGCCTA
TCGCTCTTATTATCACCCCTGTGAGGATCGACCCCGAGATCTATAAGGCTATCCAGAGGTCATCTCGGCTGGCTTTC
GATGATACTTTGGCTGAGCTTTGGGCCTCTAGATCTCCTGGTCCAGGTCCTGCTGCTGCAGAAACTACTTCTTCTTC
ACCTACCACCGGCAGGTCATCTAGAGGTTCG

**pAGT7290 (UL12(1-126) module: AATG_UL12(1-126)_TTCG)**

GG-overhang_UL12(1-126)_GG-overhang

AATGGAATCTACTGTGGGTCCTGCTTGTCCTCCTGGTAGGACTGTTACTAAGAGGCCTTGGGCTCTTGCTGAGGATA
CTCCTAGAGGTCCTGACTCTCCACCAAAGAGGCCTAGACCTAACTCTCTTCCTCTGACTACTACCTTCAGGCCTTTG
CCACCTCCTCCACAAACTACCTCTGCTGTGGATCCTTCTTCTCACTCTCCTGTGAATCCTCCAAGGGATCAGCATGC
TACTGATACCGCTGATGAGAAGCCTAGAGCTGCTTCTCCTGCTCTGTCTGATGCTTCTGGTCCTCCTACTCCTGATA
TCCCTCTTTCTCCTGGTGGTACTCATGCTAGGGATCCTGATGCAGATCCTGATAGCCCTGATCTGGACTCTGGTTCG

**pAGT7286 (AtExo1 module A: AATG_AtExo1-A_CTTG)**

GG-overhang_AtExo1-A_GG-overhang

AATGGGTATACAAGGGCTTTTGCCGTTGTTGAAATCGATAATGGTACCGATTCATATCAAGGAGCTCGAGGGTTGTA
TCGTCGCCGTCGATACATATTCATGGCTACACAAAGGGGCTTTATCTTGTAGCCGAGAGCTCTGCAAGGGATTACCC
ACCAAGAGGCATATTCAATACTGTATGCATAGAGTGAATTTACTTCGCCATCATGGAGTGAAGCCTATCATGGTCTT
TGATGGAGGTCCGTTACCGATGAAACTAGAACAAGAGAATAAGCGTGCTAGGTCCAGAAAGGAAAATCTTGCTCGTG
CATTGGAGCATGAAGCAAATGGAAATTCCTCAGCTGCTTATGAGTGCTACTCAAAGGCCGTAGATATTTCACCTTCT
ATTGCACATGAGTTGATACAGGTTCTGAGGCAGGAGAATGTTGATTACGTTGTTGCTCCTTACGAAGCTGATGCGCA
GATGGCATTCTTAGCTATTACTAAGCAAGTTGATGCGATTATCACTGAGGATTCTGATCTCATACCTTTTGGCTGCC
TAAGAATCATTTTCAAAATGGACAAGTTTGGTCATGGTGTTGAGTTTCAAGCCTCCAAGCTACCCAAAAATAAGGAT
CTCAGTCTATCAGGATTTTCAAGCCAAATGCTTCTCGAAATGTGCATATTAAGTGGCTGCGATTATTTGCAGTCACT
TCCAGGAATGGGACTCAAAAGAGCACATGCACTCATTACAAAATTCAAAAGCTATGACAGGGTAATTAAGCACTTAA
AGTACAGCACAGTTTCAGTTCCTCCTCTTTATGAAGAGTCATTCAAGAGAGCTTTACTGACTTTCAAGCATCAACGT
GTCTATGACCCAAATGCTGAAGATATCATACACTTGTGTGACATTTCTGACAACCTTGGTGAAGATTCAGATTTTGT

AGGCCCATCGATGCCACAAGATATTGCTAAGGGTATAGCTCTAGGCCAGCTTGATCCCTTCACACAGTTGCCATTCC
AGGCTGAGAGTGTTACTCCTAAATTGGCTGTTGATGATATATCCCGTCCCAAAAGTTTCAAACCTGAAACTGTAAAG
AAAAAGCTTG

**pAGT7287 (AtExo1 module B1: CTTG_AtExo1-B1-stop_GCTT)**

GG-overhang_AtExo1-B1_GG-overhang

CTTGATTTGCCAGTCCAAAAAAACCTTCTCACCAAGTACTTCTGCTTTGCGTCTGTTGAGGCAAAAAGAAAATTCAA
GGCTCCTAGGATATCACCAATGTCTCTGACCCCAACTGATGAGTCTCCAAGCATTCCTGATGACAACACACCAGATT
TAGATGCTCTTTCAAGCCAGACAACAAATGAATCCCCGGTTTATAGCTTGGGTGAGAATCCATGTGTTAGCGAAGTT
GCAGAGAAAAGAGATTCACCCGATGACGATGCAGTGGAAAGGAATCACAAGGACCTTCATCATAAATACTGCGAACG
GGAAGTGGATAGGCCGAAGTCAGATAGTTTGAAGGTTATAGTAAGAAGCAAGTATTTTAAGCAGAAACAGGAGGACA
AAAGTCTGAAACAATCGATCCCATGTCTTAATGATTGCTCTGTGATTGGTCAGAGAAAGGCTGTTAAAACTGTGATT
AACATGTCAAGCGCGTCTAAAAGAGAAGAGAGTCACAGAGCTATAGCAACAAGTCCATGTTTACACCATGACCGAAT
CTACAACGATCATGAAGATGCTAAAGAAGCGAGTTTCTCTGCCATGAACGAGGTGGCTGAGAGAACAATAAACACCC
ACAAGATTAACCATCAGATCAATGAAGAAGAACAGAACCCATCCGTTGAAATTCCTTCTGCCTTTAGTACTCCCGAG
AATGTTATACCTCTCTCTTCCATTGCTATTGATAGTTGCCATGGAGTTGCAACAGGGAAGAGGAAGCTTGACTCAGA
TGAAAACCTTCACAAGGAAAATTTGAAATCCAAGCACATGCGCATGGATGAAACTGATACTGCTCTTAATGCAGAAA
CACCATTAGAAACCGATGATGTTGAGAAGTTTGGATCGAACATTTCACATATAGGGCATTACTCAGAAATAGCAGAG
AAATCAGTGGAAAGATTTGTCTCAGCTATATCGTCTTTCAAATATTCTGGGACTGGCTCTCGTGCTAGCGGACTCCG
TGCCCCTCTCAAAGATATCCGCAACACTTGTCCGTCCAAAGGGTTATCTCTTAAACCAGACATCAGCAAGTTTGGCT
ATGCGTCAAGCAACCGACACATGGTGACAAAGTCAAGGAGAATGTGAGCTT

**pAGT7288 (AtExo1 module B2: CTTG_AtExo1-B2-no stop_TTCG)**

GG-overhang_AtExo1-B2_GG-overhang

CTTGATTTGCCAGTCCAAAAAAACCTTCTCACCAAGTACTTCTGCTTTGCGTCTGTTGAGGCAAAAAGAAAATTCAA
GGCTCCTAGGATATCACCAATGTCTCTGACCCCAACTGATGAGTCTCCAAGCATTCCTGATGACAACACACCAGATT
TAGATGCTCTTTCAAGCCAGACAACAAATGAATCCCCGGTTTATAGCTTGGGTGAGAATCCATGTGTTAGCGAAGTT
GCAGAGAAAAGAGATTCACCCGATGACGATGCAGTGGAAAGGAATCACAAGGACCTTCATCATAAATACTGCGAACG
GGAAGTGGATAGGCCGAAGTCAGATAGTTTGAAGGTTATAGTAAGAAGCAAGTATTTTAAGCAGAAACAGGAGGACA
AAAGTCTGAAACAATCGATCCCATGTCTTAATGATTGCTCTGTGATTGGTCAGAGAAAGGCTGTTAAAACTGTGATT
AACATGTCAAGCGCGTCTAAAAGAGAAGAGAGTCACAGAGCTATAGCAACAAGTCCATGTTTACACCATGACCGAAT
CTACAACGATCATGAAGATGCTAAAGAAGCGAGTTTCTCTGCCATGAACGAGGTGGCTGAGAGAACAATAAACACCC
ACAAGATTAACCATCAGATCAATGAAGAAGAACAGAACCCATCCGTTGAAATTCCTTCTGCCTTTAGTACTCCCGAG
AATGTTATACCTCTCTCTTCCATTGCTATTGATAGTTGCCATGGAGTTGCAACAGGGAAGAGGAAGCTTGACTCAGA
TGAAAACCTTCACAAGGAAAATTTGAAATCCAAGCACATGCGCATGGATGAAACTGATACTGCTCTTAATGCAGAAA
CACCATTAGAAACCGATGATGTTGAGAAGTTTGGATCGAACATTTCACATATAGGGCATTACTCAGAAATAGCAGAG
AAATCAGTGGAAAGATTTGTCTCAGCTATATCGTCTTTCAAATATTCTGGGACTGGCTCTCGTGCTAGCGGACTCCG
TGCCCCTCTCAAAGATATCCGCAACACTTGTCCGTCCAAAGGGTTATCTCTTAAACCAGACATCAGCAAGTTTGGCT
ATGCGTCAAGCAACCGACACATGGTGACAAAGTCAAGGAGAATGGGTTCG

**pAGT7295 (Lambda Exo module: AATG_LaExo-stop_GCTT)**

GG-overhang_LaExo1_GG-overhang

AATGACCCCTGATATCATTCTGCAGAGGACCGGTATTGACGTGAGAGCTGTTGAACAGGGTGATGACGCTTGGCACA
AGCTTAGGCTTGGTGTGATTACCGCTAGCGAGGTGCACAATGTGATTGCTAAGCCTCGGAGCGGTAAGAAATGGCCG
GATATGAAGATGAGCTACTTCCACACCTTGCTGGCTGAAGTGTGTACTGGTGTTGCTCCTGAGGTTAACGCTAAGGC
TCTTGCTTGGGGTAAGCAGTACGAGAATGATGCTAGGACCCTGTTCGAGTTCACCAGCGGTGTTAATGTGACCGAGT
CTCCGATCATCTACCGGGATGAGTCTATGAGGACTGCTTGCTCTCCTGATGGTCTGTGCTCTGATGGTAACGGTCTT
GAGCTTAAGTGCCCGTTCACCTCCAGGGATTTCATGAAGTTCAGGCTTGGAGGCTTCGAGGCTATCAAGTCTGCTTA
CATGGCTCAGGTGCAGTACTCTATGTGGGTGACCAGAAAGAACGCTTGGTACTTCGCTAACTACGACCCGAGGATGA
AGCGTGAGGGACTTCATTACGTTGTGATCGAGAGGGACGAGAAGTACATGGCTAGCTTCGATGAGATCGTGCCCGAG
TTCATCGAGAAGATGGATGAAGCTCTTGCCGAGATCGGTTTCGTTTTTGGTGAACAGTGGCGGTGAGCTT

**pAGT7322 (Lambda Exo module: AATG_LaExo-no stop_TTCG)**

GG-overhang_LaExo1_GG-overhang

AATGACCCCTGATATCATTCTGCAGAGGACCGGTATTGACGTGAGAGCTGTTGAACAGGGTGATGACGCTTGGCACA
AGCTTAGGCTTGGTGTGATTACCGCTAGCGAGGTGCACAATGTGATTGCTAAGCCTCGGAGCGGTAAGAAATGGCCG
GATATGAAGATGAGCTACTTCCACACCTTGCTGGCTGAAGTGTGTACTGGTGTTGCTCCTGAGGTTAACGCTAAGGC
TCTTGCTTGGGGTAAGCAGTACGAGAATGATGCTAGGACCCTGTTCGAGTTCACCAGCGGTGTTAATGTGACCGAGT

CTCCGATCATCTACCGGGATGAGTCTATGAGGACTGCTTGCTCTCCTGATGGTCTGTGCTCTGATGGTAACGGTCTT
GAGCTTAAGTGCCCGTTCACCTCCAGGGATTTCATGAAGTTCAGGCTTGGAGGCTTCGAGGCTATCAAGTCTGCTTA
CATGGCTCAGGTGCAGTACTCTATGTGGGTGACCAGAAAGAACGCTTGGTACTTCGCTAACTACGACCCGAGGATGA
AGCGTGAGGGACTTCATTACGTTGTGATCGAGAGGGACGAGAAGTACATGGCTAGCTTCGATGAGATCGTGCCCGAG
TTCATCGAGAAGATGGATGAAGCTCTTGCCGAGATCGGTTTCGTTTTTGGTGAACAGTGGCGGGGTTCG

**pAGT7296 (T5 Exo module: AATG_T5Exo-stop_GCTT)**

GG-overhang_T5Exo_GG-overhang

AATGTCTAAGAGCTGGGGCAAGTTCATCGAAGAGGAAGAGGCTGAGATGGCCTCTCGGAGGAATCTTATGATCGTGG
ATGGTACTAACCTCGGCTTCCGGTTCAAGCACAACAACAGCAAGAAGCCGTTCGCCTCCTCTTACGTGTCAACCATT
CAGAGCCTGGCCAAGTCTTACTCTGCTAGGACTACCATTGTGCTGGGCGATAAGGGAAAGTCTGTGTTCAGGCTTGA
GCACCTTCCTGAGTACAAGGGCAACCGTGATGAGAAGTATGCTCAGAGGACTGAGGAAGAGAAGGCTCTTGACGAGC
AGTTCTTCGAGTACCTGAAGGATGCTTTCGAGCTGTGCAAGACCACCTTTCCTACCTTCACCATTAGGGGCGTTGAG
GCTGATGATATGGCTGCCTACATTGTGAAGCTGATCGGCCACCTTTACGATCACGTGTGGCTTATCTCTACCGATGG
CGATTGGGATACCCTGCTTACCGATAAGGTGAGCAGGTTCTCATTCACTACCAGGCGTGAGTACCACCTGAGGGATA
TGTACGAGCACCATAACGTGGACGACGTTGAGCAGTTCATCTCCCTGAAGGCTATCATGGGTGATCTGGGTGATAAC
ATCAGGGGCGTCGAAGGTATTGGTGCTAAGAGGGGTTACAACATCATCCGTGAGTTCGGCAACGTGCTGGATATTAT
CGATCAGCTGCCTCTGCCTGGCAAGCAGAAGTACATTCAGAACCTGAACGCCAGCGAGGAACTGCTGTTTAGGAACC
TTATTCTGGTGGACCTGCCGACTTACTGCGTTGACGCTATTGCTGCTGTTGGTCAGGATGTGCTCGACAAGTTCACC
AAGGACATCCTTGAGATTGCCGAGCAGTGAGCTT

**pAGT7323 (T5 Exo module: AATG_T5Exo-no stop_TTCG)**

GG-overhang_T5Exo_GG-overhang

AATGTCTAAGAGCTGGGGCAAGTTCATCGAAGAGGAAGAGGCTGAGATGGCCTCTCGGAGGAATCTTATGATCGTGG
ATGGTACTAACCTCGGCTTCCGGTTCAAGCACAACAACAGCAAGAAGCCGTTCGCCTCCTCTTACGTGTCAACCATT
CAGAGCCTGGCCAAGTCTTACTCTGCTAGGACTACCATTGTGCTGGGCGATAAGGGAAAGTCTGTGTTCAGGCTTGA
GCACCTTCCTGAGTACAAGGGCAACCGTGATGAGAAGTATGCTCAGAGGACTGAGGAAGAGAAGGCTCTTGACGAGC
AGTTCTTCGAGTACCTGAAGGATGCTTTCGAGCTGTGCAAGACCACCTTTCCTACCTTCACCATTAGGGGCGTTGAG
GCTGATGATATGGCTGCCTACATTGTGAAGCTGATCGGCCACCTTTACGATCACGTGTGGCTTATCTCTACCGATGG
CGATTGGGATACCCTGCTTACCGATAAGGTGAGCAGGTTCTCATTCACTACCAGGCGTGAGTACCACCTGAGGGATA
TGTACGAGCACCATAACGTGGACGACGTTGAGCAGTTCATCTCCCTGAAGGCTATCATGGGTGATCTGGGTGATAAC
ATCAGGGGCGTCGAAGGTATTGGTGCTAAGAGGGGTTACAACATCATCCGTGAGTTCGGCAACGTGCTGGATATTAT
CGATCAGCTGCCTCTGCCTGGCAAGCAGAAGTACATTCAGAACCTGAACGCCAGCGAGGAACTGCTGTTTAGGAACC
TTATTCTGGTGGACCTGCCGACTTACTGCGTTGACGCTATTGCTGCTGTTGGTCAGGATGTGCTCGACAAGTTCACC
AAGGACATCCTTGAGATTGCCGAGCAGGGTTCG

**pAGT7297 (T7 Exo module: AATG_T7Exo-stop_GCTT)**

GG-overhang_T7Exo_GG-overhang

AATGGCTCTGCTTGACCTGAAGCAGTTCTATGAGCTTAGAGAGGGCTGCGACGATAAGGGTATTCTGGTGATGGATG
GTGACTGGCTTGTGTTCCAAGCTATGTCTGCTGCTGAGTTCGACGCCTCTTGGGAAGAAGAAATTTGGCACCGTTGC
TGCGATCACGCTAAGGCTAGACAGATCCTTGAGGACAGCATCAAGAGCTACGAGACTCGGAAGAAAGCTTGGGCTGG
TGCTCCTATTGTGCTGGCTTTCACCGATTCTGTGAACTGGCGGAAAGAGCTGGTTGACCCTAACTACAAGGCTAACC
GGAAGGCTGTGAAGAAGCCTGTTGGTTACTTCGAGTTCCTGGACGCTCTTTTCGAGCGGGAAGAGTTCTACTGCATC
AGGGAACCTATGCTTGAGGGCGACGATGTGATGGGTGTGATTGCTTCTAACCCTAGCGCTTTCGGTGCTAGGAAGGC
CGTTATTATCAGCTGCGACAAGGACTTCAAGACCATTCCGAACTGCGACTTTCTGTGGTGCACTACCGGTAACATTC
TTACCCAGACCGAAGAGTCTGCTGATTGGTGGCATCTTTTCCAGACCATCAAGGGCGATATCACCGATGGCTACTCT
GGTATTGCTGGTTGGGGTGATACTGCTGAGGACTTCCTTAACAACCCGTTCATTACCGAGCCTAAGACCAGCGTGTT
GAAGTCCGGTAAGAACAAGGGTCAAGAGGTGACCAAGTGGGTGAAGAGGGATCCTGAACCTCATGAGACTCTGTGGG
ACTGCATCAAGTCTATCGGTGCTAAGGCTGGTATGACCGAGGAAGATATCATCAAGCAGGGTCAGATGGCTCGGATC
CTTAGGTTCAACGAGTACAACTTCATCGACAAAGAGATCTACCTCTGGCGGCCTTGAGCTT

**pAGT7324 (T7 Exo module: AATG_T7Exo-no stop_TTCG)**

GG-overhang_T7Exo_GG-overhang

AATGGCTCTGCTTGACCTGAAGCAGTTCTATGAGCTTAGAGAGGGCTGCGACGATAAGGGTATTCTGGTGATGGATG
GTGACTGGCTTGTGTTCCAAGCTATGTCTGCTGCTGAGTTCGACGCCTCTTGGGAAGAAGAAATTTGGCACCGTTGC
TGCGATCACGCTAAGGCTAGACAGATCCTTGAGGACAGCATCAAGAGCTACGAGACTCGGAAGAAAGCTTGGGCTGG
TGCTCCTATTGTGCTGGCTTTCACCGATTCTGTGAACTGGCGGAAAGAGCTGGTTGACCCTAACTACAAGGCTAACC
GGAAGGCTGTGAAGAAGCCTGTTGGTTACTTCGAGTTCCTGGACGCTCTTTTCGAGCGGGAAGAGTTCTACTGCATC

AGGGAACCTATGCTTGAGGGCGACGATGTGATGGGTGTGATTGCTTCTAACCCTAGCGCTTTCGGTGCTAGGAAGGC
CGTTATTATCAGCTGCGACAAGGACTTCAAGACCATTCCGAACTGCGACTTTCTGTGGTGCACTACCGGTAACATTC
TTACCCAGACCGAAGAGTCTGCTGATTGGTGGCATCTTTTCCAGACCATCAAGGGCGATATCACCGATGGCTACTCT
GGTATTGCTGGTTGGGGTGATACTGCTGAGGACTTCCTTAACAACCCGTTCATTACCGAGCCTAAGACCAGCGTGTT
GAAGTCCGGTAAGAACAAGGGTCAAGAGGTGACCAAGTGGGTGAAGAGGGATCCTGAACCTCATGAGACTCTGTGGG
ACTGCATCAAGTCTATCGGTGCTAAGGCTGGTATGACCGAGGAAGATATCATCAAGCAGGGTCAGATGGCTCGGATC
CTTAGGTTCAACGAGTACAACTTCATCGACAAAGAGATCTACCTCTGGCGGCCTGGTTCG

**pAGT7298 (EcExo3 module: AATG_Exo3-stop_GCTT)**
GG-overhang_Exo3_GG-overhang
AATGAAGTTCGTGAGCTTCAACATCAACGGCCTGAGAGCTAGACCTCATCAGCTTGAGGCTATCGTTGAAAAGCACC
AGCCTGATGTGATCGGCCTGCAAGAAACTAAGGTGCACGATGACATGTTCCCGCTTGAGGAAGTTGCTAAGCTGGGC
TACAATGTGTTCTACCACGGTCAAAAGGGACACTACGGTGTGGCTTTGCTGACCAAAGAGACTCCTATCGCTGTTAG
AAGGGGCTTCCCTGGTGATGATGAAGAAGCTCAGCGGAGGATTATCATGGCTGAGATTCCTAGCCTGCTGGGTAATG
TGACCGTGATCAATGGTTACTTTCCGCAGGGTGAGAGCAGGGATCACCCTATTAAGTTTCCAGCTAAGGCCCAGTTC
TACCAGAACCTGCAGAACTACCTTGAGACTGAGCTGAAGAGGGATAACCCGGTGCTTATCATGGGCGACATGAACAT
CTCTCCAACCGATCTGGATATCGGCATCGGCGAAGAGAATAGGAAGAGATGGCTTAGGACCGGCAAGTGCTCATTCT
TGCCTGAAGAACGTGAGTGGATGGACAGGCTTATGTCTTGGGGTCTTGTGGATACTTTCCGGCATGCTAATCCTCAG
ACCGCTGATAGGTTCAGCTGGTTCGATTACAGGTCCAAGGGCTTCGATGATAACAGAGGCCTGAGGATTGATCTGCT
GCTTGCTTCTCAACCTCTTGCTGAGTGTTGCGTTGAGACTGGCATCGATTACGAGATCCGGTCCATGGAAAAGCCTT
CAGATCATGCTCCTGTGTGGGCTACTTTCAGAAGGTGAGCTT

**pAGT7325 (EcExo3 module: AATG_Exo3-no stop_TTCG)**
GG-overhang_Exo3_GG-overhang
AATGAAGTTCGTGAGCTTCAACATCAACGGCCTGAGAGCTAGACCTCATCAGCTTGAGGCTATCGTTGAAAAGCACC
AGCCTGATGTGATCGGCCTGCAAGAAACTAAGGTGCACGATGACATGTTCCCGCTTGAGGAAGTTGCTAAGCTGGGC
TACAATGTGTTCTACCACGGTCAAAAGGGACACTACGGTGTGGCTTTGCTGACCAAAGAGACTCCTATCGCTGTTAG
AAGGGGCTTCCCTGGTGATGATGAAGAAGCTCAGCGGAGGATTATCATGGCTGAGATTCCTAGCCTGCTGGGTAATG
TGACCGTGATCAATGGTTACTTTCCGCAGGGTGAGAGCAGGGATCACCCTATTAAGTTTCCAGCTAAGGCCCAGTTC
TACCAGAACCTGCAGAACTACCTTGAGACTGAGCTGAAGAGGGATAACCCGGTGCTTATCATGGGCGACATGAACAT
CTCTCCAACCGATCTGGATATCGGCATCGGCGAAGAGAATAGGAAGAGATGGCTTAGGACCGGCAAGTGCTCATTCT
TGCCTGAAGAACGTGAGTGGATGGACAGGCTTATGTCTTGGGGTCTTGTGGATACTTTCCGGCATGCTAATCCTCAG
ACCGCTGATAGGTTCAGCTGGTTCGATTACAGGTCCAAGGGCTTCGATGATAACAGAGGCCTGAGGATTGATCTGCT
GCTTGCTTCTCAACCTCTTGCTGAGTGTTGCGTTGAGACTGGCATCGATTACGAGATCCGGTCCATGGAAAAGCCTT
CAGATCATGCTCCTGTGTGGGCTACTTTCAGAAGGGGTTCG

**pAGT7299 (huTREX1 module: AATG_TREX1-stop_GCTT)**
GG-overhang_TREX1_GG-overhang
AATGCAGACCCTGATCTTCTTCGATATGGAAGCTACCGGCCTGCCATTCTCTCAGCCTAAGGTTACAGAGCTTTGCC
TTCTGGCTGTTCACAGATGCGCTCTTGAATCTCCTCCAACTTCTCAAGGTCCTCCTCCTACTGTTCCTCCACCTCCT
AGAGTTGTGGATAAGCTGTCTCTTTGTGTGGCTCCTGGTAAGGCTTGTTCTCCTGCTGCTTCTGAGATTACCGGTCT
TTCTACTGCTGTGCTTGCTGCTCATGGTAGGCAGTGCTTCGATGATAACCTTGCTAACCTGCTGCTGGCTTTCCTTA
GAAGGCAACCTCAACCTTGGTGCCTTGTGGCTCATAACGGTGATAGGTACGATTTCCCACTTCTGCAGGCTGAGCTT
GCTATGCTTGGTCTTACCTCTGCTCTGGATGGTGCTTTCTGCGTGGACTCTATTACCGCTCTTAAGGCTCTTGAGCG
GGCTTCTTCTCCTTCTGAACATGGTCCTCGGAAGTCCTACTCTCTGGGTTCTATCTACACCAGGCTGTACGGTCAGT
CTCCTCCTGATTCTCATACTGCTGAGGGTGATGTGCTGGCTCTGCTTTCTATTTGTCAATGGCGTCCTCAGGCTCTG
CTGAGATGGGTTGACGCTCATGCTAGACCTTTCGGAACCATCAGGCCTATGTACGGTGTTACTGCTTCTGCTAGGAC
TAAGCCTAGGCCTTCTGCTGTGACTACTACTGCTCATCTGGCTACTACCCGGAACACCTCTCCATCTCTTGGTGAGT
CTAGGGGTACTAAGGATCTGCCTCCTGTTAAGGATCCTGGCGCTCTTTCTAGAGAGGGTTTGCTTGCTCCTCTTGGT
CTGCTTGCTATTCTTACCCTTGCTGTGGCTACCCTTTACGGTTTGTCTCTTGCTACTCCTGGTGAGTGAGCTT

**pAGT7326 (huTREX1 module: AATG_TREX1-no stop_TTCG)**
GG-overhang_TREX1_GG-overhang
AATGCAGACCCTGATCTTCTTCGATATGGAAGCTACCGGCCTGCCATTCTCTCAGCCTAAGGTTACAGAGCTTTGCC
TTCTGGCTGTTCACAGATGCGCTCTTGAATCTCCTCCAACTTCTCAAGGTCCTCCTCCTACTGTTCCTCCACCTCCT
AGAGTTGTGGATAAGCTGTCTCTTTGTGTGGCTCCTGGTAAGGCTTGTTCTCCTGCTGCTTCTGAGATTACCGGTCT
TTCTACTGCTGTGCTTGCTGCTCATGGTAGGCAGTGCTTCGATGATAACCTTGCTAACCTGCTGCTGGCTTTCCTTA
GAAGGCAACCTCAACCTTGGTGCCTTGTGGCTCATAACGGTGATAGGTACGATTTCCCACTTCTGCAGGCTGAGCTT

GCTATGCTTGGTCTTACCTCTGCTCTGGATGGTGCTTTCTGCGTGGACTCTATTACCGCTCTTAAGGCTCTTGAGCG
GGCTTCTTCTCCTTCTGAACATGGTCCTCGGAAGTCCTACTCTCTGGGTTCTATCTACACCAGGCTGTACGGTCAGT
CTCCTCCTGATTCTCATACTGCTGAGGGTGATGTGCTGGCTCTGCTTTCTATTTGTCAATGGCGTCCTCAGGCTCTG
CTGAGATGGGTTGACGCTCATGCTAGACCTTTCGGAACCATCAGGCCTATGTACGGTGTTACTGCTTCTGCTAGGAC
TAAGCCTAGGCCTTCTGCTGTGACTACTACTGCTCATCTGGCTACTACCCGGAACACCTCTCCATCTCTTGGTGAGT
CTAGGGGTACTAAGGATCTGCCTCCTGTTAAGGATCCTGGCGCTCTTTCTAGAGAGGGTTTGCTTGCTCCTCTTGGT
CTGCTTGCTATTCTTACCCTTGCTGTGGCTACCCTTTACGGTTTGTCTCTTGCTACTCCTGGTGAGGGTTCG

**pAGT7846 (UL12-2 (AQZ56199.2; Human alphaherpesvirus 2) module A: AATG_UL12-2-A_GCAG)**
GG-overhang_UL12-2-A_GG-overhang
AATGGCTGCTGCTGCTACTCCTGGTGCTAAGAGGCCTGCTGATCCTGCTAGAGATCCTGACTCTCCACCAAAGAGGC
CAAGGCCTAACTCTCTTGATCTTGCTACTGTGTTCGGTCCTAGACCTGCTCCTCCTCATCCTACTTCTCCAGGTGCT
CCTGGTTCTCATTCTCCTCAATCTCCACCTAGAGGTCAGCCTGATGGTGGTGCTCCAGGTGAAAAGGCTAGACCAGC
TTCTCCTGCTCTTAGCGAAGCTTCTTCTGGTCCTCCTACTCCTGATATCCCTCTTTCTCCTGGTGGCGCTCATGCTA
TTGATCCTGATTGCTCTCCTGGACCTCCTGATCCAGATCCTATGTGGTCTGCTTCTGCTATCCCTAACGCTCTGCCT
CCTCATATTCTGGCTGAGACTTTCGAGAGGCACCTTAGGGGTTTGCTTAGGGGTGTTAGATCTCCTCTTGCTATCGG
TCCTCTTTGGGCTAGACTTGACTACCTTTGCTCTCTGGTGGTGTCTCTTGAAGCTGCTGGTATGGTGGATAGAGGTC
TTGGTAGACACCTTTGGAGGCTTACTAGAAGGGCTCCTCCATCTGCTGCTGAAGCTGTTGCTCCTAGACCACTTATG
GGATTCTACGAGGCTGCTACCCAGAACCAGGCTGATTGTCAACTTTGGGCTCTGCTTAGAAGGGGTCTTACTACCGC
TTCTACTCTTAGATGGGGTGCTCAGGGACCTTGCTTTTCATCTCAGTGGCTTACCCACAACGCTAGCCTTAGGCTTG
ATGCTCAGTCAAGCGCTGTGATGTTCGGTAGAGTGAATGAGCCTACCGCTAGGAACCTTCTGTTCAGGTATTGCGTT
GGTAGGGCTGATGCTGGCGTTAACGATGATGCAGATGCTGGCAG

**pAGT8430 (UL12-2 (AQZ56199.2; Human alphaherpesvirus 2) module B1: GCAG_UL12-2-B1-stop_GCTT)**
GG-overhang_UL12-2-B1_GG-overhang
GCAGATTCGTGTTCCATCAGCCAGGTGATCTGGCCGAAGAGAATGTTCATGCTTGCGGTGTGCTTATGGATGGTCAC
ACTGGAATGGTGGGCGCTTCTCTTGATATTCTTGTGTGCCCTAGGGATCCTCACGGTTATCTTGCTCCAGCTCCTCA
AACTCCTCTGGCCTTTTATGAGGTTAAGTGCCGGGCTAAGTACGCTTTCGATCCAGCTGACCCTGGCGCTCCTGCTG
CTTCAGCTTATGAAGATCTGATGGCTAGGCGTAGCCCTGAGGCTTTTAGGGCTTTCATTCGGAGCATCCCTAATCCG
GGTGTGAGATATTTTGCTCCTGGAAGAGTTCCTGGGCCTGAAGAGGCTCTTGTTACCCAAGATAGAGACTGGCTGGA
TTCTAGAGCAGCTGGTGAGAAGAGAAGATGCTCTGCTCCTGATAGGGCACTCGTTGAGCTTAATTCTGGCGTGGTGT
CTGAGGTGTTGCTTTTCGGTGTTCCTGATCTCGAGCGGAGGACTATTTCTCCTGTTGCTTGGTCATCTGGTGAGCTT
GTGAGAAGGGAACCGATCTTCGCTAATCCTAGGCACCCTAACTTCAAGCAGATTCTGGTGCAGGGTTACGTGCTGGA
TTCACACTTTCCAGATTGCCCTCTTCAGCCTCACCTGGTGACTTTTCTTGGTAGGCATAGAGCTGGTGCTGAGGAAG
GTGTTACTTTCAGGCTTGAGGATGGTAGAGGTGCACCAGCTGGAAGAGGTGGTGCCCCTGGTCCTGCTAAGGCTTCT
ATTCTTCCTGATCAAGCTGTGCCGATCGCTCTGATTATTACCCCTGTTAGAGTTGAGCCGGGCATCTACAGAGATAT
CCGTAGGAATTCTCGGCTGGCTTTCGATGATACCCTTGCTAAGCTTTGGGCCTCTAGATCTCCAGGTAGAGGTCCTG
CTGCAGCTGATACAACTTCTTCTTCACCTACTGCCGGCCGTTCTTCTAGATGAGCTT

**pAGT7847 (UL12-2 (AQZ56199.2; Human alphaherpesvirus 2) module B2: GCAG_UL12-2-B2-no stop_TTCG)**
GG-overhang_UL12-2-B2_GG-overhang
GCAGATTCGTGTTCCATCAGCCAGGTGATCTGGCCGAAGAGAATGTTCATGCTTGCGGTGTGCTTATGGATGGTCAC
ACTGGAATGGTGGGCGCTTCTCTTGATATTCTTGTGTGCCCTAGGGATCCTCACGGTTATCTTGCTCCAGCTCCTCA
AACTCCTCTGGCCTTTTATGAGGTTAAGTGCCGGGCTAAGTACGCTTTCGATCCAGCTGACCCTGGCGCTCCTGCTG
CTTCAGCTTATGAAGATCTGATGGCTAGGCGTAGCCCTGAGGCTTTTAGGGCTTTCATTCGGAGCATCCCTAATCCG
GGTGTGAGATATTTTGCTCCTGGAAGAGTTCCTGGGCCTGAAGAGGCTCTTGTTACCCAAGATAGAGACTGGCTGGA
TTCTAGAGCAGCTGGTGAGAAGAGAAGATGCTCTGCTCCTGATAGGGCACTCGTTGAGCTTAATTCTGGCGTGGTGT
CTGAGGTGTTGCTTTTCGGTGTTCCTGATCTCGAGCGGAGGACTATTTCTCCTGTTGCTTGGTCATCTGGTGAGCTT
GTGAGAAGGGAACCGATCTTCGCTAATCCTAGGCACCCTAACTTCAAGCAGATTCTGGTGCAGGGTTACGTGCTGGA
TTCACACTTTCCAGATTGCCCTCTTCAGCCTCACCTGGTGACTTTTCTTGGTAGGCATAGAGCTGGTGCTGAGGAAG
GTGTTACTTTCAGGCTTGAGGATGGTAGAGGTGCACCAGCTGGAAGAGGTGGTGCCCCTGGTCCTGCTAAGGCTTCT
ATTCTTCCTGATCAAGCTGTGCCGATCGCTCTGATTATTACCCCTGTTAGAGTTGAGCCGGGCATCTACAGAGATAT
CCGTAGGAATTCTCGGCTGGCTTTCGATGATACCCTTGCTAAGCTTTGGGCCTCTAGATCTCCAGGTAGAGGTCCTG
CTGCAGCTGATACAACTTCTTCTTCACCTACTGCCGGCCGTTCTTCTAGAGGTTCG

**pAGT9046 (PapE (AHM96060.1_FL; Papiine alphaherpesvirus 2) module: AATG_PapE-no stop_TTCG)**

GG-overhang_PapE_GG-overhang

AATGCAGACTACTACCCCTGTGGATCCTCCATCTTCTCGGTCTGAAAACAAGAGGCCTCCTGCTCCTGCTGGTGATG
AAGGTGCTGGTCCTGGTAGAGGTCTTGTTGATCCTGCTAGACCTCCTAAGAGGCCAAGGCCTGATTCTCTTCCTCTT
GCTGCTGTGTGTAGACCTGCTACTCCTCCATCACCTGGTAGACCAGAAACTCCTCCTACTCCTGATCTGCCTTTGTC
TCCTAGAGGAACCCATGGTATTGCAGCTCCAGCTGGCGAACCTGAACCTGGTTCTCCTAGTCTGCTTGAGAACTATG
TTCCTCCAGCTCCTGATGCTGGTGACGCTGGTTCTACTCCAGAGCCTGGTTGGTCTGCTGTTGCTATTCCAGATGCT
CTGCCTTCTCATGTGCTGGCTGAGACTTTTGAGAGGCACCTTTGCGGTCTTTTGAGGGGTGTTAGAAGGCCTTTGGA
TGTTGAGCCTCTTAGAGCTAGGCTGGGCTACCTTTTCTCTCTTGCTACTGCTCTTGAAGAGGCTGGCATGGTGGATA
GAGGTGTTGGTGGTCATCTTCTGAGGCTGTCTAGAAGGGCTGCTGCTGCAGATCCTAGACCTCTTATGGCTTTTTTC
GAGGCCGCTACTCAGAACCAGGCTGAATCTCAACTTTGGGGTCTGCTTAGAAGGGGTCTTACTACCGCTTCTACCCT
TAAGTGGGGACCTAGAGGACCTTGCTTTAGCCCTAGATGGCTGAAGAACAACGGTGATCCTAGGCTGGACTTCCAGT
CATCTGCTGTTATGTTCGGTAGGACCAATGAGCCTGCTGCTAGGGCTCTTTTGTTCAGGTATTGCGTTGGCAGGGCT
GATGATAGAGATGCTGAAGGTGATGATGCTGGCAGGCGTTTTGTTTTTTGCGAGCCTGGTGATGCTCCTGCAGCTTC
TGTTCATGCTTGCGGTGTTCTTGTGGATGCTCACACTGGTATGGTGGGCGCTTCTCTTGATATTCTTGTGTGCCCTA
GAGATAGGCACGGCTGCCTTAATCCTGCTCCAGGTACTCCTCTTAGGTTCTACGAGGTTAAGTGCCGGGCTAAGTAC
GCTTTCGATCCTGCTGATGCAGGCGATCCTGTTGTTGCTGCTCATAGAAGGCTTGTGGCTAGGCGTTCTCCATCTGA
TTTCAGGGCTTTCTTGCGGTCTATTGCTAGGCCTGGTGTGAGATACTTTGCTCCTGGAAGAGTTCCTGGTCCTGAGG
AAGCTCTTGTTTCTGATCATGCTGTGTGGGCTGATGCAAGAGCCGGTGATGAGAAGAGAAGGTGCTCTGCTTTGGAC
AGGGCTCTTGTGGGTCTTAATTCTGGTGTGGCTTCCGATGTGCTGCTTTTCGGAGATCCTGATCCAGAGAGAAGGAC
CGTTTCTCCTTTGGCTTGGTCATCTGGTGCTCTTGTGCACAGGGAACCTATCTTCGCTAATCCTAGGCACCCGAACT
TCAAGCAGATTCTGGTTCAGGCTTACGTGCTGGCTTCTCATTTTCCTGAATGCCCTCTTCATCCTCACCTCGTGACT
TTCATTGGTAGACATAGGACCCCTGATGAGGAAGGACTTTCTCTGAGACTTGAGGACGCTCCAGCTTCTGCTCCAGC
AGCTGTTAGGGCTGCAGCTGGTGCTTCTATTCTTCCAGATCAGGCTGTTCCTGTGGCTCTGATTATTACCCCTGTTA
GGGTTGACGCTGCCATCTACGATCTTATCCGTAGGAACTCTAGGCTCGCCTTCGATGAGACTCTTGCTAGACTTTGG
GCTTCTAGGGCTCCTGCTTCTGATCCTGCCGCTGCTGGCGAAACTTCTTCG

**pAGT9047 (PiE (BBM13184.1_FL; Pteropus lylei-associated alphaherpesvirus) module: AATG_PiE-no stop_TTCG)**

GG-overhang_PiE_GG-overhang

AATGACCTCTACCAGCAGCTCTAGTCCTCTTCATCCTCCTAGTCCTCAGAAGCGGAAGTCTCTTTCTGCTGATGGTG
TTGCTGGTCTTGCTACTCCTACTAAGAGAGCTAGGCCTCACTCTCTTCCACCTCTTGTGCTTCATTGGTCACCTCCA
TCTCCATCTTTGCTGCCTCAGGACGGGACTTTCATCTTCCCATCTGATGATACCAAGACCAGGGCTGCTGAAGAAAC
TGGTCCTCCTGCTCCTCAATCTCAGAATGCTCCTGTTAGCCCTCTGGGCGATAAGTTTTCTCCTGCTTGTGCTCCTA
CCAGCGGTCTGATTTGTTCCGATTCTGAAGAGACTGAGGACCTGGTCGAGTCTACTCAGGTTTTGTCATCTGAGGCT
GCTACCCCTCTGTCTAGAGTTGAGGCTTGTGATCTTCCTCCTCCGCTTATGTGGTCCGCTACCTCTATTCCTAATGC
TCTGCCTCCTGAGATCTTCACCAAGACCTTCGCTAAGTACCTGCGGAAGCTTCTGATCGGTATTGATCACCCTCTGG
ACATTGAGCCTCTTCAGGCTAGGCTTGGTTACCTGTACTCTCTGATGAGGGCTCTTGAAGGTGGTGGTATGCTTTCT
GAGGGCCTGTCTAGATACCTGATCTGCCAATCTAGACCTCAGGCCAGCAGATCTAAGTTGCCTAGACCTGGTTTGCC
TGTGGTGAACCCTAAGCCACTGATGAGATTTTTCGAGGCCGCTACTCAGTCTCAGGGCGATTCTCAACTTTGGGCTC
TTTTGAGAAGGGGCCTTGCTACCGCTACTACTCTTAAGTGGGGTTCTCAGGGTCCAGCTTTTGCTCCTCAGTGGTTG
GATGGTGTGGTTGATCAATCTGCTGGTGGTAAGGGTGCTGCTATTGCTTTCGGTAGGATTAACGAGCTTACCGCCAG
GACCATTCTGTTCAGGTATTGCGTTGGTAGGGCTGATCATACCGCTGATGCTGATCCTGAGGAACGGTTCATTTTCC
ACCAGCCTGATGACATGGCCGAAGAGAATGTTCATACCTGCGGTGTGCTTATGGATACCCACACTGGTATGGTGGGC
GCTTCTCTTGATATTCTTGTGTGCCCTAGAGATCAGCACGGTTGCTTGTCTCCACCTCCTAAGTACCCTCTTGCTTT
CTACGAGGTTAAGTGCCGGGCTAAGTACGCTTTCGATCCTATGGATCTTCAGAGCCCAACCACCTTGGCTTACAACC
AGCTGATGGATAGAAGGTCCCCTGCTGCTTTCAGGCGGTTCATGCACTCTATTTCTAAGCCTGGCGTGCAGTTCATC
AGCCACGGTAATTTTCCAGGACCTGAGGAAGCTCTGGTCACCACTTCTAGTCTTTGGGATCAGTCATCTGGTGCCCC
GACCAAGAAAAGAAGATGTCCTGCTGCTGAGCAGGCTCTGGTGAAGCTTAACAAGTCTGTGACCAGCAGCATCCTTC
TTTTCGGCACTCCTGATCTTGAGCAGAGGACTATTACACCTGTGAGGTGGGATTCTGGCTGCCTGTTTTACAGAGAG
GCTCTGTTCGCTAATCCTCGGCATCCTAACTTCAGGCAGATTCTGGTTCAGGCTTACGTGCTGACTTCTCACTTCCC
TGATTCTCCAGTGTCTCCTCACCTTGTGACCTTCATTGGAAGGCAGAGAACTGTGGCTGAAGAGGGTGTGAATTTCT
GGCTCGAAACCCCTTCTCCTTCTGTTGCTTGCCCTCCTAATCACGATCCTTCTCCACCACCTGTGTCTAACAGGGCT
TCTATTGCTGCCGATCAGGCTATTCCTGTGGCTGTGATTATTACCCCAGTGAGGCTTGATGTGGCCGTGTATAAGGT
GTTGCAGAGGAACTCTAGGCTGGCTTTCGATGCTACTCTTGCTCAGTTGTGGGCTTCTAGGACTCCTAAGTCTGTTC
TCGCTGCTGACGAGACTAGCTCATCTCCTACTACTGAGTCTCCTTCG

**pAGT8098 (AB4P (YP_053046.1; Equid alphaherpesvirus 1) module-A: AATG_AB4P-A_GACC)**

GG-overhang_AB4P-A_GG-overhang

AATGGACTCTTCTCCGGTGACTTATTCTGGTGAGCCTCCTTACAAGCTGCGGAGGCTTTCTCCTTCTTACCCGTACG
TTAGCAAGCTGAGAGAGAGGTGCGCTAGCAAGATTGAGACTCTGTCTGAGGGTAGCGCTAGGGATTCTCTGGAAGAG
GAAGATGTGTCTGAGGCTATGGCTACCGGTGCTTTTCTTGCTACCAGGCTTTACCTTCCTAGCGTGTTGCCTCAGAG
GATTACTACCCTGACCTTCCTGGACCACTTCAAGAAGTCTAGGCCTCTGCCTAACAGCGACAAGAGGCTTAACCCTA
TCTTCTACCGGCTGGCTTACATCAGGGATCTTGTGGGTGAAATGGAACTCGAGGGTATCGTTGAGAGGGGTACTGCT
TCTAGACTGCTTGGTGCTTCTTCTCCTGCTGGTTTCGTTGCTGGTACTTACACCCATGCTAGGGACCTGAGCAAGAC
TATGTCTCTTGCTTCTGTGAGGGATGCTGTGCTTGCTATTGAGGCTCAGACTAGGGATCAGTCTGAGTCTCAGCTTT
GGGCTCTTTTGAGAAGGGGTCTTGCTACTGCCTCTACCATGAAGTGGGGTGCTCTTGGTCCTCAGTATCATCCTCAG
TGGTGCGAGGTGTCAACTAACGCTAAGGGTATCCCTAACAACCCGGCTCTGCAATTCGGTCAGACTAATGAGAGGAC
CGCTCGGAGCCTTATCTCTGCTCTTTATGTTGCTAGGTCCGAGGCTGCTACCCCTGATCTTCTTGTTGATCCTGGTT
GCGGTCAGTGCTTCGTGTTTGATGAGTCTGCTTCAGTGCCTGGGGATGCTTATGCTTGCGGTCTTTTGATGGATGCT
AGGACC

**pAGT8098 (AB4P (YP_053046.1; Equid alphaherpesvirus 1) module-B: GACC_AB4P-B-no stop_TTCG)**

GG-overhang_AB4P-B_GG-overhang

GACCGGTGTTGTGGGCGCTTCTCTTGATATGCTTGTGTGCGATAGGGACCCTTCTGGTGTTCTTTCTCCACATTCTA
CCCAGACTACCCTGGACTTCTTCGAGATTAAGTGCCGGGCTAAGTACCTGTTCGATCCGGATTTGTTCTCTCCTGTG
GCTACCGCTTACGCCAACCTTCTTAAACATAGGACCGCTGTGTGCCTGAGGAAGTTCCTGCGGTCTATTAAGAACCC
TGCCGTCGAGTACTTCGCTCCTACTTCTGTTCCTGGTGCTACCGAGGCTCTTATTACCTGCAACTCTTCTTGGAAGC
CGCGTGAGGTGAACGAGACTAATAGAAGGTGCGGCGATTTCGACAGGGACCACATTGCTCTTAACCTGGACGCTTCT
TCTGACGTGTGGTTGTTCTCTGAGCCTGACCTTGAGTCTGAGACTATTACTCCTGCTAGGTGGGATACCGGTGAGCT
TGCTCTTTCTGTGCCTGTGTTTGCTAATCCTCGGCACCCTAACTTCAAGCAGATTCTGGTTCAGGCCTACGTGCTGT
CTGGTCATTTCCCTGATCATCAGCTGAGGCCTTTCTTGGTGACCTTCATTGGTAGACATAGGAAGAGGTGCGAAGAG
GGTAAGACCTTCACCATTTGCGATAGGCCTGAGGGCTCTCCATACAACCTTAATGAGGTGGTGCACAGCTCTTGCGC
TATTCCTATTCTTCTGTTCGTGACCCCTGTGATCGTGGATAGAGAAGGATGCTGGGAAGATATCGAGATCGAGTCTC
TGACCGCCTTCAACAAGACCGCTGATGCTATCTGGGATAGCGACTCTCCAGCTGATGTTTCTGAGCCAACCTCTTCG

**pAGT8096 (MD5 (YP_001033940.1; Gallid alphaherpesvirus 2) module-A: AATG_MD5-A_GGTG)**

GG-overhang_MD5-A_GG-overhang

AATGGAACTGGGCACTAAGTTCCCGCTGTCTAAGTCTTGCAAGGACGAGTCTAGAAAGCGGAAGAGGGGTATCACCA
TCGATTGCGATTCTCAGATCCTGGTGGGTGACGAGCAGTCTAATTCTACCAAGACCAAGCCGTACGACGAGATCTGC
GAGAACATTGTGCCTAACTACACCTTCGGCAACTACATCCTGCAAAAGATCGACCCTAACGACTGCAGGCATTCTCT
TCACCCACTTTACCACCGGCTGTTCTACATTGCTGACGTGATCAAGCAAGGGATCTCTGAGGGTTCACTGCTCGAGA
ACAAGTACAGCTACATCCTCGAGACTGAGCACATCCTGTTGGACGAGAGCAGGATCAACAACCTGTCTCCTTCTATC
CACGCTTCCAGGTGGTGTAAGATGGTTGAGTCTCTTACCAGGCTGCAGGCTAACTCTGAGCTTTGGCATATTTTCAG
ACAGTGCCTGCTGACCGCTTCTTCTGTTAAGTGGTCACCTAACGGCACCATCAACACCGCTGGTCTGATTACCAACG
ATCTGCCTTCTAGAGGTCAAACCGAGTCTATTCTGTTCGGCACTAGGAACGAGAGCCTGGCTAAGTCTCTTATCGCT
GCTCTTTGCGTGAGCCAGTCATCTGTGAGGACCATCGATAACAGCGACAAGAAGAACGAGTTCGACAACACCACCAC
CGGCATTCTGGATATCGAGAAGTACTCTTGCGGCCTGATGATCGATATTCGGACCGGTATGCTTGGCGCTTCTCTGG
ATATGGTG

**pAGT8097 (MD5 (YP_001033940.1; Gallid alphaherpesvirus 2) module-B: GGTG_MD5-B-no stop_TTCG)**

GG-overhang_MD5-B_GG-overhang

GGTGATGTGCAATAGGAACAGGCACGGTATTCTTGCTCCATGCCTGACCGATAACAACATCGAGACTTACGAGATCA
AGTGCCGGTTCAAGTACGCTTTCTGTCCCGAGATGAGGTCCGAGCTTTCTCAGTGCTATGAGAGGCTTATGGCCACT
AAGACTGTGCAGTGGTTTCGGCGGTTCCTTTACACCATTGATTGCCCTTGCGTGGACTACTTCAGGCCAGATAATTA
CCCTCGGGCTAAAGAGGCTCTGATCACCTCTGATGACGATTGGAAGGTGGGACACTCTGCTTATCATGCTGCTCAGT
CCCGGATTAAGTGCAATGAGTTCGAGATGCACCACTTGACCCTGAACAAGAACATGTCCTCTCGTGTGTGGCTTTTC
GGTGAGCCTGATCTTCAGACCAACAGCATCTACCCTCTGCTTTGGAATACCGGTGAGAGGGTGCTGTCTATCCCTAT
CTTTGCTAATCCGCGGCACCAGAACTTCAAGCAGATTTTCCTGCAGAGCTACGTGGCCTCTGGTTACTTCGGTAACA
GAAAGATCGTGCCGTTCCTGGCTACCTTCATTGGTAGGCATAGACGGCAGACTGAGCTTGGCAGATGCTTCTCTCTT
TTCGTGGATGACACCGAGGCTAGCGAGGTTGTGTATGAGATTACTCCTGAGCAGGCTATCCCCGTGATTCTGATTAT
TACCCCGGTGATCATTGACAACACCTTCTACGTGGGCATCGAAGAGTCTGGCTACAGAGCTTTTGGTGAGTTGGTGG
ATCACCTGTGGGCTAAGCAGTGCAGAATTTCG

**pAGT8094 (Dumas (NP_040170.1; Human alphaherpesvirus 3) module-A: AATG_Dumas-A_CCAC)**

GG-overhang_Dumas-A_GG-overhang

AATGGCTAGGTCTGGTCTGGATAGGATCGATATCTCTCCACAGCCTGCCAAGAAAATCGCTAGAGTTGGTGGACTTC
AGCACCCGTTCGTTAAGACCGATATCAACACCATCAACGTCGAGCACCACTTCATCGATACCCTGCAAAAGACCTCT
CCGAACATGGATTGCAGGGGTATGACCGCTGGTATCTTCATTAGGCTGAGCCACATGTACAAGATCCTGACCACTCT
CGAGTCCCCAAACGATGTGACTTACACTACTCCTGGCTCTACCAACGCTCTGTTCTTCAAGACTTCTACCCAGCCTC
AAGAGCCTAGGCCTGAAGAACTTGCTTCTAAGCTGACCCAGGACGATATCAAGAGGATCCTTCTGACCATCGAGAGC
GAGACTAGAGGTCAGGGTGATAACGCTATTTGGACCCTTCTGCGGAGGAACCTTATTACCGCTTCTACCCTTAAGTG
GTCCGTGTCTGGTCCTGTTATTCCTCCTCAGTGGTTCTACCATCACAACACCACTGATACCTACGGGGATGCTGCTG
CTATGGCTTTCGGTAAGACTAATGAGCCTGCTGCTAGGGCAATCGTTGAGGCTCTTTTCATCGACCCTGCTGACATT
AGGACCCCTGATCATCTTACACCTGAGGCCACCACCAAGTTTTTCAACTTCGATATGCTGAATACCAAGTCTCCCTC
TCTTCTTGTGGGCACCCCTAGGATTGGTACTTATGAGTGCGGTCTGCTGATCGATGTGAGGACCGGTCTTATTGGTG
CTTCTCTTGATGTGCTGGTGTGCGATAGGGATCCAC

**pAGT8095 (Dumas (NP_040170.1; Human alphaherpesvirus 3) module-B: CCAC_Dumas-B-no stop_TTCG)**

GG-overhang_Dumas-B_GG-overhang

CCACTTACCGGTACTCTTAACCCTCATCCTGCTGAGACTGACATCTCATTCTTCGAGATCAAGTGCCGGGCCAAGTA
CCTGTTCGATCCTGATGATAAGAACAACCCGCTTGGCAGGACTTACACCACCTTGATTAACAGGCCTACCATGGCTA
ACCTGCGGGATTTCCTGTACACCATTAAGAACCCGTGCGTGAGCTTCTTCGGTCCTTCTGCTAATCCTTCTACCAGA
GAGGCTCTGATCACCGATCATGTTGAGTGGAAGAGGCTGGGCTTTAAAGGTGGTAGGGCTCTTACTGAGCTTGACGC
TCATCACCTTGGTCTGAACCGGACCATTTCTTCTAGAGTGTGGGTGTTCAACGACCCGGATATTCAGAAGGGAACCA
TCACCACTATTGCTTGGGCTACTGGTGATACCGCTCTGCAGATTCCTGTGTTCGCTAATCCTAGGCACGCCAACTTC
AAGCAGATTGCCGTTCAGACCTACGTGCTGTCTGGTTACTTCCCGGCTCTTAAGCTTAGGCCTTTCCTGGTGACTTT
CATCGGTAGAGTTAGAAGGCCACATGAGGTTGGAGTTCCTCTTAGAGTGGATACTCAGGCTGCTGCCATCTACGAGT
ACAACTGGCCTACTATTCCTCCACATTGCGCTGTGCCTGTGATTGCTGTGCTTACCCCTATTGAAGTGGATGTGCCT
AGGGTGACCCAGATCTTGAAGGATACTGGCAACAACGCCATCACCAGCGCTCTTAGATCTCTGAGGTGGGATAATCT
TCACCCTGCCGTCGAAGAGGAATCTGTGGATTGTGCTAACGGAACCACCTCTTTGTTGAGGGCTACTGAGAAGCCTC
TTCTTTCG

**pAGT7848 (BGLF5 (AHA36455.1; Human gammaherpesvirus 4) module-A: AATG_BGLF5-A_GCTC)**

GG-overhang_BGLF5-A_GG-overhang

AATGGCTGATGTGGATGAGCTTGAGGACCCTATGGAAGAGATGACCTCTTACACCTTCGCTCGGTTCCTTAGGTCAC
CTGAGACTGAGGCTTTCGTGAGGAATCTTGATAGGCCTCCTCAGATGCCTGCCATGAGATACGTTTACCTTTACTGC
CTGTGCAAGCAGATCCAAGAGTTCTCTGGTGAGACTGGCTTCTGCGACTTCGTGTCTAGTCTTGTGCAAGAGAACGA
CAGCAAGGATGGGCCTTCTCTGAAGTCTATCTACTGGGGACTTCAAGAGGCTACCGATGAGCAGAGAACTGTGCTTT
GCTCCTACGTCGAGTCTATGACTAGAGGCCAGTCTGAGAACCTGATGTGGGATATTCTGCGGAACGGCATTATCAGC
TCCAGCAAGCTTCTGAGCACCATCAAGAACGGTCCTACCAAGGTTTTCGAGCCTGCTCCTATTAGCACCAACCACTA
CTTTGGTGGTCCTGTGGCTTTTGGTCTTAGGTGCGAGGATACCGTGAAGGACATTGTCTGCAAGCTGATCTGCGGTG
ACGCTTCAGCTAATAGGCAGTTCGGCTTCATGATCAGCCCTACCGATGGTATCTTCGGTGTGTCTCTTGATCTGTGC
GTGAACGTTGAAAGCCAGGGCGATTTCATCCTGTTCACCGATAGGTCTTGCATCTACGAGATCAAGTGCCGGTTCAA
GTACCTGTTCAGCAAGAGCGAGTTCGACCCTATCTACCCTTCTTACACCGCTC

**pAGT7849 (BGLF5 (AHA36455.1; Human gammaherpesvirus 4) module-B: GCTC_BGLF5-B-no stop_TTCG)**

GG-overhang_BGLF5-B_GG-overhang

GCTCTGTATAAGCGGCCCTGCAAGAGATCCTTCATCCGGTTCATTAACTCTATCGCTCGGCCGACCGTTGAGTACGT
TCCAGATGGTAGATTGCCTAGCGAGGGCGATTACCTTCTTACTCAGGATGAGGCTTGGAACCTGAAGGATGTGAGGA
AGAGAAAGCTTGGTCCTGGTCACGATCTGGTGGCTGATTCTCTTGCTGCTAACAGGGGTGTTGAGAGCATGCTTTAC
GTGATGACCGATCCTTCTGAGAACGCTGGTAGGATCGGAATCAAGGATAGGGTGCCAGTGAACATCTTCATCAACCC
GAGGCACAACTACTTCTACCAGGTGCTGCTTCAGTACAAGATCGTGGGAGATTACGTGAGGCATAGCGGTGGTGGTA
AGCCTGGTAGAGATTGCTCTCCTAGGGTGAACATTGTGACCGCATTCTTCAGAAAGAGGTCCCCACTTGATCCTGCT
ACCTGTACCTTGGGTTCTGATCTTCTGCTGGACGCCTCTGTTGAGATTCCTGTTGCTGTTCTTGTGACCCCTGTGGT
GCTTCCTGATAGCGTGATCAGAAAGACCCTTTCTACCGCTGCTGGTAGCTGGAAGGCTTACGCTGATAATACCTTCG
ATACCGCTCCTTGGGTGCCATCTGGTCTTTTTGCTGATGATGAGAGCACCCCTTCG

**pAGT7850 (SOX (YP_001129390.1; Human gammaherpesvirus 8) module-A: AATG_SOX-A_CGGT)**

GG-overhang_SOX-A_GG-overhang
AATGGAAGCTACTCCTACTCCTGCTGACCTGTTCTCTGAGGATTACCTTGTGGATACCCTGGATGGTCTGACCGTTG
ATGATCAGCAAGCTGTGCTTGCCAGCCTGAGCTTCTCTAAGTTTCTGAAGCACGCCAAGGTGAGAGATTGGTGTGCT
CAGGCTAAGATCCAGCCTTCTATGCCTGCTCTTAGGATGGCCTACAACTACTTCCTGTTCAGCAAGGTGGGCGAGTT
CATCGGTTCTGAGGATGTGTGCAACTTCTTCGTGGATAGAGTGTTCGGTGGTGTGAGGCTTCTTGATGTGGCTTCTG
TTTACGCTGCCTGCTCACAGATGAATGCTCATCAGAGGCATCACATCTGCTGCCTTGTTGAGAGGGCTACCTCTTCT
CAGTCTCTTAACCCTGTGTGGGATGCTCTGCGTGATGGCATCATCAGCTCATCTAAGTTCCACTGGGCTGTGAAGCA
GCAGAACACCAGCAAGAAAATCTTCAGCCCTTGGCCGATCACCAACAACCATTTTGTTGCTGGCCCTCTGGCTTTCG
GTCTTAGATGTGAGGAAGTGGTTAAGACCCTTCTGGCTACCTTGCTTCATCCTGATGAGGCTAACTGCCTGGACTAC
GGTTTTATGCAGTCTCCACAGAACGGCATCTTCGGTGTGTCTCTTGATTTCGCCGCTAACGTTAAGACCGATACCGA
GGGTAGACTGCAGTTCGACCCTAATTGCAAGGTGTACGAGATCAAGTGCCGGT

**pAGT7850 (SOX (YP_001129390.1; Human gammaherpesvirus 8) module-B: CGGT _SOX-B-no stop_TTCG)**

GG-overhang_SOX-B_GG-overhang
CGGTTCAAGTACACCTTCGCCAAGATGGAATGCGACCCTATCTATGCTGCTTACCAGAGGCTTTATGAGGCTCCTGG
TAAGCTGGCTCTGAAGGACTTCTTCTACAGCATCTCTAAGCCGGCCGTTGAGTACGTTGGTCTTGGTAAGCTTCCAT
CCGAGAGCGATTACCTGGTGGCTTATGATCAAGAGTGGGAAGCTTGCCCTCGGAAGAAGAGAAAGCTTACCCCTCTT
CACAACCTGATCCGTGAGTGCATTCTGCACAATAGCACCACCGAGTCCGATGTGTACGTGTTGACTGATCCTCAGGA
TACCAGGGGCCAGATCTCTATTAAGGCTCGGTTCAAGGCTAACCTGTTCGTGAATGTGCGGCACAGCTACTTCTACC
AGGTGCTCCTTCAGTCCTCTATCGTGGAAGAGTACATCGGTCTGGATAGCGGTATTCCTAGGCTGGGTTCTCCGAAG
TACTACATTGCTACCGGTTTCTTCCGTAAGCGGGGTTACCAAGATCCTGTGAACTGCACCATTGGCGGTGATGCTCT
TGATCCTCACGTTGAGATTCCGACTCTTCTGATTGTGACCCCTGTGTACTTTCCTAGGGGTGCTAAACACAGGCTGC
TTCATCAGGCTGCCAATTTCTGGTCTAGGTCCGCTAAGGACACTTTCCCGTACATCAAGTGGGACTTCAGCTACCTG
TCTGCTAACGTGCCACATTCTCCTTCG

**pAGT8740 (ME15 (YP_006990226; Stenotrophomonas phage IME15) module: AATG_ME15-stop_GCTT)**

GG-overhang_ME15_GG-overhang
AATGGCTGTGCTGAGCCTTAAAGAGTTCAGGGATATCAGAAAGGGCTGCGACGATAAGGGTATCCTGGTGATGGATG
GTGATTGGCTTGTGTTCCAGGCTATGTCTGCTGCTGAGTTCGATGCCTCTTGGGAAGAAGAGATTTGGCACCGTTGT
TGCGATCACGCTAAGGCTAGGCAGATCCTGGATGACAGCATCAAGAGCTACAGCACCCGTAAAAAGGCTTGGAACGG
TGCTCCTATTGTGCTGGCTTTCACCGACACTATCAACTGGCGGAAAGAGCTGGTTGACCCGACCTACAAAGAGAATA
GGAAGGCTACCAAGAAGCCGGTCGGTTACTTCGAGTTCCTTGACGCTCTTTTTGAGCGGCCTGAGTTCTACTGCGTG
AGGGAAGATATGCTCGAGGGTGATGATGTGATGGGCATTATCGGCTCTAACCCTTCTGCTTTCGGTGCTAGAAAGGC
CGTGATCATCAGCTGCGATAAGGACTTCAAGACCATTCCGGACTGCGACTTTCTTTGGTGCACTACCGGTAACATCC
TGACTCAGACTCAAGAGTCTGCTGATTGGTGGCACCTTTTCCAGACCATCAAGGGCGATATCACCGATGGCTACTCT
GGTATTGCTGGTTGGGGAGATTCTGCTGAGGGTTTCCTTAACGCTCCTTTCATCACTGAGCCTCAGGTGTCCGTTCT
GAAGTCCGGTAAGAACAAGGGTCAAGAGGTTACCAAGTGGGTGAAGAGGGCTCCTACTGAGTCTGAAACTCTGTGGG
ATTGCATCGTGAGCATCGGTGCTAAGGCTGGTATGACTGAAGAGGACGTTATCAAGCAGGGTCAGATGGCTAGGATC
CTGAGGTTCAACGACTACAACATCGATACCAAAGAGATTACCCTCTGGCGGCCTTCAGCTTCTTTTTGAGCTT

**pAGT9278 (ME15 (YP_006990226; Stenotrophomonas phage IME15) module: AATG_ME15-no stop_TTCG)**

GG-overhang_ME15_GG-overhang
AATGGCTGTGCTGAGCCTTAAAGAGTTCAGGGATATCAGAAAGGGCTGCGACGATAAGGGTATCCTGGTGATGGATG
GTGATTGGCTTGTGTTCCAGGCTATGTCTGCTGCTGAGTTCGATGCCTCTTGGGAAGAAGAGATTTGGCACCGTTGT
TGCGATCACGCTAAGGCTAGGCAGATCCTGGATGACAGCATCAAGAGCTACAGCACCCGTAAAAAGGCTTGGAACGG
TGCTCCTATTGTGCTGGCTTTCACCGACACTATCAACTGGCGGAAAGAGCTGGTTGACCCGACCTACAAAGAGAATA
GGAAGGCTACCAAGAAGCCGGTCGGTTACTTCGAGTTCCTTGACGCTCTTTTTGAGCGGCCTGAGTTCTACTGCGTG
AGGGAAGATATGCTCGAGGGTGATGATGTGATGGGCATTATCGGCTCTAACCCTTCTGCTTTCGGTGCTAGAAAGGC
CGTGATCATCAGCTGCGATAAGGACTTCAAGACCATTCCGGACTGCGACTTTCTTTGGTGCACTACCGGTAACATCC
TGACTCAGACTCAAGAGTCTGCTGATTGGTGGCACCTTTTCCAGACCATCAAGGGCGATATCACCGATGGCTACTCT
GGTATTGCTGGTTGGGGAGATTCTGCTGAGGGTTTCCTTAACGCTCCTTTCATCACTGAGCCTCAGGTGTCCGTTCT
GAAGTCCGGTAAGAACAAGGGTCAAGAGGTTACCAAGTGGGTGAAGAGGGCTCCTACTGAGTCTGAAACTCTGTGGG
ATTGCATCGTGAGCATCGGTGCTAAGGCTGGTATGACTGAAGAGGACGTTATCAAGCAGGGTCAGATGGCTAGGATC
CTGAGGTTCAACGACTACAACATCGATACCAAAGAGATTACCCTCTGGCGGCCTTCAGCTTCTTTTTCG

**pAGT8741 (O3-12 (NP_052100; Yersinia phage phiYeO3-12) module: AATG_O3-12-stop_GCTT)**

GG-overhang_O3-12_GG-overhang
AATGTCTCTGATCACCCTGAAGGACTTCGCTGAGATGAGAGAAGGCAAGCCGATGGAAAAGGGTGTGCTTGTGATGG
ATGGTGACTGGCTTGTGTACCAGTCTATGGCTGCTGCTGAGGTTGAAACCGATTGGGGTGATGATATCTGGACCCTT
GAGTGCGATCACGCTAAGGCTCGGTCTATTCTGGATTCCGCTATCGAGTCTTACCGGACCAGAAAGAAGGCTTGGTC
TGACGCTATGGTGGTGCTTGCTTTCACCGACGATGTGAACTGGCGGAAGGTTTTGGTGGACGAGACTTACAAAGAGA
ACCGGAAGGCTACCAGAAAGCCTGTTGGTTACAGGGACTTCCTGTCTAAGCTGTGGGAGCGTGATGAGTTCATCCAC
ATCAAAGAGGACATGCTCGAGGGTGATGACGTGATGGGTATTATCGGTTCTGGTCACGAGGTGTTCGGCTTCAAGAA
GGCTGTTCTGGTTAGCTGCGACAAGGACTTCAAGACCATTCCGGATGTGGACTTTCTGTGGTGCACTACCGGTAACA
TTCTGACCCAGACCAAAGAAACCGCTGATTGGTGGCATCTGTTCCAGACCATCAAGGGCGATATGACCGATGGTTAC
TCTGGTATTCCTGGTTGGGGAGATACTGCTGAGGCTTTCCTTAACGACCCGTTCATCGTTGAGCCTGTTGAGTCTGT
GCTGAAGTCCGGAAAGAACAAGGGTCAGACTGTGACCAAGTGGGTTAAGAGGGCTCCTGATGCTACTGAGACTCTGT
GGGATTGCATCAAGAGCATCGGTGCTAAGGCTGGTATGACCGAGCAAGAGATTATCAAGCAGGGTCAGATGGCTCGG
ATTCTGAGGTTCGAAGAGTACAACTACATCGACAAAGAGATCTACCTCTGGACCCCGAGATCTTGAGCTT

**pAGT8742 (SpiPh (NBK20419; Spirochaetia bacterium) module: AATG_SpiPh-stop_GCTT)**
GG-overhang_SpiPh_GG-overhang
AATGAGCATCAAGAGCCTTGCTCAGTTCGAGGCTATGGGCCTTTCTGGTAAGGGTCTGCTTGTGATGGATGGTGATT
GGCTTGTGTTCCAGGCTATGTCTGCTGCTGAGTTCGATGCCTCTTGGGAAGAAGAGATTTGGCACCGTTGTTGCGAT
CACGCTAAGGCTAGGCAGATTCTGGACCAGTCTATCAGCGGTTACGCCAACAGAAAGAAGGCTTGGGTTGGAGCACC
TATCGTGCTTGCTTTCACCAGCGATACCAACTGGCGGAAGGATGTGCTTGAGAGCTACAAGAGCAACCGGAAAAAGA
CCAAGAAGCCGGTTGGCTACTTCGAGTTCCTTGATGCTGTGTTCGAGGACGACCGGTACATTTGCGTGAGGGAAGAT
AACCTCGAGGGCGACGATGTGATGGGTATCATTGGTTCTAACCCTGTGCCGTTCGGCTTCAAGAAGGCTGTTCTTGT
TAGCTGCGACAAGGACTTCAAGACCATTCCGAACTGCGATTTCTTCCACGTGACCGCTGGTAAGCTTCTTGAGCAGA
ATGAGAAGTCCGCTGACTACTGGTGGATGTTCCAGACCATCAAGGGCGATATCACCGATGGCTACTCTGGTATTGCT
GGTATGGGTGAGACTGGCGCTCTTGAGTTTCTTAACGCTCCTTACAAGCTGGTGCAAGAGACTAGCCTTATCAAGGC
CGGTAAGAACAAGGGTCAAGAGAGGACTGTTTGGACCAAGAGAGAGCTGGAAGAGTCCGATTCTCTGTGGGACGCTA
TCAAGTCTATGGGTGCTAAGGCTGGGATGAGCGAAGAAGATGTTAGGGCTCAAGCTCTGGTGGCTAGGATTCTTAGG
CACAACGACTACAACTGGATCGACCGTGAGATCTACTTCCCCGAGATTTGAGCTT

**pAGT8743 (PhBO2 (YP_009790759; Pasteurella phage PHB02) module: AATG_PhBO2-stop_GCTT)**
GG-overhang_PhBO2_GG-overhang
AATGAAGTTCAACCTGAACGAGCTGAAGGACCACCTGAAGCCTTCTAAGAACCTTCTGGTGCTGGATGGTGATTGGC
TTGTGTTCCAAGCTATGAGCGCCTCTGAACAAGAGGTGGACTGGGGTAATGATATCTGGACCCTTACTTGCGACCAC
GCTAACGCTCTTGATATCCTGCAGAACTCTATCGAGGCTTGGACTACTAGACGGTCCACTTGGAAGAACGCTACCAT
TGTGGTGGCTTTCAGCGACGATACCAACTGGCGTAAGGATCTGGTGGACGAGAACTACAAGACCAACCGGAAGAAAA
CTAGGAAGCCTTGCGGTTACAGGCACTTCGTGGATACCTACATGGAACGTGAGGACACCATTTGCGTGGTGCATCCT
AATCTTGAGGCTGATGATTGCATGGGCATCATCGGTTCTGGTGGTCATCATTTCGGAACCCAGAAGGTGACCCTGAT
CAGCATCGATAAGGATTTCAGGACCGTGCCGAACTGCGATTTCCTTTGGTGCTCTACCAACAACATCCTGCCTCAGG
ATCAAGAGAGCGCTGATTTCTGGCATCTGTACCAGACCATCAAGGGCGATATCACCGATGGTTACAGCGGTATCAAA
GGTTGGGGTGAGACTGCTGAGGATTTCCTGCTTGATCCTTACATGCTGGTGCGGCAAGAGTCTACTCTTCAGAGCGG
TAAGAACAAGGGTCAGCTGAAGGTTCAGTACGTGAAGGCTGATAAGGGCGACAACTCTCTGTGGGATTGCATTGTGA
GCCTGGGTTCTAAGGTGGACATGAGCGAAGAGGACATCATTAAGCAGGCTCGGATGGCTAGGATCCTCAGGTACTCT
GATTACGACTTCAAGAACCAGCAGGTCATCCTGTGGACCCCTGATAAGTTGAATCAGTGAGCTT

**pAGT8744 (RaTL1 (YP_009785083; Ralstonia phage phiITL-1) module: AATG_RaTL1-stop_GCTT)**
GG-overhang_RaTL1_GG-overhang
AATGTCTGAGCAGAGGCTTGGTCTGCTGATCGATGCTGATTTCCTTGCTTTTCAGGCTGCTGCTAACGCTACCAGAG
TTGTTGAATGGGAGGATGGTGTGCTTACCACCTGGGCTAATATGGAAGATTGCACCCAGGCCTTTCTGTCCTCTTTG
GAGGCTCTTACCTCTAGGAACAGAAGGTGGTCTACCGCCAAGCTGATTATGTGCTTCACCGACGATCACAACTGGCG
GAAGGATATTCTGCCTAGCTACAAGGCTAACAGGTCCGGTGTTGGTAAGGGTAAGCCTATCGCTTACTGGAAGCTTG
TTGAGTGGGTGCACCAGAACTTCGAGTGCTTTGTTAGACCTGGCCTTGAGGGTGATGATTGCATGGGTATTCTGAGC
ACCAAGCCTTCTCTTGTGGGTTGCACTCATACCGTGATCGTGAGCCCTGATAAGGACTTCAAGACTGTGCCTGGTGA
GTTCTTCTGGATGACTACCGGTGAGTCTCTTGTGCTGTCTGAAGAGGATGCTAACTACTGGCACATGTACCAGACCT
TGATGGGCGATACCACTGATGGTTATGCTGGTTGTCCTGGTGTGGGTCCTACTTCTGCTGCTGAATTTCTTGCCGAG
CCGTACATTGCTTACGAGGCTTCTAAGGTGCTGAAGTCCGGTCCTAGAAAGGGTGAAGAGGTTACCTATTGGACCCA
GAGGCCTTTGGAAGCTGGTGAGGATCTTTGGGATGGTATCGTGTCCCTGTTCAAGAAGGCTGGTTTGACCGAGGAAG
ATGCTCTGGTTCAAGCTAGAGTGGCTAGGATTCTGAGGGCTAGCGATTTCGACTTCAAGGCTAAGACCCCTATTCTG
TGGGAGCGTCCACCTAAAGAGGATGTTGGTACTGACTGAGCTT

**<u>Level 0 Modules (frame underlined)</u>**

**pICH41388 (Promoter: GGAG_35Ss_TACT)**

GG-overhang-35Ss-GG-overhang

GGAGGTCAACATGGTGGAGCACGACACTCTGGTCTACTCCAAAAATGTCAAAGATACAGTCTCAGAAGATCAAAGGG
CTATTGAGACTTTTCAACAAAGGATAATTTCGGGAAACCTCCTCGGATTCCATTGCCCAGCTATCTGTCACTTCATC
GAAAGGACAGTAGAAAAGGAAGGTGGCTCCTACAAATGCCATCATTGCGATAAAGGAAAGGCTATCATTCAAGATCT
CTCTGCCGACAGTGGTCCCAAAGATGGACCCCCACCCACGAGGAGCATCGTGGAAAAAGAAGAGGTTCCAACCACGT
CTACAAAGCAAGTGGATTGATGTGACATCTCCACTGACGTAAGGGATGACGCACAATCCCACTATCCTTCGCAAGAC
CCTTCCTCTATATAAGGAAGTTCATTTCATTTGGAGAGGACACGCTACT

**pICH45089 (Promoter: GGAG_2x35Ss_TACT)**

GG-overhang-2x35Ss-GG-overhang

GGAGGTCAACATGGTGGAGCACGACACTCTGGTCTACTCCAAAAATGTCAAAGATACAGTCTCAGAAGATCAAAGGG
CTATTGAGACTTTTCAACAAAGGATAATTTCGGGAAACCTCCTCGGATTCCATTGCCCAGCTATCTGTCACTTCATC
GAAAGGACAGTAGAAAAGGAAGGTGGCTCCTACAAATGCCATCATTGCGATAAAGGAAAGGCTATCATTCAAGATCT
CTCTGCCGACAGTGGTCCCAAAGATGGACCCCCACCCACGAGGAGCATCGTGGAAAAAGAAGAGGTTCCAACCACGT
CTACAAAGCAAGTGGATTGATGTGATAACATGGTGGAGCACGACACTCTGGTCTACTCCAAAAATGTCAAAGATACA
GTCTCAGAAGATCAAAGGGCTATTGAGACTTTTCAACAAAGGATAATTTCGGGAAACCTCCTCGGATTCCATTGCCC
AGCTATCTGTCACTTCATCGAAAGGACAGTAGAAAAGGAAGGTGGCTCCTACAAATGCCATCATTGCGATAAAGGAA
AGGCTATCATTCAAGATCTCTCTGCCGACAGTGGTCCCAAAGATGGACCCCCACCCACGAGGAGCATCGTGGAAAAA
GAAGAGGTTCCAACCACGTCTACAAAGCAAGTGGATTGATGTGACATCTCCACTGACGTAAGGGATGACGCACAATC
CCACTATCCTTCGCAAGACCCTTCCTCTATATAAGGAAGTTCATTTCATTTGGAGAGGACACGCTACT

**pICH41402 (Translational enhancer: Ω-enhancer)**

GG-overhang-Ω-enhancer-GG-overhang

TACTGTATTTTTACAACAATTACCAACAACAACAAACAACAAACAACATTACAATTACTATTTACAATTACA<u>ATG</u>

**pJOG603 (Promoter: AtRPS5a)**

GG-overhang-AtRPS5a-GG-overhang

GGAGCTCAACTTTTGATTCGCTATTTGCAGTGCACCTGTGGCGTTCATCACATCTTTTGTGACACTGTTTGCACTGG
TCATTGCTATTACAAAGGACCTTCCTGATGTTGAAGGAGATCGAAAGTAAGTAACTGCACGCATAACCATTTTCTTT
CCGCTCTTTGGCTCAATCCATTTGACAGTCAAAGACAATGTTTAACCAGCTCCGTTTGATATATTGTCTTTATGTGT
TTGTTCAAGCATGTTTAGTTAATCATGCCTTTGATTGATCTTGAATAGGTTCCAAATATCAACCCTGGCAACAAAAC
TTGGAGTGAGAAACATTGCATTCCTCGGTTCTGGACTTCTGCTAGTAAATTATGTTTCAGCCATATCACTAGCTTTC
TACATGCCTCAGGTGAATTCATCTATTTCCGTCTTAACTATTTCGGTTAATTAAAGCACGAACACCATTACTGCATG
TAGAAGCTTGATAAACTATCGCCACCAATTTATTTTTGTTGCGATATTGTTACTTTCCTCAGTATGCAGCTTTGAAA
AGACCAACCCTCTTATCCTTTAACAATGAACAGGTTTTTAGAGGTAGCTTGATGATTCCTGCACATGTGATCTTGGC
TTCAGGCTTAATTTTCCAGGTAAAGCATTATGAGATACTCTTATATCTCTTACATACTTTTGAGATAATGCACAAGA
ACTTCATAACTATATGCTTTAGTTTCTGCATTTGACACTGCCAAATTCATTAATCTCTAATATCTTTGTTGTTGATC
TTTGGTAGACATGGGTACTAGAAAAAGCAAACTACACCAAGGTAAAATACTTTTGTACAAACATAAACTCGTTATCA
CGGAACATCAATGGAGTGTATATCTAACGGAGTGTAGAAACATTTGATTATTGCAGGAAGCTATCTCAGGATATTAT
CGGTTTATATGGAATCTCTTCTACGCAGAGTATCTGTTATTCCCCTTCCTCTAGCTTTCAATTTCATGGTGAGGATA
TGCAGTTTTCTTTGTATATCATTCTTCTTCTTCTTTGTAGCTTGGAGTCAAAATCGGTTCCTTCATGTACATACATC
AAGGATATGTCCTTCTGAATTTTTATATCTTGCAATAAAAATGCTTGTACCAATTGAAACACCAGCTTTTTGAGTTC
TATGATCACTGACTTGGTTCTAACCAAAAAAAAAAAAAATGTTTAATTTACATATCTAAAAGTAGGTTTAGGGAAACC
TAAACAGTAAAATATTTGTATATTATTCGAATTTCACTCATCATAAAAACTTAAATTGCACCATAAAATTTTGTTTT
ACTATTAATGATGTAATTTGTGTAACTTAAGATAAAAATAATATTCCGTAAGTTAACCGGCTAAAACCACGTATAAA
CCAGGGAACCTGTTAAACCGGTTCTTTACTGGATAAAGAAATGAAAGCCCATGTAGACAGCTCCATTAGAGCCCAAA
CCCTAAATTTCTCATCTATATAAAAGGAGTGACATTAGGGTTTTTGTTCGTCCTCTTAAAGCTTCTCGTTTTCTCTG
CCGTCTCTCTCATTCGCGCGACGCAAACGATCTTCAGGTGATCTTCTTTCTCCAAATCCTCTCTCATAACTCTGATT
TCGTACTTGTGTATTTGAGCTCACGCTCTGTTTCTCTCACCACAGCA<u>ATG</u>

**pAGT6471 (Endonuclease: AGGT_NLS-SpCas9i-NLS_TTCG)**

GG-overhang-NLS-SpCas9i(*introns*)-NLS-GG-overhang

AGGTATGGCTTCTAGCCCACCGAAGAAGAAGCGGAAGGTCAGCTGGAAAATGGACAAGAAGTACAGCATTGGACTTGATATTGGTACGAACTCAGTTGGGTGGGCCGTTATCACCGATGAATACAAGGTACCTTCGAAGAAATTTAAAGTGCTGGGCAACACAGATAGGCACAGCATTAAGAAGAACTTGATCGGAGCTCTGCTCTTTGACTCTGGAGAAACCGCGGAGGCGACAAGGCTTAAACGTACTGCGAGGAGAAGGTACACTCGCAGGAAGAACAGAATCTGTTATCTCCAAGAGATCTTTAGCAACGAGATGGCGAAG*GTAAGGATTTTTATGATATACTATGCTTATGTATTTTGTACTGAAAGCATATCCTGCTTCATTGGGATATTACTGAAAGCATTTAACTACATGTAAACTCACTTGATGATCAATAAACTTGATTTTGCAG*GTTGACGACTCGTTCTTCCATCGCCTCGAGGAATCCTTCCTGGTAGAGGAAGATAAGAAACACGAGCGTCACCCCATCTTTGGGAATATTGTTGACGAAGTAGCCTATCATGAAAAGTATCCGACTATATACCACCTTCGCAAGAAGCTGGTGGACTCAACCGATAAGGCAGACCTTCGGCTCATATACCTGGCTCTCGCGCACATGATAAAGTTTCGTGGCCATTTCTTGATCGAAGGGGACCTCAACCCGGATAACTCCGATGTGGATAAACTGTTCATTCAGCTCGTCCAAACCTACAATCAGCTGTTCGAGGAGAACCCCATCAATGCATCAG*GTAACATTCCTTAGTTACCTTTCTTTTCTTTTTCCATCATAAGTTTATAGATTGTACATGCTTTGAGATTTTTCTTTGCAAACAATCTCAG*GTGTCGACGCCAAGGCAATACTGTCTGCCAGACTTTCGAAGTCCAGACGGCTTGAGAATCTGATCGCTCAATTGCCAGGCGAGAAGAAGAACGGCTTGTTCGGGAATCTGATTGCACTGTCTCTGGGCCTCACCCCTAACTTCAAAAGCAACTTTGACCTCGCCGAGGACGCGAAGCTGCAGCTGTCAAAGGATACATACGATGATGATCTGGACAATCTGCTCGCCCAAATAG*GTAATCTTGAAATTGGAACTCTTCTTTTGTTGTCTAAACCTATCAATTTCTTTGCGGAAATTTATTTGAAGCTGTAGAGTTAAAATTGAGTCTTTTAAACTTTTGTAG*GTGATCAGTATGCCGACCTGTTCTTGGCTGCCAAGAATCTGTCAGACGCTATCTTGCTCAGTGACATTCTGCGGGTCAACACGGAGATAACCAAAGCGCCACTTAGCGCCTCCATGATCAAGAGGTACGACGAGCATCACCAGGATCTGACCCTTCTGAAGGCTTTGGTTCGCCAGCAACTCCCCGAGAAGTACAAGGAGATTTTCTTTGACCAATCGAAGAATGGCTACGCAGGGTACATTGATGGAG*GTAAGTTGTTACTTATGATTGTTTTCCTCTCTGCTACATGTATTTTGTTGTTCATTTCTGTAAGATATAAGAATTGAGTTTTCCTCTGATGATATTATTAG*GTGCAAGTCAGGAGGAATTCTACAAATTCATCAAGCCTATTCTGGAAAAGATGGACGGTACAGAGGAGCTGCTCGTTAAATTGAACCGCGAAGATTTGCTTCGGAAGCAGCGTACCTTCGACAATGGCAGCATACCGCACCAGATCCACCTCGGTGAGCTGCATGCTATCTTGAGGAGGCAAGAGGACTTCTATCCGTTCCTGAAAGACAACAGAGAGAAGATTGAAAAGATCCTCACGTTCCGCATTCCCTACTATGTAG*GTTAGTATCATATGAAGAAATACCTAGTTTCAGTTGATGAATGCTATTTTCTGACCTCAGTTGTTCTCTTTTGAGAATTATTTCTTTTCTAATTTGCCTGATTTTTCTATTAATTCATTAG*GTCCACTCGCACGCGGGAACTCGCGGTTTGCGTGGATGACACGCAAATCCGAGGAGACTATCACGCCTTGGAACTTCGAAGAGGTCGTGGACAAGGGTGCGAGTGCACAGTCCTTCATCGAAAGGATGACCAACTTCGATAAGAATCTCCCAAATGAGAAAGTCCTGCCCAAGCATAGTCTCCTGTACGAATACTTCACGGTCTACAACGAGCTGACGAAGGTGAAATATGTGACGGAGGGGATGCGCAAACCGGCCTTCCTGTCAG*GTAAATCCTGGTCCACACTTTTACGATAAAAACACAAGATTTTAAACTATGAACTGATCAATAATCATTCCTAAAAGACCACACTTTTGTTTTGTTTCTAAAGTAATTTTTACTGTTATAACAG*GTGAGCAGAAGAAGGCCATTGTCGATCTCTTGTTCAAAACCAATCGGAAGGTCACTGTGAAACAGCTTAAAGAGGACTACTTTAAGAAGATCGAATGCTTTGATTCTGTGGAAATCAGCGGCGTTGAGGATAGGTTCAATGCCTCTCTTGGCACATACCATGACCTGTTGAAAATCATCAAGGACAAGGACTTCCTTGACAACGAGGAGAACGAGGACATCCTCGAGGACATCGTGCTGACTCTCACGCTGTTTGAGGACAGAGAAATGATCGAGGAGCGCCTTAAGACTTATGCGCATCTGTTCGATGACAAGGTCATGAAGCAGTTGAAGAGGAGGAGATATACAG*GTAAGAGGTCAAAAGGTTTCCGCAATGATCCCTCTTTTTTTGTTTCTCTAGTTTCAAGAATTTGGGTATATGACTAACTTCTGAGTGTTCCTTGATGCATATTTGTGATGAGACAAATGTTTGTTCTATGTTTTAG*GTTGGGGAAGGCTCTCCAGGAAGCTCATCAACGGCATCCGCGACAAGCAATCCGGCAAGACTATACTGGACTTTCTCAAATCCGACGGTTTTGCGAATCGGAACTTCATGCAGCTTATTCACGATGACTCACTGACCTTCAAAGAAGATATCCAGAAGGCCCAAGTGTCAGGTCAGGGCGATAGCCTTCACGAACACATAGCCAACCTGGCTGGATCGCCAGCTATAAAGAAGGGCATACTGCAGACAGTGAAGGTTGTGGATGAGCTGGTGAAG*GTAAGTTCTGCATTTGGTTATGCTCCTTGCATTTTAGGTGTTCGTCGCACTTCCATTTCCATGAATAGCTAAGATTTTTTTTCTCTGCATTCATTCTTCTTGCCTCAGTTCTAACTGTTTGTGGTATTTTTGTTTTAATTATTGCTACAG*GTCATGGGCCGCCATAAGCCGGAGAACATCGTCATCGAGATGGCGAGGGAAAACCAGACGACTCAGAAAGGGCAGAAGAACTCACGGGAGCGCATGAAGCGGATAGAGGAAGGCATCAAGGAGCTTGGGAGTCAGATTCTGAAAGAGCACCCAGTCGAAAATACTCAACTCCAGAACGAGAAGCTGTACCTCTATTACCTCCAGAATGGGAGAGATATGTACGTCGACCAAGAGCTCGACATTAACAGACTCTCCGACTATGATGTGGATCACATTGTCCCTCAATCTTTCCTGAAGGACGATAGTATTGACAACAAG*GTAAAGCAACTGTGTTTTAATCAATTTCTTGTCAGGATATATGGATTATAACTTAATTTTTGAGAAATCTGTAGTATTTGGCGTGAAATGAGTTTGCTTTTTGGTTTCTCCCGTGTTATAG*GTCCTTACGCGCTCAGACAAGAACCGCGGAAAATCCGACAATGTACCCAGCGAGGAGGTTGTGAAGAAGATGAAGAACTATTGGAGGCAGCTTTTGAATGCTAAGCTCATAACCCAACGGAAATTCGACAATCTCACGAAGGCAGAAAGGGGCG

GACTGTCTGAGCTCGACAAAGCCGGCTTCATCAAGCGCCAGTTGGTTGAAACTCGTCAGATTACGAAACATGTGGCC
CAGATACTCGATTCGCGTATGAATACGAAGTATGATGAGAATGACAAACTTATCAGGGAGGTAAAG*GTAAAGTTTCC*
*AACTTTCCTTTACCATATCAAACTAAAGTTCGAAACTTTTTATTTGATCAACTTCAAGGCCACCCGATCTTTCTATT*
*CCTGATTAATTTGTGATGAATCCATATTGACTTTTGATGGTTACGCAG*GTGATCACCCTCAAGAGCAAACTGGTTAG
TGACTTCCGGAAGGACTTCCAGTTTTACAAGGTTCGCGAGATCAACAACTACCATCATGCCCATGACGCCTACCTGA
ACGCCGTTGTTGGCACTGCTCTCATCAAGAAGTATCCGAAACTGGAGTCTGAGTTTGTGTACGGGGATTACAAGGTG
TACGACGTTAGGAAGATGATCGCGAAGTCAGAACAAGAGATCGGCAAGGCTACCGCGAAATACTTCTTTTACTCGAA
TATCATGAACTTCTTCAAGACAGAGATCACTCTGGCGAATGGTGAAATCCGGAAGAGGCCTCTGATCGAGACAAATG
GCGAAACAG*GTCTGTCTTTCCTATTTCATATGTTTAATCCTAGGAATTTGATCAATTGATTGTATGTATGTCGATCC*
*CAAGACTTTCTTGTTCACTTATATCTTAACTCTCTCTTTGCTGTTTCTTGCAG*GTGAGATTGTCTGGGATAAGGGCA
GGGATTTTGCGACTGTGCGTAAGGTTCTCAGCATGCCCCAAGTCAACATAGTCAAGAAAACGGAGGTTCAAACCGGT
GGTTTCTCCAAGGAGTCCATTCTCCCTAAGCGCAACTCCGACAAACTGATTGCGAGGAAGAAGGATTGGGATCCGAA
GAAATACGGAGGCTTTGATAGCCCTACCGTGGCATACAGCGTACTGGTAGTGGCCAAGGTGGAGAAGGGCAAGAGCA
AGAAACTGAAAAGCGTCAAGGAACTGCTTGGAATTACCATAATGGAAAGGTCCTCGTTCGAGAAGAATCCGATCGAC
TTCCTCGAGGCTAAAG*GTAAAATATTGGATGCCAGACGATATTCTTTCTTTTGATTTGTAACTTTTTCCTGTCAAGG*
*TCGATAAATTTTATTTTTTTTGGTAAAAGGTCGATAATTTTTTTTTGGAGCCATTATGTAATTTTCCTAATTAACTG*
*AACCAAAATTATACTTTGCAG*GTTACAAAGAGGTGAAGAAAGACCTCATTATCAAACTGCCCAAGTATTCGCTTTTC
GAATTGGAAAATGGCAGAAAACGCATGCTGGCATCTGCCGGAGAACTGCAGAAGGGCAACGAGCTGGCATTGCCCAG
TAAGTACGTCAACTTCCTGTACTTGGCCTCACACTATGAGAAGCTGAAGGGGTCACCAGAGGACAACGAGCAGAAGC
AGTTGTTTGTCGAGCAGCACAAGCACTATCTTGATGAGATCATAGAGCAGATCAGCGAATTTTCCAAGCGGGTCATT
CTTGCAGACGCTAACCTCGATAAG*GTAAGGACTTCTCATGAATATTAGTGGCAGATTAGTGTTGTTAAAGTCTTTGG*
*TTAGATAATCGATGCCTCCTAATTGTCCATGTTTTACTGGTTTTCTACAATTACAG*GTGCTTTCCGCGTACAACAAG
CACAGAGATAAGCCGATAAGGGAACAAGCGGAAAACATCATCCACCTGTTCACACTGACCAATCTGGGAGCCCCAGC
AGCCTTTAAGTACTTCGATACCACTATCGACAGAAAGCGCTACACATCAACCAAGGAAGTGTTGGACGCTACCCTTA
TTCACCAATCTATTACAGGGCTCTATGAGACAAGGATAGATCTGTCGCAGTTGGGTGGTGACTCTAGGGCTGACCCA
AAGAAGAAGCGTAAAGTCGGTTCG

**pAGT6472 (Endonuclease: AGGT_NLS-SpCas9i(D10A)-NLS_TTCG)**

GG-overhang-NLS-SpCas9i(D10A)(*introns*)-NLS-GG-overhang

AGGTATGGCTTCTAGCCCACCGAAGAAGAAGCGGAAGGTCAGCTGGAAAATGGACAAGAAGTACAGCATTGGACTTG
CAATTGGTACGAACTCAGTTGGGTGGGCCGTTATCACCGATGAATACAAGGTACCTTCGAAGAAATTTAAAGTGCTG
GGCAACACAGATAGGCACAGCATTAAGAAGAACTTGATCGGAGCTCTGCTCTTTGACTCTGGAGAAACCGCGGAGGC
GACAAGGCTTAAACGTACTGCGAGGAGAAGGTACACTCGCAGGAAGAACAGAATCTGTTATCTCCAAGAGATCTTTA
GCAACGAGATGGCGAAG*GTAAGGATTTTTATGATATACTATGCTTATGTATTTTGTACTGAAAGCATATCCTGCTTC*
*ATTGGGATATTACTGAAAGCATTTAACTACATGTAAACTCACTTGATGATCAATAAACTTGATTTTGCAG*GTTGACG
ACTCGTTCTTCCATCGCCTCGAGGAATCCTTCCTGGTAGAGGAAGATAAGAAACACGAGCGTCACCCCATCTTTGGG
AATATTGTTGACGAAGTAGCCTATCATGAAAAGTATCCGACTATATACCACCTTCGCAAGAAGCTGGTGGACTCAAC
CGATAAGGCAGACCTTCGGCTCATATACCTGGCTCTCGCGCACATGATAAAGTTTCGTGGCCATTTCTTGATCGAAG
GGGACCTCAACCCGGATAACTCCGATGTGGATAAACTGTTCATTCAGCTCGTCCAAACCTACAATCAGCTGTTCGAG
GAGAACCCCATCAATGCATCAG*GTAACATTCCTTAGTTACCTTTCTTTTCTTTTTCCATCATAAGTTTATAGATTGT*
*ACATGCTTTGAGATTTTTCTTTGCAAACAATCTCAG*GTGTCGACGCCAAGGCAATACTGTCTGCCAGACTTTCGAAG
TCCAGACGGCTTGAGAATCTGATCGCTCAATTGCCAGGCGAGAAGAAGAACGGCTTGTTCGGGAATCTGATTGCACT
GTCTCTGGGCCTCACCCCTAACTTCAAAAGCAACTTTGACCTCGCCGAGGACGCGAAGCTGCAGCTGTCAAAGGATA
CATACGATGATGATCTGGACAATCTGCTCGCCCAAATAG*GTAATCTTGAAATTGGAACTCTTCTTTTGTTGTCTAAA*
*CCTATCAATTTCTTTGCGGAAATTTATTTGAAGCTGTAGAGTTAAAATTGAGTCTTTTAAACTTTTGTAG*GTGATCA
GTATGCCGACCTGTTCTTGGCTGCCAAGAATCTGTCAGACGCTATCTTGCTCAGTGACATTCTGCGGGTCAACACGG
AGATAACCAAAGCGCCACTTAGCGCCTCCATGATCAAGAGGTACGACGAGCATCACCAGGATCTGACCCTTCTGAAG
GCTTTGGTTCGCCAGCAACTCCCCGAGAAGTACAAGGAGATTTTCTTTGACCAATCGAAGAATGGCTACGCAGGGTA
CATTGATGGAG*GTAAGTTGTTACTTATGATTGTTTTCCTCTCTGCTACATGTATTTTGTTGTTCATTTCTGTAAGAT*
*ATAAGAATTGAGTTTTCCTCTGATGATATTATTAG*GTGCAAGTCAGGAGGAATTCTACAAATTCATCAAGCCTATTC
TGGAAAAGATGGACGGTACAGAGGAGCTGCTCGTTAAATTGAACCGCGAAGATTTGCTTCGGAAGCAGCGTACCTTC
GACAATGGCAGCATACCGCACCAGATCCACCTCGGTGAGCTGCATGCTATCTTGAGGAGGCAAGAGGACTTCTATCC
GTTCCTGAAAGACAACAGAGAGAAGATTGAAAAGATCCTCACGTTCCGCATTCCCTACTATGTAG*GTTAGTATCATA*
*TGAAGAAATACCTAGTTTCAGTTGATGAATGCTATTTTCTGACCTCAGTTGTTCTCTTTTGAGAATTATTTCTTTTC*

*TAATTTGCCTGATTTTTCTATTAATTCATTAG*GTCCACTCGCACGCGGGAACTCGCGGTTTGCGTGGATGACACGCA
AATCCGAGGAGACTATCACGCCTTGGAACTTCGAAGAGGTCGTGGACAAGGGTGCGAGTGCACAGTCCTTCATCGAA
AGGATGACCAACTTCGATAAGAATCTCCCAAATGAGAAAGTCCTGCCCAAGCATAGTCTCCTGTACGAATACTTCAC
GGTCTACAACGAGCTGACGAAGGTGAAATATGTGACGGAGGGGATGCGCAAACCGGCCTTCCTGTCAG*GTAAATCCT*
*GGTCCACACTTTTACGATAAAAACACAAGATTTTAAACTATGAACTGATCAATAATCATTCCTAAAAGACCACACTT*
*TTGTTTTGTTTCTAAAGTAATTTTTACTGTTATAACAG*GTGAGCAGAAGAAGGCCATTGTCGATCTCTTGTTCAAAA
CCAATCGGAAGGTCACTGTGAAACAGCTTAAAGAGGACTACTTTAAGAAGATCGAATGCTTTGATTCTGTGGAAATC
AGCGGCGTTGAGGATAGGTTCAATGCCTCTCTTGGCACATACCATGACCTGTTGAAAATCATCAAGGACAAGGACTT
CCTTGACAACGAGGAGAACGAGGACATCCTCGAGGACATCGTGCTGACTCTCACGCTGTTTGAGGACAGAGAAATGA
TCGAGGAGCGCCTTAAGACTTATGCGCATCTGTTCGATGACAAGGTCATGAAGCAGTTGAAGAGGAGGAGATATACA
G*GTAAGAGGTCAAAAGGTTTCCGCAATGATCCCTCTTTTTTTGTTTCTCTAGTTTCAAGAATTTGGGTATATGACTA*
*ACTTCTGAGTGTTCCTTGATGCATATTTGTGATGAGACAAATGTTTGTTCTATGTTTTAG*GTTGGGGAAGGCTCTCC
AGGAAGCTCATCAACGGCATCCGCGACAAGCAATCCGGCAAGACTATACTGGACTTTCTCAAATCCGACGGTTTTGC
GAATCGGAACTTCATGCAGCTTATTCACGATGACTCACTGACCTTCAAAGAAGATATCCAGAAGGCCCAAGTGTCAG
GTCAGGGCGATAGCCTTCACGAACACATAGCCAACCTGGCTGGATCGCCAGCTATAAAGAAGGGCATACTGCAGACA
GTGAAGGTTGTGGATGAGCTGGTGAAG*GTAAGTTCTGCATTTGGTTATGCTCCTTGCATTTTAGGTGTTCGTCGCAC*
*TTCCATTTCCATGAATAGCTAAGATTTTTTTTCTCTGCATTCATTCTTCTTGCCTCAGTTCTAACTGTTTGTGGTAT*
*TTTTGTTTTAATTATTGCTACAG*GTCATGGGCCGCCATAAGCCGGAGAACATCGTCATCGAGATGGCGAGGGAAAAC
CAGACGACTCAGAAAGGGCAGAAGAACTCACGGGAGCGCATGAAGCGGATAGAGGAAGGCATCAAGGAGCTTGGGAG
TCAGATTCTGAAAGAGCACCCAGTCGAAAATACTCAACTCCAGAACGAGAAGCTGTACCTCTATTACCTCCAGAATG
GGAGAGATATGTACGTCGACCAAGAGCTCGACATTAACAGACTCTCCGACTATGATGTGGATCACATTGTCCCTCAA
TCTTTCCTGAAGGACGATAGTATTGACAACAAG*GTAAAGCAACTGTGTTTTAATCAATTTCTTGTCAGGATATATGG*
*ATTATAACTTAATTTTTGAGAAATCTGTAGTATTTGGCGTGAAATGAGTTTGCTTTTTGGTTTCTCCCGTGTTATAG*
GTCCTTACGCGCTCAGACAAGAACCGCGGAAAATCCGACAATGTACCCAGCGAGGAGGTTGTGAAGAAGATGAAGAA
CTATTGGAGGCAGCTTTTGAATGCTAAGCTCATAACCCAACGGAAATTCGACAATCTCACGAAGGCAGAAAGGGGCG
GACTGTCTGAGCTCGACAAAGCCGGCTTCATCAAGCGCCAGTTGGTTGAAACTCGTCAGATTACGAAACATGTGGCC
CAGATACTCGATTCGCGTATGAATACGAAGTATGATGAGAATGACAAACTTATCAGGGAGGTAAAG*GTAAAGTTTCC*
*AACTTTCCTTTACCATATCAAACTAAAGTTCGAAACTTTTTATTTGATCAACTTCAAGGCCACCCGATCTTTCTATT*
*CCTGATTAATTTGTGATGAATCCATATTGACTTTTGATGGTTACGCAG*GTGATCACCCTCAAGAGCAAACTGGTTAG
TGACTTCCGGAAGGACTTCCAGTTTTACAAGGTTCGCGAGATCAACAACTACCATCATGCCCATGACGCCTACCTGA
ACGCCGTTGTTGGCACTGCTCTCATCAAGAAGTATCCGAAACTGGAGTCTGAGTTTGTGTACGGGGATTACAAGGTG
TACGACGTTAGGAAGATGATCGCGAAGTCAGAACAAGAGATCGGCAAGGCTACCGCGAAATACTTCTTTTACTCGAA
TATCATGAACTTCTTCAAGACAGAGATCACTCTGGCGAATGGTGAAATCCGGAAGAGGCCTCTGATCGAGACAAATG
GCGAAACAG*GTCTGTCTTTCCTATTTCATATGTTTAATCCTAGGAATTTGATCAATTGATTGTATGTATGTCGATCC*
*CAAGACTTTCTTGTTCACTTATATCTTAACTCTCTCTTTGCTGTTTCTTGCAG*GTGAGATTGTCTGGGATAAGGGCA
GGGATTTTGCGACTGTGCGTAAGGTTCTCAGCATGCCCCAAGTCAACATAGTCAAGAAAACGGAGGTTCAAACCGGT
GGTTTCTCCAAGGAGTCCATTCTCCCTAAGCGCAACTCCGACAAACTGATTGCGAGGAAGAAGGATTGGGATCCGAA
GAAATACGGAGGCTTTGATAGCCCTACCGTGGCATACAGCGTACTGGTAGTGGCCAAGGTGGAGAAGGGCAAGAGCA
AGAAACTGAAAAGCGTCAAGGAACTGCTTGGAATTACCATAATGGAAAGGTCCTCGTTCGAGAAGAATCCGATCGAC
TTCCTCGAGGCTAAAG*GTAAAATATTGGATGCCAGACGATATTCTTTCTTTTGATTTGTAACTTTTTCCTGTCAAGG*
*TCGATAAATTTTATTTTTTTTGGTAAAAGGTCGATAATTTTTTTTTGGAGCCATTATGTAATTTTCCTAATTAACTG*
*AACCAAAATTATACTTTGCAG*GTTACAAAGAGGTGAAGAAAGACCTCATTATCAAACTGCCCAAGTATTCGCTTTTC
GAATTGGAAAATGGCAGAAAACGCATGCTGGCATCTGCCGGAGAACTGCAGAAGGGCAACGAGCTGGCATTGCCCAG
TAAGTACGTCAACTTCCTGTACTTGGCCTCACACTATGAGAAGCTGAAGGGGTCACCAGAGGACAACGAGCAGAAGC
AGTTGTTTGTCGAGCAGCACAAGCACTATCTTGATGAGATCATAGAGCAGATCAGCGAATTTTCCAAGCGGGTCATT
CTTGCAGACGCTAACCTCGATAAG*GTAAGGACTTCTCATGAATATTAGTGGCAGATTAGTGTTGTTAAAGTCTTTGG*
*TTAGATAATCGATGCCTCCTAATTGTCCATGTTTTACTGGTTTTCTACAATTACAG*GTGCTTTCCGCGTACAACAAG
CACAGAGATAAGCCGATAAGGGAACAAGCGGAAAACATCATCCACCTGTTCACACTGACCAATCTGGGAGCCCCAGC
AGCCTTTAAGTACTTCGATACCACTATCGACAGAAAGCGCTACACATCAACCAAGGAAGTGTTGGACGCTACCCTTA
TTCACCAATCTATTACAGGGCTCTATGAGACAAGGATAGATCTGTCGCAGTTGGGTGGTGACTCTAGGGCTGACCCA
AAGAAGAAGCGTAAAGTCGGTTCG

**pAGT6473 (Endonuclease: AGGT_NLS-SpCas9i(H840A)-NLS_TTCG)**
GG-overhang-NLS-SpCas9i(H840A)(*introns*)-NLS-GG-overhang
AGGTATGGCTTCTAGCCCACCGAAGAAGAAGCGGAAGGTCAGCTGGAAAATGGACAAGAAGTACAGCATTGGACTTG
ATATTGGTACGAACTCAGTTGGGTGGGCCGTTATCACCGATGAATACAAGGTACCTTCGAAGAAATTTAAAGTGCTG
GGCAACACAGATAGGCACAGCATTAAGAAGAACTTGATCGGAGCTCTGCTCTTTGACTCTGGAGAAACCGCGGAGGC

GACAAGGCTTAAACGTACTGCGAGGAGAAGGTACACTCGCAGGAAGAACAGAATCTGTTATCTCCAAGAGATCTTTA
GCAACGAGATGGCGAAGGTAAGGATTTTTATGATATACTATGCTTATGTATTTTGTACTGAAAGCATATCCTGCTTC
ATTGGGATATTACTGAAAGCATTTAACTACATGTAAACTCACTTGATGATCAATAAACTTGATTTTGCAGGTTGACG
ACTCGTTCTTCCATCGCCTCGAGGAATCCTTCCTGGTAGAGGAAGATAAGAAACACGAGCGTCACCCCATCTTTGGG
AATATTGTTGACGAAGTAGCCTATCATGAAAAGTATCCGACTATATACCACCTTCGCAAGAAGCTGGTGGACTCAAC
CGATAAGGCAGACCTTCGGCTCATATACCTGGCTCTCGCGCACATGATAAAGTTTCGTGGCCATTTCTTGATCGAAG
GGGACCTCAACCCGGATAACTCCGATGTGGATAAACTGTTCATTCAGCTCGTCCAAACCTACAATCAGCTGTTCGAG
GAGAACCCCATCAATGCATCAGGTAACATTCCTTAGTTACCTTTCTTTTCTTTTTCCATCATAAGTTTATAGATTGT
ACATGCTTTGAGATTTTTCTTTGCAAACAATCTCAGGTGTCGACGCCAAGGCAATACTGTCTGCCAGACTTTCGAAG
TCCAGACGGCTTGAGAATCTGATCGCTCAATTGCCAGGCGAGAAGAAGAACGGCTTGTTCGGGAATCTGATTGCACT
GTCTCTGGGCCTCACCCCTAACTTCAAAAGCAACTTTGACCTCGCCGAGGACGCGAAGCTGCAGCTGTCAAAGGATA
CATACGATGATGATCTGGACAATCTGCTCGCCCAAATAGGTAATCTTGAAATTGGAACTCTTCTTTTGTTGTCTAAA
CCTATCAATTTCTTTGCGGAAATTTATTTGAAGCTGTAGAGTTAAAATTGAGTCTTTTAAACTTTTGTAGGTGATCA
GTATGCCGACCTGTTCTTGGCTGCCAAGAATCTGTCAGACGCTATCTTGCTCAGTGACATTCTGCGGGTCAACACGG
AGATAACCAAAGCGCCACTTAGCGCCTCCATGATCAAGAGGTACGACGAGCATCACCAGGATCTGACCCTTCTGAAG
GCTTTGGTTCGCCAGCAACTCCCCGAGAAGTACAAGGAGATTTTCTTTGACCAATCGAAGAATGGCTACGCAGGGTA
CATTGATGGAGGTAAGTTGTTACTTATGATTGTTTTCCTCTCTGCTACATGTATTTTGTTGTTCATTTCTGTAAGAT
ATAAGAATTGAGTTTTCCTCTGATGATATTATTAGGTGCAAGTCAGGAGGAATTCTACAAATTCATCAAGCCTATTC
TGGAAAAGATGGACGGTACAGAGGAGCTGCTCGTTAAATTGAACCGCGAAGATTTGCTTCGGAAGCAGCGTACCTTC
GACAATGGCAGCATACCGCACCAGATCCACCTCGGTGAGCTGCATGCTATCTTGAGGAGGCAAGAGGACTTCTATCC
GTTCCTGAAAGACAACAGAGAGAAGATTGAAAAGATCCTCACGTTCCGCATTCCCTACTATGTAGGTTAGTATCATA
TGAAGAAATACCTAGTTTCAGTTGATGAATGCTATTTTCTGACCTCAGTTGTTCTCTTTTGAGAATTATTTCTTTTC
TAATTTGCCTGATTTTTCTATTAATTCATTAGGTCCACTCGCACGCGGGAACTCGCGGTTTGCGTGGATGACACGCA
AATCCGAGGAGACTATCACGCCTTGGAACTTCGAAGAGGTCGTGGACAAGGGTGCGAGTGCACAGTCCTTCATCGAA
AGGATGACCAACTTCGATAAGAATCTCCCAAATGAGAAAGTCCTGCCCAAGCATAGTCTCCTGTACGAATACTTCAC
GGTCTACAACGAGCTGACGAAGGTGAAATATGTGACGGAGGGGATGCGCAAACCGGCCTTCCTGTCAGGTAAATCCT
GGTCCACACTTTTACGATAAAAACACAAGATTTTAAACTATGAACTGATCAATAATCATTCCTAAAAGACCACACTT
TTGTTTTGTTTCTAAAGTAATTTTTACTGTTATAACAGGTGAGCAGAAGAAGGCCATTGTCGATCTCTTGTTCAAAA
CCAATCGGAAGGTCACTGTGAAACAGCTTAAAGAGGACTACTTTAAGAAGATCGAATGCTTTGATTCTGTGGAAATC
AGCGGCGTTGAGGATAGGTTCAATGCCTCTCTTGGCACATACCATGACCTGTTGAAAATCATCAAGGACAAGGACTT
CCTTGACAACGAGGAGAACGAGGACATCCTCGAGGACATCGTGCTGACTCTCACGCTGTTTGAGGACAGAGAAATGA
TCGAGGAGCGCCTTAAGACTTATGCGCATCTGTTCGATGACAAGGTCATGAAGCAGTTGAAGAGGAGGAGATATACA
GGTAAGAGGTCAAAAGGTTTCCGCAATGATCCCTCTTTTTTTGTTTCTCTAGTTTCAAGAATTTGGGTATATGACTA
ACTTCTGAGTGTTCCTTGATGCATATTTGTGATGAGACAAATGTTTGTTCTATGTTTTAGGTTGGGGAAGGCTCTCC
AGGAAGCTCATCAACGGCATCCGCGACAAGCAATCCGGCAAGACTATACTGGACTTTCTCAAATCCGACGGTTTTGC
GAATCGGAACTTCATGCAGCTTATTCACGATGACTCACTGACCTTCAAAGAAGATATCCAGAAGGCCCAAGTGTCAG
GTCAGGGCGATAGCCTTCACGAACACATAGCCAACCTGGCTGGATCGCCAGCTATAAAGAAGGGCATACTGCAGACA
GTGAAGGTTGTGGATGAGCTGGTGAAGGTAAGTTCTGCATTTGGTTATGCTCCTTGCATTTTAGGTGTTCGTCGCAC
TTCCATTTCCATGAATAGCTAAGATTTTTTTTCTCTGCATTCATTCTTCTTGCCTCAGTTCTAACTGTTTGTGGTAT
TTTTGTTTTAATTATTGCTACAGGTCATGGGCCGCCATAAGCCGGAGAACATCGTCATCGAGATGGCGAGGGAAAAC
CAGACGACTCAGAAAGGGCAGAAGAACTCACGGGAGCGCATGAAGCGGATAGAGGAAGGCATCAAGGAGCTTGGGAG
TCAGATTCTGAAAGAGCACCCAGTCGAAAATACTCAACTCCAGAACGAGAAGCTGTACCTCTATTACCTCCAGAATG
GGAGAGATATGTACGTCGACCAAGAGCTCGACATTAACAGACTCTCCGACTATGATGTGGATGCAATTGTCCCTCAA
TCTTTCCTGAAGGACGATAGTATTGACAACAAGGTAAAGCAACTGTGTTTTAATCAATTTCTTGTCAGGATATATGG
ATTATAACTTAATTTTTGAGAAATCTGTAGTATTTGGCGTGAAATGAGTTTGCTTTTTGGTTTCTCCCGTGTTATAG
GTCCTTACGCGCTCAGACAAGAACCGCGGAAAATCCGACAATGTACCCAGCGAGGAGGTTGTGAAGAAGATGAAGAA
CTATTGGAGGCAGCTTTTGAATGCTAAGCTCATAACCCAACGGAAATTCGACAATCTCACGAAGGCAGAAAGGGGCG
GACTGTCTGAGCTCGACAAAGCCGGCTTCATCAAGCGCCAGTTGGTTGAAACTCGTCAGATTACGAAACATGTGGCC
CAGATACTCGATTCGCGTATGAATACGAAGTATGATGAGAATGACAAACTTATCAGGGAGGTAAAGGTAAAGTTTCC
AACTTTCCTTTACCATATCAAACTAAAGTTCGAAACTTTTTATTTGATCAACTTCAAGGCCACCCGATCTTTCTATT

*CCTGATTAATTTGTGATGAATCCATATTGACTTTTGATGGTTACGCAG*GTGATCACCCTCAAGAGCAAACTGGTTAG
TGACTTCCGGAAGGACTTCCAGTTTTACAAGGTTCGCGAGATCAACAACTACCATCATGCCCATGACGCCTACCTGA
ACGCCGTTGTTGGCACTGCTCTCATCAAGAAGTATCCGAAACTGGAGTCTGAGTTTGTGTACGGGGATTACAAGGTG
TACGACGTTAGGAAGATGATCGCGAAGTCAGAACAAGAGATCGGCAAGGCTACCGCGAAATACTTCTTTTACTCGAA
TATCATGAACTTCTTCAAGACAGAGATCACTCTGGCGAATGGTGAAATCCGGAAGAGGCCTCTGATCGAGACAAATG
GCGAAACAG*GTCTGTCTTTCCTATTTCATATGTTTAATCCTAGGAATTTGATCAATTGATTGTATGTATGTCGATCC*
*CAAGACTTTCTTGTTCACTTATATCTTAACTCTCTCTTTGCTGTTTCTTGCAG*GTGAGATTGTCTGGGATAAGGGCA
GGGATTTTGCGACTGTGCGTAAGGTTCTCAGCATGCCCCAAGTCAACATAGTCAAGAAAACGGAGGTTCAAACCGGT
GGTTTCTCCAAGGAGTCCATTCTCCCTAAGCGCAACTCCGACAAACTGATTGCGAGGAAGAAGGATTGGGATCCGAA
GAAATACGGAGGCTTTGATAGCCCTACCGTGGCATACAGCGTACTGGTAGTGGCCAAGGTGGAGAAGGGCAAGAGCA
AGAAACTGAAAAGCGTCAAGGAACTGCTTGGAATTACCATAATGGAAAGGTCCTCGTTCGAGAAGAATCCGATCGAC
TTCCTCGAGGCTAAAG*GTAAAATATTGGATGCCAGACGATATTCTTTCTTTTGATTTGTAACTTTTTCCTGTCAAGG*
*TCGATAAATTTTATTTTTTTTGGTAAAAGGTCGATAATTTTTTTTTGGAGCCATTATGTAATTTTCCTAATTAACTG*
*AACCAAAATTATACTTTGCAG*GTTACAAAGAGGTGAAGAAAGACCTCATTATCAAACTGCCCAAGTATTCGCTTTTC
GAATTGGAAAATGGCAGAAAACGCATGCTGGCATCTGCCGGAGAACTGCAGAAGGGCAACGAGCTGGCATTGCCCAG
TAAGTACGTCAACTTCCTGTACTTGGCCTCACACTATGAGAAGCTGAAGGGGTCACCAGAGGACAACGAGCAGAAGC
AGTTGTTTGTCGAGCAGCACAAGCACTATCTTGATGAGATCATAGAGCAGATCAGCGAATTTTCCAAGCGGGTCATT
CTTGCAGACGCTAACCTCGATAAG*GTAAGGACTTCTCATGAATATTAGTGGCAGATTAGTGTTGTTAAAGTCTTTGG*
*TTAGATAATCGATGCCTCCTAATTGTCCATGTTTTACTGGTTTTCTACAATTACAG*GTGCTTTCCGCGTACAACAAG
CACAGAGATAAGCCGATAAGGGAACAAGCGGAAAACATCATCCACCTGTTCACACTGACCAATCTGGGAGCCCCAGC
AGCCTTTAAGTACTTCGATACCACTATCGACAGAAAGCGCTACACATCAACCAAGGAAGTGTTGGACGCTACCCTTA
TTCACCAATCTATTACAGGGCTCTATGAGACAAGGATAGATCTGTCGCAGTTGGGTGGTGACTCTAGGGCTGACCCA
AAGAAGAAGCGTAAAGTCGGTTCG

**pAGT6468 (Endonuclease: AATG_NLS-SpCas9i-NLS_TTCG)**

GG-overhang-NLS-SpCas9i(*introns*)-NLS-GG-overhang

AATGGCTTCTAGCCCACCGAAGAAGAAGCGGAAGGTCAGCTGGAAAATGGACAAGAAGTACAGCATTGGACTTGATA
TTGGTACGAACTCAGTTGGGTGGGCCGTTATCACCGATGAATACAAGGTACCTTCGAAGAAATTTAAAGTGCTGGGC
AACACAGATAGGCACAGCATTAAGAAGAACTTGATCGGAGCTCTGCTCTTTGACTCTGGAGAAACCGCGGAGGCGAC
AAGGCTTAAACGTACTGCGAGGAGAAGGTACACTCGCAGGAAGAACAGAATCTGTTATCTCCAAGAGATCTTTAGCA
ACGAGATGGCGAAG*GTAAGGATTTTTATGATATACTATGCTTATGTATTTTGTACTGAAAGCATATCCTGCTTCATT*
*GGGATATTACTGAAAGCATTTAACTACATGTAAACTCACTTGATGATCAATAAACTTGATTTTGCAG*GTTGACGACT
CGTTCTTCCATCGCCTCGAGGAATCCTTCCTGGTAGAGGAAGATAAGAAACACGAGCGTCACCCCATCTTTGGGAAT
ATTGTTGACGAAGTAGCCTATCATGAAAAGTATCCGACTATATACCACCTTCGCAAGAAGCTGGTGGACTCAACCGA
TAAGGCAGACCTTCGGCTCATATACCTGGCTCTCGCGCACATGATAAAGTTTCGTGGCCATTTCTTGATCGAAGGGG
ACCTCAACCCGGATAACTCCGATGTGGATAAACTGTTCATTCAGCTCGTCCAAACCTACAATCAGCTGTTCGAGGAG
AACCCCATCAATGCATCAG*GTAACATTCCTTAGTTACCTTTCTTTTCTTTTTCCATCATAAGTTTATAGATTGTACA*
*TGCTTTGAGATTTTTCTTTGCAAACAATCTCAG*GTGTCGACGCCAAGGCAATACTGTCTGCCAGACTTTCGAAGTCC
AGACGGCTTGAGAATCTGATCGCTCAATTGCCAGGCGAGAAGAAGAACGGCTTGTTCGGGAATCTGATTGCACTGTC
TCTGGGCCTCACCCCTAACTTCAAAAGCAACTTTGACCTCGCCGAGGACGCGAAGCTGCAGCTGTCAAAGGATACAT
ACGATGATGATCTGGACAATCTGCTCGCCCAAATAG*GTAATCTTGAAATTGGAACTCTTCTTTTGTTGTCTAAACCT*
*ATCAATTTCTTTGCGGAAATTTATTTGAAGCTGTAGAGTTAAAATTGAGTCTTTTAAACTTTTGTAG*GTGATCAGTA
TGCCGACCTGTTCTTGGCTGCCAAGAATCTGTCAGACGCTATCTTGCTCAGTGACATTCTGCGGGTCAACACGGAGA
TAACCAAAGCGCCACTTAGCGCCTCCATGATCAAGAGGTACGACGAGCATCACCAGGATCTGACCCTTCTGAAGGCT
TTGGTTCGCCAGCAACTCCCCGAGAAGTACAAGGAGATTTTCTTTGACCAATCGAAGAATGGCTACGCAGGGTACAT
TGATGGAG*GTAAGTTGTTACTTATGATTGTTTTCCTCTCTGCTACATGTATTTTGTTGTTCATTTCTGTAAGATATA*
*AGAATTGAGTTTTCCTCTGATGATATTATTAG*GTGCAAGTCAGGAGGAATTCTACAAATTCATCAAGCCTATTCTGG
AAAAGATGGACGGTACAGAGGAGCTGCTCGTTAAATTGAACCGCGAAGATTTGCTTCGGAAGCAGCGTACCTTCGAC
AATGGCAGCATACCGCACCAGATCCACCTCGGTGAGCTGCATGCTATCTTGAGGAGGCAAGAGGACTTCTATCCGTT

CCTGAAAGACAACAGAGAGAAGATTGAAAAGATCCTCACGTTCCGCATTCCCTACTATGTAG*GTTAGTATCATATGAAGAAATACCTAGTTTCAGTTGATGAATGCTATTTTCTGACCTCAGTTGTTCTCTTTTGAGAATTATTTCTTTTCTAATTTGCCTGATTTTTCTATTAATTCATTAG*GTCCACTCGCACGCGGGAACTCGCGGTTTGCGTGGATGACACGCAAATCCGAGGAGACTATCACGCCTTGGAACTTCGAAGAGGTCGTGGACAAGGGTGCGAGTGCACAGTCCTTCATCGAAAGGATGACCAACTTCGATAAGAATCTCCCAAATGAGAAAGTCCTGCCCAAGCATAGTCTCCTGTACGAATACTTCACGGTCTACAACGAGCTGACGAAGGTGAAATATGTGACGGAGGGGATGCGCAAACCGGCCTTCCTGTCAG*GTAAATCCTGGTCCACACTTTTACGATAAAAACACAAGATTTTAAACTATGAACTGATCAATAATCATTCCTAAAAGACCACACTTTTGTTTTGTTTCTAAAGTAATTTTTACTGTTATAACAG*GTGAGCAGAAGAAGGCCATTGTCGATCTCTTGTTCAAAACCAATCGGAAGGTCACTGTGAAACAGCTTAAAGAGGACTACTTTAAGAAGATCGAATGCTTTGATTCTGTGGAAATCAGCGGCGTTGAGGATAGGTTCAATGCCTCTCTTGGCACATACCATGACCTGTTGAAAATCATCAAGGACAAGGACTTCCTTGACAACGAGGAGAACGAGGACATCCTCGAGGACATCGTGCTGACTCTCACGCTGTTTGAGGACAGAGAAATGATCGAGGAGCGCCTTAAGACTTATGCGCATCTGTTCGATGACAAGGTCATGAAGCAGTTGAAGAGGAGGAGATATACAG*GTAAGAGGTCAAAAGGTTTCCGCAATGATCCCTCTTTTTTTGTTTCTCTAGTTTCAAGAATTTGGGTATATGACTAACTTCTGAGTGTTCCTTGATGCATATTTGTGATGAGACAAATGTTTGTTCTATGTTTTAG*GTTGGGGAAGGCTCTCCAGGAAGCTCATCAACGGCATCCGCGACAAGCAATCCGGCAAGACTATACTGGACTTTCTCAAATCCGACGGTTTTGCGAATCGGAACTTCATGCAGCTTATTCACGATGACTCACTGACCTTCAAAGAAGATATCCAGAAGGCCCAAGTGTCAGGTCAGGGCGATAGCCTTCACGAACACATAGCCAACCTGGCTGGATCGCCAGCTATAAAGAAGGGCATACTGCAGACAGTGAAGGTTGTGGATGAGCTGGTGAAG*GTAAGTTCTGCATTTGGTTATGCTCCTTGCATTTTAGGTGTTCGTCGCACTTCCATTTCCATGAATAGCTAAGATTTTTTTTCTCTGCATTCATTCTTCTTGCCTCAGTTCTAACTGTTTGTGGTATTTTTGTTTTAATTATTGCTACAG*GTCATGGGCCGCCATAAGCCGGAGAACATCGTCATCGAGATGGCGAGGGAAAACCAGACGACTCAGAAAGGGCAGAAGAACTCACGGGAGCGCATGAAGCGGATAGAGGAAGGCATCAAGGAGCTTGGGAGTCAGATTCTGAAAGAGCACCCAGTCGAAAATACTCAACTCCAGAACGAGAAGCTGTACCTCTATTACCTCCAGAATGGGAGAGATATGTACGTCGACCAAGAGCTCGACATTAACAGACTCTCCGACTATGATGTGGATCACATTGTCCCTCAATCTTTCCTGAAGGACGATAGTATTGACAACAAG*GTAAAGCAACTGTGTTTTAATCAATTTCTTGTCAGGATATATGGATTATAACTTAATTTTTGAGAAATCTGTAGTATTTGGCGTGAAATGAGTTTGCTTTTTGGTTTCTCCCGTGTTATAG*GTCCTTACGCGCTCAGACAAGAACCGCGGAAAATCCGACAATGTACCCAGCGAGGAGGTTGTGAAGAAGATGAAGAACTATTGGAGGCAGCTTTTGAATGCTAAGCTCATAACCCAACGGAAATTCGACAATCTCACGAAGGCAGAAAGGGGCGGACTGTCTGAGCTCGACAAAGCCGGCTTCATCAAGCGCCAGTTGGTTGAAACTCGTCAGATTACGAAACATGTGGCCCAGATACTCGATTCGCGTATGAATACGAAGTATGATGAGAATGACAAACTTATCAGGGAGGTAAAG*GTAAAGTTTCCAACTTTCCTTTACCATATCAAACTAAAGTTCGAAACTTTTTATTTGATCAACTTCAAGGCCACCCGATCTTTCTATTCCTGATTAATTTGTGATGAATCCATATTGACTTTTGATGGTTACGCAG*GTGATCACCCTCAAGAGCAAACTGGTTAGTGACTTCCGGAAGGACTTCCAGTTTTACAAGGTTCGCGAGATCAACAACTACCATCATGCCCATGACGCCTACCTGAACGCCGTTGTTGGCACTGCTCTCATCAAGAAGTATCCGAAACTGGAGTCTGAGTTTGTGTACGGGGATTACAAGGTGTACGACGTTAGGAAGATGATCGCGAAGTCAGAACAAGAGATCGGCAAGGCTACCGCGAAATACTTCTTTTACTCGAATATCATGAACTTCTTCAAGACAGAGATCACTCTGGCGAATGGTGAAATCCGGAAGAGGCCTCTGATCGAGACAAATGGCGAAACAG*GTCTGTCTTTCCTATTTCATATGTTTAATCCTAGGAATTTGATCAATTGATTGTATGTATGTCGATCCCAAGACTTTCTTGTTCACTTATATCTTAACTCTCTCTTTGCTGTTTCTTGCAG*GTGAGATTGTCTGGGATAAGGGCAGGGATTTTGCGACTGTGCGTAAGGTTCTCAGCATGCCCCAAGTCAACATAGTCAAGAAAACGGAGGTTCAAACCGGTGGTTTCTCCAAGGAGTCCATTCTCCCTAAGCGCAACTCCGACAAACTGATTGCGAGGAAGAAGGATTGGGATCCGAAGAAATACGGAGGCTTTGATAGCCCTACCGTGGCATACAGCGTACTGGTAGTGGCCAAGGTGGAGAAGGGCAAGAGCAAGAAACTGAAAAGCGTCAAGGAACTGCTTGGAATTACCATAATGGAAAGGTCCTCGTTCGAGAAGAATCCGATCGACTTCCTCGAGGCTAAAG*GTAAAATATTGGATGCCAGACGATATTCTTTCTTTTGATTTGTAACTTTTTCCTGTCAAGGTCGATAAATTTTATTTTTTTTGGTAAAAGGTCGATAATTTTTTTTTGGAGCCATTATGTAATTTTCCTAATTAACTGAACCAAAATTATACTTTGCAG*GTTACAAAGAGGTGAAGAAAGACCTCATTATCAAACTGCCCAAGTATTCGCTTTTCGAATTGGAAAATGGCAGAAAACGCATGCTGGCATCTGCCGGAGAACTGCAGAAGGGCAACGAGCTGGCATTGCCCAGTAAGTACGTCAACTTCCTGTACTTGGCCTCACACTATGAGAAGCTGAAGGGGTCACCAGAGGACAACGAGCAGAAGCAGTTGTTTGTCGAGCAGCACAAGCACTATCTTGATGAGATCATAGAGCAGATCAGCGAATTTTCCAAGCGGGTCATTCTTGCAGACGCTAACCTCGATAAG*GTAAGGACTTCTCATGAATATTAGTGGCAGATTAGTGTTGTTAAAGTCTTTGGTTAGATAATCGATGCCTCCTAATTGTCCATGTTTTACTGGTTTTCTACAATTACAG*GTGCTTTCCGCGTACAACAAGCACAGAGATAAGCCGATAAGGGAACAAGCGGAAAACATCATCCACCTGTTCACACTGACCAATCTGGGAGCCCCAGCAGC

CTTTAAGTACTTCGATACCACTATCGACAGAAAGCGCTACACATCAACCAAGGAAGTGTTGGACGCTACCCTTATTC
ACCAATCTATTACAGGGCTCTATGAGACAAGGATAGATCTGTCGCAGTTGGGTGGTGACTCTAGGGCTGACCCAAAG
AAGAAGCGTAAAGTCGGTTCG

**pAGT6469 (Endonuclease: AATG_NLS-SpCas9i(D10A)-NLS_TTCG)**
GG-overhang-NLS-SpCas9i(D10A)(*introns*)-NLS-GG-overhang
AATGGCTTCTAGCCCACCGAAGAAGAAGCGGAAGGTCAGCTGGAAAATGGACAAGAAGTACAGCATTGGACTTGCAA
TTGGTACGAACTCAGTTGGGTGGGCCGTTATCACCGATGAATACAAGGTACCTTCGAAGAAATTTAAAGTGCTGGGC
AACACAGATAGGCACAGCATTAAGAAGAACTTGATCGGAGCTCTGCTCTTTGACTCTGGAGAAACCGCGGAGGCGAC
AAGGCTTAAACGTACTGCGAGGAGAAGGTACACTCGCAGGAAGAACAGAATCTGTTATCTCCAAGAGATCTTTAGCA
ACGAGATGGCGAAG*GTAAGGATTTTTATGATATACTATGCTTATGTATTTTGTACTGAAAGCATATCCTGCTTCATT*
*GGGATATTACTGAAAGCATTTAACTACATGTAAACTCACTTGATGATCAATAAACTTGATTTTGCAG*GTTGACGACT
CGTTCTTCCATCGCCTCGAGGAATCCTTCCTGGTAGAGGAAGATAAGAAACACGAGCGTCACCCCATCTTTGGGAAT
ATTGTTGACGAAGTAGCCTATCATGAAAAGTATCCGACTATATACCACCTTCGCAAGAAGCTGGTGGACTCAACCGA
TAAGGCAGACCTTCGGCTCATATACCTGGCTCTCGCGCACATGATAAAGTTTCGTGGCCATTTCTTGATCGAAGGGG
ACCTCAACCCGGATAACTCCGATGTGGATAAACTGTTCATTCAGCTCGTCCAAACCTACAATCAGCTGTTCGAGGAG
AACCCCATCAATGCATCAG*GTAACATTCCTTAGTTACCTTTCTTTTCTTTTTCCATCATAAGTTTATAGATTGTACA*
*TGCTTTGAGATTTTTCTTTGCAAACAATCTCAG*GTGTCGACGCCAAGGCAATACTGTCTGCCAGACTTTCGAAGTCC
AGACGGCTTGAGAATCTGATCGCTCAATTGCCAGGCGAGAAGAAGAACGGCTTGTTCGGGAATCTGATTGCACTGTC
TCTGGGCCTCACCCCTAACTTCAAAAGCAACTTTGACCTCGCCGAGGACGCGAAGCTGCAGCTGTCAAAGGATACAT
ACGATGATGATCTGGACAATCTGCTCGCCCAAATAG*GTAATCTTGAAATTGGAACTCTTCTTTTGTTGTCTAAACCT*
*ATCAATTTCTTTGCGGAAATTTATTTGAAGCTGTAGAGTTAAAATTGAGTCTTTTAAACTTTTGTAG*GTGATCAGTA
TGCCGACCTGTTCTTGGCTGCCAAGAATCTGTCAGACGCTATCTTGCTCAGTGACATTCTGCGGGTCAACACGGAGA
TAACCAAAGCGCCACTTAGCGCCTCCATGATCAAGAGGTACGACGAGCATCACCAGGATCTGACCCTTCTGAAGGCT
TTGGTTCGCCAGCAACTCCCCGAGAAGTACAAGGAGATTTTCTTTGACCAATCGAAGAATGGCTACGCAGGGTACAT
TGATGGAG*GTAAGTTGTTACTTATGATTGTTTTCCTCTCTGCTACATGTATTTTGTTGTTCATTTCTGTAAGATATA*
*AGAATTGAGTTTTCCTCTGATGATATTATTAG*GTGCAAGTCAGGAGGAATTCTACAAATTCATCAAGCCTATTCTGG
AAAAGATGGACGGTACAGAGGAGCTGCTCGTTAAATTGAACCGCGAAGATTTGCTTCGGAAGCAGCGTACCTTCGAC
AATGGCAGCATACCGCACCAGATCCACCTCGGTGAGCTGCATGCTATCTTGAGGAGGCAAGAGGACTTCTATCCGTT
CCTGAAAGACAACAGAGAGAAGATTGAAAAGATCCTCACGTTCCGCATTCCCTACTATGTAG*GTTAGTATCATATGA*
*AGAAATACCTAGTTTCAGTTGATGAATGCTATTTTCTGACCTCAGTTGTTCTCTTTTGAGAATTATTTCTTTTCTAA*
*TTTGCCTGATTTTTCTATTAATTCATTAG*GTCCACTCGCACGCGGGAACTCGCGGTTTGCGTGGATGACACGCAAAT
CCGAGGAGACTATCACGCCTTGGAACTTCGAAGAGGTCGTGGACAAGGGTGCGAGTGCACAGTCCTTCATCGAAAGG
ATGACCAACTTCGATAAGAATCTCCCAAATGAGAAAGTCCTGCCCAAGCATAGTCTCCTGTACGAATACTTCACGGT
CTACAACGAGCTGACGAAGGTGAAATATGTGACGGAGGGGATGCGCAAACCGGCCTTCCTGTCAG*GTAAATCCTGGT*
*CCACACTTTTACGATAAAAACACAAGATTTTAAACTATGAACTGATCAATAATCATTCCTAAAAGACCACACTTTTG*
*TTTTGTTTCTAAAGTAATTTTTACTGTTATAACAG*GTGAGCAGAAGAAGGCCATTGTCGATCTCTTGTTCAAAACCA
ATCGGAAGGTCACTGTGAAACAGCTTAAAGAGGACTACTTTAAGAAGATCGAATGCTTTGATTCTGTGGAAATCAGC
GGCGTTGAGGATAGGTTCAATGCCTCTCTTGGCACATACCATGACCTGTTGAAAATCATCAAGGACAAGGACTTCCT
TGACAACGAGGAGAACGAGGACATCCTCGAGGACATCGTGCTGACTCTCACGCTGTTTGAGGACAGAGAAATGATCG
AGGAGCGCCTTAAGACTTATGCGCATCTGTTCGATGACAAGGTCATGAAGCAGTTGAAGAGGAGGAGATATACAG*GT*
*AAGAGGTCAAAAGGTTTCCGCAATGATCCCTCTTTTTTTGTTTCTCTAGTTTCAAGAATTTGGGTATATGACTAACT*
*TCTGAGTGTTCCTTGATGCATATTTGTGATGAGACAAATGTTTGTTCTATGTTTTAG*GTTGGGGAAGGCTCTCCAGG
AAGCTCATCAACGGCATCCGCGACAAGCAATCCGGCAAGACTATACTGGACTTTCTCAAATCCGACGGTTTTGCGAA
TCGGAACTTCATGCAGCTTATTCACGATGACTCACTGACCTTCAAAGAAGATATCCAGAAGGCCCAAGTGTCAGGTC
AGGGCGATAGCCTTCACGAACACATAGCCAACCTGGCTGGATCGCCAGCTATAAAGAAGGGCATACTGCAGACAGTG
AAGGTTGTGGATGAGCTGGTGAAG*GTAAGTTCTGCATTTGGTTATGCTCCTTGCATTTTAGGTGTTCGTCGCACTTC*
*CATTTCCATGAATAGCTAAGATTTTTTTTCTCTGCATTCATTCTTCTTGCCTCAGTTCTAACTGTTTGTGGTATTTT*
*TGTTTTAATTATTGCTACAG*GTCATGGGCCGCCATAAGCCGGAGAACATCGTCATCGAGATGGCGAGGGAAAACCAG
ACGACTCAGAAAGGGCAGAAGAACTCACGGGAGCGCATGAAGCGGATAGAGGAAGGCATCAAGGAGCTTGGGAGTCA
GATTCTGAAAGAGCACCCAGTCGAAAATACTCAACTCCAGAACGAGAAGCTGTACCTCTATTACCTCCAGAATGGGA

GAGATATGTACGTCGACCAAGAGCTCGACATTAACAGACTCTCCGACTATGATGTGGATCACATTGTCCCTCAATCT
TTCCTGAAGGACGATAGTATTGACAACAAG*GTAAAGCAACTGTGTTTTAATCAATTTCTTGTCAGGATATATGGATT*
*ATAACTTAATTTTTGAGAAATCTGTAGTATTTGGCGTGAAATGAGTTTGCTTTTTGGTTTCTCCCGTGTTATAG*GTC
CTTACGCGCTCAGACAAGAACCGCGGAAAATCCGACAATGTACCCAGCGAGGAGGTTGTGAAGAAGATGAAGAACTA
TTGGAGGCAGCTTTTGAATGCTAAGCTCATAACCCAACGGAAATTCGACAATCTCACGAAGGCAGAAAGGGGCGGAC
TGTCTGAGCTCGACAAAGCCGGCTTCATCAAGCGCCAGTTGGTTGAAACTCGTCAGATTACGAAACATGTGGCCCAG
ATACTCGATTCGCGTATGAATACGAAGTATGATGAGAATGACAAACTTATCAGGGAGGTAAAG*GTAAAGTTTCCAAC*
*TTTCCTTTACCATATCAAACTAAAGTTCGAAACTTTTTATTTGATCAACTTCAAGGCCACCCGATCTTTCTATTCCT*
*GATTAATTTGTGATGAATCCATATTGACTTTTGATGGTTACGCAG*GTGATCACCCTCAAGAGCAAACTGGTTAGTGA
CTTCCGGAAGGACTTCCAGTTTTACAAGGTTCGCGAGATCAACAACTACCATCATGCCCATGACGCCTACCTGAACG
CCGTTGTTGGCACTGCTCTCATCAAGAAGTATCCGAAACTGGAGTCTGAGTTTGTGTACGGGGATTACAAGGTGTAC
GACGTTAGGAAGATGATCGCGAAGTCAGAACAAGAGATCGGCAAGGCTACCGCGAAATACTTCTTTTACTCGAATAT
CATGAACTTCTTCAAGACAGAGATCACTCTGGCGAATGGTGAAATCCGGAAGAGGCCTCTGATCGAGACAAATGGCG
AAACAG*GTCTGTCTTTCCTATTTCATATGTTTAATCCTAGGAATTTGATCAATTGATTGTATGTATGTCGATCCCAA*
*GACTTTCTTGTTCACTTATATCTTAACTCTCTCTTTGCTGTTTCTTGCAG*GTGAGATTGTCTGGGATAAGGGCAGGG
ATTTTGCGACTGTGCGTAAGGTTCTCAGCATGCCCCAAGTCAACATAGTCAAGAAAACGGAGGTTCAAACCGGTGGT
TTCTCCAAGGAGTCCATTCTCCCTAAGCGCAACTCCGACAAACTGATTGCGAGGAAGAAGGATTGGGATCCGAAGAA
ATACGGAGGCTTTGATAGCCCTACCGTGGCATACAGCGTACTGGTAGTGGCCAAGGTGGAGAAGGGCAAGAGCAAGA
AACTGAAAAGCGTCAAGGAACTGCTTGGAATTACCATAATGGAAAGGTCCTCGTTCGAGAAGAATCCGATCGACTTC
CTCGAGGCTAAAG*GTAAAATATTGGATGCCAGACGATATTCTTTCTTTTGATTTGTAACTTTTTCCTGTCAAGGTCG*
*ATAAATTTTATTTTTTTTGGTAAAAGGTCGATAATTTTTTTTTGGAGCCATTATGTAATTTTCCTAATTAACTGAAC*
*CAAAATTATACTTTGCAG*GTTACAAAGAGGTGAAGAAAGACCTCATTATCAAACTGCCCAAGTATTCGCTTTTCGAA
TTGGAAAATGGCAGAAAACGCATGCTGGCATCTGCCGGAGAACTGCAGAAGGGCAACGAGCTGGCATTGCCCAGTAA
GTACGTCAACTTCCTGTACTTGGCCTCACACTATGAGAAGCTGAAGGGGTCACCAGAGGACAACGAGCAGAAGCAGT
TGTTTGTCGAGCAGCACAAGCACTATCTTGATGAGATCATAGAGCAGATCAGCGAATTTTCCAAGCGGGTCATTCTT
GCAGACGCTAACCTCGATAAG*GTAAGGACTTCTCATGAATATTAGTGGCAGATTAGTGTTGTTAAAGTCTTTGGTTA*
*GATAATCGATGCCTCCTAATTGTCCATGTTTTACTGGTTTTCTACAATTACAG*GTGCTTTCCGCGTACAACAAGCAC
AGAGATAAGCCGATAAGGGAACAAGCGGAAAACATCATCCACCTGTTCACACTGACCAATCTGGGAGCCCCAGCAGC
CTTTAAGTACTTCGATACCACTATCGACAGAAAGCGCTACACATCAACCAAGGAAGTGTTGGACGCTACCCTTATTC
ACCAATCTATTACAGGGCTCTATGAGACAAGGATAGATCTGTCGCAGTTGGGTGGTGACTCTAGGGCTGACCCAAAG
AAGAAGCGTAAAGTCGGTTCG

**pAGT6470 (Endonuclease: AATG_NLS-SpCas9i(H840A)-NLS_TTCG)**
GG-overhang-NLS-SpCas9i(H840A)(*introns*)-NLS-GG-overhang
AATGGCTTCTAGCCCACCGAAGAAGAAGCGGAAGGTCAGCTGGAAAATGGACAAGAAGTACAGCATTGGACTTGATA
TTGGTACGAACTCAGTTGGGTGGGCCGTTATCACCGATGAATACAAGGTACCTTCGAAGAAATTTAAAGTGCTGGGC
AACACAGATAGGCACAGCATTAAGAAGAACTTGATCGGAGCTCTGCTCTTTGACTCTGGAGAAACCGCGGAGGCGAC
AAGGCTTAAACGTACTGCGAGGAGAAGGTACACTCGCAGGAAGAACAGAATCTGTTATCTCCAAGAGATCTTTAGCA
ACGAGATGGCGAAG*GTAAGGATTTTTATGATATACTATGCTTATGTATTTTGTACTGAAAGCATATCCTGCTTCATT*
*GGGATATTACTGAAAGCATTTAACTACATGTAAACTCACTTGATGATCAATAAACTTGATTTTGCAG*GTTGACGACT
CGTTCTTCCATCGCCTCGAGGAATCCTTCCTGGTAGAGGAAGATAAGAAACACGAGCGTCACCCCATCTTTGGGAAT
ATTGTTGACGAAGTAGCCTATCATGAAAAGTATCCGACTATATACCACCTTCGCAAGAAGCTGGTGGACTCAACCGA
TAAGGCAGACCTTCGGCTCATATACCTGGCTCTCGCGCACATGATAAAGTTTCGTGGCCATTTCTTGATCGAAGGGG
ACCTCAACCCGGATAACTCCGATGTGGATAAACTGTTCATTCAGCTCGTCCAAACCTACAATCAGCTGTTCGAGGAG
AACCCCATCAATGCATCAG*GTAACATTCCTTAGTTACCTTTCTTTTCTTTTTCCATCATAAGTTTATAGATTGTACA*
*TGCTTTGAGATTTTTCTTTGCAAACAATCTCAG*GTGTCGACGCCAAGGCAATACTGTCTGCCAGACTTTCGAAGTCC
AGACGGCTTGAGAATCTGATCGCTCAATTGCCAGGCGAGAAGAAGAACGGCTTGTTCGGGAATCTGATTGCACTGTC
TCTGGGCCTCACCCCTAACTTCAAAAGCAACTTTGACCTCGCCGAGGACGCGAAGCTGCAGCTGTCAAAGGATACAT
ACGATGATGATCTGGACAATCTGCTCGCCCAAATAG*GTAATCTTGAAATTGGAACTCTTCTTTTGTTGTCTAAACCT*
*ATCAATTTCTTTGCGGAAATTTATTTGAAGCTGTAGAGTTAAAATTGAGTCTTTTAAACTTTTGTAG*GTGATCAGTA
TGCCGACCTGTTCTTGGCTGCCAAGAATCTGTCAGACGCTATCTTGCTCAGTGACATTCTGCGGGTCAACACGGAGA

TAACCAAAGCGCCACTTAGCGCCTCCATGATCAAGAGGTACGACGAGCATCACCAGGATCTGACCCTTCTGAAGGCT
TTGGTTCGCCAGCAACTCCCCGAGAAGTACAAGGAGATTTTCTTTGACCAATCGAAGAATGGCTACGCAGGGTACAT
TGATGGAG*GTAAGTTGTTACTTATGATTGTTTTCCTCTCTGCTACATGTATTTTGTTGTTCATTTCTGTAAGATATA*
*AGAATTGAGTTTTCCTCTGATGATATTATTAG*GTGCAAGTCAGGAGGAATTCTACAAATTCATCAAGCCTATTCTGG
AAAAGATGGACGGTACAGAGGAGCTGCTCGTTAAATTGAACCGCGAAGATTTGCTTCGGAAGCAGCGTACCTTCGAC
AATGGCAGCATACCGCACCAGATCCACCTCGGTGAGCTGCATGCTATCTTGAGGAGGCAAGAGGACTTCTATCCGTT
CCTGAAAGACAACAGAGAGAAGATTGAAAAGATCCTCACGTTCCGCATTCCCTACTATGTAG*GTTAGTATCATATGA*
*AGAAATACCTAGTTTCAGTTGATGAATGCTATTTTCTGACCTCAGTTGTTCTCTTTTGAGAATTATTTCTTTTCTAA*
*TTTGCCTGATTTTTCTATTAATTCATTAG*GTCCACTCGCACGCGGGAACTCGCGGTTTGCGTGGATGACACGCAAAT
CCGAGGAGACTATCACGCCTTGGAACTTCGAAGAGGTCGTGGACAAGGGTGCGAGTGCACAGTCCTTCATCGAAAGG
ATGACCAACTTCGATAAGAATCTCCCAAATGAGAAAGTCCTGCCCAAGCATAGTCTCCTGTACGAATACTTCACGGT
CTACAACGAGCTGACGAAGGTGAAATATGTGACGGAGGGGATGCGCAAACCGGCCTTCCTGTCAG*GTAAATCCTGGT*
*CCACACTTTTACGATAAAAACACAAGATTTTAAACTATGAACTGATCAATAATCATTCCTAAAAGACCACACTTTTG*
*TTTTGTTTCTAAAGTAATTTTTACTGTTATAACAG*GTGAGCAGAAGAAGGCCATTGTCGATCTCTTGTTCAAAACCA
ATCGGAAGGTCACTGTGAAACAGCTTAAAGAGGACTACTTTAAGAAGATCGAATGCTTTGATTCTGTGGAAATCAGC
GGCGTTGAGGATAGGTTCAATGCCTCTCTTGGCACATACCATGACCTGTTGAAAATCATCAAGGACAAGGACTTCCT
TGACAACGAGGAGAACGAGGACATCCTCGAGGACATCGTGCTGACTCTCACGCTGTTTGAGGACAGAGAAATGATCG
AGGAGCGCCTTAAGACTTATGCGCATCTGTTCGATGACAAGGTCATGAAGCAGTTGAAGAGGAGGAGATATACAG*GT*
*AAGAGGTCAAAAGGTTTCCGCAATGATCCCTCTTTTTTTGTTTCTCTAGTTTCAAGAATTTGGGTATATGACTAACT*
*TCTGAGTGTTCCTTGATGCATATTTGTGATGAGACAAATGTTTGTTCTATGTTTTAG*GTTGGGGAAGGCTCTCCAGG
AAGCTCATCAACGGCATCCGCGACAAGCAATCCGGCAAGACTATACTGGACTTTCTCAAATCCGACGGTTTTGCGAA
TCGGAACTTCATGCAGCTTATTCACGATGACTCACTGACCTTCAAAGAAGATATCCAGAAGGCCCAAGTGTCAGGTC
AGGGCGATAGCCTTCACGAACACATAGCCAACCTGGCTGGATCGCCAGCTATAAAGAAGGGCATACTGCAGACAGTG
AAGGTTGTGGATGAGCTGGTGAAG*GTAAGTTCTGCATTTGGTTATGCTCCTTGCATTTTAGGTGTTCGTCGCACTTC*
*CATTTCCATGAATAGCTAAGATTTTTTTTCTCTGCATTCATTCTTCTTGCCTCAGTTCTAACTGTTTGTGGTATTTT*
*TGTTTTAATTATTGCTACAG*GTCATGGGCCGCCATAAGCCGGAGAACATCGTCATCGAGATGGCGAGGGAAAACCAG
ACGACTCAGAAAGGGCAGAAGAACTCACGGGAGCGCATGAAGCGGATAGAGGAAGGCATCAAGGAGCTTGGGAGTCA
GATTCTGAAAGAGCACCCAGTCGAAAATACTCAACTCCAGAACGAGAAGCTGTACCTCTATTACCTCCAGAATGGGA
GAGATATGTACGTCGACCAAGAGCTCGACATTAACAGACTCTCCGACTATGATGTGGATGCAATTGTCCCTCAATCT
TTCCTGAAGGACGATAGTATTGACAACAAG*GTAAAGCAACTGTGTTTTAATCAATTTCTTGTCAGGATATATGGATT*
*ATAACTTAATTTTTGAGAAATCTGTAGTATTTGGCGTGAAATGAGTTTGCTTTTTGGTTTCTCCCGTGTTATAG*GTC
CTTACGCGCTCAGACAAGAACCGCGGAAAATCCGACAATGTACCCAGCGAGGAGGTTGTGAAGAAGATGAAGAACTA
TTGGAGGCAGCTTTTGAATGCTAAGCTCATAACCCAACGGAAATTCGACAATCTCACGAAGGCAGAAAGGGGCGGAC
TGTCTGAGCTCGACAAAGCCGGCTTCATCAAGCGCCAGTTGGTTGAAACTCGTCAGATTACGAAACATGTGGCCCAG
ATACTCGATTCGCGTATGAATACGAAGTATGATGAGAATGACAAACTTATCAGGGAGGTAAAG*GTAAAGTTTCCAAC*
*TTTCCTTTACCATATCAAACTAAAGTTCGAAACTTTTTATTTGATCAACTTCAAGGCCACCCGATCTTTCTATTCCT*
*GATTAATTTGTGATGAATCCATATTGACTTTTGATGGTTACGCAG*GTGATCACCCTCAAGAGCAAACTGGTTAGTGA
CTTCCGGAAGGACTTCCAGTTTTACAAGGTTCGCGAGATCAACAACTACCATCATGCCCATGACGCCTACCTGAACG
CCGTTGTTGGCACTGCTCTCATCAAGAAGTATCCGAAACTGGAGTCTGAGTTTGTGTACGGGGATTACAAGGTGTAC
GACGTTAGGAAGATGATCGCGAAGTCAGAACAAGAGATCGGCAAGGCTACCGCGAAATACTTCTTTTACTCGAATAT
CATGAACTTCTTCAAGACAGAGATCACTCTGGCGAATGGTGAAATCCGGAAGAGGCCTCTGATCGAGACAAATGGCG
AAACAG*GTCTGTCTTTCCTATTTCATATGTTTAATCCTAGGAATTTGATCAATTGATTGTATGTATGTCGATCCCAA*
*GACTTTCTTGTTCACTTATATCTTAACTCTCTCTTTGCTGTTTCTTGCAG*GTGAGATTGTCTGGGATAAGGGCAGGG
ATTTTGCGACTGTGCGTAAGGTTCTCAGCATGCCCCAAGTCAACATAGTCAAGAAAACGGAGGTTCAAACCGGTGGT
TTCTCCAAGGAGTCCATTCTCCCTAAGCGCAACTCCGACAAACTGATTGCGAGGAAGAAGGATTGGGATCCGAAGAA
ATACGGAGGCTTTGATAGCCCTACCGTGGCATACAGCGTACTGGTAGTGGCCAAGGTGGAGAAGGGCAAGAGCAAGA
AACTGAAAAGCGTCAAGGAACTGCTTGGAATTACCATAATGGAAAGGTCCTCGTTCGAGAAGAATCCGATCGACTTC
CTCGAGGCTAAAG*GTAAAATATTGGATGCCAGACGATATTCTTTCTTTTGATTTGTAACTTTTTCCTGTCAAGGTCG*
*ATAAATTTTATTTTTTTTGGTAAAAGGTCGATAATTTTTTTTTGGAGCCATTATGTAATTTTCCTAATTAACTGAAC*
*CAAAATTATACTTTGCAG*GTTACAAAGAGGTGAAGAAAGACCTCATTATCAAACTGCCCAAGTATTCGCTTTTCGAA

TTGGAAAATGGCAGAAAACGCATGCTGGCATCTGCCGGAGAACTGCAGAAGGGCAACGAGCTGGCATTGCCCAGTAA
GTACGTCAACTTCCTGTACTTGGCCTCACACTATGAGAAGCTGAAGGGGTCACCAGAGGACAACGAGCAGAAGCAGT
TGTTTGTCGAGCAGCACAAGCACTATCTTGATGAGATCATAGAGCAGATCAGCGAATTTTCCAAGCGGGTCATTCTT
GCAGACGCTAACCTCGATAAG*GTAAGGACTTCTCATGAATATTAGTGGCAGATTAGTGTTGTTAAAGTCTTTGGTTA*
*GATAATCGATGCCTCCTAATTGTCCATGTTTTACTGGTTTTCTACAATTACAG*GTGCTTTCCGCGTACAACAAGCAC
AGAGATAAGCCGATAAGGGAACAAGCGGAAAACATCATCCACCTGTTCACACTGACCAATCTGGGAGCCCCAGCAGC
CTTTAAGTACTTCGATACCACTATCGACAGAAAGCGCTACACATCAACCAAGGAAGTGTTGGACGCTACCCTTATTC
ACCAATCTATTACAGGGCTCTATGAGACAAGGATAGATCTGTCGCAGTTGGGTGGTGACTCTAGGGCTGACCCAAAG
AAGAAGCGTAAAGTCGGTTCG

**pAGT6395 (Endonuclease: AATG_NLS-dSpCas9i(D10A; H840A)-NLS_TTCG)**
GG-overhang-NLS-dSpCas9i(D10A; H840A)(*introns*)-NLS-GG-overhang
AATGGCTTCTAGCCCACCGAAGAAGAAGCGGAAGGTCAGCTGGAAAATGGACAAGAAGTACAGCATTGGACTTGCAA
TTGGTACGAACTCAGTTGGGTGGGCCGTTATCACCGATGAATACAAGGTACCTTCGAAGAAATTTAAAGTGCTGGGC
AACACAGATAGGCACAGCATTAAGAAGAACTTGATCGGAGCTCTGCTCTTTGACTCTGGAGAAACCGCGGAGGCGAC
AAGGCTTAAACGTACTGCGAGGAGAAGGTACACTCGCAGGAAGAACAGAATCTGTTATCTCCAAGAGATCTTTAGCA
ACGAGATGGCGAAG*GTAAGGATTTTTATGATATACTATGCTTATGTATTTTGTACTGAAAGCATATCCTGCTTCATT*
*GGGATATTACTGAAAGCATTTAACTACATGTAAACTCACTTGATGATCAATAAACTTGATTTTGCAG*GTTGACGACT
CGTTCTTCCATCGCCTCGAGGAATCCTTCCTGGTAGAGGAAGATAAGAAACACGAGCGTCACCCCATCTTTGGGAAT
ATTGTTGACGAAGTAGCCTATCATGAAAAGTATCCGACTATATACCACCTTCGCAAGAAGCTGGTGGACTCAACCGA
TAAGGCAGACCTTCGGCTCATATACCTGGCTCTCGCGCACATGATAAAGTTTCGTGGCCATTTCTTGATCGAAGGGG
ACCTCAACCCGGATAACTCCGATGTGGATAAACTGTTCATTCAGCTCGTCCAAACCTACAATCAGCTGTTCGAGGAG
AACCCCATCAATGCATCAG*GTAACATTCCTTAGTTACCTTTCTTTTCTTTTTCCATCATAAGTTTATAGATTGTACA*
*TGCTTTGAGATTTTTCTTTGCAAACAATCTCAG*GTGTCGACGCCAAGGCAATACTGTCTGCCAGACTTTCGAAGTCC
AGACGGCTTGAGAATCTGATCGCTCAATTGCCAGGCGAGAAGAAGAACGGCTTGTTCGGGAATCTGATTGCACTGTC
TCTGGGCCTCACCCCTAACTTCAAAAGCAACTTTGACCTCGCCGAGGACGCGAAGCTGCAGCTGTCAAAGGATACAT
ACGATGATGATCTGGACAATCTGCTCGCCCAAATAG*GTAATCTTGAAATTGGAACTCTTCTTTTGTTGTCTAAACCT*
*ATCAATTTCTTTGCGGAAATTTATTTGAAGCTGTAGAGTTAAAATTGAGTCTTTTAAACTTTTGTAG*GTGATCAGTA
TGCCGACCTGTTCTTGGCTGCCAAGAATCTGTCAGACGCTATCTTGCTCAGTGACATTCTGCGGGTCAACACGGAGA
TAACCAAAGCGCCACTTAGCGCCTCCATGATCAAGAGGTACGACGAGCATCACCAGGATCTGACCCTTCTGAAGGCT
TTGGTTCGCCAGCAACTCCCCGAGAAGTACAAGGAGATTTTCTTTGACCAATCGAAGAATGGCTACGCAGGGTACAT
TGATGGAG*GTAAGTTGTTACTTATGATTGTTTTCCTCTCTGCTACATGTATTTTGTTGTTCATTTCTGTAAGATATA*
*AGAATTGAGTTTTCCTCTGATGATATTATTAG*GTGCAAGTCAGGAGGAATTCTACAAATTCATCAAGCCTATTCTGG
AAAAGATGGACGGTACAGAGGAGCTGCTCGTTAAATTGAACCGCGAAGATTTGCTTCGGAAGCAGCGTACCTTCGAC
AATGGCAGCATACCGCACCAGATCCACCTCGGTGAGCTGCATGCTATCTTGAGGAGGCAAGAGGACTTCTATCCGTT
CCTGAAAGACAACAGAGAGAAGATTGAAAAGATCCTCACGTTCCGCATTCCCTACTATGTAG*GTTAGTATCATATGA*
*AGAAATACCTAGTTTCAGTTGATGAATGCTATTTTCTGACCTCAGTTGTTCTCTTTTGAGAATTATTTCTTTTCTAA*
*TTTGCCTGATTTTTCTATTAATTCATTAG*GTCCACTCGCACGCGGGAACTCGCGGTTTGCGTGGATGACACGCAAAT
CCGAGGAGACTATCACGCCTTGGAACTTCGAAGAGGTCGTGGACAAGGGTGCGAGTGCACAGTCCTTCATCGAAAGG
ATGACCAACTTCGATAAGAATCTCCCAAATGAGAAAGTCCTGCCCAAGCATAGTCTCCTGTACGAATACTTCACGGT
CTACAACGAGCTGACGAAGGTGAAATATGTGACGGAGGGGATGCGCAAACCGGCCTTCCTGTCAG*GTAAATCCTGGT*
*CCACACTTTTACGATAAAAACACAAGATTTTAAACTATGAACTGATCAATAATCATTCCTAAAAGACCACACTTTTG*
*TTTTGTTTCTAAAGTAATTTTTACTGTTATAACAG*GTGAGCAGAAGAAGGCCATTGTCGATCTCTTGTTCAAAACCA
ATCGGAAGGTCACTGTGAAACAGCTTAAAGAGGACTACTTTAAGAAGATCGAATGCTTTGATTCTGTGGAAATCAGC
GGCGTTGAGGATAGGTTCAATGCCTCTCTTGGCACATACCATGACCTGTTGAAAATCATCAAGGACAAGGACTTCCT
TGACAACGAGGAGAACGAGGACATCCTCGAGGACATCGTGCTGACTCTCACGCTGTTTGAGGACAGAGAAATGATCG
AGGAGCGCCTTAAGACTTATGCGCATCTGTTCGATGACAAGGTCATGAAGCAGTTGAAGAGGAGGAGATATACAG*GT*
*AAGAGGTCAAAAGGTTTCCGCAATGATCCCTCTTTTTTTGTTTCTCTAGTTTCAAGAATTTGGGTATATGACTAACT*
*TCTGAGTGTTCCTTGATGCATATTTGTGATGAGACAAATGTTTGTTCTATGTTTTAG*GTTGGGGAAGGCTCTCCAGG
AAGCTCATCAACGGCATCCGCGACAAGCAATCCGGCAAGACTATACTGGACTTTCTCAAATCCGACGGTTTTGCGAA
TCGGAACTTCATGCAGCTTATTCACGATGACTCACTGACCTTCAAAGAAGATATCCAGAAGGCCCAAGTGTCAGGTC

AGGGCGATAGCCTTCACGAACACATAGCCAACCTGGCTGGATCGCCAGCTATAAAGAAGGGCATACTGCAGACAGTG
AAGGTTGTGGATGAGCTGGTGAAG*GTAAGTTCTGCATTTGGTTATGCTCCTTGCATTTTAGGTGTTCGTCGCACTTC*
*CATTTCCATGAATAGCTAAGATTTTTTTTCTCTGCATTCATTCTTCTTGCCTCAGTTCTAACTGTTTGTGGTATTTT*
*TGTTTTAATTATTGCTACAG*GTCATGGGCCGCCATAAGCCGGAGAACATCGTCATCGAGATGGCGAGGGAAAACCAG
ACGACTCAGAAAGGGCAGAAGAACTCACGGGAGCGCATGAAGCGGATAGAGGAAGGCATCAAGGAGCTTGGGAGTCA
GATTCTGAAAGAGCACCCAGTCGAAAATACTCAACTCCAGAACGAGAAGCTGTACCTCTATTACCTCCAGAATGGGA
GAGATATGTACGTCGACCAAGAGCTCGACATTAACAGACTCTCCGACTATGATGTGGATGCAATTGTCCCTCAATCT
TTCCTGAAGGACGATAGTATTGACAACAAG*GTAAAGCAACTGTGTTTTAATCAATTTCTTGTCAGGATATATGGATT*
*ATAACTTAATTTTTGAGAAATCTGTAGTATTTGGCGTGAAATGAGTTTGCTTTTTGGTTTCTCCCGTGTTATAG*GTC
CTTACGCGCTCAGACAAGAACCGCGGAAAATCCGACAATGTACCCAGCGAGGAGGTTGTGAAGAAGATGAAGAACTA
TTGGAGGCAGCTTTTGAATGCTAAGCTCATAACCCAACGGAAATTCGACAATCTCACGAAGGCAGAAAGGGGCGGAC
TGTCTGAGCTCGACAAAGCCGGCTTCATCAAGCGCCAGTTGGTTGAAACTCGTCAGATTACGAAACATGTGGCCCAG
ATACTCGATTCGCGTATGAATACGAAGTATGATGAGAATGACAAACTTATCAGGGAGGTAAAG*GTAAAGTTTCCAAC*
*TTTCCTTTACCATATCAAACTAAAGTTCGAAACTTTTTATTTGATCAACTTCAAGGCCACCCGATCTTTCTATTCCT*
*GATTAATTTGTGATGAATCCATATTGACTTTTGATGGTTACGCAG*GTGATCACCCTCAAGAGCAAACTGGTTAGTGA
CTTCCGGAAGGACTTCCAGTTTTACAAGGTTCGCGAGATCAACAACTACCATCATGCCCATGACGCCTACCTGAACG
CCGTTGTTGGCACTGCTCTCATCAAGAAGTATCCGAAACTGGAGTCTGAGTTTGTGTACGGGGATTACAAGGTGTAC
GACGTTAGGAAGATGATCGCGAAGTCAGAACAAGAGATCGGCAAGGCTACCGCGAAATACTTCTTTTACTCGAATAT
CATGAACTTCTTCAAGACAGAGATCACTCTGGCGAATGGTGAAATCCGGAAGAGGCCTCTGATCGAGACAAATGGCG
AAACAG*GTCTGTCTTTCCTATTTCATATGTTTAATCCTAGGAATTTGATCAATTGATTGTATGTATGTCGATCCCAA*
*GACTTTCTTGTTCACTTATATCTTAACTCTCTCTTTGCTGTTTCTTGCAG*GTGAGATTGTCTGGGATAAGGGCAGGG
ATTTTGCGACTGTGCGTAAGGTTCTCAGCATGCCCCAAGTCAACATAGTCAAGAAAACGGAGGTTCAAACCGGTGGT
TTCTCCAAGGAGTCCATTCTCCCTAAGCGCAACTCCGACAAACTGATTGCGAGGAAGAAGGATTGGGATCCGAAGAA
ATACGGAGGCTTTGATAGCCCTACCGTGGCATACAGCGTACTGGTAGTGGCCAAGGTGGAGAAGGGCAAGAGCAAGA
AACTGAAAAGCGTCAAGGAACTGCTTGGAATTACCATAATGGAAAGGTCCTCGTTCGAGAAGAATCCGATCGACTTC
CTCGAGGCTAAAG*GTAAAATATTGGATGCCAGACGATATTCTTTCTTTTGATTTGTAACTTTTTCCTGTCAAGGTCG*
*ATAAATTTTATTTTTTTTGGTAAAAGGTCGATAATTTTTTTTTGGAGCCATTATGTAATTTTCCTAATTAACTGAAC*
*CAAAATTATACTTTGCAG*GTTACAAAGAGGTGAAGAAAGACCTCATTATCAAACTGCCCAAGTATTCGCTTTTCGAA
TTGGAAAATGGCAGAAAACGCATGCTGGCATCTGCCGGAGAACTGCAGAAGGGCAACGAGCTGGCATTGCCCAGTAA
GTACGTCAACTTCCTGTACTTGGCCTCACACTATGAGAAGCTGAAGGGGTCACCAGAGGACAACGAGCAGAAGCAGT
TGTTTGTCGAGCAGCACAAGCACTATCTTGATGAGATCATAGAGCAGATCAGCGAATTTTCCAAGCGGGTCATTCTT
GCAGACGCTAACCTCGATAAG*GTAAGGACTTCTCATGAATATTAGTGGCAGATTAGTGTTGTTAAAGTCTTTGGTTA*
*GATAATCGATGCCTCCTAATTGTCCATGTTTTACTGGTTTTCTACAATTACAG*GTGCTTTCCGCGTACAACAAGCAC
AGAGATAAGCCGATAAGGGAACAAGCGGAAAACATCATCCACCTGTTCACACTGACCAATCTGGGAGCCCCAGCAGC
CTTTAAGTACTTCGATACCACTATCGACAGAAAGCGCTACACATCAACCAAGGAAGTGTTGGACGCTACCCTTATTC
ACCAATCTATTACAGGGCTCTATGAGACAAGGATAGATCTGTCGCAGTTGGGTGGTGACTCTAGGGCTGACCCAAAG
AAGAAGCGTAAAGTCGGTTCG

**pAGM47523 (Endonuclease: AATG_NLS-SpCas9i-NLS_GCTT)**
GG-overhang-NLS-SpCas9i(H840A)(*introns*)-NLS-GG-overhang
AATGGCTTCTAGCCCACCGAAGAAGAAGCGGAAGGTCAGCTGGAAAATGGACAAGAAGTACAGCATTGGACTTGATA
TTGGTACGAACTCAGTTGGGTGGGCCGTTATCACCGATGAATACAAGGTACCTTCGAAGAAATTTAAAGTGCTGGGC
AACACAGATAGGCACAGCATTAAGAAGAACTTGATCGGAGCTCTGCTCTTTGACTCTGGAGAAACCGCGGAGGCGAC
AAGGCTTAAACGTACTGCGAGGAGAAGGTACACTCGCAGGAAGAACAGAATCTGTTATCTCCAAGAGATCTTTAGCA
ACGAGATGGCGAAG*GTAAGGATTTTTATGATATAGTATGCTTATGTATTTTGTACTGAAAGCATATCCTGCTTCATT*
*GGGATATTACTGAAAGCATTTAACTACATGTAAACTCACTTGATGATCAATAAACTTGATTTTGCAG*GTTGACGACT
CGTTCTTCCATCGCCTCGAGGAATCCTTCCTGGTAGAGGAAGATAAGAAACACGAGCGTCACCCCATCTTTGGGAAT
ATTGTTGACGAAGTAGCCTATCATGAAAAGTATCCGACTATATACCACCTTCGCAAGAAGCTGGTGGACTCAACCGA
TAAGGCAGACCTTCGGCTCATATACCTGGCTCTCGCGCACATGATAAAGTTTCGTGGCCATTTCTTGATCGAAGGGG
ACCTCAACCCGGATAACTCCGATGTGGATAAACTGTTCATTCAGCTCGTCCAAACCTACAATCAGCTGTTCGAGGAG
AACCCCATCAATGCATCAG*GTAACATTCCTTAGTTACCTTTCTTTTCTTTTTCCATCATAAGTTTATAGATTGTACA*
*TGCTTTGAGATTTTTCTTTGCAAACAATCTCAG*GTGTCGACGCCAAGGCAATACTGTCTGCCAGACTTTCGAAGTCC

AGACGGCTTGAGAATCTGATCGCTCAATTGCCAGGCGAGAAGAAGAACGGCTTGTTCGGGAATCTGATTGCACTGTC
TCTGGGCCTCACCCCTAACTTCAAAAGCAACTTTGACCTCGCCGAGGACGCGAAGCTGCAGCTGTCAAAGGATACAT
ACGATGATGATCTGGACAATCTGCTCGCCCAAATAG*GTGCTCTTGAAATTGGAACTCTTCTTTTGTTGTCTAAACCT*
*ATCAATTTCTTTGCGGAAATTTATTTGAAGCTGTAGAGTTAAAATTGAGTCTTTTAAACTTTTGTAG*GTGATCAGTA
TGCCGACCTGTTCTTGGCTGCCAAGAATCTGTCAGACGCTATCTTGCTCAGTGACATTCTGCGGGTCAACACGGAGA
TAACCAAAGCGCCACTTAGCGCCTCCATGATCAAGAGGTACGACGAGCATCACCAGGATCTGACCCTTCTGAAGGCT
TTGGTTCGCCAGCAACTCCCCGAGAAGTACAAGGAGATTTTCTTTGACCAATCGAAGAATGGCTACGCAGGGTACAT
TGATGGAG*GTAAGTTGTTACTTATGATTGTTTTCCTCTCTGCTACATGTATTTTGTTGTTCATTTCTGTAAGATATA*
*AGAATTGAGTTTTCCTCTGATGATATTATTAG*GTGCAAGTCAGGAGGAATTCTACAAATTCATCAAGCCTATTCTGG
AAAAGATGGACGGTACAGAGGAGCTGCTCGTTAAATTGAACCGCGAAGATTTGCTTCGGAAGCAGCGTACCTTCGAC
AATGGCAGCATACCGCACCAGATCCACCTCGGTGAGCTGCATGCTATCTTGAGGAGGCAAGAGGACTTCTATCCGTT
CCTGAAAGACAACAGAGAGAAGATTGAAAAGATCCTCACGTTCCGCATTCCCTACTATGTAG*GTTAGTATCATATGA*
*AGAAATACCTAGTTTCAGTTGATGAATGCTATTTTCTGACCTCAGTTGTTCTCTTTTGAGAATTATTTCTTTTCTAA*
*TTTGCCTGATTTTTCTATTAATTCATTAG*GTCCACTCGCACGCGGGAACTCGCGGTTTGCGTGGATGACACGCAAAT
CCGAGGAGACTATCACGCCTTGGAACTTCGAAGAGGTCGTGGACAAGGGTGCGAGTGCACAGTCCTTCATCGAAAGG
ATGACCAACTTCGATAAGAATCTCCCAAATGAGAAAGTCCTGCCCAAGCATAGTCTCCTGTACGAATACTTCACGGT
CTACAACGAGCTGACGAAGGTGAAATATGTGACGGAGGGGATGCGCAAACCGGCCTTCCTGTCAG*GTAAATCCTGGT*
*CCACACTTTTACGATAAAAACACAAGATTTTAAACTATGAACTGATCAATAATCATTCCTAAAAGACCACACTTTTG*
*TTTTGTTTCTAAAGTAATTTTTACTGTTATAACAG*GTGAGCAGAAGAAGGCCATTGTCGATCTCTTGTTCAAAACCA
ATCGGAAGGTCACTGTGAAACAGCTTAAAGAGGACTACTTTAAGAAGATCGAATGCTTTGATTCTGTGGAAATCAGC
GGCGTTGAGGATAGGTTCAATGCCTCTCTTGGCACATACCATGACCTGTTGAAAATCATCAAGGACAAGGACTTCCT
TGACAACGAGGAGAACGAGGACATCCTCGAGGACATCGTGCTGACTCTCACGCTGTTTGAGGACAGAGAAATGATCG
AGGAGCGCCTTAAGACTTATGCGCATCTGTTCGATGACAAGGTCATGAAGCAGTTGAAGAGGAGGAGATATACAG*GT*
*AAGAGGTCAAAAGGTTTCCGCAATGATCCCTCTTTTTTTGTTTCTCTAGTTTCAAGAATTTGGGTATATGACTAACT*
*TCTGAGTGTTCCTTGATGCATATTTGTGATGAGACAAATGTTTGTTCTATGTTTTAG*GTTGGGGAAGGCTCTCCAGG
AAGCTCATCAACGGCATCCGCGACAAGCAATCCGGCAAGACTATACTGGACTTTCTCAAATCCGACGGTTTTGCGAA
TCGGAACTTCATGCAGCTTATTCACGATGACTCACTGACCTTCAAAGAAGATATCCAGAAGGCCCAAGTGTCAGGTC
AGGGCGATAGCCTTCACGAACACATAGCCAACCTGGCTGGATCGCCAGCTATAAAGAAGGGCATACTGCAGACAGTG
AAGGTTGTGGATGAGCTGGTGAAG*GTAAGTTCTGCATTTGGTTATGCTCCTTGCATTTTAGGTGTTCGTCGCACTTC*
*CATTTCCATGAATAGCTAAGATTTTTTTTCTCTGCATTCATTCTTCTTGCCTCAGTTCTAACTGTTTGTGGTATTTT*
*TGTTTTAATTATTGCTACAG*GTCATGGGCCGCCATAAGCCGGAGAACATCGTCATCGAGATGGCGAGGGAAAACCAG
ACGACTCAGAAAGGGCAGAAGAACTCACGGGAGCGCATGAAGCGGATAGAGGAAGGCATCAAGGAGCTTGGGAGTCA
GATTCTGAAAGAGCACCCAGTCGAAAATACTCAACTCCAGAACGAGAAGCTGTACCTCTATTACCTCCAGAATGGGA
GAGATATGTACGTCGACCAAGAGCTCGACATTAACAGACTCTCCGACTATGATGTGGATCACATTGTCCCTCAATCT
TTCCTGAAGGACGATAGTATTGACAACAAG*GTAAAGCAACTGTGTTTTAATCAATTTCTTGTCAGGATATATGGATT*
*ATAACTTAATTTTTGAGAAATCTGTAGTATTTGGCGTGAAATGAGTTTGCTTTTTGGTTTCTCCCGTGTTATAG*GTC
CTTACGCGCTCAGACAAGAACCGCGGAAAATCCGACAATGTACCCAGCGAGGAGGTTGTGAAGAAGATGAAGAACTA
TTGGAGGCAGCTTTTGAATGCTAAGCTCATAACCCAACGGAAATTCGACAATCTCACGAAGGCAGAAAGGGGCGGAC
TGTCTGAGCTCGACAAAGCCGGCTTCATCAAGCGCCAGTTGGTTGAAACTCGTCAGATTACGAAACATGTGGCCCAG
ATACTCGATTCGCGTATGAATACGAAGTATGATGAGAATGACAAACTTATCAGGGAGGTAAAG*GTAAAGTTTCCAAC*
*TTTCCTTTACCATATCAAACTAAAGTTCGAAACTTTTTATTTGATCAACTTCAAGGCCACCCGATCTTTCTATTCCT*
*GATTAATTTGTGATGAATCCATATTGACTTTTGATGGTTACGCAG*GTGATCACCCTCAAGAGCAAACTGGTTAGTGA
CTTCCGGAAGGACTTCCAGTTTTACAAGGTTCGCGAGATCAACAACTACCATCATGCCCATGACGCCTACCTGAACG
CCGTTGTTGGCACTGCTCTCATCAAGAAGTATCCGAAACTGGAGTCTGAGTTTGTGTACGGGGATTACAAGGTGTAC
GACGTTAGGAAGATGATCGCGAAGTCAGAACAAGAGATCGGCAAGGCTACCGCGAAATACTTCTTTTACTCGAATAT
CATGAACTTCTTCAAGACAGAGATCACTCTGGCGAATGGTGAAATCCGGAAGAGGCCTCTGATCGAGACAAATGGCG
AAACAG*GTCTGTCTTTCCTATTTCATATGTTTAATCCTAGGAATTTGATCAATTGATTGTATGTATGTCGATCCCAA*
*GACTTTCTTGTTCACTTATATCTTAACTCTCTCTTTGCTGTTTCTTGCAG*GTGAGATTGTCTGGATAAGGGCAGGG
ATTTTGCGACTGTGCGTAAGGTTCTCAGCATGCCCCAAGTCAACATAGTCAAGAAAACGGAGGTTCAAACCGGTGGT
TTCTCCAAGGAGTCCATTCTCCCTAAGCGCAACTCCGACAAACTGATTGCGAGGAAGAAGGATTGGGATCCGAAGAA
ATACGGAGGCTTTGATAGCCCTACCGTGGCATACAGCGTACTGGTAGTGGCCAAGGTGGAGAAGGGCAAGAGCAAGA
AACTGAAAAGCGTCAAGGAACTGCTTGGAATTACCATAATGGAAAGGTCCTCGTTCGAGAAGAATCCGATCGACTTC
CTCGAGGCTAAAG*GTAAAATATTGGATGCCAGACGATATTCTTTCTTTTGATTTGTAACTTTTTCCTGTCAAGGTCG*
*ATAAATTTTATTTTTTTTGGTAAAAGGTCGATAATTTTTTTTTGGAGCCATTATGTAATTTTCCTAATTAACTGAAC*
*CAAAATTATACAAACCAG*GTTACAAAGAGGTGAAGAAAGACCTCATTATCAAACTGCCCAAGTATTCGCTTTTCGAA
TTGGAAAATGGCAGAAAACGCATGCTGGCATCTGCCGGAGAACTGCAGAAGGGCAACGAGCTGGCATTGCCCAGTAA
GTACGTCAACTTCCTGTACTTGGCCTCACACTATGAGAAGCTGAAGGGGTCACCAGAGGACAACGAGCAGAAGCAGT
TGTTTGTCGAGCAGCACAAGCACTATCTTGATGAGATCATAGAGCAGATCAGCGAATTTTCCAAGCGGGTCATTCTT

GCAGACGCTAACCTCGATAAG*GTAAGGACTTCTCATGAATATTAGTGGCAGATTAGTGTTGTTAAAGTCTTTGGTTA*
*GATAATCGATGCCTCCTAATTGTCCATGTTTTACTGGTTTTCTACAATTAAAG*GTGCTTTCCGCGTACAACAAGCAC
AGAGATAAGCCGATAAGGAACAAGCGGAAAACATCATCCACCTGTTCACACTGACCAATCTGGGAGCCCCAGCAGC
CTTTAAGTACTTCGATACCACTATCGACAGAAAGCGCTACACATCAACCAAGGAAGTGTTGGACGCTACCCTTATTC
ACCAATCTATTACAGGGCTCTATGAGACAAGGATAGATCTGTCGCAGTTGGGTGGTGACTCTAGGGCTGACCCAAAG
AAGAAGCGTAAAGTCTTGAGCTT

**pAGT8163 (Endonuclease: AATG_NLS-ttLbCas12a-i-NLS_TTCG)**
GG-overhang-NLS-ttLbCas12a-i(*introns*)-NLS-GG-overhang
AATGCTGCAGCCTAAGAAGAAGAGAAAGGTTGGAGGAGTCGACTCGAGTGCGGCCGCCACAATGAGCAAGCTCGAGA
AGTTTACCAACTGCTACAGCCTGTCTAAGACCCTGAGGTTCAAGGCTATTCCTGTGGGTAAGACCCAAGAGAATATC
GACAACAAGCGGCTGCTGGTTGAGGATGAGAAGAGAGCTGAGGATTACAAGGGCGTGAAGAAGCTGCTGGATCGGTA
CTACCTGAGCTTCATCAACGATGTGCTGCACAGCATCAAGCTGAAGAACCTGAACAACTACATCAGCCTGTTCCGGA
AGAAAACCCGGACCGAGAAAGAGAACAAAGAGCTTGAGAACCTCGAGATCAACCTGCGGAAAGAGATCGCTAAGGCT
TTCAAGGGTAACGAAG*GTAAGGATTTTTATGATATACTATGCTTATGTATTTTGTACTGAAAGCATATCCTGCTTCA*
*TTGGGATATTACTGAAAGCATTTAACTACATGTAAACTCACTTGATGATCAATAAACTTGATTTTGCAG*GTTACAAG
AGCCTGTTCAAGAAGGATATTATCGAGACTATCCTGCCTGAGTTCCTGGACGATAAGGATGAGATTGCCCTGGTGAA
CAGCTTCAACGGTTTCACTACTGCCTTCACCGGTTTCTTCAGAAACCGGGAAAACATGTTCAGCGAAGAGGCCAAGT
CTACCTCTATCGCTTTCCGGTGCATTAACGAGAACTTGACCCGGTACATCAGCAACATGGACATCTTCGAGAAGGTG
GACGCCATCTTCGATAAGCACGAGGTGCAAGAAATCAAAGAGAAGATCCTGAACTCCGACTACGACGTCGAGGATTT
TTTTGAGGGCGAGTTCTTCAACTTCGTGCTCACCCAAGAAG*GTAACATTCCTTAGTTACCTTTCTTTTCTTTTTCCA*
*TCATAAGTTTATAGATTGTACATGCTTTGAGATTTTTCTTTGCAAACAATCTCAG*GTATCGATGTGTACAACGCTAT
CATCGGTGGTTTCGTGACTGAGAGCGGTGAGAAGATTAAGGGCCTGAACGAGTACATTAACCTGTACAATCAAAAGA
CCAAGCAGAAGCTGCCGAAGTTCAAGCCGCTTTACAAGCAGGTTCTGAGCGATCGTGAGAGCCTGTCTTTTTACGGA
GAGGGATACACCTCTGATGAAGAGGTTTTGGAG*GTAATCTTGAAATTGGAACTCTTCTTTTGTTGTCTAAACCTATC*
*AATTTCTTTGCGGAAATTTATTTGAAGCTGTAGAGTTAAAATTGAGTCTTTTAAACTTTTGTAG*GTGTTCCGTAACA
CCCTGAACAAGAACAGCGAGATCTTCAGCTCCATCAAGAAGCTGGAAAAGCTGTTTAAGAACTTCGACGAGTACAGC
AGCGCTGGCATCTTCGTTAAGAACGGTCCTGCTATCAGCACCATCAGCAAGGATATTTTCGGCGAGTGGAACGTGAT
CCGGGATAAGTGGAATGCTGAGTACGATGACATCCACCTGAAGAAAAAGGCTGTGGTGACCGAGAAGTACGAGGATG
ATAGGCGGAAGTCCTTCAAGAAGATAG*GTAAGTTGTTACTTATGATTGTTTTCCTCTCTGCTACATGTATTTTGTTG*
*TTCATTTCTGTAAGATATAAGAATTGAGTTTTCCTCTGATGATATTATTAG*GTTCCTTTAGCCTCGAGCAGCTTCAA
GAGTATGCTGACGCTGATCTGTCCGTGGTCGAGAAGCTTAAAGAGATCATCATCCAGAAGGTCGACGAGATCTACAA
GGTGTACGGCAGCTCTGAGAAGCTTTTCGATGCTGACTTCGTGTTGGAGAAGTCTCTGAAGAAGAACGACGCCGTTG
TCGCTATCATGAAGGATCTGCTGGACAGCGTGAAGTCTTTCGAGAACTATATCAAGGCCTTCTTCGGCGAAG*GTTAG*
*TATCATATGAAGAAATACCTAGTTTCAGTTGATGAATGCTATTTTCTGACCTCAGTTGTTCTCTTTTGAGAATTATT*
*TCTTTTCTAATTTGCCTGATTTTTCTATTAATTCATTAG*GTAAAGAGACTAATAGGGACGAGTCATTCTACGGCGAT
TTCGTGCTGGCTTACGACATCCTTCTTAAGGTGGACCACATCTACGACGCCATCAGAAATTACGTGACCCAGAAGCC
GTACAGCAAGGACAAGTTCAAGTTGTACTTCCAGAATCCGCAGTTCATGGGCGGCTGGGACAAAGACAAAGAGACAG
ATTACAGGGCTACCATCCTGCGGTACGGCTCTAAGTACTACCTTGCCATCATGGACAAGAAATACGCCAAGTGCCTG
CAAAAGATCGACAAGGATGATGTGAACGGCAACTACGAGAAGATCAACTACAAGCTCCTGCCAG*GTAAATCCTGGTC*
*CACACTTTTACGATAAAAACACAAGATTTTAAACTATGAACTGATCAATAATCATTCCTAAAAGACCACACTTTTGT*
*TTTGTTTCTAAAGTAATTTTTACTGTTATAACAG*GTCCTAACAAGATGCTTCCTAAGGTGTTCTTCTCAAAGAAATG
GATGGCCTACTACAACCCGAGCGAGGACATCCAGAAAATCTACAAGAACGGCACCTTCAAAAAGGGCGACATGTTCA
ACCTGAACGACTGCCACAAGCTGATCGATTTCTTCAAGGACAGCATCAGCCGGTATCCGAAGTGGTCTAACGCTTAC
GATTTCAACTTCAGCGAGACTGAGAAGTATAAGGATATCGCCGGCTTCTACCGTGAGGTTGAGGAACAGGGTTACAA
GGTTAGCTTCGAGAGCGCCAGCAAGAAAGAGGTGGACAAGTTGGTTGAAGAAG*GTAAGAGGTCAAAAGGTTTCCGCA*
*ATGATCCCTCTTTTTTTGTTTCTCTAGTTTCAAGAATTTGGGTATATGACTAACTTCTGAGTGTTCCTTGATGCATA*
*TTTGTGATGAGACAAATGTTTGTTCTATGTTTTAG*GTAAGCTGTACATGTTCCAAATCTATAACAAGGACTTCTCCG
ACAAGTCTCACGGCACTCCTAATCTGCATACAATGTACTTCAAGCTGCTGTTCGACGAGAACAACCACGGTCAGATT
AGGCTTTCTGGTGGTGCTGAGCTGTTCATGAGAAGGGCCTCACTGAAGAAAGAAGAGTTGGTCGTTCACCCTGCCAA
CTCTCCAATCGCTAACAAGAACCCTGACAACCCGAAAAAGACCACCACCTTGTCTTACGACGTGTACAAGGATAAGC
GGTTCAGCGAGGATCAGTACGAGCTTCACATTCCGATCGCCATCAACAAGTGCCCGAAGAACATCTTCAAGATCAAT
ACCGAGGTGCGGGTGCTGCTGAAGCACGATGATAATCCTTACGTGATCGGCATCGATAGGGGCGAGAGAAACCTTCT
TTACATCGTGGTGGTGGACGGCAAGGGCAATATCGTTGAGCAGTACTCTCTGAACGAGATTATCAACAATTTCAACG
GCATCCGGATCAAGACCGACTACCACTCTCTGCTGGATAAGAAAGAAAAAGAGCGGTTCGAGGCCAGGCAGAACTGG
ACTTCTATCGAAAACATCAAAGAGCTGAAGGCCGGCTACATCTCTCAGGTGGTGCATAAGATTTGCGAGCTGGTGGA
AAAGTACGACGCTGTGATTGCTCTCGAGGATCTGAACAGCGGCTTCAAGAACTCACGTGTGAAG*GTAAAGCAACTGT*

GTTTTAATCAATTTCTTGTCAGGATATATGGATTATAACTTAATTTTTGAGAAATCTGTAGTATTTGGCGTGAAATG
AGTTTGCTTTTTGGTTTCTCCCGTGTTATAGGTTGAGAAGCAGGTCTACCAAAAGTTCGAGAAGATGCTCATCGACA
AGCTGAACTACATGGTGGACAAAAAGAGCAACCCTTGCGCTACCGGTGGTGCTCTTAAGGGTTACCAGATCACTAAC
AAGTTCGAGTCTTTCAAGAGCATGAGCACCCAGAACGGCTTCATCTTCTACATCCCTGCTTGGCTGACCAGCAAGAT
CGATCCTTCTACTGGCTTCGTCAACCTGCTCAAGACCAAGTACACCAGCATTGCCGACAGCAAGAAGTTCATCAGCT
CATTCGACCGGATCATGTACGTGCCAGAAGAGGATCTTTTCGAGTTCGCCCTCGATTACAAGAACTTCTCTAGGACC
GACGCCGACTACATTAAGAAGTGGAAGCTGTACTCCTACGGCAACCGGATTCGGATCTTTCGGAACCCGAAGAAAAA
CAACGTGTTCGACTGGGAAGAGGTAAAGTTTCCAACTTTCCTTTACCATATCAAACTATAGTTCGAAACTTTTTATT
TGATCAACTTCAAGGCCACCCGATCTTTCTATTCCTGATTAATTTGTGATGAATCCATATTGACTTTTGATGGTTAC
GCAGGTGTGCCTGACCTCTGCCTACAAAGAACTGTTCAACAAGTACGGCATCAACTACCAGCAGGGTGATATTAGGG
CTCTGCTTTGCGAGCAGTCTGACAAGGCTTTCTACAGCTCTTTCATGGCCCTGATGTCTCTGATGCTGCAAATGAGG
AACTCTATCACCGGTAGGACCGATGTGGACTTCCTTATCTCTCCGGTGAAGAACAGTGACGGGATCTTCTACGACAG
CCGGAATTATGAGGCTCAAGAGAACGCAATCCTGCCGAAGAATGCTGATGCTAACGGCGCTTACAACATTGCCAGAA
AGGTGCTGTGGGCTATCGGCCAGTTTAAGAAAGCCGAAGATGAGAAGTTGGACAAGGTCTGTCTTTCCTATTTCATA
TGTTTAATCCTAGGAATTTGATCAATTGATTGTATGTATGTCGATCCCAAGACTTTCTTGTTCACTTATATCTTAAC
TCTCTCTTTGCTGTTTCTTGCAGGTGAAGATCGCTATCTCCAACAAAGAGTGGCTCGAGTACGCTCAGACTAGCGTT
AAGCATAAAAGGCCGGCGGCCACGAAAAAGGCCGGCCAGGCAAAAAAGAAAAAGGGTTCG

**pAGT8162 (Endonuclease: AATG_NLS-dttLbCas12a(E925Q)-i-NLS_TTCG)**

GG-overhang-NLS-ttLbCas12a-i(E925Q)(*introns*)-NLS-GG-overhang

AATGCTGCAGCCTAAGAAGAAGAGAAAGGTTGGAGGAGTCGACTCGAGTGCGGCCGCCACAATGAGCAAGCTCGAGA
AGTTTACCAACTGCTACAGCCTGTCTAAGACCCTGAGGTTCAAGGCTATTCCTGTGGGTAAGACCCAAGAGAATATC
GACAACAAGCGGCTGCTGGTTGAGGATGAGAAGAGAGCTGAGGATTACAAGGGCGTGAAGAAGCTGCTGGATCGGTA
CTACCTGAGCTTCATCAACGATGTGCTGCACAGCATCAAGCTGAAGAACCTGAACAACTACATCAGCCTGTTCCGGA
AGAAAACCCGGACCGAGAAAGAGAACAAAGAGCTTGAGAACCTCGAGATCAACCTGCGGAAAGAGATCGCTAAGGCT
TTCAAGGGTAACGAAGGTAAGGATTTTTATGATATACTATGCTTATGTATTTTGTACTGAAAGCATATCCTGCTTCA
TTGGGATATTACTGAAAGCATTTAACTACATGTAAACTCACTTGATGATCAATAAACTTGATTTTGCAGGTTACAAG
AGCCTGTTCAAGAAGGATATTATCGAGACTATCCTGCCTGAGTTCCTGGACGATAAGGATGAGATTGCCCTGGTGAA
CAGCTTCAACGGTTTCACTACTGCCTTCACCGGTTTCTTCAGAAACCGGGAAAACATGTTCAGCGAAGAGGCCAAGT
CTACCTCTATCGCTTTCCGGTGCATTAACGAGAACTTGACCCGGTACATCAGCAACATGGACATCTTCGAGAAGGTG
GACGCCATCTTCGATAAGCACGAGGTGCAAGAAATCAAAGAGAAGATCCTGAACTCCGACTACGACGTCGAGGATTT
TTTTGAGGGCGAGTTCTTCAACTTCGTGCTCACCCAAGAAGGTAACATTCCTTAGTTACCTTTCTTTTCTTTTTCCA
TCATAAGTTTATAGATTGTACATGCTTTGAGATTTTTCTTTGCAAACAATCTCAGGTATCGATGTGTACAACGCTAT
CATCGGTGGTTTCGTGACTGAGAGCGGTGAGAAGATTAAGGGCCTGAACGAGTACATTAACCTGTACAATCAAAAGA
CCAAGCAGAAGCTGCCGAAGTTCAAGCCGCTTTACAAGCAGGTTCTGAGCGATCGTGAGAGCCTGTCTTTTTACGGA
GAGGGATACACCTCTGATGAAGAGGTTTTGGAGGTAATCTTGAAATTGGAACTCTTCTTTTGTTGTCTAAACCTATC
AATTTCTTTGCGGAAATTTATTTGAAGCTGTAGAGTTAAAATTGAGTCTTTTAAACTTTTGTAGGTGTTCCGTAACA
CCCTGAACAAGAACAGCGAGATCTTCAGCTCCATCAAGAAGCTGGAAAAGCTGTTTAAGAACTTCGACGAGTACAGC
AGCGCTGGCATCTTCGTTAAGAACGGTCCTGCTATCAGCACCATCAGCAAGGATATTTTCGGCGAGTGGAACGTGAT
CCGGGATAAGTGGAATGCTGAGTACGATGACATCCACCTGAAGAAAAAGGCTGTGGTGACCGAGAAGTACGAGGATG
ATAGGCGGAAGTCCTTCAAGAAGATAGGTAAGTTGTTACTTATGATTGTTTTCCTCTCTGCTACATGTATTTTGTTG
TTCATTTCTGTAAGATATAAGAATTGAGTTTTCCTCTGATGATATTATTAGGTTCCTTTAGCCTCGAGCAGCTTCAA
GAGTATGCTGACGCTGATCTGTCCGTGGTCGAGAAGCTTAAAGAGATCATCATCCAGAAGGTCGACGAGATCTACAA
GGTGTACGGCAGCTCTGAGAAGCTTTTCGATGCTGACTTCGTGTTGGAGAAGTCTCTGAAGAAGAACGACGCCGTTG
TCGCTATCATGAAGGATCTGCTGGACAGCGTGAAGTCTTTCGAGAACTATATCAAGGCCTTCTTCGGCGAAGGTTAG
TATCATATGAAGAAATACCTAGTTTCAGTTGATGAATGCTATTTTCTGACCTCAGTTGTTCTCTTTTGAGAATTATT
TCTTTTCTAATTTGCCTGATTTTTCTATTAATTCATTAGGTAAAGAGACTAATAGGGACGAGTCATTCTACGGCGAT
TTCGTGCTGGCTTACGACATCCTTCTTAAGGTGGACCACATCTACGACGCCATCAGAAATTACGTGACCCAGAAGCC
GTACAGCAAGGACAAGTTCAAGTTGTACTTCCAGAATCCGCAGTTCATGGGCGGCTGGGACAAAGACAAAGAGACAG
ATTACAGGGCTACCATCCTGCGGTACGGCTCTAAGTACTACCTTGCCATCATGGACAAGAAATACGCCAAGTGCCTG
CAAAAGATCGACAAGGATGATGTGAACGGCAACTACGAGAAGATCAACTACAAGCTCCTGCCAGGTAAATCCTGGTC
CACACTTTTACGATAAAAACACAAGATTTTAAACTATGAACTGATCAATAATCATTCCTAAAAGACCACACTTTTGT
TTTGTTTCTAAAGTAATTTTTACTGTTATAACAGGTCCTAACAAGATGCTTCCTAAGGTGTTCTTCTCAAAGAAATG
GATGGCCTACTACAACCCGAGCGAGGACATCCAGAAAATCTACAAGAACGGCACCTTCAAAAAGGGCGACATGTTCA
ACCTGAACGACTGCCACAAGCTGATCGATTTCTTCAAGGACAGCATCAGCCGGTATCCGAAGTGGTCTAACGCTTAC
GATTTCAACTTCAGCGAGACTGAGAAGTATAAGGATATCGCCGGCTTCTACCGTGAGGTTGAGGAACAGGGTTACAA
GGTTAGCTTCGAGAGCGCCAGCAAGAAAGAGGTGGACAAGTTGGTTGAAGAAGGTAAGAGGTCAAAAGGTTTCCGCA
ATGATCCCTCTTTTTTTGTTTCTCTAGTTTCAAGAATTTGGGTATATGACTAACTTCTGAGTGTTCCTTGATGCATA

TTTGTGATGAGACAAATGTTTGTTCTATGTTTTAGGTAAGCTGTACATGTTCCAAATCTATAACAAGGACTTCTCCGACAAGTCTCACGGCACTCCTAATCTGCATACAATGTACTTCAAGCTGCTGTTCGACGAGAACAACCACGGTCAGATTAGGCTTTCTGGTGGTGCTGAGCTGTTCATGAGAAGGGCCTCACTGAAGAAAGAAGAGTTGGTCGTTCACCCTGCCAACTCTCCAATCGCTAACAAGAACCCTGACAACCCGAAAAAGACCACCACCTTGTCTTACGACGTGTACAAGGATAAGCGGTTCAGCGAGGATCAGTACGAGCTTCACATTCCGATCGCCATCAACAAGTGCCCGAAGAACATCTTCAAGATCAATACCGAGGTGCGGGTGCTGCTGAAGCACGATGATAATCCTTACGTGATCGGCATCGATAGGGGCGAGAGAAACCTTCTTTACATCGTGGTGGTGGACGGCAAGGGCAATATCGTTGAGCAGTACTCTCTGAACGAGATTATCAACAATTTCAACGGCATCCGGATCAAGACCGACTACCACTCTCTGCTGGATAAGAAAGAAAAAGAGCGGTTCGAGGCCAGGCAGAACTGGACTTCTATCGAAAACATCAAAGAGCTGAAGGCCGGCTACATCTCTCAGGTGGTGCATAAGATTTGCGAGCTGGTGGAAAAGTACGACGCTGTGATTGCTCTCCAGGATCTGAACAGCGGCTTCAAGAACTCACGTGTGAAGGTAAAGCAACTGTGTTTTAATCAATTTCTTGTCAGGATATATGGATTATAACTTAATTTTTGAGAAATCTGTAGTATTTGGCGTGAAATGAGTTTGCTTTTTGGTTTCTCCCGTGTTATAGGTTGAGAAGCAGGTCTACCAAAAGTTCGAGAAGATGCTCATCGACAAGCTGAACTACATGGTGGACAAAAAGAGCAACCCTTGCGCTACCGGTGGTGCTCTTAAGGGTTACCAGATCACTAACAAGTTCGAGTCTTTCAAGAGCATGAGCACCCAGAACGGCTTCATCTTCTACATCCCTGCTTGGCTGACCAGCAAGATCGATCCTTCTACTGGCTTCGTCAACCTGCTCAAGACCAAGTACACCAGCATTGCCGACAGCAAGAAGTTCATCAGCTCATTCGACCGGATCATGTACGTGCCAGAAGAGGATCTTTTCGAGTTCGCCCTCGATTACAAGAACTTCTCTAGGACCGACGCCGACTACATTAAGAAGTGGAAGCTGTACTCCTACGGCAACCGGATTCGGATCTTTCGGAACCCGAAGAAAAACAACGTGTTCGACTGGGAAGAGGTAAAGTTTCCAACTTTCCTTTACCATATCAAACTATAGTTCGAAACTTTTTATTTGATCAACTTCAAGGCCACCCGATCTTTCTATTCCTGATTAATTTGTGATGAATCCATATTGACTTTTGATGGTTACGCAGGTGTGCCTGACCTCTGCCTACAAAGAACTGTTCAACAAGTACGGCATCAACTACCAGCAGGGTGATATTAGGGCTCTGCTTTGCGAGCAGTCTGACAAGGCTTTCTACAGCTCTTTCATGGCCCTGATGTCTCTGATGCTGCAAATGAGGAACTCTATCACCGGTAGGACCGATGTGGACTTCCTTATCTCTCCGGTGAAGAACAGTGACGGGATCTTCTACGACAGCCGGAATTATGAGGCTCAAGAGAACGCAATCCTGCCGAAGAATGCTGATGCTAACGGCGCTTACAACATTGCCAGAAAGGTGCTGTGGGCTATCGGCCAGTTTAAGAAAGCCGAAGATGAGAAGTTGGACAAGGTCTGTCTTTCCTATTTCATATGTTTAATCCTAGGAATTTGATCAATTGATTGTATGTATGTCGATCCCAAGACTTTCTTGTTCACTTATATCTTAACTCTCTCTTTGCTGTTTCTTGCAGGTGAAGATCGCTATCTCCAACAAAGAGTGGCTCGAGTACGCTCAGACTAGCGTTAAGCATAAAAGGCCGGCGGCCACGAAAAAGGCCGGCCAGGCAAAAAAGAAAAAGGGTTCG

**pAGT8173 (Endonuclease: AGGT_NLS-ttLbCas12a-i-NLS_TTCG)**

GG-overhang-NLS-ttLbCas12a-i(*introns*)-NLS-GG-overhang

AGGTCTGCAGCCTAAGAAGAAGAGAAAGGTTGGAGGAGTCGACTCGAGTGCGGCCGCCACAATGAGCAAGCTCGAGAAGTTTACCAACTGCTACAGCCTGTCTAAGACCCTGAGGTTCAAGGCTATTCCTGTGGGTAAGACCCAAGAGAATATCGACAACAAGCGGCTGCTGGTTGAGGATGAGAAGAGAGCTGAGGATTACAAGGGCGTGAAGAAGCTGCTGGATCGGTACTACCTGAGCTTCATCAACGATGTGCTGCACAGCATCAAGCTGAAGAACCTGAACAACTACATCAGCCTGTTCCGGAAGAAAACCCGGACCGAGAAAGAGAACAAAGAGCTTGAGAACCTCGAGATCAACCTGCGGAAAGAGATCGCTAAGGCTTTCAAGGGTAACGAAGGTAAGGATTTTTATGATATACTATGCTTATGTATTTTGTACTGAAAGCATATCCTGCTTCATTGGGATATTACTGAAAGCATTTAACTACATGTAAACTCACTTGATGATCAATAAACTTGATTTTGCAGGTTACAAGAGCCTGTTCAAGAAGGATATTATCGAGACTATCCTGCCTGAGTTCCTGGACGATAAGGATGAGATTGCCCTGGTGAACAGCTTCAACGGTTTCACTACTGCCTTCACCGGTTTCTTCAGAAACCGGGAAAACATGTTCAGCGAAGAGGCCAAGTCTACCTCTATCGCTTTCCGGTGCATTAACGAGAACTTGACCCGGTACATCAGCAACATGGACATCTTCGAGAAGGTGGACGCCATCTTCGATAAGCACGAGGTGCAAGAAATCAAAGAGAAGATCCTGAACTCCGACTACGACGTCGAGGATTTTTTTGAGGGCGAGTTCTTCAACTTCGTGCTCACCCAAGAAGGTAACATTCCTTAGTTACCTTTCTTTTCTTTTTCCATCATAAGTTTATAGATTGTACATGCTTTGAGATTTTCTTTGCAAACAATCTCAGGTATCGATGTGTACAACGCTATCATCGGTGGTTTCGTGACTGAGAGCGGTGAGAAGATTAAGGGCCTGAACGAGTACATTAACCTGTACAATCAAAAGACCAAGCAGAAGCTGCCGAAGTTCAAGCCGCTTTACAAGCAGGTTCTGAGCGATCGTGAGAGCCTGTCTTTTTACGGAGAGGGATACACCTCTGATGAAGAGGTTTTGGAGGTAATCTTGAAATTGGAACTCTTCTTTTGTTGTCTAAACCTATCAATTTCTTTGCGGAAATTTATTTGAAGCTGTAGAGTTAAAATTGAGTCTTTTAAACTTTTGTAGGTGTTCCGTAACACCCTGAACAAGAACAGCGAGATCTTCAGCTCCATCAAGAAGCTGGAAAAGCTGTTTAAGAACTTCGACGAGTACAGCAGCGCTGGCATCTTCGTTAAGAACGGTCCTGCTATCAGCACCATCAGCAAGGATATTTTCGGCGAGTGGAACGTGATCCGGGATAAGTGGAATGCTGAGTACGATGACATCCACCTGAAGAAAAAGGCTGTGGTGACCGAGAAGTACGAGGATGATAGGCGGAAGTCCTTCAAGAAGATAGGTAAGTTGTTACTTATGATTGTTTTCCTCTCTGCTACATGTATTTTGTTGTTCATTTCTGTAAGATATAAGAATTGAGTTTTCCTCTGATGATATTATTAGGTTCCTTTAGCCTCGAGCAGCTTCAAGAGTATGCTGACGCTGATCTGTCCGTGGTCGAGAAGCTTAAAGAGATCATCATCCAGAAGGTCGACGAGATCTACAAGGTGTACGGCAGCTCTGAGAAGCTTTTCGATGCTGACTTCGTGTTGGAGAAGTCTCTGAAGAAGAACGACGCCGTTGTCGCTATCATGAAGGATCTGCTGGACAGCGTGAAGTCTTTCGAGAACTATATCAAGGCCTTCTTCGGCGAAGGTTAGTATCATATGAAGAAATACCTAGTTTCAGTTGATGAATGCTATTTTCTGACCTCAGTTGTTCTCTTTTGAGAATTATTTCTTTTCTAATTTGCCTGATTTTTCTATTAATTCATTAGGTAAAGAGACTAATAGGGACGAGTCATTCTACGGCGATTTCGTGCTGGCTTACGACATCCTTCTTAAGGTGGACCACATCTACGACGCCATCAGAAATTACGTGACCCAGAAGCC

GTACAGCAAGGACAAGTTCAAGTTGTACTTCCAGAATCCGCAGTTCATGGGCGGCTGGGACAAAGACAAAGAGACAG
ATTACAGGGCTACCATCCTGCGGTACGGCTCTAAGTACTACCTTGCCATCATGGACAAGAAATACGCCAAGTGCCTG
CAAAAGATCGACAAGGATGATGTGAACGGCAACTACGAGAAGATCAACTACAAGCTCCTGCCAG*GTAAATCCTGGTC*
*CACACTTTTACGATAAAAACACAAGATTTTAAACTATGAACTGATCAATAATCATTCCTAAAAGACCACACTTTTGT*
*TTTGTTTCTAAAGTAATTTTTACTGTTATAACAG*GTCCTAACAAGATGCTTCCTAAGGTGTTCTTCTCAAAGAAATG
GATGGCCTACTACAACCCGAGCGAGGACATCCAGAAAATCTACAAGAACGGCACCTTCAAAAAGGGCGACATGTTCA
ACCTGAACGACTGCCACAAGCTGATCGATTTCTTCAAGGACAGCATCAGCCGGTATCCGAAGTGGTCTAACGCTTAC
GATTTCAACTTCAGCGAGACTGAGAAGTATAAGGATATCGCCGGCTTCTACCGTGAGGTTGAGGAACAGGGTTACAA
GGTTAGCTTCGAGAGCGCCAGCAAGAAAGAGGTGGACAAGTTGGTTGAAGAAG*GTAAGAGGTCAAAAGGTTTCCGCA*
*ATGATCCCTCTTTTTTTGTTTCTCTAGTTTCAAGAATTTGGGTATATGACTAACTTCTGAGTGTTCCTTGATGCATA*
*TTTGTGATGAGACAAATGTTTGTTCTATGTTTTAG*GTAAGCTGTACATGTTCCAAATCTATAACAAGGACTTCTCCG
ACAAGTCTCACGGCACTCCTAATCTGCATACAATGTACTTCAAGCTGCTGTTCGACGAGAACAACCACGGTCAGATT
AGGCTTTCTGGTGGTGCTGAGCTGTTCATGAGAAGGGCCTCACTGAAGAAAGAAGAGTTGGTCGTTCACCCTGCCAA
CTCTCCAATCGCTAACAAGAACCCTGACAACCCGAAAAAGACCACCACCTTGTCTTACGACGTGTACAAGGATAAGC
GGTTCAGCGAGGATCAGTACGAGCTTCACATTCCGATCGCCATCAACAAGTGCCCGAAGAACATCTTCAAGATCAAT
ACCGAGGTGCGGGTGCTGCTGAAGCACGATGATAATCCTTACGTGATCGGCATCGATAGGGGCGAGAGAAACCTTCT
TTACATCGTGGTGGTGGACGGCAAGGGCAATATCGTTGAGCAGTACTCTCTGAACGAGATTATCAACAATTTCAACG
GCATCCGGATCAAGACCGACTACCACTCTCTGCTGGATAAGAAAGAAAAAGAGCGGTTCGAGGCCAGGCAGAACTGG
ACTTCTATCGAAAACATCAAAGAGCTGAAGGCCGGCTACATCTCTCAGGTGGTGCATAAGATTTGCGAGCTGGTGGA
AAAGTACGACGCTGTGATTGCTCTCGAGGATCTGAACAGCGGCTTCAAGAACTCACGTGTGAAG*GTAAAGCAACTGT*
*GTTTTAATCAATTTCTTGTCAGGATATATGGATTATAACTTAATTTTTGAGAAATCTGTAGTATTTGGCGTGAAATG*
*AGTTTGCTTTTTGGTTTCTCCCGTGTTATAG*GTTGAGAAGCAGGTCTACCAAAAGTTCGAGAAGATGCTCATCGACA
AGCTGAACTACATGGTGGACAAAAAGAGCAACCCTTGCGCTACCGGTGGTGCTCTTAAGGGTTACCAGATCACTAAC
AAGTTCGAGTCTTTCAAGAGCATGAGCACCCAGAACGGCTTCATCTTCTACATCCCTGCTTGGCTGACCAGCAAGAT
CGATCCTTCTACTGGCTTCGTCAACCTGCTCAAGACCAAGTACACCAGCATTGCCGACAGCAAGAAGTTCATCAGCT
CATTCGACCGGATCATGTACGTGCCAGAAGAGGATCTTTTCGAGTTCGCCCTCGATTACAAGAACTTCTCTAGGACC
GACGCCGACTACATTAAGAAGTGGAAGCTGTACTCCTACGGCAACCGGATTCGGATCTTTCGGAACCCGAAGAAAAA
CAACGTGTTCGACTGGGAAGAG*GTAAAGTTTCCAACTTTCCTTTACCATATCAAACTATAGTTCGAAACTTTTTATT*
*TGATCAACTTCAAGGCCACCCGATCTTTCTATTCCTGATTAATTTGTGATGAATCCATATTGACTTTTGATGGTTAC*
*GCAG*GTGTGCCTGACCTCTGCCTACAAAGAACTGTTCAACAAGTACGGCATCAACTACCAGCAGGGTGATATTAGGG
CTCTGCTTTGCGAGCAGTCTGACAAGGCTTTCTACAGCTCTTTCATGGCCCTGATGTCTCTGATGCTGCAAATGAGG
AACTCTATCACCGGTAGGACCGATGTGGACTTCCTTATCTCTCCGGTGAAGAACAGTGACGGGATCTTCTACGACAG
CCGGAATTATGAGGCTCAAGAGAACGCAATCCTGCCGAAGAATGCTGATGCTAACGGCGCTTACAACATTGCCAGAA
AGGTGCTGTGGGCTATCGGCCAGTTTAAGAAAGCCGAAGATGAGAAGTTGGACAAG*GTCTGTCTTTCCTATTTCATA*
*TGTTTAATCCTAGGAATTTGATCAATTGATTGTATGTATGTCGATCCCAAGACTTTCTTGTTCACTTATATCTTAAC*
*TCTCTCTTTGCTGTTTCTTGCAG*GTGAAGATCGCTATCTCCAACAAAGAGTGGCTCGAGTACGCTCAGACTAGCGTT
AAGCATAAAAGGCCGGCGGCCACGAAAAAGGCCGGCCAGGCAAAAAAGAAAAAGGGTTCG

**pAGT7406 (UL12: AATG_UL12_TTCG)**

GG-overhang-UL12-GG-overhang

AATGGAATCTACTGTGGGTCCTGCTTGTCCTCCTGGTAGGACTGTTACTAAGAGGCCTTGGGCTCTTGCTGAGGATA
CTCCTAGAGGTCCTGACTCTCCACCAAAGAGGCCTAGACCTAACTCTCTTCCTCTGACTACTACCTTCAGGCCTTTG
CCACCTCCTCCACAAACTACCTCTGCTGTGGATCCTTCTTCTCACTCTCCTGTGAATCCTCCAAGGGATCAGCATGC
TACTGATACCGCTGATGAGAAGCCTAGAGCTGCTTCTCCTGCTCTGTCTGATGCTTCTGGTCCTCCTACTCCTGATA
TCCCTCTTTCTCCTGGTGGTACTCATGCTAGGGATCCTGATGCAGATCCTGATAGCCCTGATCTGGACTCTATGTGG
TCTGCTTCTGTGATCCCTAACGCTCTGCCTTCTCACATTCTGGCTGAGACTTTCGAGAGGCACCTTAGGGGTTTGCT
TAGAGGTGTTAGGGCTCCTCTTGCTATTGGTCCTCTTTGGGCTAGACTGGACTACCTTTGCTCTCTTGCTGTGGTGC
TTGAAGAGGCTGGTATGGTGGATAGAGGTCTTGGTAGACACCTTTGGAGGCTTACTAGAAGAGGTCCTCCAGCTGCT
GCTGATGCTGTTGCTCCTAGACCTCTTATGGGATTCTACGAGGCTGCTACTCAGAACCAGGCTGATTGTCAACTTTG
GGCTCTGCTTAGAAGGGGTCTTACTACCGCTTCTACTCTTAGATGGGGTCCTCAGGGACCTTGCTTTTCTCCTCAAT
GGCTGAAACACAACGCTAGCCTTAGGCCTGATGTGCAGTCATCTGCTGTGATGTTCGGTAGGGTTAACGAGCCTACC
GCTCGGTCTTTGCTTTTCAGGTATTGCGTTGGCAGGGCTGATGATGGTGGTGAAGCTGGTGCTGATACCAGGCGGTT
TATTTTCCACGAGCCATCTGATCTGGCCGAAGAGAATGTTCATACCTGCGGTGTGCTTATGGATGGTCACACTGGAA
TGGTGGGCGCTTCTCTTGATATTCTTGTGTGCCCTAGGGACATCCACGGTTACCTTGCTCCAGTTCCTAAGACTCCT
CTGGCCTTTTACGAGGTTAAGTGCAGGGCTAAGTACGCTTTCGATCCTATGGACCCTTCTGACCCTACTGCTTCTGC
TTACGAGGATCTGATGGCTCATAGAAGCCCTGAGGCTTTCAGGGCTTTCATCCGGTCTATTCCTAAGCCGAGCGTGA
GATACTTTGCTCCTGGAAGAGTTCCTGGTCCTGAGGAAGCTCTTGTTACTCAAGATCAGGCTTGGTCTGAGGCTCAT
GCTTCAGGTGAGAAGAGAAGATGCTCAGCTGCTGATAGGGCACTCGTTGAGCTTAATTCTGGCGTGGTGTCTGAGGT

GTTGCTTTTCGGTGCTCCTGATCTCGGTAGGCACACTATTTCTCCAGTGAGCTGGTCCTCTGGTGATCTTGTTAGAA
GGGAACCCGTGTTCGCTAATCCTAGGCACCCTAACTTCAAGCAGATTCTGGTGCAGGGTTACGTGCTGGATTCTCAC
TTTCCAGATTGCCCTCCACATCCTCACCTTGTGACTTTCATTGGTCGGCATAGGACCTCAGCTGAAGAGGGTGTTAC
TTTCAGGCTTGAGGATGGTGCTGGTGCTCTTGGTGCTGCTGGTCCTTCTAAGGCTTCTATTCTTCCTAACCAGGCCG
TGCCTATCGCTCTTATTATCACCCCTGTGAGGATCGACCCCGAGATCTATAAGGCTATCCAGAGGTCATCTCGGCTG
GCTTTCGATGATACTTTGGCTGAGCTTTGGGCCTCTAGATCTCCTGGTCCAGGTCCTGCTGCTGCAGAAACTACTTC
TTCTTCACCTACCACCGGCAGGTCATCTAGAGGTTCG

**pAGT8081 (nucleoplasm NLS: AATG_nucNLS_TTCG)**
GG-overhang-nucNLS-GG-overhang
AATGGGTAAAAGGCCGGCGGCCACGAAAAAGGCCGGCCAGGCAAAAAAGAAAAAGGGTTCG

**pAGT8080 (nucleoplasm NLS: AGGT_nucNLS_TTCG)**
GG-overhang-nucNLS-GG-overhang
AGGTAAAAGGCCGGCGGCCACGAAAAAGGCCGGCCAGGCAAAAAAGAAAAAGGGTTCG

**pAGT7355 (SSAP: AATG_ICP8i_TTCG)**
GG-overhang-ICP8-i(*introns*)-GG-overhang
AATGGAAACTAAGCCTAAGACCGCTACTACCATCAAGGTTCCACCTGGTCCTCTTGGTTACGTTTACGCTAGAGCTT
GTCCGTCTGAGGGTATTGAGCTTCTTGCTCTGCTTAGCGCTAGGTCCGGTGATTCTGATGTTGCTGTTGCTCCTCTT
GTGGTGGGTCTTACTGTTGAGTCTGGCTTCGAGGCTAATGTGGCTGTTGTTGTTGGTTCTAGGACCACCGGTCTTGG
TGGTACTGCTGTGTCTCTTAAGCTTACCCCGAGCCACTACAGCAGCTCTGTTTACGTTTTCCATGGTGGCAGGCACC
TGGATCCTTCTACTCAAGCTCCTAACCTTACCAGGCTTTGTGAGAGGGCTAGAAGGCACTTCGGCTTCTCTGATTAT
ACTCCTAGGCCTGGCGATCTGAAGCACGAAACTACTGGTGAAGCTCTTTGCGAGAGGCTTGGTCTTGATCCTGATAG
GGCTCTGCTTTACCTTGTGGTGACCGAGGGTTTCAAAGAGGCTGTCTGCATCAACAACACCTTTCTGCACCTTGGCG
GCTCTGATAAGGTAACATTCCTTAGTTACCTTTCTTTTCTTTTTCCATCATAAGTTTATAGATTGTACATGCTTTGA
GATTTTTCTTTGCAAACAATCTCAGGTTACCATTGGTGGTGCTGAGGTGCACAGGATTCCAGTTTATCCTCTGCAGC
TGTTCATGCCGGATTTCTCTAGAGTGATCGCTGAGCCTTTCAACGCCAACCACAGATCTATCGGCGAGAACTTCACT
TACCCTCTGCCGTTCTTCAACAGGCCTCTTAACAGGCTTTTGTTCGAGGCTGTTGTGGGTCCTGCTGCTGTTGCACT
TAGGTGTAGGAATGTTGATGCTGTGGCTAGGGCTGCTGCTCATCTTGCTTTCGATGAGAATCATGAGGGTGCTGCTC
TGCCTGCTGATATTACTTTCACTGCTTTCGAGGCCAGCCAGGGTAAGACTCCTAGAGGTGGTAGAGATGGTGGTGGT
AAGGGACCTGCTGGTGGTTTTGAACAGAGGCTTGCTTCTGTGATGGCTGGTGATGCTGCTCTTGCTCTTGAGAGCAT
TGTGAGCATGGCTGTGTTTGATGAGCCTCCGACCGATATTTCTGCTTGGCCTTTGTTCGAAGGCCAGGATACTGCTG
CTGCTAGGGCTAATGCTGTTGGTGCTTATCTTGCTCGTGCTGCTGGTCTTGTTGGAGCCATGGTGTTCTCTACTAAC
TCTGCTCTCCACCTTACCGAGGTTGACGATGCTGGTCCAGCTGATCCAAAGGATCACAGCAAGCCAAGCTTCTACCG
GTTCTTCCTTGTGCCTGGTACTCATGTGGCTGCTAATCCTCAGGTTGACCGTGAAGGTCATGTGGTGCCTGGTTTTG
AGGGTAGACCTACTGCTCCTTTGGTTGGTGGAACTCAAGAGTTCGCTGGTGAGCACCTTGCTATGCTTTGCGGTTTT
TCTCCTGCTCTGCTGGCCAAGATGCTGTTCTATCTTGAGAGGTAAGTTGTTACTTATGATTGTTTTCCTCTCTGCTA
CATGTATTTTGTTGTTCATTTCTGTAAGATATAAGAATTGAGTTTTCCTCTGATGATATTATTAGGTGTGATGGCGG
AGTGATCGTTGGTAGACAAGAGATGGACGTGTTCCGGTACGTGGCAGATTCTAATCAGACTGACGTGCCATGCAACT
TGTGCACCTTCGATACAAGGCATGCTTGCGTGCACACTACCCTTATGAGACTTAGAGCTAGGCACCCGAAGTTCGCT
TCAGCTGCTAGAGGTGCTATTGGTGTGTTCGGCACCATGAACTCCATGTACAGCGACTGTGATGTCCTGGGAAACTA
CGCTGCTTTCAGCGCTCTTAAGCGTGCTGATGGTTCTGAGACTGCTCGGACCATTATGCAAGAGACTTACAGGGCTG
CTACCGAGAGAGTTATGGCTGAGCTTGAGACTCTTCAGTACGTTGACCAGGCTGTGCCTACTGCTATGGGTAGACTT
GAGACTATCATCACCAACCGTGAGGCTCTGCATACCGTGGTTAACAATGTTAGGCAGGTTGTGGACCGTGAGGTTGA
GCAGCTTATGAGGAATCTTGTTGAGGGCCGTAACTTCAAGTTCAGAGATGGTCTTGGAGAGGCCAACCACGCTATGT
CTCTTACCCTTGATCCTTACGCTTGCGGTCCTTGTCCTCTGCTTCAACTTCTTGGTCGGAGGTCTAACTTGGCTGTG
TACCAGGATTTGGCTCTGTCTCAGTGCCACGGTGTTTTTGCTGGTCAATCTGTGGAAGGTCGGAACTTCAGAAATCA
GTTCCAGCCTGTGCTTCGGAGGCGTGTGATGGATATGTTCAACAACGGTTTCCTGAGCGCTAAGACCCTTACCGTTG
CTCTTTCTGAGGGTGCAGCTATTTGCGCTCCTTCTCTTACTGCTGGTCAAACCGCTCCTGCTGAGTCATCATTTGAG
GGTGATGTGGCTAGAGTGACCCTGGGTTTCCTAAAGAGCTGCGGGTTAAGTCTAGGGTGCTGTTTGCTGGTGCTTC
CGCTAATGCTTCTGAAGCTGCTAAGGCTAGGGTTGCCTCTCTTCAGTCTGCTTACCAGAAGCCGGATAAGAGGGTTG
ACATTCTTCTTGGCCCTCTGGGTTTCCTGCTGAAGCAATTCCATGCTGCTATCTTCCCTAACGGTAAGCCTCCTGGA
AGCAACCAGCCTAATCCACAGTGGTTTTGGACTGCCCTTCAGAGGAATCAGCTTCCTGCTAGGCTTCTGAGCAGAGA
GGATATCGAGACTATCGCCTTTATCAAGAAATTCAGCCTGGACTACGGCGCCATCAACTTCATTAACCTGGCTCCGA
ACAACGTGAGCGAGCTGGCTATGTACTACATGGCTAACCAGATCCTGAGGTTAGTATCATATGAAGAAATACCTAGT
TTCAGTTGATGAATGCTATTTTCTGACCTCAGTTGTTCTCTTTTGAGAATTATTTCTTTTCTAATTTGCCTGATTTT

TCTATTAATTCATTAGGTACTGCGACCACTCTACCTACTTCATCAACACCCTGACCGCTATTATCGCTGGCTCTAGA
AGGCCTCCATCTGTTCAAGCTGCTGCCGCTTGGTCTGCTCAAGGTGGTGCAGGTCTTGAAGCTGGTGCTAGAGCACT
TATGGATGCTGTTGACGCTCATCCTGGTGCTTGGACTTCTATGTTCGCTAGCTGCAACCTTCTCAGGCCTGTTATGG
CAGCTAGGCCTATGGTTGTGCTGGGCCTTTCTATCAGCAAGTACTACGGTATGGCTGGCAACGATAGGGTTTTCCAG
GCTGGTAATTGGGCTAGCCTTATGGGTGGTAAGAACGCTTGCCCTCTGCTGATTTTCGACCGGACTAGGAAGTTCGT
TCTGGCTTGTCCTAGAGCTGGTTTCGTTTGCGCTGCTTCATCTCTTGGAGGTGGTGCTCATGAGTCTAGTTTGTGCG
AACAGCTGAGGGGCATTATCTCTGAAGGTAAATCCTGGTCCACACTTTTACGATAAAAACACAAGATTTTAAACTAT
GAACTGATCAATAATCATTCCTAAAAGACCACACTTTTGTTTTGTTTCTAAAGTAATTTTTACTGTTATAACAGGTG
GCGCTGCAGTTGCTAGCTCTGTGTTTGTTGCTACCGTGAAGTCACTTGGCCCTAGAACTCAGCAGCTCCAGATTGAG
GATTGGCTGGCTCTTTTGGAGGACGAGTACCTGTCTGAAGAGATGATGGAACTTACCGCTAGGGCTCTTGAGCGTGG
TAATGGTGAATGGTCTACCGATGCTGCTTTGGAGGTTGCACATGAAGCTGAGGCTCTTGTGTCTCAGCTTGGTAATG
CTGGTGAGGTGTTCAACTTCGGTGATTTCGGTTGCGAGGATGACAACGCTACTCCTTTTGGTGGACCTGGTGCTCCT
GGTCCTGCTTTTGCTGGAAGAAAAAGGGCTTTCCACGGCGACGATCCTTTTGGAGAAGGTCCTCCTGATAAGAAGGG
TGATCTGACCCTTGACATGCTTTCG

**pAGT7354 (EcSSB: AATG_SSB_TTCG)**

GG-overhang-SSB-GG-overhang

AATGGCTTCTAGGGGCGTTAACAAGGTGATCCTTGTGGGTAACCTTGGTCAGGATCCCGAAGTTCGGTACATGCCTA
ATGGTGGTGCTGTGGCCAATATTACCCTTGCCACTTCTGAGAGCTGGCGTGATAAGGCTACTGGCGAAATGAAGGAA
CAGACCGAGTGGCATAGGGTTGTGCTTTTCGGTAAGCTTGCTGAGGTGGCATCTGAGTACCTTAGGAAGGGTAGCCA
GGTTTACATTGAGGGTCAGCTGAGAACTAGGAAGTGGACCGATCAGTCTGGACAGGATAGGTACACTACTGAGGTGG
TGGTGAACGTTGGTGGAACCATGCAAATGCTTGGTGGTAGACAAGGTGGTGGCGCTCCTGCTGGTGGAAATATTGGT
GGTGGTCAACCTCAAGGCGGTTGGGGTCAGCCTCAACAACCACAAGGTGGAAATCAGTTCTCAGGCGGTGCTCAATC
TAGACCTCAACAATCTGCTCCAGCTGCTCCTTCTAATGAGCCTCCTATGGATTTCGACGACGACATCCCTTTCGGTT
CG

**pAGT6575 (AtRad51i genomic sequence: AATG_Rad51i_GCTT)**

GG-overhang-AtRad51(*intron*)-GG-overhang

AATGACGACGATGGAGCAGCGTAGAAACCAGAATGCTGTCCAACAACAAGACGATGAAGAAACCCAGCACGGACCTT
TCCCTGTCGAACAG*CTTCAGGTTCAATTTCTGCAGATTTCTTTCATTTGTTGTGGAAATTTACCTTCGCAGTCTATA*
*CGCTTCATCAGATCTTGCGCTAATTGAAATGGGTTCGGCTTTAATTCACTAAAAATTTCTGCTTTTTTTGTACATCG*
*CGATGTGAAATAGGTGTGAATCTGGGCGAGAATTGAGTTTCGCTACTGTCTGTTAGGCGAATTGGTTTTAGGGGTTC*
*GTTTAATTCAGAAAAAAATCGACACTTTTTGGGGGTTTTATCCTTTTCTGGGGAACTTCTCTATTGCTGTGACTCCA*
*GTGTCATATAATCTCTATAGTTCTTCTCAATTCGTGTTCATAAAAGGGAGATGCATCAGTGTGTTTTGTGTTATGAT*
*TATATGCTATTTCATCTCTTTAAATCTTCAAAA*CTTCAGGCAGCAGGTATTGCTTCTGTTGATGTAAAGAAGCTTAG
GGATGCTGGACTCTGTACTGTTGAAGGTGTTGCTTATACTCCGAGGAAGGATCTCTTGCAGATTAAAGGAATTAGTG
ATGCCAAGGTTGACAAGATTGTAGAAGCA*GGTATTACACATGTAGAAACTTTTTGCTTTCCTTCTCTGTTAAATACA*
*TAACACCTCTTTGATACTCTTAGAGTTAATGTGTGTTCTTATTTGTGTTTTCTTCTGTGATA*GCTTCAAAGCTAGTT
CCTCTGGGGTTCACTAGTGCGAGCCAGCTCCATGCTCAGAGACAGGAAATTATTCAGATTACCTCTGGATCACGGGA
GCTCGATAAAGTTCTAGA*AGGTTGGTGCTGGTTTCTGGTGCATCTTATAACGGCATTTTGTTATTGTGATCTTACTT*
*TGCCGTATGCTCAAC*AGGAGGTATTGAAACTGGTTCCATCACAGAGTTATATGGTGAGTTCCGCTCTGGAAAGACTC
AGCTGTGCCATACACTGTGTGTGACTTGTCAA*GTAAGAACCCTGTTCCTATACCACTCAGTTATTGTTGCTTATCTA*
*GCAGAAATCAGTGATCTTGTCTTTCTTCTTACAATTTCCAAACCTGAGAAAGATATTCACATTAAGGTTCACGTTTG*
*AATGATTAG*CTTCCCATGGATCAAGGAGGTGGAGAGGGAAAGGCCATGTACATTGATGCTGAGGGAACATTCAGGCC
ACAAAGACTCTTACAGATAGCTGA*CAGGTATGTTATTCTTGTAACACACATGAGATCACAATAGCGTTCTTGAGACG*
*GAATCATCTGTAAGAAATCAAATGAAATGATG*CAGGTTTGGATTAAATGGAGCTGATGTACTAGAAAACGTTGCCTA
TGCGAGGGCGTATAATACAGATCATCAGTCAAGGCTTTTGCTTGAAGCAGCATCAATGATGATTGAAACAAG*GTGTG*
*TTGAGTTTATTTTGGTGTTCAGTTCCGTTATGTCTATCATGAGAGTGCTAACTTTTAAAGTTACGATGACCCTATTG*
*GCACAG*GTTTGCTCTCCTGATTGTCGATAGTGCTACCGCTCTCTACAGAACAGATTTCTCTGGAAGGGGAGAGCTTT
CGGCTCGACAAATGCATCTTGCAAAGTTCTTGAGATCTCTTCAGAAGTTAGCAGATGAG*GTGAACATTCAGTACTAA*
*CTGTTTCTTTTCCAATATTGCTCTGAGACACCATCCTGAAGTATTCTCTCTTATGTTTAATGGCTCTAG*TTTGGTGT
GGCTGTTGTTATAACAAACCAAGTAGTTGCGCAAGTAGATGGTTCAGCTCTTTTTGCTGGTCCCCAATTTAAGCCGA
TTGGTGGGAATATCATGGCTCATGCCACCACAACAAG*GTCTGTAATGTTTGCAAATTCGACAAATCCATGTTTCTTA*
*GTGTTTTTTGTTAGGGTTCGTTAGGAAAGTGTGTTTGTTATGATAATTGATACAG*GTTGGCGTTGAGGAAAGGAAGA
GCAGAGGAGAGAATCTGTAAAGTGATAAGCTCGCCATGTTTGCCAGAAGCGGAAGCTCGATTTCAAATATCTACAGA
AGGTGTAACAGATTGCAAGGATTGAGCTT

**pAGT7883 (AtRad52-1i genomic sequence: AATG_Rad52-1i_GCTT)**

GG-overhang-AtRad52-1(*intron*)-GG-overhang

AATGGCGGGGTTAGGGTTACGGCTAAAGGCGGCAAAATGGACGTTGAGGAGCGGATCAGGTGCTGTATCGAGAGAAT
GGTCGTCGGAGATGGGGAAAGGTGTTCGTCGATTCTCTACTGAAACAGAGAACGATGTTCCAACTTCAGGAATCAGT
AGACCACTCGCTGAGATTCTCAAGGAGCTTAACAAAAAGGTTCCAGATTCAGTCATCCGAACTCGAGTTGAAGATGG
GTGCTCCATCAAATACATCCCTTG*GTAATCTCTTTCCTCAGGGTTTAGGGTTCTTTAATTGGGGTTTTTGGGTCAAT*
*ACTTATACTCTTTTCTTGTGTTGATTCTCAG*GCACATTGTGAATCGGATCATGAATATGCATGCTCCA*GGTTTGCTA*
*AACCCTCTCTTGTTCTTCTGGGGTTCGTTGTTGGGTTTTGGTTTTGACATTTTGGTGTATTATTCTCTGTTTTCA*GA
ATGGTCAGGAGAAGTTCGAAGTGTCACTTACTCTCCTGATGGCAACACTGTTACTGTGGCTTATCGTGTGACTCTCT
ACGGCACTGATGCTG*AGGTCTTTCCCTTCCTCTTGTTTCAGTGTTATGGTTTGATTTGGTTAGCCTTTAGCTCAAGT*
*CGTCTCCTGGTAGAGTTCTTTGCCTTCTTGCCTCAGAAGCTGCAAATATAGAAACCTAGTTGTTTTCTTTTGGGTTG*
*TGTCCATTTTGAATTGTATGATGAAACCCGTCAAGGAATACAGCTTAGAGAGTGTATGTTTATCAACTCTGGAGAAA*
*CACCAACTCTGGGTTATAGATTCTGTTATCAGTACTTACAAACCTTAGAGATGATATCACTTGTGAATAGGGAGGTT*
*TGACCTGTTTTCGTTAGTAAATATGCTAGTGGTTTGCTTCCTTGCATACTTTCTTGAGCATATTATGTTTTCATTGT*
*TTTGCTTGTAACTTGTACGCTGAAATCATTTGCTAATAATCATTTTTTTGGTGTTGTCA*AGATATTTAGGGAATCAA
CAGGTACCACATCTGTAGATGACAAAGGCTATGGTGATGCTGTGCAAAAGGCGGAAGCAATGGCATTTCGCCGTGCG
TGTGCAAGATTTGGATTAGGGCTTCATCTTTATCACGAGGATGCTTTGGGTTCG

**pAGT7884 (AtRad52-2i genomic sequence: AATG_Rad52-2i_GCTT)**

GG-overhang-AtRad52-2(*intron*)-GG-overhang

AATGGCTTTGCAAGTGCAGCAAACTTCAGCAGCCTTCACAATCTCCTCACCATCCACCGCCGCTGCACGCATCAAAC
TCTCCCCTTTCAGAACCGTGGCGGTTAACCGCGGAGTTAGATGCAGCGGCGGTGGAGTCGGAGGCGGAGATGCCGGG
AAGAAAAAAGCAGTTCCTAATTCGAATTATGTAGTTCCTATTGACAAATTCTCATCTTCTTCGTCGATTACTAGGCC
TTTGATTGAGATACTTCGTGACCTTAATAAGAAGATTCCTGATAACATTGTTAAGAGTCATGATCCTCCTTCTACTT
CTGCTGCTACTTCTGGTTTTATCCCTTG*GTATTAATTATCTTTTTTTTTTATTATCTTTTTGTTGTTAAGAAGATT*
*CTTGTTGGTTCTTGATCTGAGGTTTTAATTTATAG*GTACCATGCAAATCGGATGCTTAGCTTCTATGCTCCTGGTTG
GTGTGGTGAAGTTAGAGATGTTATTTTCTCTGAGAATGGAAATGTTACTGTTGTTTATCGTCTCACTATTCGTGGCT
CTGATGGTGAG*GTAAAAGTTCTTCAACATTCCTCAATTTCTTTCTTTCTTTCTTTCCTTGGGTGTAGTTGTGTGTGA*
*TTGAGTAGGTAAGCAAAGAAACTTATGTCTGATTTGACTAATTTTGTGAAAAATTAAACCTTTGCATTGCATACATT*
*GTGTCTCTCTTATTGTATCTTTGTAGAGGAAAAAACTTTGATGGTAGTTTTCAAGAGTTTATTAATGATTAAGGAAA*
*AGATTCTAGTTCATGTATTGATGCTTGTTTGTCTTGTTTTTGTGCAAAATCAAACCTTTCTAGAGGTTCATTTTGCT*
*AATGGTATATTTCACGAGTATAACAACGATTATAAGCATATGCTTTGGTAGATTGAAACCAAAGGTAATGTGATCGT*
*GGAAGTCCTCGAGCCTTATGACTTTGGTCATCTATGGCAATGCCTAAACCAGCAAAATCCCATTTTTATGACTTTGA*
*TATGATAGAAGGTTGTTTTGTGGAGAGCGAGTTGGATTGGTTGTTGAAGTTGTAGATTAGTGAACCAACATATGTAC*
*CATTGATGACAGGACTAAGTTTTGAAGTTTTATGTAATGCAG*GCACATCGTGAATCAACTGGGACAGTAACGACGAC
TGATGATCACATCGAGGATCCAGTTACTGCAGCTGAGGAAATAGCATTCTGCAGAGCATGTGCTCGATTTGGACTCG
GCTTGTATCTATACCACGAAGGTTCG

**pAGT8089 (XTEN16-nucleoplasm NLS: TTCG_XTEN16-nNLS_GCTT)**

GG-overhang-XTEN16-nNLS-GG-overhang

TTCGGGTTCAGAAACCCCTGGTACTAGCGAGTCAGCTACACCAGAGTCAGGTAAAAGGCCGGCGGCCACGAAAAAGG
CCGGCCAGGCAAAAAAGAAAAAGGGTTGAGCTT

**pAGT5439 (tOCS: TTCG_STOP-tOCS_CGCT)**

GG-overhang-STOP-tOCS-GG-overhang

TTCGTGAGCTTGTCCTGCTTTAATGAGATATGCGAGAAGCCTATGATCGCATGATATTTGCTTTCAATTCTGTTGTG
CACGTTGTAAAAAACCTGAGCATGTGTAGCTCAGATCCTTACCGCCGGTTTCGGTTCATTCTAATGAATATATCACC
CGTTACTATCGTATTTTTATGAATAATATTCTCCGTTCAATTTACTGATTGTACCCTACTACTTATATGTACAATAT
TAAAATGAAAACAATATATTGTGCTGAATAGGTTTATAGCGACATCTATGATAGAGCGCCACAATAACAAACAATTG
CGTTTTATTATTACAAATCCAATTTTAAAAAAAGCGGCAGAACCGGTCAAACCTAAAAGACTGATTACATAAATCTT
ATTCAAATTTCAAAAGTGCCCCAGGGGCTAGTATCTACGACACACCGAGCGGCGAACTAATAACGCTCACTGAAGGG
AACTCCGGTTCCCCGCCGGCGCGCATGGGTGAGATTCCTTGAAGTTGAGTATTGGCCGTCCGCTCTACCGAAAGTTA
CGGGCACCATTCAACCCGGTCCAGCACGGCGGCCGGGTAACCGACTTGCTGCCCCGAGAATTATGCAGCATTTTTTT
GGTGTATGTGGGCCCCAAATGAAGTGCAGGTCAAACCTTGACAGTGACGACAAATCGTTGGGCGGGTCCAGGGCGAA
TTTTGCGACAACATGTCGAGGCTCAGCAGGACCGCT

**pICH41432 (tOCS: GCTT_tOCS_CGCT)**

GG-overhang-tOCS-GG-overhang
GCTTGTCCTGCTTTAATGAGATATGCGAGAAGCCTATGATCGCATGATATTTGCTTTCAATTCTGTTGTGCACGTTG
TAAAAAACCTGAGCATGTGTAGCTCAGATCCTTACCGCCGGTTTCGGTTCATTCTAATGAATATATCACCCGTTACT
ATCGTATTTTTATGAATAATATTCTCCGTTCAATTTACTGATTGTACCCTACTACTTATATGTACAATATTAAAATG
AAAACAATATATTGTGCTGAATAGGTTTATAGCGACATCTATGATAGAGCGCCACAATAACAAACAATTGCGTTTTA
TTATTACAAATCCAATTTTAAAAAAAGCGGCAGAACCGGTCAAACCTAAAAGACTGATTACATAAATCTTATTCAAA
TTTCAAAAGTGCCCCAGGGGCTAGTATCTACGACACACCGAGCGGCGAACTAATAACGCTCACTGAAGGGAACTCCG
GTTCCCCGCCGGCGCGCATGGGTGAGATTCCTTGAAGTTGAGTATTGGCCGTCCGCTCTACCGAAAGTTACGGGCAC
CATTCAACCCGGTCCAGCACGGCGGCCGGGTAACCGACTTGCTGCCCCGAGAATTATGCAGCATTTTTTTGGTGTAT
GTGGGCCCCAAATGAAGTGCAGGTCAAACCTTGACAGTGACGACAAATCGTTGGGCGGGTCCAGGGCGAATTTTGCG
ACAACATGTCGAGGCTCAGCAGGACCGCT

**pAGT8149 (tOCS: TTCG_STOP-tNOS_CGCT)**
GG-overhang-STOP-tNOS-GG-overhang
TTCGTGATCCTCTAGAGTCAAGCAGATCGTTCAAACATTTGGCAATAAAGTTTCTTAAGATTGAATCCTGTTGCCGG
TCTTGCGATGATTATCATATAATTTCTGTTGAATTACGTTAAGCATGTAATAATTAACATGTAATGCATGACGTTAT
TTATGAGATGGGTTTTTATGATTAGAGTCCCGCAATTATACATTTAATACGCGATAGAAAACAAAATATAGCGCGCA
AACTAGGATAAATTATCGCGCGCGGTGTCATCTATGTTACTAGATCGACGCT

**pAGT8149 (tOCS: GCTT_tNOS_CGCT)**
GG-overhang-tNOS-GG-overhang
GCTTGATCCTCTAGAGTCAAGCAGATCGTTCAAACATTTGGCAATAAAGTTTCTTAAGATTGAATCCTGTTGCCGGT
CTTGCGATGATTATCATATAATTTCTGTTGAATTACGTTAAGCATGTAATAATTAACATGTAATGCATGACGTTATT
TATGAGATGGGTTTTTATGATTAGAGTCCCGCAATTATACATTTAATACGCGATAGAAAACAAAATATAGCGCGCAA
ACTAGGATAAATTATCGCGCGCGGTGTCATCTATGTTACTAGATCGACGCT

**pAGT5824 (Solanum leucopersicum pU6: GGAG_SlpU6_ATTG)**
GG-overhang-SlpU6-GG-overhang
GGAGCCAAAACGTAGAACCGATACCCCAATAAAACAAAAGTACCAATACCGTTATTAAAATAGTTAAACCAATAACT
CATTACCGATAAATCGCAGTTCTACTTTGTTTTTCTCCATTGAAGTCCATGAAGCAAGAGAAATAAGAAAAAGGCCG
GCCCATTTAGATTGAAGATCCAGGCCCAGGCCCGTAAAAGAAACCCAACAAGCAAATTCTCCCTCATCGCTTATACA
AAGCTACTTTGCCTCGTTTATATAGCGGAATATGAACATGTATGATTG

**pICSL90001 (synthetic pU6 from three At U6 promoters: GGAG_sAtpU6_ATTG)**
GG-overhang-sAtpU6-GG-overhang
GGAGTGATCAAAAGTCCCACATCGATCAGGTGATATATAGCAGCTTAGTTTATATAATGATAGAGTCGACATAGCGA
TTG

**pAGT6271 (At tU6-26 67nt (t67): GGAG_t67_ATTG)**
GG-overhang-t67-GG-overhang
TTTTTTTTGCAAAATTTTCCAGATCGATTTCTTCTTCCTCTGTTCTTCGGCGTTCAATTTCTGGGGTCGCT

**General SpCas9 sgRNA module (sgRNA: ATTG_Spacer-sgRNA(flip+extension)-t67_CGCT)**
GG-overhang-Spacer(20nt)-sgRNA-t67-GG-overhang
ATTGNNNNNNNNNNNNNNNNNNNNGTTTAAGAGCTATGCTGGAAACAGCATAGCAAGTTTAAATAAGGCTAGTCCGT
TATCAACTTGAAAAAGTGGCACCGAGTCGGTGCTTTTTTTTGCAAAATTTTCCAGATCGATTTCTTCTTCCTCTGTT
CTTCGGCGTTCAATTTCTGGGGTCGCT

**General ttLbCas12a-I crRNA module (crRNA: ATTG_DR-spacer_TTTT)**
GG-overhang-DR-Spacer(20 nt)-GG-overhang
ATTGTAATTTCTACTAAGTGTAGATNNNNNNNNNNNNNNNNNNNNTTTT

**Table 2. SpCas9 targets used in this study**

| sgRNA | Target sequence | Figure |
|---|---|---|
| TMV-In1 | TGTTGATCAAAAGATGGGTG AGG | Fig.1, Fig.4 |
| TMV-In2 | AAGCCGCGGTGCGGGTGCCA GGG | Fig.1, Fig.4 |

| | | |
|---|---|---|
| TMV-Out1 | CACCGCGGCTTCGAGAAAAG AGG | Fig.1, Fig.4 |
| TMV-Out2 | GGCGTGCCCTTGGGCTCCCC GGG | Fig.1, Fig.4 |
| NbPGK-1 | AGCCACATATCCACTGGTGG TGG | Fig.2 |
| NbPGK-1 | CCACTGATTATGCTGATCAC CGG | Fig.2 |
| NbTPR-1 | CGTAGCTACAGCGTCTATAA CGG | Fig.2 |
| NbTPR-2 | TAGTCAGTCGGCGACGACAG CGG | Fig.2 |
| D1 | CAAGCTCTTGTTGCTCAAGG AGG | Fig.5 |
| sgR-ThiC1 | ACAACTACTTATATTCGCCA TGG | Fig.5 |

**Table 3. ttLbCas12a-i targets used in this study**

| crRNA | Target sequence | Figure |
|---|---|---|
| TMV-In1 | CTTC GAGAAAAGAGGTCAGAAAAT | Fig.4 |
| TMV-In2 | TTTG ATCAACATCAAAATTAGGTT | Fig.4 |
| TMV-Out1 | TTTC TCGAAGCCGCGGTGCGGGTG | Fig.4 |
| TMV-Out2 | TTTG ATGTTGATCAAAAGATGGGT | Fig.4 |

**TMV-locus (nbi775)**
**TMV transgene (pICH38191 in *Nicotiana benthamiana*)**
RB_tNOS-Kan-pNOS_pAct2-Ω-RdRP(*intron*)-AttB-MP-GFP-3´NTR-tNOS_LB

CCTGTGGTTGGCACATACAAATGGACGAACGGATAAACCTTTTCACGCCCTTTTAAATATCCGATTATTCTAATAAA
CGCTCTTTTCTCTTAGGTTTACCCGCCAATATATCCTGTCAAACACTGATAGTTTAAACTGAAGGCGGGAAACGACA
ATCTGATCTAAGCTTGCATGCCTGCAGGTCGATCTAGTAACATAGATGACACCGCGCGCGATAATTTATCCTAGTTT
GCGCGCTATATTTTGTTTTCTATCGCGTATTAAATGTATAATTGCGGGACTCTAATCATAAAAACCCATCTCATAAA
TAACGTCATGCATTACATGTTAATTATTACATGCTTAACGTAATTCAACAGAAATTATATGATAATCATCGCAAGAC
CGGCAACAGGATTCAATCTTAAGAAACTTTATTGCCAAATGTTTGAACGATCTGCTTGACTCTAGATCCAGAGTCCC
GCTCAGAAGAACTCGTCAAGAAGGCGATAGAAGGCGATGCGCTGCGAATCGGGAGCGGCGATACCGTAAAGCACGAG
GAAGCGGTCAGCCCATTCGCCGCCAAGCTCTTCAGCAATATCACGGGTAGCCAACGCTATGTCCTGATAGCGGTCCG
CCACACCCAGCCGGCCACAGTCGATGAATCCAGAAAAGCGGCCATTTTCCACCATGATATTCGGCAAGCAGGCATCG
CCATGAGTCACGACGAGATCCTCGCCGTCGGGCATACGCGCCTTGAGCCTGGCGAACAGTTCGGCTGGCGCGAGCCC
CTGATGCTCTTCGTCCAGATCATCCTGATCGACAAGACCGGCTTCCATCCGAGTACGTGCTCGCTCGATGCGATGTT
TCGCTTGGTGGTCGAATGGGCAGGTAGCCGGATCAAGCGTATGCAGCCGCCGCATTGCATCAGCCATGATGGATACT
TTCTCGGCAGGAGCAAGGTGAGATGACAGGAGATCCTGCCCCGGCACTTCGCCCAATAGCAGCCAGTCCCTTCCCGC
TTCAGTGACAACGTCGAGCACAGCTGCGCAAGGAACGCCCGTCGTGGCCAGCCACGATAGCCGCGCTGCCTCGTCCT
GGAGTTCATTCAGGGCACCGGACAGGTCGGTCTTGACAAAAAGAACCGGGCGCCCCTGCGCTGACAGCCGGAACACG
GCGGCATCAGAGCAGCCGATTGTCTGTTGTGCCCAGTCATAGCCGAATAGCCTCTCCACCCAAGCGGCCGGAGAACC
TGCGTGCAATCCATCTTGTTCAATCATGCGAAACGATCCAGATCCGGTGCAGATTATTTGGATTGAGAGTGAATATG
AGACTCTAATTGGATACCGAGGGGAATTTATGGAACGTCAGTGGAGCATTTTTGACAAGAAATATTTGCTAGCTGAT
AGTGACCTTAGGCGACTTTTGAACGCGCAATAATGGTTTCTGACGTATGTGCTTAGCTCATTAAACTCCAGAAACCC
GCGGCTGAGTGGCTCCTTCAACGTTGCGGTTCTGTCAGTTCCAAACGTAAAACGGCTTGTCCCGCGTCATCGGCGGG
GGTCATAACGTGACTCCCTTAATTCTCCGCTCATGGTACCACGCGTTTCGACAAAATTTAGAACGAACTTAATTATG
ATCTCAAATACATTGATACATATCTCATCTAGATCTAGGTTATCATTATGTAAGAAAGTTTTGACGAATATGGCACG
ACAAAATGGCTAGACTCGATGTAATTGGTATCTCAACTCAACATTATACTTATACCAAACATTAGTTAGACAAAATT
TAAACAACTATTTTTTATGTATGCAAGAGTCAGCATATGTATAATTGATTCAGAATCGTTTTGACGAGTTCGGATGT
AGTAGTAGCCATTATTTAATGTACATACTAATCGTGAATAGTGAATATGATGAAACATTGTATCTTATTGTATAAAT
ATCCATAAACACATCATGAAAGACACTTTCTTTCACGGTCTGAATTAATTATGATACAATTCTAATAGAAAACGAAT
TAAATTACGTTGAATTGTATGAAATCTAATTGAACAAGCCAACCACGACGACGACTAACGTTGCCTGGATTGACTCG
GTTTAAGTTAACCACTAAAAAAACGGAGCTGTCATGTAACACGCGGATCGAGCAGGTCACAGTCATGAAGCCATCAA
AGCAAAAGAACTAATCCAAGGGCTGAGATGATTAATTAGTTTAAAAATTAGTTAACACGAGGGAAAAGGCTGTCTGA
CAGCCAGGTCACGTTATCTTTACCTGTGGTCGAAATGATTCGTGTCTGTCGATTTTAATTATTTTTTTGAAAGGCCG
AAAATAAAGTTGTAAGAGATAAACCCGCCTATATAAATTCATATATTTCCTCTCCGCTTTGAAGTTTTAGTTTTAT
TGCAACAACAACAACAAATTACAATAACAACAAACAAAATACAAACAACAACAACATGGCACAATTTCAACAAACAA
TTGACATGCAAACTCTCCAAGCCGCTGCGGGACGCAACAGCTTGGTGAATGATTTGGCATCTCGTCGCGTTTACGAT
AATGCAGTCGAGGAGCTGAATGCTCGTTCCAGACGTCCCAAGGTAATAGGAACTTTCTGGATCTACTTTATTTGCTG
GATCTCGATCTTGTTTTCTCAATTTCCTTGAGATCTGGAATTCGTTTAATTTGGATCTGTGAACCTCCACTAAATCT
TTTGGTTTTACTAGAATCGATCTAAGTTGACCGATCAGTTAGCTCGATTATAGCTACCAGAATTTGGCTTGACCTTG
ATGGAGAGATCCATGTTCATGTTACCTGGGAAATGATTTGTATATGTGAATTGAAATCTGAACTGTTGAAGTTAGAT
TGAATCTGAACACTGTCAATGTTAGATTGAATCTGAACACTGTTTAAGGTTAGATGAAGTTTGTGTATAGATTCTTC
GAAACTTTAGGATTTGTAGTGTCGTACGTTGAACAGAAAGCTATTTCTGATTCAATCAGGGTTTATTTGACTGTATT
GAACTCTTTTTGTGTGTTTGCAGGTCCACTTCTCCAAGGCAGTGTCTACGGAACAGACCCTGATTGCAACAAACGCA
TATCCGGAGTTCGAGATTTCCTTTACTCATACGCAATCCGCTGTGCACTCCTTGGCCGGAGGCCTTCGGTCACTTGA
GTTGGAGTATCTCATGATGCAAGTTCCGTTCGGTTCTCTGACGTACGACATCGGCGGTAACTTTTCCGCGCACCTTT
TCAAAGGGCGCGATTACGTTCACTGCTGCATGCCTAATCTGGATGTACGTGACATTGCTCGCCATGAAGGACACAAG
GAAGCTATTTACAGTTATGTGAATCGTTTGAAAAGGCAGCAGCGTCCTGTGCCTGAATACCAGAGGGCAGCTTTCAA
CAACTACGCTGAGAACCCGCACTTCGTCCATTGCGACAAACCTTTCCAACAGTGTGAATTGACGACAGCGTATGGCA
CTGACACCTACGCTGTAGCTCTCCATAGCATTTATGATATCCCTGTTGAGGAGTTCGGTTCTGCGCTACTCAGGAAG
AATGTGAAAACTTGTTTCGCGGCCTTTCATTTCCATGAGAATATGCTTCTAGATTGTGATACAGTCACACTCGATGA
GATTGGAGCTACGTTCCAGAAATCAGGTAACATTCCTTAGTTACCTTTCTTTTCTTTTTCCATCATAAGTTTATAGA
TTGTACATGCTTTGAGATTTTTCTTTGCAAACAATCTCAGGTGATAACCTGAGCTTCTTCTTCCATAATGAGAGCAC
TCTCAATTACACCCACAGCTTCAGCAACATCATCAAGTACGTGTGCAAGACGTTCTTCCCTGCTAGTCAACGCTTCG
TGTACCACAAGGAGTTCCTGGTCACTAGAGTCAACACTTGGTACTGCAAGTTCACGAGAGTGGATACGTTCACTCTG
TTCCGTGGTGTGTACCACAACAATGTGGATTGCGAAGAGTTTTACAAGGCTATGGACGATGCGTGGCACTACAAAAA
GACGTTAGCAATGCTTAATGCCGAGAGGACCATCTTCAAGGATAACGCTGCGTTAAACTTCTGGTTCCCGAAGGTGC
TCTTGAAATTGGAAGTCTTCTTTTGTTGTCTAAACCTATCAATTTCTTTGCGGAAATTTATTTGAAGCTGTAGAGTT
AAAATTGAGTCTTTTAAACTTTTGTAGGTGAGAGACATGGTTATCGTCCCTCTCTTTGACGCTTCTATCACAACTGG

TAGGATGTCTAGGAGAGAGGTTATGGTGAACAAGGACTTCGTCTACACGGTCCTAAATCACATCAAGACCTATCAAG
CTAAGGCACTGACGTACGCAAACGTGCTGAGCTTCGTGGAGTCTATTAGGTCTAGAGTGATAATTAACGGTGTCACT
GCCAG*GTAAGTTGTTACTTATGATTGTTTTCCTCTCTGCTACATGTATTTTGTTGTTCATTTCTGTAAGATATAAGA*
*ATTGAGTTTTCCTCTGATGATATTATTAG*GTCTGAATGGGACACAGACAAGGCAATTCTAGGTCCATTAGCAATGAC
ATTCTTCCTGATCACGAAGCTGGGTCATGTGCAAGATGAAATAATCCTGAAAAAGTTCCAGAAGTTCGACAGAACCA
CCAATGAGCTGATTTGGACAAGTCTCTGCGATGCCCTGATGGGGGTTATTCCCTCGGTCAAGGAGACGCTTGTGCGC
GGTGGTTTTGTGAAAGTAGCAGAACAAGCCTTAGAGATCAAG*GTTAGTATCATATGAAGAAATACCTAGTTTCAGTT*
*GATGAATGCTATTTTCTGACCTCTTTTCTCGAAGCCGCGGTGCGGGTGCCAGGGCGTGCCCTTGGGCTCCCCGGGCG*
*CGTACTCCACCTCACCCATCTTTTGATCAACATCAAAATTAGGTTCAATTTTCATCAACCAAATAATATTTTTCATG*
*TATATATAG*GTCACAGAAAAACGACCTTGAAAGATTATACGGCCGGAATCAAAACATGTCTTTGGTATCAAAGGAAA
AGTGGTGATGTGACAACCTTTATTGGTAATACCATCATCATTGCCGCATGTTTGAGCTCAATGATCCCCATGGACAA
AGTGATAAAGGCAGCTTTTTGTGGAGACGATAGCCTGATTTACATTCCTAAAGGTTTAGACTTGCCTGATATTCAGG
CGGGCGCGAACCTCATGTGGAACTTCGAGGCCAAACTCTTCAGGAAGAAGTATGGTTACTTCTGTGGTCGTTATGTT
ATTCACCATGATAGAGGAGCCATTGTGTATTACGATCCGCTTAAACTAATATCTAAGTTAGGTTGTAAACATATTAG
AGATGTTGTTCACTTAGAAGAGTTACGCGAGTCTTTGTGTGATGTAGCTAGTAACTTAAATAATTGTGCGTATTTTT
CACAGTTAGATGAGGCCGTTGCCGAGGTTCATAAGACCGCGGTAGGCGGTTCGTTTGCTTTTTGTAGTATAATTAAG
TATTTGTCAGATAAGAGATTGTTTAGAGATTTGTTCTTTGTTTGATAATGTCGATAGTCTCGTACGAACCTAAGGTG
AGTGATTTCCTCAATCTTTCGAAGAAGGAAGAGATCTTGCCGAAGGCTCTAACGAGGTTAAAAACCGTGTCTATTAG
TACTAAAGATATTATATCTGTCAAGGAGTCGGAGACTTTGTGTGATATAGATTTGTTAATCAATGTGCCATTAGATA
AGTATAGATATGTGGGTATCCTAGGAGCCGTTTTTACCGGAGAGTGGCTAGTGCCAGACTTCGTTAAAGGTGGAGTG
ACGATAAGTGTGATAGATAAGCGTCTGGTGAACTCAAAGGAGTGCGTGATTGGTACGTACAGAGCCGCAGCCAAGAG
TAAGAGGTTCCAGTTCAAATTGGTTCCAAATTACTTTGTGTCCACCGTGGACGCAAAGAGGAAGCCGTGGCAG*GTAA*
*GGATTTTTATGATATAGTATGCTTATGTATTTTGTACTGAAAGCATATCCTGCTTCATTGGGATATTACTGAAAGCA*
*TTTAACTACATGTAAACTCACTTGATGATCAATAAACTTGATTTTGCAG*GTTCATGTTCGTATACAAGACTTGAAGA
TTGAGGCGGGTTGGCAGCCGTTAGCTCTGGAAGTAGTTTCAGTTGCTATGGTCACCAATAACGTTGTCATGAAGGGT
TTGAGGGAAAAGGTCGTCGCAATAAATGATCCGGACGTCGAAGGTTTCGAAG*GTAAGCCATCTTCCTGCTTATTTTT*
*ATAATGAACATAGAAATAGGAAGTTGTGCAGAGAAACTAATTAACCTGACTCAAAATCTACCCTCATAATTGTTGTT*
*TGATATTGGTCTTGTATTTTGCAG*GTGTGGTTGACGAATTCGTCGATTCGGTTGCAGCATTTAAAGCGGTTGACAAC
TTTAAAAGAAGGAAAAAGAAGGTTGAAGAAAAGGGTGTAGTAAGTAAGTATAAGTACAGACCGGAGAAGTACGCCGG
TCCTGATTCGTTTAATTTGAAAGAAGAAAACGTCTTACAACATTACAAACCCGAATCAGTACCAGTATTTCGATAAG
AAACAAGAAAGGTATGGTGAGCAAGGGCGAGGAGCTGTTCACCGGGGTGGTGCCCATCCTGGTCGAGCTGGACGGCG
ACGTAAACGGCCACAAGTTCAGCGTGTCCGGCGAGGGCGAGGGCGATGCCACCTACGGCAAGCTGACCCTGAAGTTC
ATCTGCACCACCGGCAAGCTGCCCGTGCCCTGGCCCACCCTCGTGACCACCTTCAGCTACGGCGTGCAGTGCTTCAG
CCGCTACCCCGACCACATGAAGCAGCACGACTTCTTCAAGTCCGCCATGCCCGAAGGCTACGTCCAGGAGCGCACCA
TCTTCTTCAAGGACGACGGCAACTACAAGACCCGCGCCGAGGTGAAGTTCGAGGGCGACACCCTGGTGAACCGCATC
GAGCTGAAGGGCATCGACTTCAAGGAGGACGGCAACATCCTGGGGCACAAGCTGGAGTACAACTACAACAGCCACAA
CGTCTATATCATGGCCGACAAGCAGAAGAACGGCATCAAGGTGAACTTCAAGATCCGCCACAACATCGAGGACGGCA
GCGTGCAGCTCGCCGACCACTACCAGCAGAACACCCCCATCGGCGACGGCCCCGTGCTGCTGCCCGACAACCACTAC
CTGAGCACCCAGTCCGCCCTGAGCAAAGACCCCAACGAGAAGCGCGATCACATGGTCCTGCTGGAGTTCGTGACCGC
CGCCGGGATCACTCACGGCATGGACGAGCTGTACAAGTAAGCTTACTAGAGCGTGGTGCGCACGATAGCGCATAGTG
TTTTTCTCTCCACTTGAATCGAAGAGATAGACTTACGGTGTAAATCCGTAGGGGTGGCGTAAACCAAATTACGCAAT
GTTTTGGGTTCCATTTAAATCGAAACCCCTTATTTCCTGGATCACCTGTTAACGCACGTTTGACGTGTATTACAGTG
GGAATAAGTAAAAGTGAGAGGTTCGAATCCTCCCTAACCCCGGGTAGGGGCCCAGCGGCCGCTCTAGCTAGAGTCAA
GCAGATCGTTCAAACATTTGGCAATAAAGTTTCTTAAGATTGAATCCTGTTGCCGGTCTTGCGATGATTATCATATA
ATTTCTGTTGAATTACGTTAAGCATGTAATAATTAACATGTAATGCATGACGTTATTTATGAGATGGGTTTTTATGA
TTAGAGTCCCGCAATTATACATTTAATACGCGATAGAAAACAAAATATAGCGCGCAAACTAGGATAAATTATCGCGC
GCGGTGTCATCTATGTTACTAGATCGACCTGCATCCACCCCAGTACATTAAAAACGTCCGCAATGTGTTATTAAGTT
GTCTAAGCGTCAATTTGTTTACACCACAATATATCCTGCCACCAGCCAGCCAACAGCTCCCCGACCGGCAGCTCGGC
ACAAAATCACCACTCGATACAGGCAGCCCATCAG

**TMV-Donor**

**pAGT6917 (Donor: 5´HA – 1348bp; 3´HA – 777bp)**

GG-overhang_sgRNA target D1 (**PAM**)_(5´HA(1348 bp)); RdRP(*intron*)- MP; (3´HA(777bp))_ sgRNA target D1 (**PAM**)_GG-overhang

GGAGGGAAAGCTTCATCAAGCTCTTGTTGCTCAAGG**AGG**AGATAGAATTGGCTTGGTAATTTACGTTCTGTACGCGA
CGACTGAATACGACAGCGAATGGCACTGACACCTACGCTGTAGCTCTCCATAGCATTTATGATATCCCTGTTGAGGA
GTTCGGTTCTGCGCTACTCAGGAAGAATGTGAAAACTTGTTTCGCGGCCTTTCATTTCCATGAGAATATGCTTCTAG
ATTGTGATACAGTCACACTCGATGAGATTGGAGCTACGTTCCAGAAATCAG*GTAACATTCCTTAGTTACCTTTCTTT*

TCTTTTTCCATCATAAGTTTATAGATTGTACATGCTTTGAGATTTTTCTTTGCAAACAATCTCAGGTGATAACCTGA
GCTTCTTCTTCCATAATGAGAGCACTCTCAATTACACCCACAGCTTCAGCAACATCATCAAGTACGTGTGCAAGACG
TTCTTCCCTGCTAGTCAACGCTTCGTGTACCACAAGGAGTTCCTGGTCACTAGAGTCAACACTTGGTACTGCAAGTT
CACGAGAGTGGATACGTTCACTCTGTTCCGTGGTGTGTACCACAACAATGTGGATTGCGAAGAGTTTTACAAGGCTA
TGGACGATGCGTGGCACTACAAAAAGACGTTAGCAATGCTTAATGCCGAGAGGACCATCTTCAAGGATAACGCTGCG
TTAAACTTCTGGTTCCCGAAGGTGCTCTTGAAATTGGAACTCTTCTTTTGTTGTCTAAACCTATCAATTTCTTTGCG
GAAATTTATTTGAAGCTGTAGAGTTAAAATTGAGTCTTTTAAACTTTTGTAGGTGAGAGACATGGTTATCGTCCCTC
TCTTTGACGCTTCTATCACAACTGGTAGGATGTCTAGGAGAGAGGTTATGGTGAACAAGGACTTCGTCTACACGGTC
CTAAATCACATCAAGACCTATCAAGCTAAGGCACTGACGTACGCAAACGTGCTGAGCTTCGTGGAGTCTATTAGGTC
TAGAGTGATAATTAACGGTGTCACTGCCAGGTAAGTTGTTACTTATGATTGTTTTCCTCTCTGCTACATGTATTTTG
TTGTTCATTTCTGTAAGATATAAGAATTGAGTTTTCCTCTGATGATATTATTAGGTCTGAATGGGACACAGACAAGG
CAATTCTAGGTCCATTAGCAATGACATTCTTCCTGATCACGAAGCTGGGTCATGTGCAAGATGAAATAATCCTGAAA
AAGTTCCAGAAGTTCGACAGAACCACCAATGAGCTGATTTGGACAAGTCTCTGCGATGCCCTGATGGGGGTTATTCC
CTCGGTCAAGGAGACACTTGTGCGCGGTGGTTTTGTGAAAGTAGCAGAACAAGCCTTAGAGATCAAGGTTAGTATCA
TATGAAGAAATACCTAGTTTCAGTTGATGAATGCTATTTTCTGACCTCAGTTGTTCTCTTTTGAGAATTATTTCTTT
TCTAATTTGCCTGATTTTTCTATTAATTCATTAGGTTCCCGAGCTATACTGTACCTTCGCCGACCGATTGGTACTAC
AGTACAAGAAGGCGGAGGAGTTCCAATCGTGTGATCTTTCCAAACCTCTAGAAGAGTCAGAGAAGTACTACAACGCA
TTATCCGAGCTATCAGTGCTTGAGAATCTCGACTCTTTTGACTTAGAGGCGTTTAAGACTTTATGTCAGCAGAAGAA
TGTGGACCCGGATATGGCAGCAAAGGTAAATCCTGGTCCACACTTTTACGATAAAAACACAAGATTTTAAACTATGA
ACTGATCAATAATCATTCCTAAAAGACCACACTTTTGTTTTGTTTCTAAAGTAATTTTTACTGTTATAACAGGTGGT
CGTAGCAATCATGAAGTCAGAATTGACGTTGCCTTTCAAGAAACCTACAGAAGAGGAAATCTCGGAGTCGCTAAAAC
CAGGAGAGGGGTCGTGTGCAGAGCATAAGGAAGTGTTGAGCTTACAAAATGATGCTCCGTTCCCGTGTGTGAAAAAT
CTAGTTGAAGGTTCCGTGCCGGCGTATGGAATGTGTCCTAAGGGTGGTGGTTTCGACAAATTGGATGTGGACATTGC
TGATTTCCATCTCAAGAGTGTAGATGCAGTTAAAAAGGGAACTATGATGTCTGCGGTGTACACAGGGTCTATCAAAG
TTCAACAAATGAAGAACTACATAGATTACTTAAGTGCGTCGCTGGCAGCTACAGTCTCAAACCTCTGCAAGGTAAGA
GGTCAAAAGGTTTCCGCAATGATCCCTCTTTTTTTGTTTCTCTAGTTTCAAGAATTTGGGTATATGACTAACTTCTG
AGTGTTCCTTGATGCATATTTGTGATGAGACAAATGTTTGTTCTATGTTTTAGGTGCTTAGAGATGTTCACGGCGTT
GACCCAGAGTCACAGGAGAAATCTGGAGTGTGGGATGTTAGGAGAGGACGTTGGTTACTTAAACCTAATGCGAAAAG
TCACGCGTGGGGTGTGGCAGAAGATGCCAACCACAAGTTGGTTATTGTGTTACTCAACTGGGATGACGGAAAGCCGG
TTTGTGATGAGACATGGTTCAGGGTGGCGGTGTCAAGCGATTCCTTGATATATTCGGATATGGGAAAACTTAAGACG
CTCACGTCTTGCAGTCCAAATGGTGAGCCACCGGAGCCTAACGCCAAAGTAATTTTGGTCGATGGTGTTCCCGGTTG
TGGAAAAACGAAGGAGATTATCGAAAAGGTAAGTTCTGCATTTGGTTATGCTCCTTGCATTTTAGGTGTTCGTCGCA
CTTCCATTTCCATGAATAGCTAAGATTTTTTTTCTCTGCATTCATTCTTCTTGCCTCAGTTCTAACTGTTTGTGGTA
TTTTTGTTTTAATTATTGCTACAGGTAAACTTCTCTGAGGACTTGATCTTAGTCCCTGGGAAGGAAGCTTCTAAGAT
GATCATCCGGAGGGCCAACCAAGCTGGTGTGATAAGAGCGGATAAGGACAATGTTAGAACGGTGGATTCCTTCTTGA
TGCATCCTTCTAGAAGGGTGTTTAAGAGGTTGTTTATCGATGAAGGACTAATGCTGCATACAGGTTGTGTAAATTTC
CTACTGCTGCTATCTCAATGTGACGTCGCATATGTGTATGGGGACACAAAGCAAATTCCGTTCATTTGCAGAGTCGC
GAACTTTCCGTATCCAGCGCATTTTGCAAAACTCGTCGCTGATGAGAAGGAAGTCAGAAGAGTTACGCTCAGGTAAA
GCAACTGTGTTTTAATCAATTTCTTGTCAGGATATATGGATTATAACTTAATTTTTGAGAAATCTGTAGTATTTGGC
GTGAAATGAGTTTGCTTTTTGGTTTCTCCCGTGTTATAGGTGCCCGGCTGATGTTACGTATTTCCTTAACAAGAAGT
ATGACGGGGCGGTGATGTGTACCAGCGCGGTAGAGAGATCCGTGAAGGCAGAAGTGGTGAGAGGAAAGGGTGCATTG
AACCCAATAACCTTACCGTTGGAGGGTAAAATTTTGACCTTCACACAAGCTGACAAGTTCGAGTTACTGGAGAAGGG
TTACAAGGTAAAGTTTCCAACTTTCCTTTACCATATCAAACTAAAGTTCGAAACTTTTTATTTGATCAACTTCAAGG
CCACCCGATCTTTCTATTCCTGATTAATTTGTGATGAATCCATATTGACTTTTGATGGTTACGCAGGATGTGAACAC
TGTGCACGAGGTGCAAGGGGAGACATACGAGAAAACTGCTATTGTGCGCTTGACATCAACTCCGTTAGAGATCATAT
CGAGTGCGTCACCTCATGTTTTGGTGGCGCTGACAAGACACACAACGTGTTGTAAATATTACACCGTTGTGTTGGAC
CCGATGGTGAATGTGATTTCAGAAATGGAGAAGTTGTCCAATTTCCTTCTTGACATGTATAGAGTTGAAGCAGGTCT
GTCTTTCCTATTTCATATGTTTAATCCTAGGAATTTGATCAATTGATTGTATGTATGTCGATCCCAAGACTTTCTTG
TTCACTTATATCTTAACTCTCTCTTTGCTGTTTCTTGCAGGTGTCCAATAGCAATTACAAATCGATGCAGTATTCAG
GGGACAGAACTTGTTTGTTCAGACGCCCAAGTCAGGAGATTGGCGAGATATGCAATTTTACTATGACGCACTTCTTC
CCGGAAACAGTACTATTCTCAATGAATTTGATGCTGTTACGATGAATTTGAGGGATATTTCCTTAAACGTCAAAGAT
TGCAGAATCGACTTCTCCAAATCCGTGCAACTTCCTAAAGAACAACCTATTTTCCTCAAGCCTAAAATAAGAACTGC
GGCAGAAATGCCGAGAACTGCAGGTAAAATATTGGATGCCAGACGATATTCTTTCTTTTGATTTGTAACTTTTTCCT
GTCAAGGTCGATAAATTTTATTTTTTTTGGTAAAAGGTCGATAATTTTTTTTTGGAGCCATTATGTAATTTTCCTAA
TTAACTGAACCAAAATTATACAAACCAGGTTTGCTGGAAAATTTGGTTGCAATGATCAAAAGAAACATGAATGCGCC
GGATTTGACAGGGACAATTGACATTGAGGATACTGCATCTCTGGTGGTTGAAAAGTTTTGGGATTCGTATGTTGACA
AGGAATTTAGTGGAACGAACGAAATGACCATGACAAGGGAGAGCTTCTCCAGGTAAGGACTTCTCATGAATATTAGT
GGCAGATTAGTGTTGTTAAAGTCTTTGGTTAGATAATCGATGCCTCCTAATTGTCCATGTTTTACTGGTTTTCTACA

*ATTAAAG*GTGGCTTTCGAAACAAGAGTCATCTACAGTTGGTCAGTTAGCGGACTTTAACTTTGTGGATTTGCCGGCA
GTAGATGAGTACAAGCATATGATCAAGAGTCAACCAAAGCAAAAGTTAGACTTGAGTATTCAAGACGAATATCCTGC
ATTGCAGACGATAGTCTACCATTCGAAAAAGATCAATGCGATTTTCGGTCCAATGTTTTCAGAACTTACGAGGATGT
TACTCGAAAGGATTGACTCTTCGAAGTTTCTGTTCTACACCAGAAAGACACCTGCACAAATAGAGGACTTCTTTTCT
GACCTAGACTCAACCCAGGCGATGGAAATTCTGGAACTCGACATTTCGAAGTACGATAAGTCACAAAACGAGTTCCA
TTGTGCTGTAGAGTACAAGATCTGGGAAAAGTTAGGAATTGATGAGTGGCTAGCTGAGGTCTGGAAACAAG*GTGAGT*
*TCCTAAGTTCCATTTTTTTGTAATCCTTCAATGTTATTTTAACTTTTCAGATCAACATCAAAATTAGGTTCAATTTT*
*CATCAACCAAATAATATTTTTCATGTATATATAG*GTCACAGAAAAACGACCTTGAAAGATTATACGGCCGGAATCAA
AACATGTCTTTGGTATCAAAGGAAAAGTGGTGATGTGACAACCTTTATTGGTAATACCATCATCATTGCCGCATGTT
TGAGCTCAATGATCCCCATGGACAAAGTGATAAAGGCAGCTTTTTGTGGCGACGATAGCCTGATTTACATTCCTAAA
GGTTTAGACTTGCCTGATATTCAGGCGGGCGCGAACCTCATGTGGAACTTCGAGGCCAAACTCTTCAGGAAGAAGTA
TGGTTACTTCTGTGGTCGTTATGTTATTCACCATGATAGAGGAGCCATTGTGTATTACGATCCGCTTAAACTAATAT
CTAAGTTAGGTTGTAAACATATTAGAGATGTTGTTCACTTAGAAGAGTTACGCGAGTCTTTGTGTGATGTAGCTAGT
AACTTAAATAATTGTGCGTATTTTTCACAGTTAGATGAGGCCGTTGCCGAGGTTCATAAGACCGCGGTAGGCGGTTC
GTTTGCTTTTTGTAGTATAATTAAGTATTTGTCAGATAAGAGATTGTTTAGAGATTTGTTCTTTGTTTGATAATGTC
GATAGTCTCGTACGAACCTAAGGTGAGTGATTTCCTCAATCTTTCGAAGAAGGAAGAGATCTTGCCGAAGGCTCTAA
CGAGGTTAAAAACCGTGTCTATTAGTACTAAAGATATTATATCTGTCAAGGAGTCGTGCTTGGTAATTTAGTCGCGC
GATTGTACCACTGGAAAGCTTCATCAAGCTCTTGTTGCTCAAGG**AGG**AGATAGAATTGGCGCT

**pAGT7390 (Donor: 5´HA – 500bp; 3´HA – 500bp)**

GG-overhang_sgRNA target D1 (**PAM**)_(5´HA(500 bp)); RdRP(*intron*); (3´HA(500bp))_sgRNA target D1 (**PAM**)_GG-overhang

GGAGGGAAAGCTTCATCAAGCTCTTGTTGCTCAAGG**AGG**AGATAGAATTGGCTTGGTAATTTACGTTCTGTACGCGA
CGACTGAATTCAAGACCTATCAAGCTAAGGCACTGACGTACGCAAACGTGCTGAGCTTCGTGGAGTCTATTAGGTCT
AGAGTGATAATTAACGGTGTCACTGCCAG*GTAAGTTGTTACTTATGATTGTTTTCCTCTCTGCTACATGTATTTTGT*
*TGTTCATTTCTGTAAGATATAAGAATTGAGTTTTCCTCTGATGATATTATTAG*GTCTGAATGGGACACAGACAAGGC
AATTCTAGGTCCATTAGCAATGACATTCTTCCTGATCACGAAGCTGGGTCATGTGCAAGATGAAATAATCCTGAAAA
AGTTCCAGAAGTTCGACAGAACCACCAATGAGCTGATTTGGACAAGTCTCTGCGATGCCCTGATGGGGGTTATTCCC
TCGGTCAAGGAGACACTTGTGCGCGGTGGTTTTGTGAAAGTAGCAGAACAAGCCTTAGAGATCAAG*GTTAGTATCAT*
*ATGAAGAAATACCTAGTTTCAGTTGATGAATGCTATTTTCTGACCTCAGTTGTTCTCTTTTGAGAATTATTTCTTTT*
*CTAATTTGCCTGATTTTTCTATTAATTCATTAG*GTTCCCGAGCTATACTGTACCTTCGCCGACCGATTGGTACTACA
GTACAAGAAGGCGGAGGAGTTCCAATCGTGTGATCTTTCCAAACCTCTAGAAGAGTCAGAGAAGTACTACAACGCAT
TATCCGAGCTATCAGTGCTTGAGAATCTCGACTCTTTTGACTTAGAGGCGTTTAAGACTTTATGTCAGCAGAAGAAT
GTGGACCCGGATATGGCAGCAAAG*GTAAATCCTGGTCCACACTTTTACGATAAAAACACAAGATTTTAAACTATGAA*
*CTGATCAATAATCATTCCTAAAAGACCACACTTTTGTTTTGTTTCTAAAGTAATTTTTACTGTTATAACAG*GTGGTC
GTAGCAATCATGAAGTCAGAATTGACGTTGCCTTTCAAGAAACCTACAGAAGAGGAAATCTCGGAGTCGCTAAAACC
AGGAGAGGGGTCGTGTGCAGAGCATAAGGAAGTGTTGAGCTTACAAAATGATGCTCCGTTCCCGTGTGTGAAAAATC
TAGTTGAAGGTTCCGTGCCGGCGTATGGAATGTGTCCTAAGGGTGGTGGTTTCGACAAATTGGATGTGGACATTGCT
GATTTCCATCTCAAGAGTGTAGATGCAGTTAAAAAGGGAACTATGATGTCTGCGGTGTACACAGGGTCTATCAAAGT
TCAACAAATGAAGAACTACATAGATTACTTAAGTGCGTCGCTGGCAGCTACAGTCTCAAACCTCTGCAAG*GTAAGAG*
*GTCAAAAGGTTTCCGCAATGATCCCTCTTTTTTTGTTTCTCTAGTTTCAAGAATTTGGGTATATGACTAACTTCTGA*
*GTGTTCCTTGATGCATATTTGTGATGAGACAAATGTTTGTTCTATGTTTTAG*GTGCTTAGAGATGTTCACGGCGTTG
ACCCAGAGTCACAGGAGAAATCTGGAGTGTGGGATGTTAGGAGAGGACGTTGGTTACTTAAACCTAATGCGAAAAGT
CACGCGTGGGGTGTGGCAGAAGATGCCAACCACAAGTTGGTTATTGTGTTACTCAACTGGGATGACGGAAAGCCGGT
TTGTGATGAGACATGGTTCAGGGTGGCGGTGTCAAGCGATTCCTTGATATATTCGGATATGGGAAAACTTAAGACGC
TCACGTCTTGCAGTCCAAATGGTGAGCCACCGGAGCCTAACGCCAAAGTAATTTTGGTCGATGGTGTTCCCGGTTGT
GGAAAAACGAAGGAGATTATCGAAAAG*GTAAGTTCTGCATTTGGTTATGCTCCTTGCATTTTAGGTGTTCGTCGCAC*
*TTCCATTTCCATGAATAGCTAAGATTTTTTTTCTCTGCATTCATTCTTCTTGCCTCAGTTCTAACTGTTTGTGGTAT*
*TTTTGTTTTAATTATTGCTACAG*GTAAACTTCTCTGAGGACTTGATCTTAGTCCCTGGGAAGGAAGCTTCTAAGATG
ATCATCCGGAGGGCCAACCAAGCTGGTGTGATAAGAGCGGATAAGGACAATGTTAGAACGGTGGATTCCTTCTTGAT
GCATCCTTCTAGAAGGGTGTTTAAGAGGTTGTTTATCGATGAAGGACTAATGCTGCATACAGGTTGTGTAAATTTCC
TACTGCTGCTATCTCAATGTGACGTCGCATATGTGTATGGGGACACAAAGCAAATTCCGTTCATTTGCAGAGTCGCG
AACTTTCCGTATCCAGCGCATTTTGCAAAACTCGTCGCTGATGAGAAGGAAGTCAGAAGAGTTACGCTCAG*GTAAAG*
*CAACTGTGTTTTAATCAATTTCTTGTCAGGATATATGGATTATAACTTAATTTTTGAGAAATCTGTAGTATTTGGCG*
*TGAAATGAGTTTGCTTTTTGGTTTCTCCCGTGTTATAG*GTGCCCGGCTGATGTTACGTATTTCCTTAACAAGAAGTA
TGACGGGGCGGTGATGTGTACCAGCGCGGTAGAGAGATCCGTGAAGGCAGAAGTGGTGAGAGGAAAGGGTGCATTGA
ACCCAATAACCTTACCGTTGGAGGGTAAAATTTTGACCTTCACACAAGCTGACAAGTTCGAGTTACTGGAGAAGGGT
TACAAG*GTAAAGTTTCCAACTTTCCTTTACCATATCAAACTAAAGTTCGAAACTTTTTATTTGATCAACTTCAAGGC*

*CACCCGATCTTTCTATTCCTGATTAATTTGTGATGAATCCATATTGACTTTTGATGGTTACGCAG*GATGTGAACACT
GTGCACGAGGTGCAAGGGGAGACATACGAGAAAACTGCTATTGTGCGCTTGACATCAACTCCGTTAGAGATCATATC
GAGTGCGTCACCTCATGTTTTGGTGGCGCTGACAAGACACACAACGTGTTGTAAATATTACACCGTTGTGTTGGACC
CGATGGTGAATGTGATTTCAGAAATGGAGAAGTTGTCCAATTTCCTTCTTGACATGTATAGAGTTGAAGCAG*GTCTG*
*TCTTTCCTATTTCATATGTTTAATCCTAGGAATTTGATCAATTGATTGTATGTATGTCGATCCCAAGACTTTCTTGT*
*TCACTTATATCTTAACTCTCTCTTTGCTGTTTCTTGCAG*GTGTCCAATAGCAATTACAAATCGATGCAGTATTCAGG
GGACAGAACTTGTTTGTTCAGACGCCCAAGTCAGGAGATTGGCGAGATATGCAATTTTACTATGACGCACTTCTTCC
CGGAAACAGTACTATTCTCAATGAATTTGATGCTGTTACGATGAATTTGAGGGATATTTCCTTAAACGTCAAAGATT
GCAGAATCGACTTCTCCAAATCCGTGCAACTTCCTAAAGAACAACCTATTTTCCTCAAGCCTAAAATAAGAACTGCG
GCAGAAATGCCGAGAACTGCAG*GTAAAATATTGGATGCCAGACGATATTCTTTCTTTTGATTTGTAACTTTTTCCTG*
*TCAAGGTCGATAAATTTTATTTTTTTTGGTAAAAGGTCGATAATTTTTTTTTGGAGCCATTATGTAATTTTCCTAAT*
*TAACTGAACCAAAATTATACAAACCAG*GTTTGCTGGAAAATTTGGTTGCAATGATCAAAAGAAACATGAATGCGCCG
GATTTGACAGGGACAATTGACATTGAGGATACTGCATCTCTGGTGGTTGAAAAGTTTTGGGATTCGTATGTTGACAA
GGAATTTAGTGGAACGAACGAAATGACCATGACAAGGGAGAGCTTCTCCAG*GTAAGGACTTCTCATGAATATTAGTG*
*GCAGATTAGTGTTGTTAAAGTCTTTGGTTAGATAATCGATGCCTCCTAATTGTCCATGTTTTACTGGTTTTCTACAA*
*TTAAAG*GTGGCTTTCGAAACAAGAGTCATCTACAGTTGGTCAGTTAGCGGACTTTAACTTTGTGGATTTGCCGGCAG
TAGATGAGTACAAGCATATGATCAAGAGTCAACCAAAGCAAAAGTTAGACTTGAGTATTCAAGACGAATATCCTGCA
TTGCAGACGATAGTCTACCATTCGAAAAAGATCAATGCGATTTTCGGTCCAATGTTTTCAGAACTTACGAGGATGTT
ACTCGAAAGGATTGACTCTTCGAAGTTTCTGTTCTACACCAGAAAGACACCTGCACAAATAGAGGACTTCTTTTCTG
ACCTAGACTCAACCCAGGCGATGGAAATTCTGGAACTCGACATTTCGAAGTACGATAAGTCACAAAACGAGTTCCAT
TGTGCTGTAGAGTACAAGATCTGGGAAAAGTTAGGAATTGATGAGTGGCTAGCTGAGGTCTGGAAACAAG*GTGAGTT*
*CCTAAGTTCCATTTTTTTGTAATCCTTCAATGTTATTTTAACTTTTCAGATCAACATCAAAATTAGGTTCAATTTTC*
*ATCAACCAAATAATATTTTTCATGTATATATAG*GTCACAGAAAAACGACCTTGAAAGATTATACGGCCGGAATCAAA
ACATGTCTTTGGTATCAAAGGAAAAGTGGTGATGTGACAACCTTTATTGGTAATACCATCATCATTGCCGCATGTTT
GAGCTCAATGATCCCCATGGACAAAGTGATAAAGGCAGCTTTTTGTGGCGACGATAGCCTGATTTACATTCCTAAAG
GTTTAGACTTGCCTGATATTCAGGCGGGCGCGAACCTCATGTGGAACTTCGAGGCCAAACTCTTCAGGAAGAAGTAT
GGTTACTTCTGTGGTCGTTATGTTATTCACCATGATAGAGGAGCCATTGTGTATTACGATCCGCTTAAACTAATATC
TAAGTTAGGTTGTAAACATATTAGAGATGTTGTTCACTTAGAAGAGTTACGCGAGTCTTTGTGTGATGTAGCTAGTA
ACTTAAATATGCTTGGTAATTTAGTCGCGCGATTGTACCACTGGAAAGCTTCATCAAGCTCTTGTTGCTCAAGG**AGG**
AGATAGAATTGGCGCT

**pAGT7391 (Donor: 5´HA – 250bp; 3´HA – 250bp)**
GG-overhang_sgRNA target D1 (**PAM**)_(5´HA(250 bp)); RdRP(*intron*); (3´HA(250bp))_ sgRNA target D1 (**PAM**)_GG-overhang
GGAGGGAAAGCTTCATCAAGCTCTTGTTGCTCAAGG**AGG**AGATAGAATTGGCTTGGTAATTTACGTTCTGTACGCGA
CGACTGAATTTCCTGATCACGAAGCTGGGTCATGTGCAAGATGAAATAATCCTGAAAAAGTTCCAGAAGTTCGACAG
AACCACCAATGAGCTGATTTGGACAAGTCTCTGCGATGCCCTGATGGGGGTTATTCCCTCGGTCAAGGAGACACTTG
TGCGCGGTGGTTTTGTGAAAGTAGCAGAACAAGCCTTAGAGATCAAG*GTTAGTATCATATGAAGAAATACCTAGTTT*
*CAGTTGATGAATGCTATTTTCTGACCTCAGTTGTTCTCTTTTGAGAATTATTTCTTTTCTAATTTGCCTGATTTTTC*
*TATTAATTCATTAG*GTTCCCGAGCTATACTGTACCTTCGCCGACCGATTGGTACTACAGTACAAGAAGGCGGAGGAG
TTCCAATCGTGTGATCTTTCCAAACCTCTAGAAGAGTCAGAGAAGTACTACAACGCATTATCCGAGCTATCAGTGCT
TGAGAATCTCGACTCTTTTGACTTAGAGGCGTTTAAGACTTTATGTCAGCAGAAGAATGTGGACCCGGATATGGCAG
CAAAG*GTAAATCCTGGTCCACACTTTTACGATAAAAACACAAGATTTTAAACTATGAACTGATCAATAATCATTCCT*
*AAAAGACCACACTTTTGTTTTGTTTCTAAAGTAATTTTTACTGTTATAACAG*GTGGTCGTAGCAATCATGAAGTCAG
AATTGACGTTGCCTTTCAAGAAACCTACAGAAGAGGAAATCTCGGAGTCGCTAAAACCAGGAGAGGGGTCGTGTGCA
GAGCATAAGGAAGTGTTGAGCTTACAAAATGATGCTCCGTTCCCGTGTGTGAAAAATCTAGTTGAAGGTTCCGTGCC
GGCGTATGGAATGTGTCCTAAGGGTGGTGGTTTCGACAAATTGGATGTGGACATTGCTGATTTCCATCTCAAGAGTG
TAGATGCAGTTAAAAAGGGAACTATGATGTCTGCGGTGTACACAGGGTCTATCAAAGTTCAACAAATGAAGAACTAC
ATAGATTACTTAAGTGCGTCGCTGGCAGCTACAGTCTCAAACCTCTGCAAG*GTAAGAGGTCAAAAGGTTTCCGCAAT*
*GATCCCTCTTTTTTTGTTTCTCTAGTTTCAAGAATTTGGGTATATGACTAACTTCTGAGTGTTCCTTGATGCATATT*
*TGTGATGAGACAAATGTTTGTTCTATGTTTTAG*GTGCTTAGAGATGTTCACGGCGTTGACCCAGAGTCACAGGAGAA
ATCTGGAGTGTGGGATGTTAGGAGAGGACGTTGGTTACTTAAACCTAATGCGAAAAGTCACGCGTGGGGTGTGGCAG
AAGATGCCAACCACAAGTTGGTTATTGTGTTACTCAACTGGGATGACGGAAAGCCGGTTTGTGATGAGACATGGTTC
AGGGTGGCGGTGTCAAGCGATTCCTTGATATATTCGGATATGGGAAAACTTAAGACGCTCACGTCTTGCAGTCCAAA
TGGTGAGCCACCGGAGCCTAACGCCAAAGTAATTTGGTCGATGGTGTTCCCGGTTGTGGAAAAACGAAGGAGATTA
TCGAAAAG*GTAAGTTCTGCATTTGGTTATGCTCCTTGCATTTTAGGTGTTCGTCGCACTTCCATTTCCATGAATAGC*
*TAAGATTTTTTTTCTCTGCATTCATTCTTCTTGCCTCAGTTCTAACTGTTTGTGGTATTTTTGTTTTAATTATTGCT*
*ACAG*GTAAACTTCTCTGAGGACTTGATCTTAGTCCCTGGGAAGGAAGCTTCTAAGATGATCATCCGGAGGGCCAACC

AAGCTGGTGTGATAAGAGCGGATAAGGACAATGTTAGAACGGTGGATTCCTTCTTGATGCATCCTTCTAGAAGGGTG
TTTAAGAGGTTGTTTATCGATGAAGGACTAATGCTGCATACAGGTTGTGTAAATTTCCTACTGCTGCTATCTCAATG
TGACGTCGCATATGTGTATGGGGACACAAAGCAAATTCCGTTCATTTGCAGAGTCGCGAACTTTCCGTATCCAGCGC
ATTTTGCAAAACTCGTCGCTGATGAGAAGGAAGTCAGAAGAGTTACGCTCAG*GTAAAGCAACTGTGTTTTAATCAAT*
*TTCTTGTCAGGATATATGGATTATAACTTAATTTTTGAGAAATCTGTAGTATTTGGCGTGAAATGAGTTTGCTTTTT*
*GGTTTCTCCCGTGTTATAG*GTGCCCGGCTGATGTTACGTATTTCCTTAACAAGAAGTATGACGGGGCGGTGATGTGT
ACCAGCGCGGTAGAGAGATCCGTGAAGGCAGAAGTGGTGAGAGGAAAGGGTGCATTGAACCCAATAACCTTACCGTT
GGAGGGTAAAATTTTGACCTTCACACAAGCTGACAAGTTCGAGTTACTGGAGAAGGGTTACAAG*GTAAAGTTTCCAA*
*CTTTCCTTTACCATATCAAACTAAAGTTCGAAACTTTTTATTTGATCAACTTCAAGGCCACCCGATCTTTCTATTCC*
*TGATTAATTTGTGATGAATCCATATTGACTTTTGATGGTTACGCAG*GATGTGAACACTGTGCACGAGGTGCAAGGGG
AGACATACGAGAAAACTGCTATTGTGCGCTTGACATCAACTCCGTTAGAGATCATATCGAGTGCGTCACCTCATGTT
TTGGTGGCGCTGACAAGACACACAACGTGTTGTAAATATTACACCGTTGTGTTGGACCCGATGGTGAATGTGATTTC
AGAAATGGAGAAGTTGTCCAATTTCCTTCTTGACATGTATAGAGTTGAAGCAG*GTCTGTCTTTCCTATTTCATATGT*
*TTAATCCTAGGAATTTGATCAATTGATTGTATGTATGTCGATCCCAAGACTTTCTTGTTCACTTATATCTTAACTCT*
*CTCTTTGCTGTTTCTTGCAG*GTGTCCAATAGCAATTACAAATCGATGCAGTATTCAGGGGACAGAACTTGTTTGTTC
AGACGCCCAAGTCAGGAGATTGGCGAGATATGCAATTTTACTATGACGCACTTCTTCCCGGAAACAGTACTATTCTC
AATGAATTTGATGCTGTTACGATGAATTTGAGGGATATTTCCTTAAACGTCAAAGATTGCAGAATCGACTTCTCCAA
ATCCGTGCAACTTCCTAAAGAACAACCTATTTTCCTCAAGCCTAAAATAAGAACTGCGGCAGAAATGCCGAGAACTG
CAG*GTAAAATATTGGATGCCAGACGATATTCTTTCTTTTGATTTGTAACTTTTTCCTGTCAAGGTCGATAAATTTTA*
*TTTTTTTTGGTAAAAGGTCGATAATTTTTTTTTGGAGCCATTATGTAATTTTCCTAATTAACTGAACCAAAATTATA*
*CAAACCAG*GTTTGCTGGAAAATTTGGTTGCAATGATCAAAAGAAACATGAATGCGCCGGATTTGACAGGGACAATTG
ACATTGAGGATACTGCATCTCTGGTGGTTGAAAAGTTTTGGGATTCGTATGTTGACAAGGAATTTAGTGGAACGAAC
GAAATGACCATGACAAGGGAGAGCTTCTCCAG*GTAAGGACTTCTCATGAATATTAGTGGCAGATTAGTGTTGTTAAA*
*GTCTTTGGTTAGATAATCGATGCCTCCTAATTGTCCATGTTTTACTGGTTTTCTACAATTAAAG*GTGGCTTTCGAAA
CAAGAGTCATCTACAGTTGGTCAGTTAGCGGACTTTAACTTTGTGGATTTGCCGGCAGTAGATGAGTACAAGCATAT
GATCAAGAGTCAACCAAAGCAAAAGTTAGACTTGAGTATTCAAGACGAATATCCTGCATTGCAGACGATAGTCTACC
ATTCGAAAAAGATCAATGCGATTTTCGGTCCAATGTTTTCAGAACTTACGAGGATGTTACTCGAAAGGATTGACTCT
TCGAAGTTTCTGTTCTACACCAGAAAGACACCTGCACAAATAGAGGACTTCTTTTCTGACCTAGACTCAACCCAGGC
GATGGAAATTCTGGAACTCGACATTTCGAAGTACGATAAGTCACAAAACGAGTTCCATTGTGCTGTAGAGTACAAGA
TCTGGGAAAAGTTAGGAATTGATGAGTGGCTAGCTGAGGTCTGGAAACAAG*GTGAGTTCCTAAGTTCCATTTTTTTG*
*TAATCCTTCAATGTTATTTTAACTTTTCAGATCAACATCAAAATTAGGTTCAATTTTCATCAACCAAATAATATTTT*
*TCATGTATATATAG*GTCACAGAAAAACGACCTTGAAAGATTATACGGCCGGAATCAAAACATGTCTTTGGTATCAAA
GGAAAAGTGGTGATGTGACAACCTTTATTGGTAATACCATCATCATTGCCGCATGTTTGAGCTCAATGATCCCCATG
GACAAAGTGATAAAGGCAGCTTTTTGTGGCGACGATAGCCTGATTTACTGCTTGGTAATTTAGTCGCGCGATTGTAC
CACTGGAAAGCTTCATCAAGCTCTTGTTGCTCAAGG**AGG**AGATAGAATTGGCGCT

**pAGT7392 (Donor: 5´HA – 100bp; 3´HA – 100bp)**
GG-overhang_sgRNA target D1 (**PAM**)_(5´HA(100 bp)); RdRP(*intron*)- MP; (3´HA(100bp))_ sgRNA target D1 (**PAM**)_GG-overhang

GGAGGGAAAGCTTCATCAAGCTCTTGTTGCTCAAGG**AGG**AGATAGAATTGGCTTGGTAATTTACGTTCTGTACGCGA
CGACTGAATGGTGGTTTTGTGAAAGTAGCAGAACAAGCCTTAGAGATCAAG*GTTAGTATCATATGAAGAAATACCTA*
*GTTTCAGTTGATGAATGCTATTTTCTGACCTCAGTTGTTCTCTTTTGAGAATTATTTCTTTTCTAATTTGCCTGATT*
*TTTCTATTAATTCATTAG*GTTCCCGAGCTATACTGTACCTTCGCCGACCGATTGGTACTACAGTACAAGAAGGCGGA
GGAGTTCCAATCGTGTGATCTTTCCAAACCTCTAGAAGAGTCAGAGAAGTACTACAACGCATTATCCGAGCTATCAG
TGCTTGAGAATCTCGACTCTTTTGACTTAGAGGCGTTTAAGACTTTATGTCAGCAGAAGAATGTGGACCCGGATATG
GCAGCAAAG*GTAAATCCTGGTCCACACTTTTACGATAAAAACACAAGATTTTAAACTATGAACTGATCAATAATCAT*
*TCCTAAAAGACCACACTTTTGTTTTGTTTCTAAAGTAATTTTTACTGTTATAACAG*GTGGTCGTAGCAATCATGAAG
TCAGAATTGACGTTGCCTTTCAAGAAACCTACAGAAGAGGAAATCTCGGAGTCGCTAAAACCAGGAGAGGGGTCGTG
TGCAGAGCATAAGGAAGTGTTGAGCTTACAAAATGATGCTCCGTTCCCGTGTGTGAAAAATCTAGTTGAAGGTTCCG
TGCCGGCGTATGGAATGTGTCCTAAGGGTGGTGGTTTCGACAAATTGGATGTGGACATTGCTGATTTCCATCTCAAG
AGTGTAGATGCAGTTAAAAAGGGAACTATGATGTCTGCGGTGTACACAGGGTCTATCAAAGTTCAACAAATGAAGAA
CTACATAGATTACTTAAGTGCGTCGCTGGCAGCTACAGTCTCAAACCTCTGCAAG*GTAAGAGGTCAAAAGGTTTCCG*
*CAATGATCCCTCTTTTTTTTGTTTCTCTAGTTTCAAGAATTTGGGTATATGACTAACTTCTGAGTGTTCCTTGATGCA*
*TATTTGTGATGAGACAAATGTTTGTTCTATGTTTTAG*GTGCTTAGAGATGTTCACGGCGTTGACCCAGAGTCACAGG
AGAAATCTGGAGTGTGGGATGTTAGGAGAGGACGTTGGTTACTTAAACCTAATGCGAAAAGTCACGCGTGGGGTGTG
GCAGAAGATGCCAACCACAAGTTGGTTATTGTGTTACTCAACTGGGATGACGGAAAGCCGGTTTGTGATGAGACATG
GTTCAGGGTGGCGGTGTCAAGCGATTCCTTGATATATTCGGATATGGGAAAACTTAAGACGCTCACGTCTTGCAGTC
CAAATGGTGAGCCACCGGAGCCTAACGCCAAAGTAATTTTGGTCGATGGTGTTCCCGGTTGTGGAAAAACGAAGGAG

ATTATCGAAAAG*GTAAGTTCTGCATTTGGTTATGCTCCTTGCATTTTAGGTGTTCGTCGCACTTCCATTTCCATGAA*
*TAGCTAAGATTTTTTTTCTCTGCATTCATTCTTCTTGCCTCAGTTCTAACTGTTTGTGGTATTTTTGTTTTAATTAT*
*TGCTACAG*GTAAACTTCTCTGAGGACTTGATCTTAGTCCCTGGGAAGGAAGCTTCTAAGATGATCATCCGGAGGGCC
AACCAAGCTGGTGTGATAAGAGCGGATAAGGACAATGTTAGAACGGTGGATTCCTTCTTGATGCATCCTTCTAGAAG
GGTGTTTAAGAGGTTGTTTATCGATGAAGGACTAATGCTGCATACAGGTTGTGTAAATTTCCTACTGCTGCTATCTC
AATGTGACGTCGCATATGTGTATGGGGACACAAAGCAAATTCCGTTCATTTGCAGAGTCGCGAACTTTCCGTATCCA
GCGCATTTTGCAAAACTCGTCGCTGATGAGAAGGAAGTCAGAAGAGTTACGCTCAG*GTAAAGCAACTGTGTTTTAAT*
*CAATTTCTTGTCAGGATATATGGATTATAACTTAATTTTTGAGAAATCTGTAGTATTTGGCGTGAAATGAGTTTGCT*
*TTTTGGTTTCTCCCGTGTTATAG*GTGCCCGGCTGATGTTACGTATTTCCTTAACAAGAAGTATGACGGGGCGGTGAT
GTGTACCAGCGCGGTAGAGAGATCCGTGAAGGCAGAAGTGGTGAGAGGAAAGGGTGCATTGAACCCAATAACCTTAC
CGTTGGAGGGTAAAATTTTGACCTTCACACAAGCTGACAAGTTCGAGTTACTGGAGAAGGGTTACAAG*GTAAAGTTT*
*CCAACTTTCCTTTACCATATCAAACTAAAGTTCGAAACTTTTTATTTGATCAACTTCAAGGCCACCCGATCTTTCTA*
*TTCCTGATTAATTTGTGATGAATCCATATTGACTTTTGATGGTTACGCAG*GATGTGAACACTGTGCACGAGGTGCAA
GGGGAGACATACGAGAAAACTGCTATTGTGCGCTTGACATCAACTCCGTTAGAGATCATATCGAGTGCGTCACCTCA
TGTTTTGGTGGCGCTGACAAGACACACAACGTGTTGTAAATATTACACCGTTGTGTTGGACCCGATGGTGAATGTGA
TTTCAGAAATGGAGAAGTTGTCCAATTTCCTTCTTGACATGTATAGAGTTGAAGCAG*GTCTGTCTTTCCTATTTCAT*
*ATGTTTAATCCTAGGAATTTGATCAATTGATTGTATGTATGTCGATCCCAAGACTTTCTTGTTCACTTATATCTTAA*
*CTCTCTCTTTGCTGTTTCTTGCAG*GTGTCCAATAGCAATTACAAATCGATGCAGTATTCAGGGGACAGAACTTGTTT
GTTCAGACGCCCAAGTCAGGAGATTGGCGAGATATGCAATTTTACTATGACGCACTTCTTCCCGGAAACAGTACTAT
TCTCAATGAATTTGATGCTGTTACGATGAATTTGAGGGATATTTCCTTAAACGTCAAAGATTGCAGAATCGACTTCT
CCAAATCCGTGCAACTTCCTAAAGAACAACCTATTTTCCTCAAGCCTAAAATAAGAACTGCGGCAGAAATGCCGAGA
ACTGCAG*GTAAAATATTGGATGCCAGACGATATTCTTTCTTTTGATTTGTAACTTTTTCCTGTCAAGGTCGATAAAT*
*TTTATTTTTTTTGGTAAAAGGTCGATAATTTTTTTTTGGAGCCATTATGTAATTTTCCTAATTAACTGAACCAAAAT*
*TATACAAACCAG*GTTTGCTGGAAAATTTGGTTGCAATGATCAAAAGAAACATGAATGCGCCGGATTTGACAGGGACA
ATTGACATTGAGGATACTGCATCTCTGGTGGTTGAAAAGTTTTGGGATTCGTATGTTGACAAGGAATTTAGTGGAAC
GAACGAAATGACCATGACAAGGGAGAGCTTCTCCAG*GTAAGGACTTCTCATGAATATTAGTGGCAGATTAGTGTTGT*
*TAAAGTCTTTGGTTAGATAATCGATGCCTCCTAATTGTCCATGTTTTACTGGTTTTCTACAATTAAAG*GTGGCTTTC
GAAACAAGAGTCATCTACAGTTGGTCAGTTAGCGGACTTTAACTTTGTGGATTTGCCGGCAGTAGATGAGTACAAGC
ATATGATCAAGAGTCAACCAAAGCAAAAGTTAGACTTGAGTATTCAAGACGAATATCCTGCATTGCAGACGATAGTC
TACCATTCGAAAAAGATCAATGCGATTTTCGGTCCAATGTTTTCAGAACTTACGAGGATGTTACTCGAAAGGATTGA
CTCTTCGAAGTTTCTGTTCTACACCAGAAAGACACCTGCACAAATAGAGGACTTCTTTTCTGACCTAGACTCAACCC
AGGCGATGGAAATTCTGGAACTCGACATTTCGAAGTACGATAAGTCACAAAACGAGTTCCATTGTGCTGTAGAGTAC
AAGATCTGGGAAAAGTTAGGAATTGATGAGTGGCTAGCTGAGGTCTGGAAACAAG*GTGAGTTCCTAAGTTCCATTTT*
*TTTGTAATCCTTCAATGTTATTTTAACTTTTCAGATCAACATCAAAATTAGGTTCAATTTTCATCAACCAAATAATA*
*TTTTTCATGTATATATAG*GTCACAGAAAAACGACCTTGAAAGATTATACGGCCGGATGCTTGGTAATTTAGTCGCGC
GATTGTACCACTGGAAAGCTTCATCAAGCTCTTGTTGCTCAAGG**AGG**AGATAGAATTGGCGCT

**NbPGK-locus (*Nicotiana benthamiana*; Niben101Scf05688g08010.1)**
5´UTR; Exon; *intron*; Stop codon(exchanged for GUSi); 3´UTR (homology arms underlined); **sgRNA-targets (PAM)**

CATAAGTGAGGGACTAGCTCTTGTAGGAATATTAGAATTATGGAGAATTTCCTTGGGAAAGCTCTAGATATTTATGG
ATTTGGTAGGAATACTCTTTTGAAAAGCCTTGGAATGTTCTGGTCATGTAGAAAATTCTAGAGGGTGTGCTTATATG
TAAACATGAAAAGACTTTGTGGAATAATTATTCTTAACATACTGACCCCTTGGTGATTAGTATAAATATGGGGCATT
CATTTGTAACTCATCAAGCAAAACAATCAAGTTTTCTACAATTAAAAGCTTCATTTTTCAAATTTCTCTTGTCTTTC
TTGTCTGACATTACGTTAGCGATATTGAGTATAATTATTTAGGCTGACTTAGCATAGTAAAATCATAAACAAATTGT
GCAAAATTGTAAGTGAGTTGTGAAGTGCCCATTTATACTTAATTAAAAAGTATGCACGGGACAATTGGCTTGGAAGA
CGATATGTGATTTTTCTCGTCTAAGTTGGTTGAGGAGGCCTGCTCATTGTTGATTGTCAATCATCCTAGCTAATTCT
CAATCATTTCTTTAACTTTATTGCTTTCATCGACTTGTATGTTCTAATAAATGAAGTTACATTTTTTAAAAAGTTAC
GTTAGTCCTTATTCAACCGTCGATACTCATTAATTAAACCTATTAACTATTAATTTCGCTTGATATTTTTCTATTGG
ATGATTTCCTTTTGTTTTGTCATTTATCTTCGCCAAAAACAATTACTACTTGTATGAGGAAAAAAAAGAAGAGATGA
CTTAATGTGCAACTAATGCCTTAATACGTCCTACGTAATACGGATTTTTTCATTTTCTATTATTTTCAGTTCTTTGG
AACTCGTAACCTATTTTCAAAAGAGAGTTTGAACTTGAGATTAACACAATGACTGTTGATCATTGATTCTACATAGT
TGAATTTAAAAGAGGTAAAATGTTACTACTTGAGATTTTGTGGTGAGAAGTAGAAAGAAAATGTAAGGCCAATGAGT
GACGATATGTTTGGGCCAAACAAATAAAAAAGAGTAGCCTGGAAATCATCGTTGAACCTCAACCTCTACTCAACGAT
AAGAAAAATCCACTTAACGCCCATTTCCAACCTCATTCCTCCACGTCCATATTACACCCTTTTATTTCCTTGTCAAC
ATACATCTGACATCTCTTCACTCTGTCTTCACACTCTCTGCTTTTGCCGTAGGACAGAATCGTCGAGTTTAGTGACG

AAGACTGTAATCAATGGCATCAGCTACAGCTTCTCACTCTTTGTGCGGCATCCCCGCCACCTCATCCTCTACTACCA
ACAAGTCTATTGCCCCTTCATCTGCTCGCTTCCTCGCCAAAACTCCTCCCCGCGGCCTCGGCTTCGCTGGCGCCGCC
GCTGATTCTCTCTTCACCAACCACGTGGCAACCAAGCTCCGATCCCTCAAGAGCTCCTCCAAGCCTGTTAGGGGCGT
TGCTTCTATGGCCAAGAAAAGCGTTGGTGACCTCACCGCTGCCGAGTTGAAGGGCAAGAATGTCTTCGTCAGGGCCG
ATTTGAATGTCCCACTTGATGATAACCAGAACATTACTGATGACACTAGAATTAGAGCTGCCGTCCCTACTATCAAG
CACTTGATGGCCAATGGTGCTAAAGTTATTCTCTCCACTCACCTGGTCTGTACCACTTTCTTCTCTTCATCTGTCCT
TTTTTCTTTAATTTTAATTGCATACCTACGATCACCCCCCCCCCCCCCCCTTTTCTTTTTGCCCCTTTCCCAGTGGA
TCTTTGTTGTGCCTTTTCTTTTTATTTTTTAATCTCAGTAACAGAGTGGAACTGAAATTGAACTTATTTACTGGCTG
ACAATATCATAGGAAATTGCGTGTCTATGGATGATTAGTCTATTTAGAATTATTAAATTTTGATAGCTTTTTACAAG
TCACTCTAATGTGCCTTCTATGTTGTCCTCCTATTTGCTTTCTCTTCTTCCGCATTTTGTTAAGGTGTTGAGGATAT
GTCAAATCTTAGTTTATTTTGCTTATTGAAATTTCAGGGACGGCCAAAAGGAGTCACTCCTAAATACAGCTTGGCAC
CCCTAGTCCCCAGGCTATCCGAACTGCTTGGAATCCAGGTACATCCTGTTGCAATTTTAGGTCTATTCATTGTTTAA
AAAGAAGATAGAAAGAAAGAATTAAAGCATAAGAAATAAGCAGTTCATGTCGATCAAAAGGTGACTGTCTACCTTAG
TTTGTATTGACTCTTGTATGCTCTACCAATAGGTTGTGAAGGCTGAGGACTGCATTGGTCCGGAAGTTGAGAAGTTG
GTTGCTTCACTTCCCGAGGGTGGTGTTCTTCTTCTCGAGAACGTGAGATTCTACAAGGAGGAAGAGAAGAATGAACC
TGAGTTTGCAAAGAAACTTGCATCATTGGCAGATCTTTACGTGAATGATGCATTCGGTACAGCTCACAGAGCACACG
CCTCTACAGAGGGAGTTACTAAATTTTTGAAGCCTTCTGTTGCAGGTTTCCTCTTACAAAAGGTTAGCTTACTTGAT
TGTCACTTCCTTTACTGAGGCAGAGCTAGAATTTCAACTTTATGGATTCTGAATTTTAGAACAATGACTTCAAGTGC
TAATAACTGGGTTCTAAATTTAATATTTGTACATATTTAATAATTTCTTTTAGCATGGTTTGAGCAAAAGCTACTGG
GTTTGGCCGAATCCATACTTGGGCTTCTAGCTGTGCTCCTGCTTCCTTACATATTGTTTTCTTTGACAAACTGATGG
GCGTAAAACCATCTATGGAAAATTGCACTCTTTAGAAATTTTATATGTCTTAATTGTTCTTTTATTTGTTTCAGGAA
TTGGACTACTTAGTCGGGGCAGTTTCAAATCCAAAGAGGCCATTTGCTGCTATTGTGGGTGGTTCAAAGGTTTCATC
CAAGATTGGAGTGATCGAATCACTTTTAGAGAAATGTGATATATTGCTTTTGGGTGGAGGAATGATCTTTACCTTCT
ACAAGGCTCAGGGTCTTTCAGTTGGTTCCTCCTTGGTTGAGGAAGACAAACTAGAACTCGCTACATCACTCCTAGAG
AAGGCCAAGGCGAAAGGAGTCAGTCTCTTGTTACCATCTGATGTTGTGATTGCAGATAAATTTGCTCCTGATGCAAA
CAGCAAGGTTTGCATGCTAAGTTTTCTCATATAAACCTATCTGACCTTAGAGCTTTTTGCTCTTGAGATTCTTTAGA
CTTTCCATCTGAAATCTGTACTGTAATTGGCTCTTAATATCAGAGTTTGTTACTTATGGATTGTGTTGAAAATGCAA
TTTTGTTTTGGTTACTGCAGATTGTGCCGGCATCTGCTATCCCAGATGGTTGGATGGGGTTGGACATTGGACCAGAC
TCTGTTAAGACTTTCAACGATGCCTTGGATACCACAAAAACAGTGATCTGGAATGGACCTATGGGGGTGTTTGAATT
TGACAAGTTTGCTGTTGGAACAGAGGTACCAATTACCATTCTTCTCTTCATATTTGTTTTACCTTACCGAATGCTGA
GCTTTATAAAAGAAATAAAAAAGGGAATAAAGCTGGTTTTACATAGCTTTAAAAGTAAAGGAAGAGGAATAATCTGG
TTGGATATGTCACTTTGTGTGTTTACCTGAGAGTAAATAGTAATAAGAATGTTGTTGTGGTGATAGGCAATTGCAAA
GAAGCTCGCGGACTTAAGTGGGAAAGGAGTGACAACTATCATTGGAGGTGGAGATTCTGTTGCAGCTGTTGAGAAAG
TTGGAGTTGCTAGCGTGATG**AGCCACATATCCACTGGTGG**TGGTGCCAGTTTGGAGCTACTGGAAGGCAAGGTGCTC
CCTGGTGTCGTTGCTCTAGATGAAGCAGATGCCCCTGTTGCTGTGTAAAACAATTTGTACTAATTCTTTTTTCT**CCG**
**GTGATCAGCATAATCAGTGG**TAATTTCCAGTTGGGAAGCATTGAGTTGATGTTGTAGATTTTTCAGGTTATATTGTT
ATATAATGTCCCCTTTCTTTAACCCATGTTATTTTGTCTAAATAAAGGGCGAGTATATCAGTTATAGACAGCTATCC
TTTTGATGTCCTTCAACAAACTCTATCCTTGATTTGGTTCAGTTTGGAGGAACTTCTTTAGACATAAGAACTTTTGC
CATGTGAACAAACTCATGCTGCGTGTTAATGTTCACCTGCCCTTCGCATTAAATGACGCCTCTAATACGTGGTGGTC
TATGTAAAGTGAGATTACTGTTAGTTATGCAGTTATGAAACTTCTGGAAAACTTGAGACACCTGATCTTTGATTTTC
AAAATATAAACCCATAATTTTCGTGAAGCATCATCAATAAAAGTAACAAAATATTTGTTACCGTACATTTATTCAAT
TTCCATTGGGCCACAAACATCAGAATATACTAAATCAAGTATATTCAATTTTCTTTCAAACGATGTCTGAAATGAGA
CTCTATGCTGCTTACCAAATAAACAGTAGTCACAGGGTTTTATTGTTGTACCTTTGGCATAAGAAATAAGTGATTTT
TTGGCAAGAATCTGCAATCCCTTGTCGCTCATATGACCCATTCTTTTTATGCCACAAATTTGCAGAAATCTCATCTT
GCATCGCATTCAATTCACCTTGGCATATTACTGCATTTGTCCTGTACAACGTGCCACGAGTAACTCCCTTTGCAATC
ACCAACGATCCCTTGGTGAGTCTCCATTTTTGATTTGCAAAATAGTTCTCGTATCCATCTCGGTCCAAAGTAATCCC
CGAGATCAAGTTCATCCACAAAACAGGTACATGCCGCACATCCTTTAGAACCAATGTGCATCCGACATTTGTCTTGA
TACAAATGTCACCAATCCCCGCAATCTTTCAGTNCCAAATTATCGTACGAAGATGGCAACGAGTTCAATAGCAAGAT
GGC

**pAGT7378 (Donor NbPGK-GUS: 5´HA – 1001bp; 3´HA – 999bp)**
GG-overhang-5´HA(1001 bp); NbPGK: Exon**;** *intron*; GUS: Exon; *intron*; 3´UTR; 3´HA(999bp)-GG-overhang; **Mutations** to prevent **sgRNA** cleavage (PAM)

GGAGTGCTATTGTGGGTGGTTCAAAGGTTTCATCCAAGATTGGAGTGATCGAATCACTTTTAGAGAAATGTGATATA
TTGCTTTTGGGTGGAGGAATGATCTTTACCTTCTACAAGGCTCAGGGTCTTTCAGTTGGTTCCTCCTTGGTTGAGGA
AGATAAACTAGAACTCGCTACATCACTCCTAGAGAAGGCCAAGGCGAAAGGAGTCAGTCTCTTGTTACCATCTGATG
TTGTGATTGCAGATAAATTTGCTCCTGATGCAAACAGCAAG*GTTTGCATGCTAAGTTTTCTCATATAAACCTATCTG*
*ACCTTAGAGCTTTTTGCTCTTGAGATTCTTTAGACTTTCCATCTGAAATCTGTACTGTAATTGGCTCTTAATATCAG*

*AGTTTGTTACTTATGGATTGTGTTGAAAATGCAATTTTGTTTTGGTTACTGCAG*ATTGTGCCGGCATCTGCTATCCC
AGATGGTTGGATGGGGTTGGACATTGGACCAGACTCTGTTAAGACTTTCAACGATGCCTTGGATACCACAAAAACAG
TGATCTGGAATGGACCTATGGGGGTGTTTGAATTTGACAAGTTTGCTGTTGGAACAGAG*GTACCAATTACCATTCTT*
*CTCTTCATATTTGTTTTACCTTACCGAATGCTGAGCTTTATAAAAGAAATAAAAAAGGGAATAAAGCTGGTTTTACA*
*TAGCTTTAAAAGTAAAGGAAGAGGAATAATCTGGTTGGATATGTCACTTTGTGTGTTTACCTGAGAGTAAATAGTAA*
*TAAGAATGTTGTTGTGGTGATAG*GCAATTGCAAAGAAGCTCGCGGACTTAAGTGGGAAAGGAGTGACAACTATCATT
GGAGGTGGAGATTCTGTTGCAGCTGTTGAGAAAGTTGGAGTTGCTAGCGTGATG**AGCCACATATCCACTGGAGGTGC**
TGCCAGTTTGGAGCTACTGGAAGGCAAGGTGCTCCCTGGTGTCGTTGCTCTAGATGAAGCAGATGCCCCTGTTGCTG
TGTCAGGTCAGTCCCTTATGTTACGTCCTGTAGAAACCCCAACCCGTGAAATCAAAAACTCGACGGCCTGTGGGCA
TTCAGTCTGGATCGCGAAAACTGTGGAATTGATCAGCGTTGGTGGGAAAGCGCGTTACAAGAAAGCCGGGCAATTGC
TGTGCCAGGCAGTTTTAACGATCAGTTCGCCGATGCAGATATTCGTAATTATGCGGGCAACGTCTGGTATCAGCGCG
AAGTCTTTATACCGAAAG*GTAAGTCTTACTCTCTCTTTTTTGGTCTGTATTTTTAATTTTTTGAAGTATACTATTTG*
*TACTGACGCTAATAATCTTTTTTCAG*GTTGGGCAGGCCAGCGTATCGTGCTGCGTTTCGATGCGGTCACTCATTACG
GCAAAGTGTGGGTCAATAATCAGGAAGTGATGGAGCATCAGGGCGGCTATACGCCATTTGAAGCCGATGTCACGCCG
TATGTTATTGCCGGGAAAAGTGTACGTATCACCGTTTGTGTGAACAACGAACTGAACTGGCAGACTATCCCGCCGGG
AATGGTGATTACCGACGAAAACGGCAAGAAAAAGCAGTCTTACTTCCATGATTTCTTTAACTATGCCGGAATCCATC
GCAGCGTAATGCTCTACACCACGCCGAACACCTGGGTGGACGATATCACCGTGGTGACGCATGTCGCGCAAGACTGT
AACCACGCGTCTGTTGACTGGCAG*GTACTTCATGCTTCAACGTGTAACTTAAGAGATACTGTGTGAAATTTTATATT*
*TCCATACATTTGCTTGACCTTTGCTTTTTGTCAATTTTTTTCCCCTTACAG*GTGGTGGCCAATGGTGATGTCAGCGT
TGAACTGCGTGATGCGGATCAACAGGTGGTTGCAACTGGACAAGGCACTAGCGGGACTTTGCAAGTGGTGAATCCGC
ACCTCTGGCAACCGGGTGAAGGTTATCTCTATGAACTGTGCGTCACAGCCAAAAGCCAGACAGAGTGTGATATCTAC
CCGCTTCGCGTCGGCATCCGGTCAGTGGCAGTGAAGGGCGAACAGTTCCTGATTAACCACAAACCGTTCTACTTTAC
TGGCTTTGGTCGTCATGAAGATGCGGACTTGCGTGGCAAAGGATTCGATAACGTGCTGATGGTGCACGACCACGCAT
TAATGGACTGGATTGGGGCCAACTCCTACCGTACCTCGCATTACCCTTACGCTGAAGAGATGCTCGACTGGGCAGAT
GAACATGGCATCGTGGTGATTGATGAAACTGCTGCTGTCGGCTTTAACCTCTCTTTAGGCATTGGTTTCGAAGCGGG
CAACAAGCCGAAAGAACTGTACAGCGAAGAGGCAGTCAACGGGGAAACTCAGCAAGCGCACTTACAGGCGATTAAAG
AGCTGATAGCGCGTGACAAAAACCACCCAAGCGTGGTGATGTGGAGTATTGCCAACGAACCGGATACCCGTCCGCAA
GGTGCACGGGAATATTTCGCGCCACTGGCGGAAGCAACGCGTAAACTCGACCCGACGCGTCCGATCACCTGCGTCAA
TGTAATGTTCTGCGACGCTCACACCGATACCATCAGCGATCTCTTTGATGTGCTGTGCCTGAACCGTTATTACGGAT
GGTATGTCCAAAGCGGCGATTTGGAAACGGCAGAGAAGGTACTGGAAAAAGAACTTCTGGCCTGGCAGGAGAAACTG
CATCAGCCGATTATCATCACCGAATACGGCGTGGATACGTTAGCCGGGCTGCACTCAATGTACACCGACATGTGGAG
TGAAGAGTATCAGTGTGCATGGCTGGATATGTATCACCGCGTCTTTGATCGCGTCAGCGCCGTCGTCGGTGAACAGG
TATGGAATTTCGCCGATTTTGCGACCTCGCAAGGCATATTGCGCGTTGGCGGTAACAAGAAAGGGATCTTCACTCGC
GACCGCAAACCGAAGTCGGCGGCTTTTCTGCTGCAAAAACGCTGGACTGGCATGAACTTCGGTGAAAAACCGCAGCA
GGGAGGCAAACAATGAGCTTAACAATTTGTACTAATTCTTTTTTCT**GCGGTCATCAGCATAATCAGTGG**TAATTTCC
AGTTGGGAAGCATTGAGTTGATGTTGTAGATTTTTCAGGTTATATTGTTATATAATGTCCCCTTTCTTTAACCCATG
TTATTTTGTCTAAATAAAGGGCGAGTATATCAGTTATAGACAGCTATCCTTTTGATGTCCTTCAACAAACTCTATCC
TTGATTTGGTTCAGTTTGGAGGAACTTCTTTAGACATAAGAACTTTTGCCATGTGAACAAACTCATGCTGCGTGTTA
ATGTTCACCTGCCCTTCGCATTAAATGACGCCTCTAATACGTGGTGGTCTATGTAAAGTGAGATTACTGTTAGTTAT
GCAGTTATGAAACTTCTGGAAAACTTGAGACACCTGATCTTTGATTTTCAAAATATAAACCCATAATTTTCGTGAAG
CATCATCAATAAAAGTAACAAAATATTTGTTACCGTACATTTATTCAATTTCCATTGGGCCACAAACATCAGAATAT
ACTAAATCAAGTATATTCAATTTTCTTTCAAACGATGTCTGAAATGAGACTCAATGCTGCTTACCAAATAAACAGTA
GTCACAGGGTTTTATTGTTGTACCTTTGGCATAAGAAATAAGTGATTTTTTGGCAAGAATCTGCAATCCCTTGTCGC
TCATATGACCCATTCTTTTTATGCCACAAATTTGCAGAAATCTCATCTTGCATCGCATTCAATTCACCTTGGCATAT
TACTGCATTTGTCCTGTACAACGTGCCACGAGTAACTCCCTTTGCAATCACCAACGATCCCTTGGTGAGTCTCCATT
TTTGATTTGCAAAATAGTTCTCGTATCCATCTCGGTCCAAAGTAATCCCCGAGATCAAGTTCATCCACAAAACAGGT
ACATGCCGCACATCCTTTAGAACCAATGTGCATCCGACATTTGTCTTGATACAAATGTCACCAATCCCCCTGCAATC
TTTGAGTAACTCGTGTTACGCT

**pAGT7879 (D1-Donor NbPGK-GUS-D1: 5´HA – 1001bp; 3´HA – 999bp)**

GG-overhang-sgRNA-target D1 (**PAM**); 5´HA(1001 bp); NbPGK: Exon**;** *intron*; GUS: Exon; *intron*; 3´UTR; 3´HA(999bp); **Mutations** to prevent **sgRNA** cleavage (PAM); sgRNA-target D1 (**PAM**)-GG-overhang

GGAGCCAATTCTATCT**CCT**CCTTGAGCAACAAGAGCTTGATGAAGCTTTCCTTGCTATTGTGGGTGGTTCAAAGGTT
TCATCCAAGATTGGAGTGATCGAATCACTTTTAGAGAAATGTGATATATTGCTTTTGGGTGGAGGAATGATCTTTAC
CTTCTACAAGGCTCAGGGTCTTTCAGTTGGTTCCTCCTTGGTTGAGGAAGATAAACTAGAACTCGCTACATCACTCC
TAGAGAAGGCCAAGGCGAAAGGAGTCAGTCTCTTGTTACCATCTGATGTTGTGATTGCAGATAAATTTGCTCCTGAT
GCAAACAGCAAG*GTTTGCATGCTAAGTTTCTCATATAAACCTATCTGACCTTAGAGCTTTTTGCTCTTGAGATTCT*
*TTAGACTTTCCATCTGAAATCTGTACTGTAATTGGCTCTTAATATCAGAGTTTGTTACTTATGGATTGTGTTGAAAA*

*TGCAATTTTGTTTTGGTTACTGCAG*ATTGTGCCGGCATCTGCTATCCCAGATGGTTGGATGGGGTTGGACATTGGAC
CAGACTCTGTTAAGACTTTCAACGATGCCTTGGATACCACAAAAACAGTGATCTGGAATGGACCTATGGGGGTGTTT
GAATTTGACAAGTTTGCTGTTGGAACAGAG*GTACCAATTACCATTCTTCTCTTCATATTTGTTTTACCTTACCGAAT*
*GCTGAGCTTTATAAAAGAAATAAAAAAGGGAATAAAGCTGGTTTTACATAGCTTTAAAAGTAAAGGAAGAGGAATAA*
*TCTGGTTGGATATGTCACTTTGTGTGTTTACCTGAGAGTAAATAGTAATAAGAATGTTGTTGTGGTGATAG*GCAATT
GCAAAGAAGCTCGCGGACTTAAGTGGGAAAGGAGTGACAACTATCATTGGAGGTGGAGATTCTGTTGCAGCTGTTGA
GAAAGTTGGAGTTGCTAGCGTGATG**AGCCACATATCCACTGGAGGTGC**TGCCAGTTTGGAGCTACTGGAAGGCAAGG
TGCTCCCTGGTGTCGTTGCTCTAGATGAAGCAGATGCCCCTGTTGCTGTGTCAGGTCAGTCCCTTATGTTACGTCCT
GTAGAAACCCCAACCCGTGAAATCAAAAAACTCGACGGCCTGTGGGCATTCAGTCTGGATCGCGAAAACTGTGGAAT
TGATCAGCGTTGGTGGGAAAGCGCGTTACAAGAAAGCCGGGCAATTGCTGTGCCAGGCAGTTTTAACGATCAGTTCG
CCGATGCAGATATTCGTAATTATGCGGGCAACGTCTGGTATCAGCGCGAAGTCTTTATACCGAAAG*GTAAGTCTTAC*
*TCTCTCTTTTTTGGTCTGTATTTTTAATTTTTTGAAGTATACTATTTGTACTGACGCTAATAATCTTTTTTCAG*GTT
GGGCAGGCCAGCGTATCGTGCTGCGTTTCGATGCGGTCACTCATTACGGCAAAGTGTGGGTCAATAATCAGGAAGTG
ATGGAGCATCAGGGCGGCTATACGCCATTTGAAGCCGATGTCACGCCGTATGTTATTGCCGGGAAAAGTGTACGTAT
CACCGTTTGTGTGAACAACGAACTGAACTGGCAGACTATCCCGCCGGGAATGGTGATTACCGACGAAAACGGCAAGA
AAAAGCAGTCTTACTTCCATGATTTCTTTAACTATGCCGGAATCCATCGCAGCGTAATGCTCTACACCACGCCGAAC
ACCTGGGTGGACGATATCACCGTGGTGACGCATGTCGCGCAAGACTGTAACCACGCGTCTGTTGACTGGCAG*GTACT*
*TCATGCTTCAACGTGTAACTTAAGAGATACTGTGTGAAATTTTATATTTCCATACATTTGCTTGACCTTTGCTTTTT*
*GTCAATTTTTTTCCCCTTACAG*GTGGTGGCCAATGGTGATGTCAGCGTTGAACTGCGTGATGCGGATCAACAGGTGG
TTGCAACTGGACAAGGCACTAGCGGACTTTGCAAGTGGTGAATCCGCACCTCTGGCAACCGGGTGAAGGTTATCTC
TATGAACTGTGCGTCACAGCCAAAAGCCAGACAGAGTGTGATATCTACCCGCTTCGCGTCGGCATCCGGTCAGTGGC
AGTGAAGGGCGAACAGTTCCTGATTAACCACAAACCGTTCTACTTTACTGGCTTTGGTCGTCATGAAGATGCGGACT
TGCGTGGCAAAGGATTCGATAACGTGCTGATGGTGCACGACCACGCATTAATGGACTGGATTGGGGCCAACTCCTAC
CGTACCTCGCATTACCCTTACGCTGAAGAGATGCTCGACTGGGCAGATGAACATGGCATCGTGGTGATTGATGAAAC
TGCTGCTGTCGGCTTTAACCTCTCTTTAGGCATTGGTTTCGAAGCGGGCAACAAGCCGAAAGAACTGTACAGCGAAG
AGGCAGTCAACGGGGAAACTCAGCAAGCGCACTTACAGGCGATTAAAGAGCTGATAGCGCGTGACAAAAACCACCCA
AGCGTGGTGATGTGGAGTATTGCCAACGAACCGGATACCCGTCCGCAAGGTGCACGGGAATATTTCGCGCCACTGGC
GGAAGCAACGCGTAAACTCGACCCGACGCGTCCGATCACCTGCGTCAATGTAATGTTCTGCGACGCTCACACCGATA
CCATCAGCGATCTCTTTGATGTGCTGTGCCTGAACCGTTATTACGGATGGTATGTCCAAAGCGGCGATTTGGAAACG
GCAGAGAAGGTACTGGAAAAAGAACTTCTGGCCTGGCAGGAGAAACTGCATCAGCCGATTATCATCACCGAATACGG
CGTGGATACGTTAGCCGGGCTGCACTCAATGTACACCGACATGTGGAGTGAAGAGTATCAGTGTGCATGGCTGGATA
TGTATCACCGCGTCTTTGATCGCGTCAGCGCCGTCGTCGGTGAACAGGTATGGAATTTCGCCGATTTTGCGACCTCG
CAAGGCATATTGCGCGTTGGCGGTAACAAGAAAGGATCTTCACTCGCGACCGCAAACCGAAGTCGGCGGCTTTTCT
GCTGCAAAAACGCTGGACTGGCATGAACTTCGGTGAAAAACCGCAGCAGGGAGGCAAACAATGAGCTTAACAATTTG
TACTAATTCTTTTTTCT**GCGGTCATCAGCATAATCAGTGG**TAATTTCCAGTTGGGAAGCATTGAGTTGATGTTGTAG
ATTTTTCAGGTTATATTGTTATATAATGTCCCCTTTCTTTAACCCATGTTATTTTGTCTAAATAAAGGGCGAGTATA
TCAGTTATAGACAGCTATCCTTTTGATGTCCTTCAACAAACTCTATCCTTGATTTGGTTCAGTTTGGAGGAACTTCT
TTAGACATAAGAACTTTTGCCATGTGAACAAACTCATGCTGCGTGTTAATGTTCACCTGCCCTTCGCATTAAATGAC
GCCTCTAATACGTGGTGGTCTATGTAAAGTGAGATTACTGTTAGTTATGCAGTTATGAAACTTCTGGAAAACTTGAG
ACACCTGATCTTTGATTTTCAAAATATAAACCCATAATTTTCGTGAAGCATCATCAATAAAAGTAACAAAATATTTG
TTACCGTACATTTATTCAATTTCCATTGGGCCACAAACATCAGAATATACTAAATCAAGTATATTCAATTTTCTTTC
AAACGATGTCTGAAATGAGACTCAATGCTGCTTACCAAATAAACAGTAGTCACAGGGTTTTATTGTTGTACCTTTGG
CATAAGAAATAAGTGATTTTTTGGCAAGAATCTGCAATCCCTTGTCGCTCATATGACCCATTCTTTTTATGCCACAA
ATTTGCAGAAATCTCATCTTGCATCGCATTCAATTCACCTTGGCATATTACTGCATTTGTCCTGTACAACGTGCCAC
GAGTAACTCCCTTTGCAATCACCAACGATCCCTTGGTGAGTCTCCATTTTTGATTTGCAAAATAGTTCTCGTATCCA
TCTCGGTCCAAAGTAATCCCCGAGATCAAGTTCATCCACAAAACAGGTACATGCCGCACATCCTTTAGAACCAATGT
GCATCCGACATTTGTCTTGATACAAATGTCACCAATCCCCCTGCAATCTTTGAGTAACTCGTGTTACCACTGGAAAG
CTTCATCAAGCTCTTGTTGCTCAAGG**AGG**AGATAGAATTGGCGCT

**NbTPR-locus (*Nicotiana benthamiana*; Niben101Scf03365g03011.1)**
5´UTR; Exon; *intron*; Stop codon(exchanged for GUSi); 3´UTR (homology arms underlined); **sgRNA-targets (PAM)**

ATGCTAAAATGTCAGACTACGATTCTTACCCTGTTACCTTCCTAGTCCAACCCAAGAAAACAAGTGGACA
ACTCTTCATTTTTTTGTTATTCCAACCTGTTAATGAATGGCAAAATCAGATTTGTTTTCTTTGGTAACCAA
CGGAGCATATCACCCCAAACACACATGTCAAAAAGCAACCACTCTTGACCTTTTCCAGCTAAAATAATAA
TTGAGAAAAGTCTGATATTTTTTGCTACAAACTTTCATTCTCCACCACTTCTTTTTTTTCTTCTTCTTCA

ATTCCAACACTATCGTAATATCTTCCATTTATTATGTCAAATCTTTTCTTTCTATATAAAGAGTCATTTC
TTCCCCAGCAAGATTATTGGTCCAGAAGATTTTCAAGAATACTCATTTTAG*GTAAATCTCTCTTTGTTCT*
*TCTGCTATTGATTTTTCCTTCTGGTGTTTGTCTGTTTATGCCTAAAATCTTTTCTCTTCTCTACAG*GAAC
TTTATAATCACATAAATAAAGATTGAATCTTTGTTTTACACTCAAACAAGAAAAGTAAAATGATGCTAAG
GAGTTCATCAACACCACTTCTTAGATCTTTACTCTCAGAAAGTCCAAGCAATCACCACCACCATCAGCAC
TCTGATTTAACCCCTAATCACACACCCAATTCCATTTTCCATAGCTATACCAAACTCTCATGTAACCATG
GTGGATATCAGAATTACACCAAGATTTCTAACTCTCCCATGTCTCCTTCAGTTTCTGAGCTTAGCAATGG
CAGACAATTAGCATCTCATGGTATTAGAAGAGCTCAATCTGAAGGGAACTTGGAAGGATTGACAAAAGCC
TCAACAGAAGAAGTTGATGAATTTAGCCTCTCAAAACTACCGAAGACGCTTGTACGTAAACCCCACAAAG
CATTCATGGAAACCATTCCATCTTTCAACTTTCACAATTCAAGGGAGTTCCATTCGGACGACGACAGCTA
TGATGAGGAAGATGAAGATAATGACTGTGGTTACATTAGTAAGCAGTACAGTTTGGGAAATAATGGTGTA
AAAGAGGAAATGAGTTATGTGAGTCAAAATTCAAAGTTAGAAATTGTTGAAGGCAGAGAAGAAATGTACC
TAGCAAGAGGAATTGGGATTGCTGATATTGGTTGTTTTGATGATGGTGGCCCTTATGGAGGCTGGGGCAA
CGGAGGAGGAGGAGGCGGCGGTTATCCGCCAGTAGCCTTCGACAGAGAAGGTGGTGGCGACAGTCAGGGG
CTCCACATAGAAGAGTATTACAAGAGGATGTTGGAACAGAATCCTGGCAATTCCCTATTCTTGAGGAATT
ATGCCAACTTTTTATACCAA*GTGAGTCTTTAACCTTCCTCAAGCTACTCTTATTTAGCATATTTTACAGG*
*CATCTTGTGAATCCACAATAGCTAACTTTGCTTTTGCTAATGTTGGGGCAG*ACAAAGAAAGATCTTAAGG
GGGCAGAGGAATACTACTCAAGAGCTATTTTAGCAGATCCAAGTGATGGTGAAATTCTTTCACAGTACGC
TAAACTAATATGGGAGCTTCACCGCGATGAAGATAGAGCCACGAGCTATTTTGAAAGAGCAGTTCAAGCT
GCCTCTAGTGACAG*GTTCGCTTCTCATACACTACTGAGAATAAGCTTTATTATAGAAAATACAATTCAAG*
*AACGGTGTTATTTATTTCAAGAACTGACCTCTTCATCTTTGATGTGCTGACAG*CCACATACATGCAGCTT
ATGCTAATTTCCTCTGGGAGATAGAAGATGAAGAAGATGAGAATGATGATATACAAGCACCACGAATGCT
GCATAC**CGTAGCTACAGCGTCTATAACGG**CTTGACTTGAAAGAAAATCATTATTCAAATGTTTAACAGCA
GTATTAGCTGAAGTTTG*GTGAGCAATTATGAAGAACCTGTAATTTAGCCTCTAAGGTAGTAATAAAACTG*
*ATGCACAGGACTTCTGTATTTCTT**CCGCTGTCGTCGCCGACTGACTA**ACCCTGTGAATGGTGAAACAGAA*
*GCACTTGTCAAACTCCAATTGTGCTAAGCGTTTATATTGACAAGTTCTTCGCGACTTTCATTGCTTGCAA*
*AACTTACATGTTTAATACCATATTACAATCAG*GAGAGTTGGGGGACATGGATTATGAAGAAGCAGCTTGA
CGTCACCACCATACCGTTTAGCTTTGCTCAAGACATTCTGACATGCTGCATTGAACAACAAAAATAACAG
TATTTCATCTTGGATTGTAGTACCATCATCGTAGGGAGGAATTAGCATTAACTGTGACTAGGGCATTAGT
CCAAATGAAACAGGAAATCTTCGTACACAGCCTAACTAAAGAATCACGACCAAGAGATTATATCCTGCAT
ACAAGGATAATGTTAATTGATTCAACCAAAGAACTACTCAAGCAAGCAAAAGATAGAACAAATCTTTAAG
AAAAGCTTAAAGAGAAATAAGTACCACAATGTGAAATTGAAATTAATTTAAGCTACAGACTATCTCCACG
AGAAACGGTACAGCTGATTGATCTACTTTCCTCAATGTAAACTTGCCTTCAAGCCAATGGCACGGGTAAT
GCCTAAGAACAGGGAAGAGAAAGCATCATATACTAGGTATACAACGGATGAGATAACAAAGTAATGTATG
AATATGAGATTGTGGCAATGATGACGTACACCTTTCACTTGCAACATTACTCAAGACAACTAACTGCATC
ATATTTCTTCTGAAAAAATGTTTTATCTACTGAATGCAAGTTCCGCAACCATTAAATAAAAGATCATCGA
TCTATGTGATCAGAAATATTTCAAGCACGCACCTTACTAATGCGATTTACTACTAGAAAGCTAGCTAAAA
TTAAGGTTGAGGGAACAATCTGCTAGGAATCATACAGGACAGAAGGTAAAAAAAAGTTAAGAGCAGCTTT
TACCAACAACTGGAAATGCAACTCAAATCCTCCACGAAAACACACAAACAAATCAATGAATCATGACTTC
AAATTTTCAAAGTTCATACCTTAATTGCATTTTGGAAACCCTGAGGTATTCAAACTCAACAAAGGTATAG
CAAATGCTGATATTCTATCCATCAATTGCAGTCGAAATATTAGAGCATGTAAGACAATTGAAGCATGTAA
TATTGACAACGGAAAAAAACTACTTCGCGCTTTTTCAAGAATACTCCATCCTGCTTATGCTTCCCAAAGT
TCTTAAACTCTTGCAGAAAATCTTGTCTAGTACATTGGATATGGCAAATTCCTCCGGTAGACAGATTTTG
ATTCCCCTGGAGAAAAGTATATAGAGTCAAATCTAGCTAAACTCGGTAATATAATGAACTAGACGCTAGA
ATGCATCTACTGAAAGAGTAGCGACATTGATTGCGAAAATGCATTCATCTTACCTTCTTGACATAAAATT
TCATCTGCCTGGGACATTGAGTTTGACGCAGAAGCAGGTTTCCACTCTGTAGCAGGCACAGCACTCTTTG
TATAA

**pAGT7958 (D1-Donor NbTPR-GUS-D1: 5´HA – 896bp; 3´HA – 1000bp)**
GG-overhang-sgRNA-target D1 (**PAM**); 5´HA(896 bp); NbPGK: Exon**;** *intron*; GUS: Exon; *intron*; 3´UTR;
3´HA(1000bp); **Mutations** to prevent **sgRNA** cleavage (PAM); sgRNA-target D1 (**PAM**)-GG-overhang

GGAGCCAATTCTATCT**CCT**CCTTGAGCAACAAGAGCTTGATGAAGCTTTCCTTGAGTTCCATTCGGACGA
CGACAGCTATGATGAGGAAGATGAAGATAATGACTGTGGTTACATTAGTAAGCAGTACAGTTTGGGAAAT
AATGGTGTAAAAGAGGAAATGAGTTATGTGAGTCAAAATTCAAAGTTAGAAATTGTTGAAGGCAGAGAAG
AAATGTACCTAGCAAGAGGAATTGGGATTGCTGATATTGGTTGTTTTGATGATGGTGGCCCTTATGGAGG
CTGGGGCAACGGAGGAGGAGGAGGCGGCGGTTATCCGCCAGTAGCCTTCGACAGAGAAGGTGGTGGCGAC
AGTCAGGGGCTCCACATAGAAGAGTATTACAAGAGGATGTTGGAACAGAATCCTGGCAATTCCCTATTCT
TGAGGAATTATGCCAACTTTTTATACCAAGTGAGTCTTTAACCTTCCTCAAGCTACTCTTATTTAGCATA
TTTTACAGGCATCTTGTGAATCCACAATAGCTAACTTTGCTTTTGCTAATGTTGGGGCAGACAAAGAAAG
ATCTTAAGGGGGCAGAGGAATACTACTCAAGAGCTATTTTAGCAGATCCAAGTGATGGTGAAATTCTTTC
ACAGTACGCTAAACTAATATGGGAGCTTCACCGCGATGAAGATAGAGCCACGAGCTATTTTGAAAGAGCA
GTTCAAGCTGCCTCTAGTGACAGGTTCGCTTCTCATACACTACTGAGAATAAGCTTTATTATAGAAAATA
CAATTCAAGAACGGTGTTATTTATTTCAAGAACTGACCTCTTCATCTTTGATGTGCTGACAGCCACATAC
ATGCAGCTTATGCTAATTTCCTCTGGGAGATAGAAGATGAAGAAGATGAGAATGATGATATACAAGCACC
ACGAATGCTGCATAC**CGTAGCTACAGCGTCAATTACCG**CAGGTCAGTCCCTTATGTTACGTCCTGTAGAA
ACCCCAACCCGTGAAATCAAAAAACTCGACGGCCTGTGGGCATTCAGTCTGGATCGCGAAAACTGTGGAA
TTGATCAGCGTTGGTGGGAAAGCGCGTTACAAGAAAGCCGGGCAATTGCTGTGCCAGGCAGTTTTAACGA
TCAGTTCGCCGATGCAGATATTCGTAATTATGCGGGCAACGTCTGGTATCAGCGCGAAGTCTTTATACCG
AAAGGTAAGTCTTACTCTCTCTTTTTTGGTCTGTATTTTTAATTTTTTGAAGTATACTATTTGTACTGAC
GCTAATAATCTTTTTTCAGGTTGGGCAGGCCAGCGTATCGTGCTGCGTTTCGATGCGGTCACTCATTACG
GCAAAGTGTGGGTCAATAATCAGGAAGTGATGGAGCATCAGGGCGGCTATACGCCATTTGAAGCCGATGT
CACGCCGTATGTTATTGCCGGGAAAAGTGTACGTATCACCGTTTGTGTGAACAACGAACTGAACTGGCAG
ACTATCCCGCCGGGAATGGTGATTACCGACGAAAACGGCAAGAAAAAGCAGTCTTACTTCCATGATTTCT
TTAACTATGCCGGAATCCATCGCAGCGTAATGCTCTACACCACGCCGAACACCTGGGTGGACGATATCAC
CGTGGTGACGCATGTCGCGCAAGACTGTAACCACGCGTCTGTTGACTGGCAGGTACTTCATGCTTCAACG
TGTAACTTAAGAGATACTGTGTGAAATTTTATATTTCCATACATTTGCTTGACCTTTGCTTTTTGTCAAT
TTTTTTCCCCTTACAGGTGGTGGCCAATGGTGATGTCAGCGTTGAACTGCGTGATGCGGATCAACAGGTG
GTTGCAACTGGACAAGGCACTAGCGGGACTTTGCAAGTGGTGAATCCGCACCTCTGGCAACCGGGTGAAG
GTTATCTCTATGAACTGTGCGTCACAGCCAAAAGCCAGACAGAGTGTGATATCTACCCGCTTCGCGTCGG
CATCCGGTCAGTGGCAGTGAAGGGCGAACAGTTCCTGATTAACCACAAACCGTTCTACTTTACTGGCTTT
GGTCGTCATGAAGATGCGGACTTGCGTGGCAAAGGATTCGATAACGTGCTGATGGTGCACGACCACGCAT
TAATGGACTGGATTGGGGCCAACTCCTACCGTACCTCGCATTACCCTTACGCTGAAGAGATGCTCGACTG
GGCAGATGAACATGGCATCGTGGTGATTGATGAAACTGCTGCTGTCGGCTTTAACCTCTCTTTAGGCATT
GGTTTCGAAGCGGGCAACAAGCCGAAAGAACTGTACAGCGAAGAGGCAGTCAACGGGGAAACTCAGCAAG
CGCACTTACAGGCGATTAAAGAGCTGATAGCGCGTGACAAAAACCACCCAAGCGTGGTGATGTGGAGTAT
TGCCAACGAACCGGATACCCGTCCGCAAGGTGCACGGGAATATTTCGCGCCACTGGCGGAAGCAACGCGT
AAACTCGACCCGACGCGTCCGATCACCTGCGTCAATGTAATGTTCTGCGACGCTCACACCGATACCATCA
GCGATCTCTTTGATGTGCTGTGCCTGAACCGTTATTACGGATGGTATGTCCAAAGCGGCGATTTGGAAAC
GGCAGAGAAGGTACTGGAAAAAGAACTTCTGGCCTGGCAGGAGAAACTGCATCAGCCGATTATCATCACC
GAATACGGCGTGGATACGTTAGCCGGGCTGCACTCAATGTACACCGACATGTGGAGTGAAGAGTATCAGT
GTGCATGGCTGGATATGTATCACCGCGTCTTTGATCGCGTCAGCGCCGTCGTCGGTGAACAGGTATGGAA
TTTCGCCGATTTTGCGACCTCGCAAGGCATATTGCGCGTTGGCGGTAACAAGAAAGGGATCTTCACTCGC
GACCGCAAACCGAAGTCGGCGGCTTTTCTGCTGCAAAAACGCTGGACTGGCATGAACTTCGGTGAAAAAC
CGCAGCAGGGAGGCAAACAATGAGCTTGAAAGAAAATCATTATTCAAATGTTTAACAGCAGTATTAGCTG
AAGTTTGGTGAGCAATTATGAAGAACCTGTAATTTAGCCTCTAAGGTAGTAATAAAACTGATGCACAGGA
CTTCTGTATTTCTT**GCGCTCTCCTCGCCGACTGACTA**ACCCTGTGAATGGTGAAACAGAAGCACTTGTCA
AACTCCAATTGTGCTAAGCGTTTATATTGACAAGTTCTTCGCGACTTTCATTGCTTGCAAAACTTACATG
TTTAATACCATATTACAATCAGGAGAGTTGGGGGACATGGATTATGAAGAAGCAGCTTGACGTCACCACC
ATACCGTTTAGCTTTGCTCAAGACATTCTGACATGCTGCATTGAACAACAAAAATAACAGTATTTCATCT
TGGATTGTAGTACCATCATCGTAGGGAGGAATTAGCATTAACTGTGACTAGGGCATTAGTCCAAATGAAA
CAGGAAATCTTCGTACACAGCCTAACTAAAGAATCACGACCAAGAGATTATATCCTGCATACAAGGATAA
TGTTAATTGATTCAACCAAAGAACTACTCAAGCAAGCAAAAGATAGAACAAATCTTTAAGAAAAGCTTAA
AGAGAAATAAGTACCACAATGTGAAATTGAAATTAATTTAAGCTACAGACTATCTCCACGAGAAACGGTA
CAGCTGATTGATCTACTTTCCTCAATGTAAACTTGCCTTCAAGCCAATGGCACGGGTAATGCCTAAGAAC

AGGGAAGAGAAAGCATCATATACTAGGTATACAACGGATGAGATAACAAAGTAATGTATGAATATGAGAT
TGTGGCAATGATGACGTACACCTTTCACTTGCAACATTACTCAAGACAACTAACTGCATCATATTTCTTC
TGAAAAAATGTTTTATCTACTGAATGCAAGTTCCGCAACCATTAAATAAAAGATCATCGATCTATGTGAT
CAGAAATATTTCAAGCACGCACCTTACTAATGCGATTTACTACTCCACTGGAAAGCTTCATCAAGCTCTT
GTTGCTCAAGG**AGG**AGATAGAATTGGCGCT

**ThiC::pDAP101 transgenic locus (T-DNA insertion)**

5´UTR-ThiC(Exon-(BsaI site)-*Intron*)-insertion-T-DNA(pDAP101) insertion-3´UTR**_sgRNA-ThiC(PAM)**_5´ (510bp) and 3´(472bp) homology arm

CGATTCAGGAGGTTCGTTCCTTTTTTAAAGGACCCTAATCACTCTGAGTACCACTGACTCACTCAGTGTG
CGCGATTCATTTCAAAAACGAGCCAGCCTCTTCTTCCTTCGTCTACTAGATCAGATCCAAAGCTTCCTCT
TCCAG*GTTCGAATCCTTGATTTCTCCATGAATGTGCATGGTAGTCCAACAATTGTCGATGTTTTTGATAG*
*AGAGTTTTGTAGATTTTCTCCGGCGAAATTCCGATTTGTTCTTCAATATTATGTGCATGAAACTTTTTT*
*TTAAGATTGTGCGTTTAGATGCAATATTCGACTCTTTGTTGTTCTCATGCTCGTCGATTTCGATGTGTTT*
*CTGTTAATCCATTGATCGTATCGGAAACTGTGATTGATTGATTCATATTTTCGTTTGTCTCCAG*CTATGG
CTGCTTCAGTACACTGTACCTTGATGTCCGTCGTATGCAACAACAAGAATCACTCTGCTCGGCCGAAACT
TCCAAACTCGTCTTTGTTACCTGGATTCGATGTTGTTGTTCAAGCTGCTGCTACTCGATTCAAGAAGGAA
ACAACAACCACAAGAGCCACTTTGACGTTTGATCCACCAACCACTAATTCTGAGAGAGCTAAGCAGAGAA
AACACACCATTGATCCTTCTTCTCCTGATTTTCAACCAATTCCATCTTTTGAAGAATGCTTTCCTAAGAG
CACTAAAGAACACAA*GTAATTGCTTCACTTAATCTACATTTTTTCATATTGGAAGAGTTGAGAAATCACT*
*GGTTGGTTTTTGGTTGTTTTCAG*GGAAGTTGTCCATGAAGAATCTGGTCATGTTCTTAAAGTTCCCTTTC
GTCGTGTTCATTTGTCTGGTGGTGAGCCAGCTTTTGATAATTATGACACTAGTGGTCCTCAAAATGTTAA
TGCACACATTG*GTATGTGATTCCACCTCGTGTTTACTTTACACATTCACCTCTCTTTTATGTGACTATCG*
*ATAAATGAAACTTACCAAGCAG*GGCTTGCTAAGCTAAGGAAGGAGTGGATTGATCGTCGGGAGAAGCTAG
GAACACCAAGATACACTCAAATGTACTACGCTAAGCAAGGGATCATAACTGAGGAAATGCTCTACTGTGC
TACTAGGGAGAAGCTAGACCCTGAGTTTGTAAGATCAGAAGTTGCACGAGGACGGGCGATTATCCCTTCC
AACAAGAAGCATTTGGAGCTGGAACCGATGATTGTTGGTAGAAAGTTCTTGGTCAAGGTCAATGCGAATA
TCGGAAACTCTGCTGTTGCCAGCTCTATTGAAGAGGAAGTCTACAAGGTTCAGTGGGCAACCATGTGGGG
AGCTGATACAATCATGGATCTCTCAACTGGTCGTCACATCCATGAGACACGTGAGTGGATCCTAAGGAAT
TCAGCTGTGCCTGTTGGTACGGTGCCTATTTATCAAGCACTTGAGAAAGTGGATGGAATTGCTGAGAATC
TTAACTGGGAGGTTTTCAGAGAGACTCTGATTGAACAAGCTGAGCAAGGTGTAGACTATTTCACAATCCA
TGCTGGAGTTTTGCTGCGTTACATTCCCTTAACTGCCAAGCGTTTGACCGGGATCGTTTCGCGTGGAGGA
TCCATTCATGCTAAATGGTGCTTAGCTTACCACAAGGAGAACTTTGCTTACGAGCACTGGGATGACATTC
TAGACATCTGTAACCAGTATGATGTGGCTCTTTCCATTGGAGATGGTCTGAGACCTGGCTCCATTTATGA
TGCTAACGACACTGCTCAGTTTGCAGAGCTCCTTACTCAAGGTGAACTAACTCGCCGAGCGTGGAAAAA
GATGTGCAG*GTATACTAC* ***ACAACTACTTAT*** **ATTCGCCATGG** CATATGCTAGCATGCATAAACTGAAGGCG
GGAAACGACAATCTGATCCAAGCTCAAGCGAGCTCCAGCTTTTGTTCCCTTTAGTGAGGGTTAATTTCGA
GCTTGGCGTAATCATGGTCATAGCTGTTTCCTGTGTGAAATTGTTATCCGCTCACAATTCCACACAACAT
ACGAGCCGGAAGCATAAAGTGTAAAGCCTGGGGTGCCTAATGAGTGAGCTAACTCACATTAATTGCGTTG
CGCTCACTGCCCGCTTTCCAGTCGGGAAACCTGTCGTGCCAGCTGCATTAATGAATCGGCCAACGCGCGG
GGAGAGGCGGTTTGCGTATTGGGCGCTCTTCCGCTTCCTCGCTCACTGACTCGCTGCGCTCGGTCGTTCG
GCTGCGGCGAGCGGTATCAGCTCACTCAAAGGCGGTAATACGGTTATCCACAGAATCAGGGGATAACGCA
GGAAAGAACATGTGAGCAAAAGGCCAGCAAAAGGCCAGGAACCGTAAAAAGGCCGCGTTGCTGGCGTTTT
TCCATAGGCTCCGCCCCCCTGACGAGCATCACAAAAATCGACGCTCAAGTCAGAGGTGGCGAAACCCGAC
AGGACTATAAAGATACCAGGCGTTTCCCCCTGGAAGCTCCCTCGTGCGCTCTCCTGTTCCGACCCTGCCG
CTTACCGGATACCTGTCCGCCTTTCTCCCTTCGGGAAGCGTGGCGCTTTCTCATAGCTCACGCTGTAGGT
ATCTCAGTTCGGTGTAGGTCGTTCGCTCCAAGCTGGGCTGTGTGCACGAACCCCCCGTTCAGCCCGACCG
CTGCGCCTTATCCGGTAACTATCGTCTTGAGTCCAACCCGGTAAGACACGACTTATCGCCACTGGCAGCA
GCCACTGGTAACAGGATTAGCAGAGCGAGGTATGTAGGCGGTGCTACAGAGTTCTTGAAGTGGTGGCCTA

ACTACGGCTACACTAGAAGGACAGTATTTGGTATCTGCGCTCTGCTGAAGCCAGTTACCTTCGGAAAAAG
AGTTGGTAGCTCTTGATCCGGCAAACAAACCACCGCTGGTAGCGGTGGTTTTTTTGTTTGCAAGCAGCAG
ATTACGCGCAGAAAAAAAGGATCTCAAGAAGATCCTTTGATCTTTTCTACGGGGTCTGACGCTCAGTGGA
ACGAAAACTCACGTTAAGGGATTTTGGTCATGAGATTATCAAAAAGGATCTTCACCTAGATCCTTTTAAA
TTAAAAATGAAGTTTTAAATCAATCTAAAGTATATATGAGTAAACTTGGTCTGACAGTTACCAATGCTTA
ATCAGTGAGGCACCTATCTCAGCGATCTGTCTATTTCGTTCATCCATAGTTGCCTGACTCCCCGTCGTGT
AGATAACTACGATACGGGAGGGCTTACCATCTGGCCCCAGTGCTGCAATGATACCGCGAGACCCACGCTC
ACCGGCTCCAGATTTATCAGCAATAAACCAGCCAGCCGGAAGGGCCGAGCGCAGAAGTGGTCCTGCAACT
TTATCCGCCTCCATCCAGTCTATTAATTGTTGCCGGGAAGCTAGAGTAAGTAGTTCGCCAGTTAATAGTT
TGCGCAACGTTGTTGCCATTGCTACAGGCATCGTGGTGTCACGCTCGTCGTTTGGTATGGCTTCATTCAG
CTCCGGTTCCCAACGATCAAGGCGAGTTACATGATCCCCCATGTTGTGCAAAAAAGCGGTTAGCTCCTTC
GGTCCTCCGATCGTTGTCAGAAGTAAGTTGGCCGCAGTGTTATCACTCATGGTTATGGCAGCACTGCATA
ATTCTCTTACTGTCATGCCATCCGTAAGATGCTTTTCTGTGACTGGTGAGTACTCAACCAAGTCATTCTG
AGAATAGTGTATGCGGCGACCGAGTTGCTCTTGCCCGGCGTCAATACGGGATAATACCGCGCCACATAGC
AGAACTTTAAAAGTGCTCATCATTGGAAAACGTTCTTCGGGGCGAAAACTCTCAAGGATCTTACCGCTGT
TGAGATCCAGTTCGATGTAACCCACTCGTGCACCCAACTGATCTTCAGCATCTTTTACTTTCACCAGCGT
TTCTGGGTGAGCAAAAACAGGAAGGCAAAATGCCGCAAAAAAGGGAATAAGGGCGACACGGAAATGTTGA
ATACTCATACTCTTCCTTTTTCAATATTATTGAAGCATTTATCAGGGTTATTGTCTCATGAGCGGATACA
TATTTGAATGTATTTAGAAAAATAAACAAATAGGGGTTCCGCGCACATTTCCCCGAAAAGTGCCACCTAA
ATTGTAAGCGTTAATATTTTGTTAAAATTCGCGTTAAATTTTTGTTAAATCAGCTCATTTTTTAACCAAT
AGGCCGAAATCGGCAAAATCCCTTATAAATCAAAAGAATAGACCGAGATAGGGTTGAGTGTTGTTCCAGT
TTGGAACAAGAGTCCACTATTAAAGAACGTGGACTCCAACGTCAAAGGGCGAAAAACCGTCTATCAGGGC
GATGGCCCACTACGTGAACCATCACCCTAATCAAGTTTTTTGGGGTCGAGGTGCCGTAAAGCACTAAATC
GGAACCCTAAAGGGAGCCCCCGATTTAGAGCTTGACGGGGAAAGCCGGCGAACGTGGCGAGAAAGGAAGG
GAAGAAAGCGAAAGGAGCGGGCGCTAGGGCGCTGGCAAGTGTAGCGGTCACGCTGCGCGTAACCACCACA
CCCGCCGCGCTTAATGCGCCGCTACAGGGCGCGTCCCATTCGCCATTCAGGCTGCGCAACTGTTGGGAAG
GGCGATCGGTGCGGGCCTCTTCGCTATTACGCCAGCTGGCGAAAGGGGGATGTGCTGCAAGGCGATTAAG
TTGGGTAACGCCAGGGTTTTCCCAGTCACGACGTTGTAAAACGACGGCCAGTGAGCGCGCGTAATACGAC
TCACTATAGGGCGAATTGGGTACCGGGCCCCCCCTCGAGGTCGACGGTATCGATAAGCTTGATATCGAAT
TCGAGCTCGGTACCCACTGGATTTTGGTTTTAGGAATTAGAAATTTTATTGATAGAAGTATTTTACAAAT
ACAAATACATACTAAGGGTTTCTTATATGCTCAACACATGAGCGAAACCCTATAAGAACCCTAATTCCCT
TATCTGGGAACTACTCACACATTATTATAGAGAGAGATAGATTTGTAGAGAGAGACTGGTGATTTCAGCG
GCATGCCTGCAGGTCGACTCTAGAGGATCCTAGACGCGTGAGATCAGATCTCGGTGACGGGCAGGACCGG
ACGGGGCGGTACCGGCAGGCTGAAGTCCAGCTGCCAGAAACCCACGTCATGCCAGTTCCCGTGCTTGAAG
CCGGCCGCCCGCAGCATGCCGCGGGGGGCATATCCGAGCGCCTCGTGCATGCGCACGCTCGGGTCGTTGG
GCAGCCCGATGACAGCGACCACGCTCTTGAAGCCCTGTGCCTCCAGGGACTTCAGCAGGTGGGTGTAGAG
CGTGGAGCCCAGTCCCGTCCGCTGGTGGCGGGGGGAGACGTACACGGTCGACTCGGCCGTCCAGTCGTAG
GCGTTGCGTGCCTTCCAGGGGCCCGCGTAGGCGATGCCGGCGACCTCGCCGTCCACCTCGGCGACGAGCC
AGGGATAGCGCTCCCGCAGACGGACGAGGTCGTCCGTCCACTCCTGCGGTTCCTGCGGCTCGGTACGGAA
GTTGACCGTGCTTGTCTCGATGTAGTGGTTGACGATGGTGCAGACCGCCGGCATGTCCGCCTCGGTGGCA
CGGCGGATGTCGGCCGGGCGTCGTTCTGGGCTCATGGATCCACGTGTGGAAGATATGAATTTTTTTGAGA
AACTAGATAAGATTAATGAATATCGGTGTTTTGGTTTTTTCTTGTGGCCGTCTTTGTTTATATTGAGATT
TTTCAAATCAGTGCGCAAGACGTGACGTAAGTATCCGAGTCAGTTTTTATTTTTCTACTAATTTGGTCGT
TTATTTCGGCGTGTAGGACATGGCAACCGGGCCTGAATTTCGCGGGTATTCTGTTTCTATTCCAACTTTT
TCTTGATCCGCAGCCATTAACGACTTTTGAATAGATACGCTGACACGCCAAGCCTCGCTAGTCAAAAGTG
TACCAAACAACGCTTTACAGCAAGAACGGAATGCGCGTGACCGTCGCGGTGACGCCATTTCGCCTTTTCA
GAAATGGATAAATAGCCTTGCTTCCTATTATATCTTCCCAAATTACCAATACATTACACTAGCATCTGAA
TTTCATAACCAATCTCGATACACCAAATCGAATTCAATTCGGCGTTAATTCAGTACATTAAAAACGTCCG
CAATGTGTTATTAAGTTGTCTAAGCGTCAATTTGTTTACACCACAATATATCCTG*GATACAACTACTTAT*
*ATCTACTTTTCCAG*GTGATGAATGAAGGGCCAGGGCATGTCCCAATGCACAAGATTCCAGAGAATATGCA
GAAGCAGTTGGAGTGGTGTAACGAGGCACCATTCTACACCCTTGGTCCTTTGACTACTGATATTGCCCCT
GGATATGATCACATTACCTCTGCCATTGGAGCTGCCAATATTGGAGCCTTGGGTACAGCTCTTCTTTGCT
ATGTAACACCAAAAGAACACCTTGGGCTACCAAACAGGGACGATGTGAAGGCCGGGGTTATAGCATACAA

GATCGCCGCTCATGCAGCTGATCTAGCCAAACAGCATCCACATGCTCAGGCATGGGACGATGCGCTGAGC
AAAGCGCGGTTTGAGTTTAGATGGATGGACCAATTTGCTCTGTCGTTGGACCCCATGACTGCTATGTCTT
TCCATGATGAAACTCTTCCAGCTGATGGAGCCAAGGTTGCACACTTTTGCTCCATGTGTGGACCAAAATT
CTGCTCTATGAAGATAACAGAAGACATCCGAAAGTATGCAGAGGAGAATGGTTATGGCAGTGCTGAAGAA
GCAATCAGACAAGGAATGGATGCTATGAGTGAAGAATTCAACATCGCAAAGAAAACCATTAGCGGAGAAC
AGCACGGTGAAGTAGGTGGAGAAATATATTTGCCAGAGAGCTATGTCAAAGCTGCTCAGAAATAAAAG*GT*
*CAGTATGTTTAGACTGTTAGTCGTTGCTTTCTCAACAAACATGTTAGTTACTGCATGCTAGTATAAAATC*
*ATTCAGGTTTATAATCTTTTCTTAAATCTGCAACATATGGTCAACTCTTAAATGAGTCCTTACTGTGATC*
*TTTGTTTTTTATCGTGTTTCTTTTTCTTCTGCTGCATCAG*GCAAATGTTTTAAACAAGACCTTGCTTACC
CAAGTCTTGGTGCCTGTTGGACTATACCTGGATAAAGGCACAAACTGTTGGTAAGCTTAGTAGTCTCTAT
GTCATGTTACTTTTAGAACTATCTATGTTGTCTGTTCATTTGAGTCAGAGTCAGCAATAAAGACAATCTA
AGTTGATGTTTCAATACTTTTTTGTGTGATTTGGTTGGTGAATTGACATGCAAAAGCACCAGGGGTGCTT
GAACCAGGATAGCCTGCGAAAAGGCGGGCTATCCGGGACCAGGCTGAGAAAGTCCCTTTGAACCTGAACA
GGGTAATGCCTGCGCAGGGAGTGTGCAGTTTTTTTTTTTTCCTGTAGCTTTCTAAAGGAGAAGAAGCTAC
TGTTGCCGCTCGAGTCTCGTTCCACGGTTTTCAACAGTTAGTTTCTTATGAGCTAAGAGATTCAGCTTAA
TTGGCTTACAGCCATAAAAGAAGTCTTTAACTGATGCACTAAGTCACTAACAGTAGGGAATAATTCAATC
AAAAAATCATCCAGATTGATAAAAATGCATTTGCACCTTTGGGGCATAAGCTGAAATTACTCTGCTCGCA
CAAATCAGATTTTTAAAACTTATGATCATGTTTTCAACTTATACTCGTTTGTTTACATAATGGGATGATC
AGTTGTTCACTTTGAGCAAAGCATACTGGGCAGGTGTAGGGAAAGTGAAGCTCATGAAACTGATCCACTG
TAATTCTCATTGTTCAGCTTTTATTTACACAAATGTGCGAGTCAACACGTAAAAGATTATGATATATTGC
AGTAGAACATAAGTAAAACGACCTTACCAAATGCGAAAGAAAGCTTTTCGAGCCGGAATGATCAAAAGGT
TACACATTCTGTACTCATTTTCCTGCTATATGAGATCACACAATTATCAAGAGCCTGAAACTCTGGAATA
CCAATATTTAGACACGTTACATAGAGTTTTGAAGGTTTCTCATTTGACCAGCTCCTGTGTTTGACATGCT
TTGGTTTCATATGGTTAATGTTTCTGCTAAATCAACCATGAGAGATGTGAACTATTTCTATATCATGTGT
CTGTAACGTTGGGGCCGCTTGTGAGCCTCTTCCTTCACCATTTTCAGTTTGATCAGTTAATAACCTCATG
TAAAAGCAGTGAGAGAGAGAAGAGATCAGAGATGGCACACTGTTTCCTTTGGAAATCCTGGGCGTTGTTA
TGTTTGATCGTAGGCTTTGTGACTGGTAAAGAAAAAAACAGCATGTATGTTATGTTCAAAATACTAAAAA
ACGACTCGTAG

**pAGT9813 (Level 1): Donor to repair ThiC::pDAP101 transgenic locus**

GG-overhang-sgRNA-target D1 (**PAM**); 5´HA(510 bp); ThiC(Exon-**mutated BsaI site;** *intron*); **sgRNA-ThiC(PAM);** Insertion**;** synThiC(Exon); tOCS; 3´UTR; T-DNA(pDAP101) insertion; 3´HA(472bp); sgRNA-target D1 (**PAM**)-GG-overhang

GGAGCCAATTCTATCT**CCT**CCTTGAGCAACAAGAGCTTGATGAAGCTTTCCTTGGGATCTCTCAACTGGT
CGTCACATCCATGAGACACGTGAGTGGATCCTAAGGAATTCAGCTGTGCCTGTTGGTACGGTGCCTATTT
ATCAAGCACTTGAGAAAGTGGATGGAATTGCTGAGAATCTTAACTGGGAGGTTTTCAGAGAGACTCTGAT
TGAACAAGCTGAGCAAGGTGTAGACTATTTCACAATCCATGCTGGAGTTTTGCTGCGTTACATTCCCTTA
ACTGCCAAGCGTTTGACCGGGATCGTTTCGCGTGGAGGATCCATTCATGCTAAATGGTGCTTAGCTTACC
ACAAGGAGAACTTTGCTTACGAGCACTGGGATGACATTCTAGACATCTGTAACCAGTATGATGTGGCTCT
TTCCATTGGAGATGGTCTGAG**G**CCTGGCTCCATTTATGATGCTAACGACACTGCTCAGTTTGCAGAGCTC
CTTACTCAAGGTGAACTAACTCGCCGAGCGTGGAAAAAGATGTGCAGGTATACTAC**ACAACTACTTATA**
**TTCG**ATATCCTGGATACAACTACTTATATCTACTTTTCCAGGTGATGAATGAAGGACCTGGTCATGTGCC
GATGCACAAGATCCCTGAGAACATGCAAAAGCAGCTCGAGTGGTGTAACGAGGCTCCTTTCTATACTCTC
GGACCGCTGACTACTGATATCGCTCCTGGATACGATCACATCACCTCTGCTATTGGGGCTGCAAACATTG
GTGCTCTTGGAACTGCTCTCCTCTGCTACGTTACCCCTAAAGAGCATCTTGGACTCCCTAACAGGGATGA
CGTTAAGGCTGGTGTGATCGCTTACAAGATCGCTGCTCATGCTGCTGACCTTGCTAAGCAACATCCTCAT
GCTCAGGCTTGGGATGACGCTCTTTCTAAGGCTAGGTTCGAGTTCCGTTGGATGGACCAGTTCGCTCTCT
CTCTCGATCCTATGACCGCTATGTCTTTCCACGATGAGACTCTCCCTGCTGACGGTGCTAAAGTTGCTCA
CTTCTGTTCTATGTGCGGGCCGAAGTTCTGCAGCATGAAGATCACTGAGGACATCCGTAAGTACGCCGAA
GAGAACGGATATGGGTCTGCTGAAGAGGCTATCAGGCAAGGTATGGATGCCATGAGCGAAGAGTTCAACA

TTGCCAAGAAAACCATCAGCGGAGAGCAGCACGGTGAAGTTGGAGGTGAAATCTACCTGCCTGAGAGCTA
TGTGAAGGCCGCTCAAAAGTGAGCTTGTCCTGCTTTAATGAGATATGCGAGAAGCCTATGATCGCATGAT
ATTTGCTTTCAATTCTGTTGTGCACGTTGTAAAAAACCTGAGCATGTGTAGCTCAGATCCTTACCGCCGG
TTTCGGTTCATTCTAATGAATATATCACCCGTTACTATCGTATTTTTATGAATAATATTCTCCGTTCAAT
TTACTGATTGTACCCTACTACTTATATGTACAATATTAAAATGAAAACAATATATTGTGCTGAATAGGTT
TATAGCGACATCTATGATAGAGCGCCACAATAACAAACAATTGCGTTTTATTATTACAAATCCAATTTTA
AAAAAAGCGGCAGAACCGGTCAAACCTAAAAGACTGATTACATAAATCTTATTCAAATTTCAAAAGTGCC
CCAGGGGCTAGTATCTACGACACACCGAGCGGCGAACTAATAACGCTCACTGAAGGGAACTCCGGTTCCC
CGCCGGCGCGCATGGGTGAGATTCCTTGAAGTTGAGTATTGGCCGTCCGCTCTACCGAAAGTTACGGGCA
CCATTCAACCCGGTCCAGCACGGCGGCCGGGTAACCGACTTGCTGCCCCGAGAATTATGCAGCATTTTTT
TGGTGTATGTGGGCCCCAAATGAAGTGCAGGTCAAACCTTGACAGTGACGACAAATCGTTGGGCGGGTCC
AGGGCGAATTTTGCGACAACATGTCGAGGCTCAGCAGGACCGCT**CCATGG**CATATGCTAGCATGCATAAA
CTGAAGGCGGGAAACGACAATCTGATCCAAGCTCAAGCGAGCTCCAGCTTTTGTTCCCTTTAGTGAGGGT
TAATTTCGAGCTTGGCGTAATCATGGTCATAGCTGTTTCCTGTGTGAAATTGTTATCCGCTCACAATTCC
ACACAACATACGAGCCGGAAGCATAAAGTGTAAAGCCTGGGGTGCCTAATGAGTGAGCTAACTCACATTA
ATTGCGTTGCGCTCACTGCCCGCTTTCCAGTCGGGAAACCTGTCGTGCCAGCTGCATTAATGAATCGGCC
AACGCGCGGGGAGAGGCGGTTTGCGTATTGGGCGCTCTTCCGCTTCCTCGCTCACTGACTCGCTGCGCTC
GGTCGTTCGGCTGCGGCGAGCGGTATCAGCTCACTCAAAGGCGGTAATACGGTTATCCACAGAATCAGGG
GATAACGCAGGAAAGAACATGTGAGCCCACTGGAAAGCTTCATCAAGCTCTTGTTGCTCAAGG**AGG**AGAT
AGAATTGGCGCT

**pAGT8225 (pET28a-based GG-compatible plasmid for expression of His-tagged proteins in E.coli; Kan^R^)**

pT7-T7 translational enhancer-GG-Overhang-**BsaI**-pLac-LacZalpha-**BsaI**-GG-overhang-***6xHis***- **tT7**

CCCGCGAAATTAATACGACTCACTATAGGGGAATTGTGAGCGGATAACAATTCCCCTCTAGAAATAATTTTGTTTAA
CTTTAAGAAGGAGATATACCATG**GAGACC**TGCAGCTGGCACGACAGGTTTGCCGACTGGAAAGCGGGCAGTGAGCGC
AACGCAATTAATGTGAGTTAGCTCACTCATTAGGCACCCCAGGCTTTACACTTTATGCTTCCGGCTCGTATGTTGTG
TGGAATTGTGAGCGGATAACAATTTCACACAGGAAACAGCTATGACCATGATTACGCCAAGCTTGCATGCCTGCAGG
TCGACTCTAGAGGATCCCCGGGTACCGAGCTCGAATTCACTGGCCGTCGTTTTACAACGTCGTGACTGGGAAAACCC
TGGCGTTACCCAACTTAATCGCCTTGCAGCACATCCCCCTTTCGCCAGCTGGCGTAATAGCGAAGAGGCCCGCACCG
ATCGCCCTTCCCAACAGTTGCGCAGCCTGAATGGCGAATGGCGCCTGATGCGGTATTTTCTCCTTACGCATCTGTGC
GGTATTTCACACCGCATATGGTGCACTCTCAGTACAATCTGCTCTGATGCCGCATAGTTAAGCCAGCCCCGACACCC
GCCAACACCCGCTGACGCGCCCTGACGGGCTTGTCTGCTCCCGGCATCCGCTTACAGACAAGCTGTGACT**GGTCTC**T
TTCGCTCGAG***CACCACCACCACCACCAC***TGAGATCCGG**CTGCTAACAAAGCCCGAAAGGAAGCTGAGTTGGCTGCTG**
**CCACCGCTGAGCAATAACTAGCATAACCCCTTGGGGCCTCTAAACGGGTCTTGAGGGGTTTTTTGCTGAAAGGAGGA**
**ACTATATCCGGAT**TGGCGAATGGGACGCGCCCTGTAGCGGCGCATTAAGCGCGGCGGGTGTGGTGGTTACGCGCAGC
GTGACCGCTACACTTGCCAGCGCCCTAGCGCCCGCTCCTTTCGCTTTCTTCCCTTCCTTTCTCGCCACGTTCGCCGG
CTTTCCCCGTCAAGCTCTAAATCGGGGGCTCCCTTTAGGGTTCCGATTTAGTGCTTTACGGCACCTCGACCCCAAAA
AACTTGATTAGGGTGATGGTTCACGTAGTGGGCCATCGCCCTGATAGACGGTTTTTCGCCCTTTGACGTTGGAGTCC
ACGTTCTTTAATAGTGGACTCTTGTTCCAAACTGGAACAACACTCAACCCTATCTCGGTCTATTCTTTTGATTTATA
AGGGATTTTGCCGATTTCGGCCTATTGGTTAAAAAATGAGCTGATTTAACAAAAATTTAACGCGAATTTTAACAAAA
TATTAACGTTTACAATTTCAGGTGGCACTTTTCGGGGAAATGTGCGCGGAACCCCTATTTGTTTATTTTTCTAAATA
CATTCAAATATGTATCCGCTCATGAATTAATTCTTAGAAAAACTCATCGAGCATCAAATGAAACTGCAATTTATTCA
TATCAGGATTATCAATACCATATTTTTGAAAAAGCCGTTTCTGTAATGAAGGAGAAAACTCACCGAGGCAGTTCCAT
AGGATGGCAAGATCCTGGTATCGGTCTGCGATTCCGACTCGTCCAACATCAATACAACCTATTAATTTCCCCTCGTC
AAAAATAAGGTTATCAAGTGAGAAATCACCATGAGTGACGACTGAATCCGGTGAGAATGGCAAAAGTTTATGCATTT
CTTTCCAGACTTGTTCAACAGGCCAGCCATTACGCTCGTCATCAAAATCACTCGCATCAACCAAACCGTTATTCATT
CGTGATTGCGCCTGAGCGAGACGAAATACGCGATCGCTGTTAAAAGGACAATTACAAACAGGAATCGAATGCAACCG
GCGCAGGAACACTGCCAGCGCATCAACAATATTTTCACCTGAATCAGGATATTCTTCTAATACCTGGAATGCTGTTT
TCCCGGGGATCGCAGTGGTGAGTAACCATGCATCATCAGGAGTACGGATAAAATGCTTGATGGTCGGAAGAGGCATA
AATTCCGTCAGCCAGTTTAGTCTGACCATCTCATCTGTAACATCATTGGCAACGCTACCTTTGCCATGTTTCAGAAA
CAACTCTGGCGCATCGGGCTTCCCATACAATCGATAGATTGTCGCACCTGATTGCCCGACATTATCGCGAGCCCATT
TATACCCATATAAATCAGCATCCATGTTGGAATTTAATCGCGGCCTAGAGCAAGACGTTTCCCGTTGAATATGGCTC
ATAACACCCCTTGTATTACTGTTTATGTAAGCAGACAGTTTTATTGTTCATGACCAAAATCCCTTAACGTGAGTTTT

CGTTCCACTGAGCGTCAGACCCCGTAGAAAAGATCAAAGGATCTTCTTGAGATCCTTTTTTTCTGCGCGTAATCTGC
TGCTTGCAAACAAAAAAACCACCGCTACCAGCGGTGGTTTGTTTGCCGGATCAAGAGCTACCAACTCTTTTTCCGAA
GGTAACTGGCTTCAGCAGAGCGCAGATACCAAATACTGTCCTTCTAGTGTAGCCGTAGTTAGGCCACCACTTCAAGA
ACTCTGTAGCACCGCCTACATACCTCGCTCTGCTAATCCTGTTACCAGTGGCTGCTGCCAGTGGCGATAAGTCGTGT
CTTACCGGGTTGGACTCAAGACGATAGTTACCGGATAAGGCGCAGCGGTCGGGCTGAACGGGGGGTTCGTGCACACA
GCCCAGCTTGGAGCGAACGACCTACACCGAACTGAGATACCTACAGCGTGAGCTATGAGAAAGCGCCACGCTTCCCG
AAGGGAGAAAGGCGGACAGGTATCCGGTAAGCGGCAGGGTCGGAACAGGAGAGCGCACGAGGGAGCTTCCAGGGGGA
AACGCCTGGTATCTTTATAGTCCTGTCGGGTTTCGCCACCTCTGACTTGAGCGTCGATTTTTGTGATGCTCGTCAGG
GGGGCGGAGCCTATGGAAAAACGCCAGCAACGCGGCCTTTTTACGGTTCCTGGCCTTTTGCTGGCCTTTTGCTCACA
TGTTCTTTCCTGCGTTATCCCCTGATTCTGTGGATAACCGTATTACCGCCTTTGAGTGAGCTGATACCGCTCGCCGC
AGCCGAACGACCGAGCGCAGCGAGTCAGTGAGCGAGGAAGCGGAAGAGCGCCTGATGCGGTATTTTCTCCTTACGCA
TCTGTGCGGTATTTCACACCGCATATATGGTGCACTCTCAGTACAATCTGCTCTGATGCCGCATAGTTAAGCCAGTA
TACACTCCGCTATCGCTACGTGACTGGGTCATGGCTGCGCCCCGACACCCGCCAACACCCGCTGACGCGCCCTGACG
GGCTTGTCTGCTCCCGGCATCCGCTTACAGACAAGCTGTGACCGTCTCCGGGAGCTGCATGTGTCAGAGGTTTTCAC
CGTCATCACCGAAACGCGCGAGGCAGCTGCGGTAAAGCTCATCAGCGTGGTCGTGAAGCGATTCACAGATGTCTGCC
TGTTCATCCGCGTCCAGCTCGTTGAGTTTCTCCAGAAGCGTTAATGTCTGGCTTCTGATAAAGCGGGCCATGTTAAG
GGCGGTTTTTTCCTGTTTGGTCACTGATGCCTCCGTGTAAGGGGGATTTCTGTTCATGGGGGTAATGATACCGATGA
AACGAGAGAGGATGCTCACGATACGGGTTACTGATGATGAACATGCCCGGTTACTGGAACGTTGTGAGGGTAAACAA
CTGGCGGTATGGATGCGGCGGGACCAGAGAAAAATCACTCAGGGTCAATGCCAGCGCTTCGTTAATACAGATGTAGG
TGTTCCACAGGGTAGCCAGCAGCATCCTGCGATGCAGATCCGGAACATAATGGTGCAGGGCGCTGACTTCCGCGTTT
CCAGACTTTACGAAACACGGAAACCGAAGACCATTCATGTTGTTGCTCAGGTCGCAGACGTTTTGCAGCAGCAGTCG
CTTCACGTTCGCTCGCGTATCGGTGATTCATTCTGCTAACCAGTAAGGCAACCCCGCCAGCCTAGCCGGGTCCTCAA
CGACAGGAGCACGATCATGCGCACCCGTGGGGCCGCCATGCCGGCGATAATGGCCTGCTTCTCGCCGAAACGTTTGG
TGGCGGGACCAGTGACGAAGGCTTGAGCGAGGGCGTGCAAGATTCCGAATACCGCAAGCGACAGGCCGATCATCGTC
GCGCTCCAGCGAAAGCGGTCCTCGCCGAAAATGACCCAGAGCGCTGCCGGCACCTGTCCTACGAGTTGCATGATAAA
GAAGACAGTCATAAGTGCGGCGACGATAGTCATGCCCCGCGCCCACCGGAAGGAGCTGACTGGGTTGAAGGCTCTCA
AGGGCATCGGTCGAGATCCCGGTGCCTAATGAGTGAGCTAACTTACATTAATTGCGTTGCGCTCACTGCCCGCTTTC
CAGTCGGGAAACCTGTCGTGCCAGCTGCATTAATGAATCGGCCAACGCGCGGGGAGAGGCGGTTTGCGTATTGGGCG
CCAGGGTGGTTTTTCTTTTCACCAGTGAGACGGGCAACAGCTGATTGCCCTTCACCGCCTGGCCCTGAGAGAGTTGC
AGCAAGCGGTCCACGCTGGTTTGCCCCAGCAGGCGAAAATCCTGTTTGATGGTGGTTAACGGCGGGATATAACATGA
GCTGTCTTCGGTATCGTCGTATCCCACTACCGAGATATCCGCACCAACGCGCAGCCCGGACTCGGTAATGGCGCGCA
TTGCGCCCAGCGCCATCTGATCGTTGGCAACCAGCATCGCAGTGGGAACGATGCCCTCATTCAGCATTTGCATGGTT
TGTTGAAAACCGGACATGGCACTCCAGTCGCCTTCCCGTTCCGCTATCGGCTGAATTTGATTGCGAGTGAGATATTT
ATGCCAGCCAGCCAGACGCAGACGCGCCGAGACAGAACTTAATGGGCCCGCTAACAGCGCGATTTGCTGGTGACCCA
ATGCGACCAGATGCTCCACGCCCAGTCGCGTACCGTCTTCATGGGAGAAAATAATACTGTTGATGGGTGTCTGGTCA
GAGACATCAAGAAATAACGCCGGAACATTAGTGCAGGCAGCTTCCACAGCAATGGCATCCTGGTCATCCAGCGGATA
GTTAATGATCAGCCCACTGACGCGTTGCGCGAGAAGATTGTGCACCGCCGCTTTACAGGCTTCGACGCCGCTTCGTT
CTACCATCGACACCACCACGCTGGCACCCAGTTGATCGGCGCGAGATTTAATCGCCGCGACAATTTGCGACGGCGCG
TGCAGGGCCAGACTGGAGGTGGCAACGCCAATCAGCAACGACTGTTTGCCCGCCAGTTGTTGTGCCACGCGGTTGGG
AATGTAATTCAGCTCCGCCATCGCCGCTTCCACTTTTTCCCGCGTTTTCGCAGAAACGTGGCTGGCCTGGTTCACCA
CGCGGGAAACGGTCTGATAAGAGACACCGGCATACTCTGCGACATCGTATAACGTTACTGGTTTCACATTCACCACC
CTGAATTGACTCTCTTCCGGGCGCTATCATGCCATACCGCGAAAGGTTTTGCGCCATTCGATGGTGTCCGGGATCTC
GACGCTCTCCCTTATGCGACTCCTGCATTAGGAAGCAGCCCAGTAGTAGGTTGAGGCCGTTGAGCACCGCCGCCGCA
AGGAATGGTGCATGCAAGGAGATGGCGCCCAACAGTCCCCCGGCCACGGGGCCTGCCACCATACCCACGCCGAAACA
AGCGCTCATGAGCCCGAAGTGGCGAGCCCGATCTTCCCCATCGGTGATGTCGGCGATATAGGCGCCAGCAACCGCAC
CTGTGGCGCCGGTGATGCCGGCCACGATGCGTCCGGCGTAGAGGATCGAGATCTCGAT

**pAGT8226 (6xHis-Thrombin: CCAT_6xHis-Thrombin_AATG)**
GG-overhang-***6xHis***-**Thrombin**-GG-overhang
CCATGGGCAGCAGC***CATCATCATCATCATCAC***AGCAGCGGC**CTGGTGCCGCGCGGCTC**AATG

**pAGT8228 (UL12 (codon optimized for E.coli): AATG_UL12-no stop_TTCG)**
GG-overhang-UL12(E.coli)-GG-overhang
AATGGAATCTACTGTGGGTCCTGCTTGTCCTCCTGGTAGGACTGTTACTAAGAGGCCTTGGGCTCTTGCTGAGGATA
CTCCTAGAGGTCCTGACTCTCCACCAAAGAGGCCTAGACCTAACTCTCTTCCTCTGACTACTACCTTCAGGCCTTTG
CCACCTCCTCCACAAACTACCTCTGCTGTGGATCCTTCTTCTCACTCTCCTGTGAATCCTCCAAGGGATCAGCATGC
TACTGATACCGCTGATGAGAAGCCTAGAGCTGCTTCTCCTGCTCTGTCTGATGCTTCTGGTCCTCCTACTCCTGATA
TCCCTCTTTCTCCTGGTGGTACTCATGCTAGGGATCCTGATGCAGATCCTGATAGCCCTGATCTGGACTCTATGTGG

TCTGCTTCTGTGATCCCTAACGCTCTGCCTTCTCACATTCTGGCTGAGACTTTCGAGAGGCACCTTAGGGGTTTGCT
TAGAGGTGTTAGGGCTCCTCTTGCTATTGGTCCTCTTTGGGCTAGACTGGACTACCTTTGCTCTCTTGCTGTGGTGC
TTGAAGAGGCTGGTATGGTGGATAGAGGTCTTGGTAGACACCTTTGGAGGCTTACTAGAAGAGGTCCTCCAGCTGCT
GCTGATGCTGTTGCTCCTAGACCTCTTATGGGATTCTACGAGGCTGCTACTCAGAACCAGGCTGATTGTCAACTTTG
GGCTCTGCTTAGAAGGGGTCTTACTACCGCTTCTACTCTTAGATGGGGTCCTCAGGGACCTTGCTTTTCTCCTCAAT
GGCTGAAACACAACGCTAGCCTTAGGCCTGATGTGCAGTCATCTGCTGTGATGTTCGGTAGGGTTAACGAGCCTACC
GCTCGGTCTTTGCTTTTCAGGTATTGCGTTGGCAGGGCTGATGATGGTGGTGAAGCTGGTGCTGATACCAGGCGGTT
TATTTTCCACGAGCCATCTGATCTGGCCGAAGAGAATGTTCATACCTGCGGTGTGCTTATGGATGGTCACACTGGAA
TGGTGGGCGCTTCTCTTGATATTCTTGTGTGCCCTAGGGACATCCACGGTTACCTTGCTCCAGTTCCTAAGACTCCT
CTGGCCTTTTACGAGGTTAAGTGCAGGGCTAAGTACGCTTTCGATCCTATGGACCCTTCTGACCCTACTGCTTCTGC
TTACGAGGATCTGATGGCTCATAGAAGCCCTGAGGCTTTCAGGGCTTTCATCCGGTCTATTCCTAAGCCGAGCGTGA
GATACTTTGCTCCTGGAAGAGTTCCTGGTCCTGAGGAAGCTCTTGTTACTCAAGATCAGGCTTGGTCTGAGGCTCAT
GCTTCAGGTGAGAAGAGAAGATGCTCAGCTGCTGATAGGGCACTCGTTGAGCTTAATTCTGGCGTGGTGTCTGAGGT
GTTGCTTTTCGGTGCTCCTGATCTCGGTAGGCACACTATTTCTCCAGTGAGCTGGTCCTCTGGTGATCTTGTTAGAA
GGGAACCCGTGTTCGCTAATCCTAGGCACCCTAACTTCAAGCAGATTCTGGTGCAGGGTTACGTGCTGGATTCTCAC
TTTCCAGATTGCCCTCCACATCCTCACCTTGTGACTTTCATTGGTCGGCATAGGACCTCAGCTGAAGAGGGTGTTAC
TTTCAGGCTTGAGGATGGTGCTGGTGCTCTTGGTGCTGCTGGTCCTTCTAAGGCTTCTATTCTTCCTAACCAGGCCG
TGCCTATCGCTCTTATTATCACCCCTGTGAGGATCGACCCCGAGATCTATAAGGCTATCCAGAGGTCATCTCGGCTG
GCTTTCGATGATACTTTGGCTGAGCTTTGGGCCTCTAGATCTCCTGGTCCAGGTCCTGCTGCTGCAGAAACTACTTC
TTCTTCACCTACCACCGGCAGGTCATCTAGAGGTTCG

**pAGT78232 (pAGT8225-based expression vector: pT7_6xHis-Thrombin-UL12-6xHis_tT7; Kan$^{R}$)**

pT7-T7 translational enhancer-GG-Overhang-***6xHis***-**Thrombin**-UL12-***6xHis*** –GG-overhang-**tT7**

TAATACGACTCACTATAGGGGAATTGTGAGCGGATAACAATTCCCCTCTAGAAATAATTTTGTTTAACTTTAAGAAG
GAGATATACCATGGGCAGCAGC***CATCATCATCATCATCAC***AGCAGCGGC**CTGGTGCCGCGCGGCTC**AATGGAATCTA
CTGTGGGTCCTGCTTGTCCTCCTGGTAGGACTGTTACTAAGAGGCCTTGGGCTCTTGCTGAGGATACTCCTAGAGGT
CCTGACTCTCCACCAAAGAGGCCTAGACCTAACTCTCTTCCTCTGACTACTACCTTCAGGCCTTTGCCACCTCCTCC
ACAAACTACCTCTGCTGTGGATCCTTCTTCTCACTCTCCTGTGAATCCTCCAAGGGATCAGCATGCTACTGATACCG
CTGATGAGAAGCCTAGAGCTGCTTCTCCTGCTCTGTCTGATGCTTCTGGTCCTCCTACTCCTGATATCCCTCTTTCT
CCTGGTGGTACTCATGCTAGGGATCCTGATGCAGATCCTGATAGCCCTGATCTGGACTCTATGTGGTCTGCTTCTGT
GATCCCTAACGCTCTGCCTTCTCACATTCTGGCTGAGACTTTCGAGAGGCACCTTAGGGGTTTGCTTAGAGGTGTTA
GGGCTCCTCTTGCTATTGGTCCTCTTTGGGCTAGACTGGACTACCTTTGCTCTCTTGCTGTGGTGCTTGAAGAGGCT
GGTATGGTGGATAGAGGTCTTGGTAGACACCTTTGGAGGCTTACTAGAAGAGGTCCTCCAGCTGCTGCTGATGCTGT
TGCTCCTAGACCTCTTATGGGATTCTACGAGGCTGCTACTCAGAACCAGGCTGATTGTCAACTTTGGGCTCTGCTTA
GAAGGGGTCTTACTACCGCTTCTACTCTTAGATGGGGTCCTCAGGGACCTTGCTTTTCTCCTCAATGGCTGAAACAC
AACGCTAGCCTTAGGCCTGATGTGCAGTCATCTGCTGTGATGTTCGGTAGGGTTAACGAGCCTACCGCTCGGTCTTT
GCTTTTCAGGTATTGCGTTGGCAGGGCTGATGATGGTGGTGAAGCTGGTGCTGATACCAGGCGGTTTATTTTCCACG
AGCCATCTGATCTGGCCGAAGAGAATGTTCATACCTGCGGTGTGCTTATGGATGGTCACACTGGAATGGTGGGCGCT
TCTCTTGATATTCTTGTGTGCCCTAGGGACATCCACGGTTACCTTGCTCCAGTTCCTAAGACTCCTCTGGCCTTTTA
CGAGGTTAAGTGCAGGGCTAAGTACGCTTTCGATCCTATGGACCCTTCTGACCCTACTGCTTCTGCTTACGAGGATC
TGATGGCTCATAGAAGCCCTGAGGCTTTCAGGGCTTTCATCCGGTCTATTCCTAAGCCGAGCGTGAGATACTTTGCT
CCTGGAAGAGTTCCTGGTCCTGAGGAAGCTCTTGTTACTCAAGATCAGGCTTGGTCTGAGGCTCATGCTTCAGGTGA
GAAGAGAAGATGCTCAGCTGCTGATAGGGCACTCGTTGAGCTTAATTCTGGCGTGGTGTCTGAGGTGTTGCTTTTCG
GTGCTCCTGATCTCGGTAGGCACACTATTTCTCCAGTGAGCTGGTCCTCTGGTGATCTTGTTAGAAGGGAACCCGTG
TTCGCTAATCCTAGGCACCCTAACTTCAAGCAGATTCTGGTGCAGGGTTACGTGCTGGATTCTCACTTTCCAGATTG
CCCTCCACATCCTCACCTTGTGACTTTCATTGGTCGGCATAGGACCTCAGCTGAAGAGGGTGTTACTTTCAGGCTTG
AGGATGGTGCTGGTGCTCTTGGTGCTGCTGGTCCTTCTAAGGCTTCTATTCTTCCTAACCAGGCCGTGCCTATCGCT
CTTATTATCACCCCTGTGAGGATCGACCCCGAGATCTATAAGGCTATCCAGAGGTCATCTCGGCTGGCTTTCGATGA
TACTTTGGCTGAGCTTTGGGCCTCTAGATCTCCTGGTCCAGGTCCTGCTGCTGCAGAAACTACTTCTTCTTCACCTA
CCACCGGCAGGTCATCTAGAGGTTCGCTCGAG***CACCACCACCACCACCAC***TGAGATCCGG**CTGCTAACAAAGCCCGA**
**AAGGAAGCTGAGTTGGCTGCTGCCACCGCTGAGCAATAACTAGCATAACCCCTTGGGGCCTCTAAACGGGTCTTGAG**
**GGGTTTTTTGCTGAAAGGAGGAACTATATCCGGAT**TGGCGAATGGACGCGCCCTGTAGCGGCGCATTAAGCGCGGC
GGGTGTGGTGGTTACGCGCAGCGTGACCGCTACACTTGCCAGCGCCCTAGCGCCCGCTCCTTTCGCTTTCTTCCCTT
CCTTTCTCGCCACGTTCGCCGGCTTTCCCCGTCAAGCTCTAAATCGGGGGCTCCCTTTAGGGTTCCGATTTAGTGCT
TTACGGCACCTCGACCCCAAAAAACTTGATTAGGGTGATGGTTCACGTAGTGGGCCATCGCCCTGATAGACGGTTTT
TCGCCCTTTGACGTTGGAGTCCACGTTCTTTAATAGTGGACTCTTGTTCCAAACTGGAACAACACTCAACCCTATCT
CGGTCTATTCTTTTGATTTATAAGGATTTTGCCGATTTCGGCCTATTGGTTAAAAAATGAGCTGATTTAACAAAAA

TTTAACGCGAATTTTAACAAAATATTAACGTTTACAATTTCAGGTGGCACTTTTCGGGGAAATGTGCGCGGAACCCC
TATTTGTTTATTTTTCTAAATACATTCAAATATGTATCCGCTCATGAATTAATTCTTAGAAAAACTCATCGAGCATC
AAATGAAACTGCAATTTATTCATATCAGGATTATCAATACCATATTTTTGAAAAAGCCGTTTCTGTAATGAAGGAGA
AAACTCACCGAGGCAGTTCCATAGGATGGCAAGATCCTGGTATCGGTCTGCGATTCCGACTCGTCCAACATCAATAC
AACCTATTAATTTCCCCTCGTCAAAAATAAGGTTATCAAGTGAGAAATCACCATGAGTGACGACTGAATCCGGTGAG
AATGGCAAAAGTTTATGCATTTCTTTCCAGACTTGTTCAACAGGCCAGCCATTACGCTCGTCATCAAAATCACTCGC
ATCAACCAAACCGTTATTCATTCGTGATTGCGCCTGAGCGAGACGAAATACGCGATCGCTGTTAAAAGGACAATTAC
AAACAGGAATCGAATGCAACCGGCGCAGGAACACTGCCAGCGCATCAACAATATTTTCACCTGAATCAGGATATTCT
TCTAATACCTGGAATGCTGTTTTCCCGGGGATCGCAGTGGTGAGTAACCATGCATCATCAGGAGTACGGATAAAATG
CTTGATGGTCGGAAGAGGCATAAATTCCGTCAGCCAGTTTAGTCTGACCATCTCATCTGTAACATCATTGGCAACGC
TACCTTTGCCATGTTTCAGAAACAACTCTGGCGCATCGGGCTTCCCATACAATCGATAGATTGTCGCACCTGATTGC
CCGACATTATCGCGAGCCCATTTATACCCATATAAATCAGCATCCATGTTGGAATTTAATCGCGGCCTAGAGCAAGA
CGTTTCCCGTTGAATATGGCTCATAACACCCCTTGTATTACTGTTTATGTAAGCAGACAGTTTTATTGTTCATGACC
AAAATCCCTTAACGTGAGTTTTCGTTCCACTGAGCGTCAGACCCCGTAGAAAAGATCAAAGGATCTTCTTGAGATCC
TTTTTTTCTGCGCGTAATCTGCTGCTTGCAAACAAAAAAACCACCGCTACCAGCGGTGGTTTGTTTGCCGGATCAAG
AGCTACCAACTCTTTTTCCGAAGGTAACTGGCTTCAGCAGAGCGCAGATACCAAATACTGTCCTTCTAGTGTAGCCG
TAGTTAGGCCACCACTTCAAGAACTCTGTAGCACCGCCTACATACCTCGCTCTGCTAATCCTGTTACCAGTGGCTGC
TGCCAGTGGCGATAAGTCGTGTCTTACCGGGTTGGACTCAAGACGATAGTTACCGGATAAGGCGCAGCGGTCGGGCT
GAACGGGGGGTTCGTGCACACAGCCCAGCTTGGAGCGAACGACCTACACCGAACTGAGATACCTACAGCGTGAGCTA
TGAGAAAGCGCCACGCTTCCCGAAGGGAGAAAGGCGGACAGGTATCCGGTAAGCGGCAGGGTCGGAACAGGAGAGCG
CACGAGGGAGCTTCCAGGGGGAAACGCCTGGTATCTTTATAGTCCTGTCGGGTTTCGCCACCTCTGACTTGAGCGTC
GATTTTTGTGATGCTCGTCAGGGGGGCGGAGCCTATGGAAAAACGCCAGCAACGCGGCCTTTTTACGGTTCCTGGCC
TTTTGCTGGCCTTTTGCTCACATGTTCTTTCCTGCGTTATCCCCTGATTCTGTGGATAACCGTATTACCGCCTTTGA
GTGAGCTGATACCGCTCGCCGCAGCCGAACGACCGAGCGCAGCGAGTCAGTGAGCGAGGAAGCGGAAGAGCGCCTGA
TGCGGTATTTTCTCCTTACGCATCTGTGCGGTATTTCACACCGCATATATGGTGCACTCTCAGTACAATCTGCTCTG
ATGCCGCATAGTTAAGCCAGTATACACTCCGCTATCGCTACGTGACTGGGTCATGGCTGCGCCCCGACACCCGCCAA
CACCCGCTGACGCGCCCTGACGGGCTTGTCTGCTCCCGGCATCCGCTTACAGACAAGCTGTGACCGTCTCCGGGAGC
TGCATGTGTCAGAGGTTTTCACCGTCATCACCGAAACGCGCGAGGCAGCTGCGGTAAAGCTCATCAGCGTGGTCGTG
AAGCGATTCACAGATGTCTGCCTGTTCATCCGCGTCCAGCTCGTTGAGTTTCTCCAGAAGCGTTAATGTCTGGCTTC
TGATAAAGCGGGCCATGTTAAGGGCGGTTTTTTCCTGTTTGGTCACTGATGCCTCCGTGTAAGGGGGATTTCTGTTC
ATGGGGGTAATGATACCGATGAAACGAGAGAGGATGCTCACGATACGGGTTACTGATGATGAACATGCCCGGTTACT
GGAACGTTGTGAGGGTAAACAACTGGCGGTATGGATGCGGCGGGACCAGAGAAAAATCACTCAGGGTCAATGCCAGC
GCTTCGTTAATACAGATGTAGGTGTTCCACAGGGTAGCCAGCAGCATCCTGCGATGCAGATCCGGAACATAATGGTG
CAGGGCGCTGACTTCCGCGTTTCCAGACTTTACGAAACACGGAAACCGAAGACCATTCATGTTGTTGCTCAGGTCGC
AGACGTTTTGCAGCAGCAGTCGCTTCACGTTCGCTCGCGTATCGGTGATTCATTCTGCTAACCAGTAAGGCAACCCC
GCCAGCCTAGCCGGGTCCTCAACGACAGGAGCACGATCATGCGCACCCGTGGGGCCGCCATGCCGGCGATAATGGCC
TGCTTCTCGCCGAAACGTTTGGTGGCGGGACCAGTGACGAAGGCTTGAGCGAGGGCGTGCAAGATTCCGAATACCGC
AAGCGACAGGCCGATCATCGTCGCGCTCCAGCGAAAGCGGTCCTCGCCGAAAATGACCCAGAGCGCTGCCGGCACCT
GTCCTACGAGTTGCATGATAAAGAAGACAGTCATAAGTGCGGCGACGATAGTCATGCCCCGCGCCCACCGGAAGGAG
CTGACTGGGTTGAAGGCTCTCAAGGGCATCGGTCGAGATCCCGGTGCCTAATGAGTGAGCTAACTTACATTAATTGC
GTTGCGCTCACTGCCCGCTTTCCAGTCGGGAAACCTGTCGTGCCAGCTGCATTAATGAATCGGCCAACGCGCGGGGA
GAGGCGGTTTGCGTATTGGGCGCCAGGGTGGTTTTTCTTTTCACCAGTGAGACGGGCAACAGCTGATTGCCCTTCAC
CGCCTGGCCCTGAGAGAGTTGCAGCAAGCGGTCCACGCTGGTTTGCCCCAGCAGGCGAAAATCCTGTTTGATGGTGG
TTAACGGCGGGATATAACATGAGCTGTCTTCGGTATCGTCGTATCCCACTACCGAGATATCCGCACCAACGCGCAGC
CCGGACTCGGTAATGGCGCGCATTGCGCCCAGCGCCATCTGATCGTTGGCAACCAGCATCGCAGTGGGAACGATGCC
CTCATTCAGCATTTGCATGGTTTGTTGAAAACCGGACATGGCACTCCAGTCGCCTTCCCGTTCCGCTATCGGCTGAA
TTTGATTGCGAGTGAGATATTTATGCCAGCCAGCCAGACGCAGACGCGCCGAGACAGAACTTAATGGGCCCGCTAAC
AGCGCGATTTGCTGGTGACCCAATGCGACCAGATGCTCCACGCCCAGTCGCGTACCGTCTTCATGGGAGAAAATAAT
ACTGTTGATGGGTGTCTGGTCAGAGACATCAAGAAATAACGCCGGAACATTAGTGCAGGCAGCTTCCACAGCAATGG
CATCCTGGTCATCCAGCGGATAGTTAATGATCAGCCCACTGACGCGTTGCGCGAGAAGATTGTGCACCGCCGCTTTA
CAGGCTTCGACGCCGCTTCGTTCTACCATCGACACCACCACGCTGGCACCCAGTTGATCGGCGCGAGATTTAATCGC
CGCGACAATTTGCGACGGCGCGTGCAGGGCCAGACTGGAGGTGGCAACGCCAATCAGCAACGACTGTTTGCCCGCCA
GTTGTTGTGCCACGCGGTTGGGAATGTAATTCAGCTCCGCCATCGCCGCTTCCACTTTTTCCCGCGTTTTCGCAGAA
ACGTGGCTGGCCTGGTTCACCACGCGGGAAACGGTCTGATAAGAGACACCGGCATACTCTGCGACATCGTATAACGT
TACTGGTTTCACATTCACCACCCTGAATTGACTCTCTTCCGGGCGCTATCATGCCATACCGCGAAAGGTTTTGCGCC
ATTCGATGGTGTCCGGGATCTCGACGCTCTCCCTTATGCGACTCCTGCATTAGGAAGCAGCCCAGTAGTAGGTTGAG
GCCGTTGAGCACCGCCGCCGCAAGGAATGGTGCATGCAAGGAGATGGCGCCCAACAGTCCCCCGGCCACGGGGCCTG
CCACCATACCCACGCCGAAACAAGCGCTCATGAGCCCGAAGTGGCGAGCCCGATCTTCCCCATCGGTGATGTCGGCG

ATATAGGCGCCAGCAACCGCACCTGTGGCGCCGGTGATGCCGGCCACGATGCGTCCGGCGTAGAGGATCGAGATCTC
GATCCCGCGAAAT

**Table 4. Oligonucleotides used in this study**

| Oligo | Sequence (5´→3´) | Purpose | Figure |
|---|---|---|---|
| TMV_F | TTGGTCTCTACATGTAGCAGAACAAGCCTTAGAGATCAAG | Genotype TMV (P1) | Fig.1 |
| TMV_R | AAGGTCTCAACAACGTATAATCTTTCAAGGTCGTTTTTCTGTGAC | | |
| TMV-upstream_F | TTGGTCTCTACATCGTCCTGTGCCTGAATACCAGAG | Genotype TMV (P2) | Fig.1 |
| TMV-Do-upstream_R | AAGGTCTCAACAATCGGCGAAGGTACAGTATAGCTCG | | |
| TMV-Do-downstream_F | TTGGTCTCTACATGATGAGTGGCTAGCTGAGGTCTG | Genotype TMV (P3) | Fig.1 |
| TMV_downstream_R2 | AAGGTCTCAACAACTCTTTGCGTCCACGGTGGACAC | | |
| Nb_PGK-F4 | GAGCACACGCCTCTACAGAGGG | Genotype NbPGK (5´) | Fig.2 |
| GT_GUSi_R | TTCCCACCAACGCTGATCAATTCCAC | | |
| GUSi_F3 | ACCGCAGCAGGGAGGCAAAC | Genotype NbPGK (3´) | Fig.2 |
| PGK_OUT_R1 | GCCATCTTGCTATTGAACTCGTTGC | | |
| TPR_F1 | CCCACAAAGCATTCATGGAAACCATTCC | Genotype NbTPR (5´) | Fig.2 |
| GT_GUSi_R | TTCCCACCAACGCTGATCAATTCCAC | | |
| GUSi_F3 | ACCGCAGCAGGGAGGCAAAC | Genotype NbTPR (3´) | Fig.2 |
| TPR_R1 | CCTTCTGTCCTGTATGATTCCTAGCAG | | |
| THIC_For4 | TCTACAAGGTTCAGTGGGCAA | Genotyping synTHIC (5´) | |
| SynTHIC_Rev3 | CGCTGATGGTTTTCTTGGCAA | Genotyping synTHIC (5´) | |
| SynTHIC_For2 | TTGCCAAGAAAACCATCAGCG | Genotyping | |
| pDAP101_Rev1 | CGCCTGGTATCTTTATAGTCCTGTCG | Genotyping synTHIC (3´) | |
| GL1_For2 | GGCCATAGATACAATTAAACCAACTG | GL1 genotyping | |
| GL1_Rev1 | GAGGAGCTTGTGGAGACGAAT | GL1 genotyping | |
| Cas9_6F | TTGGTCTCTACATTACGGAGGCTTTGATAGCCCTACC | Genotyping Cas9 presence | |
| tNos_Rev21 | TGTTTGAACGATCTGCTTGA | Genotyping Cas9 presence | |
| Oregon Green-labeled hairpin oligo | GGAAGGGCCCGCTGACAGTTTTTCTGTCAGCGGGCCCTTCC | Substrate for *in vitro* Exo assay | Fig.3 |

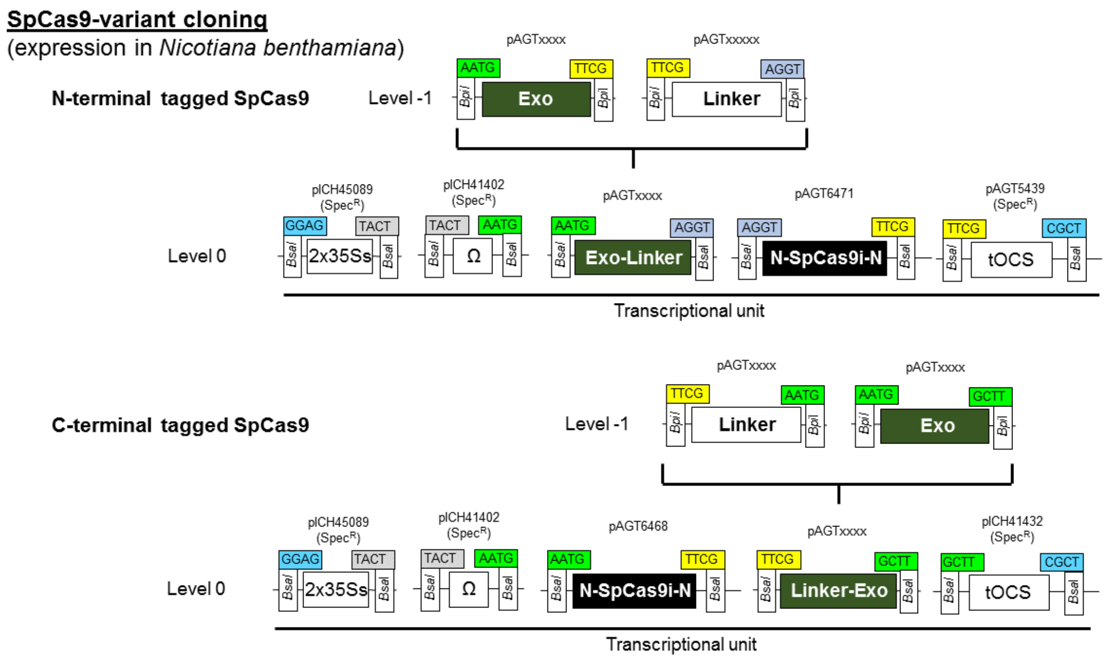
SpCas9-variant cloning
(expression in Nicotiana benthamiana)
N-terminal tagged SpCas9
Level -1
pAGTxxxx
AATG
Exo
TTCG
Bpi
pAGTxxxxx
TTCG
Linker
AGGT
Level 0
pICH45089
(SpecR)
GGAG
2x35Ss
TACT
pICH41402
(SpecR)
TACT
Ω
AATG
pAGTxxxx
AATG
Exo-Linker
AGGT
pAGT6471
AGGT
N-SpCas9i-N
TTCG
pAGT5439
(SpecR)
TTCG
tOCS
CGCT
BsaI
Transcriptional unit
C-terminal tagged SpCas9
Level -1
pAGTxxxx
TTCG
Linker
AATG
pAGTxxxx
AATG
Exo
GCTT
Level 0
pICH45089
(SpecR)
GGAG
2x35Ss
TACT
pICH41402
(SpecR)
TACT
Ω
AATG
pAGT6468
AATG
N-SpCas9i-N
TTCG
pAGTxxxx
TTCG
Linker-Exo
GCTT
pICH41432
(SpecR)
GCTT
tOCS
CGCT
Transcriptional unit

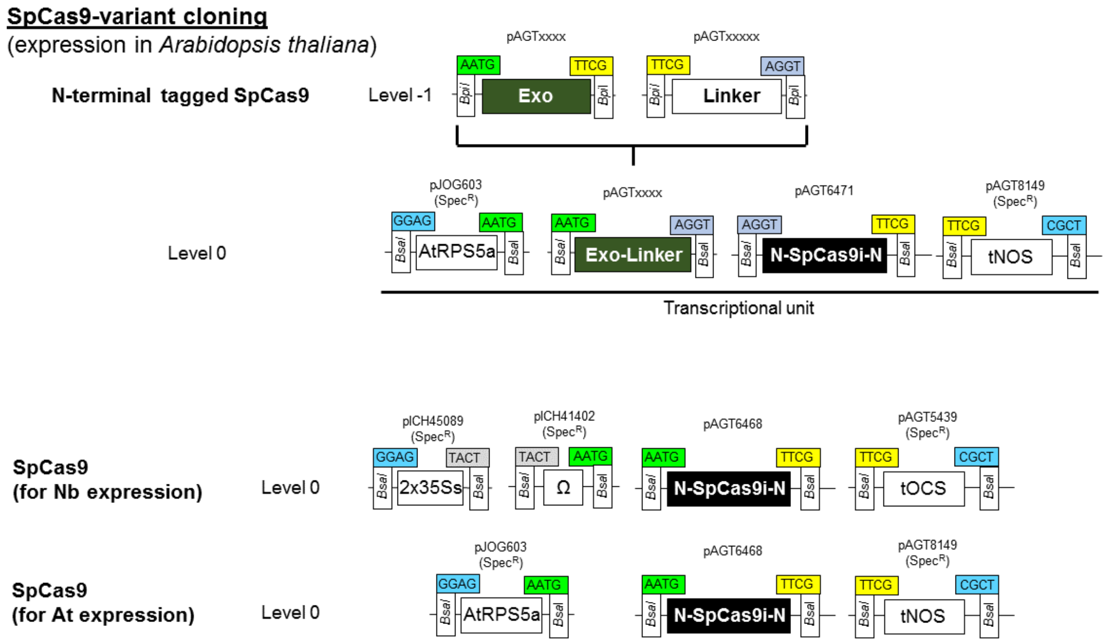
SpCas9-variant cloning
(expression in Arabidopsis thaliana)
N-terminal tagged SpCas9
Level -1
pAGTxxxx
AATG
Exo
TTCG
pAGTxxxxx
TTCG
Linker
AGGT
Level 0
pJOG603
(SpecR)
GGAG
AtRPS5a
AATG
pAGTxxxx
AATG
Exo-Linker
AGGT
pAGT6471
AGGT
N-SpCas9i-N
TTCG
pAGT8149
(SpecR)
TTCG
tNOS
CGCT
Transcriptional unit
SpCas9
(for Nb expression)
Level 0
pICH45089
(SpecR)
GGAG
2x35Ss
TACT
pICH41402
(SpecR)
TACT
Ω
AATG
pAGT6468
AATG
N-SpCas9i-N
TTCG
pAGT5439
(SpecR)
TTCG
tOCS
CGCT
SpCas9
(for At expression)
Level 0
pJOG603
(SpecR)
GGAG
AtRPS5a
AATG
pAGT6468
AATG
N-SpCas9i-N
TTCG
pAGT8149
(SpecR)
TTCG
tNOS
CGCT

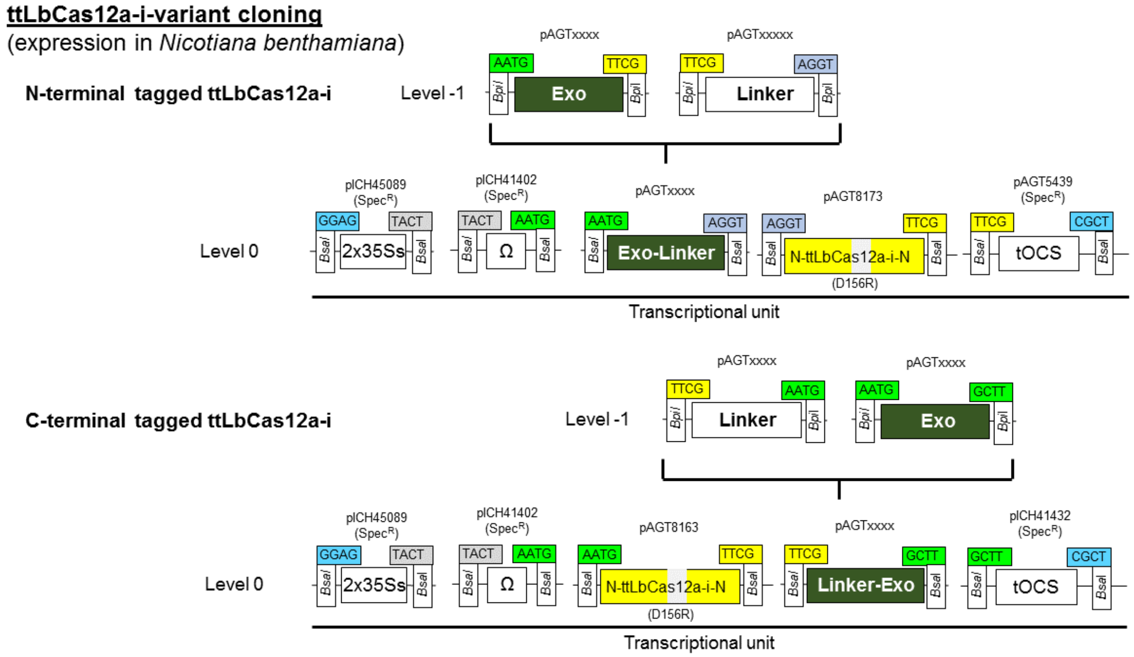
ttLbCas12a-i-variant cloning
(expression in Nicotiana benthamiana)
N-terminal tagged ttLbCas12a-i
Level -1
pAGTxxxx
pAGTxxxxx
AATG
TTCG
AGGT
Exo
Linker
Bpi
Level 0
pICH45089 (Spec^R)
pICH41402 (Spec^R)
pAGTxxxx
pAGT8173
pAGT5439 (Spec^R)
GGAG
TACT
AATG
CGCT
2x35Ss
Ω
Exo-Linker
N-ttLbCas12a-i-N
(D156R)
tOCS
BsaI
Transcriptional unit
C-terminal tagged ttLbCas12a-i
GCTT
pAGT8163
pICH41432 (Spec^R)
Linker-Exo

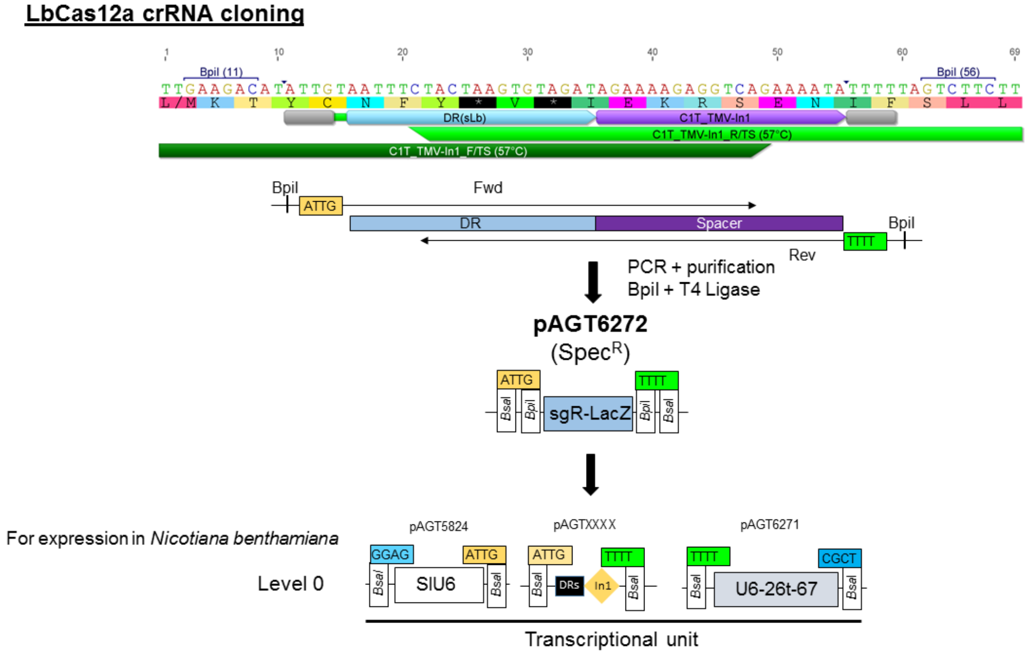
LbCas12a crRNA cloning
BpiI (11)
BpiI (56)
DR(sLb)
C1T_TMV-In1
C1T_TMV-In1_R/TS (57°C)
C1T_TMV-In1_F/TS (57°C)
BpiI
ATTG
Fwd
DR
Spacer
TTTT
Rev
PCR + purification
BpiI + T4 Ligase
pAGT6272
(Spec^R)
sgR-LacZ
For expression in Nicotiana benthamiana
Level 0
pAGT5824
pAGTXXXX
pAGT6271
GGAG
CGCT
SIU6
DRs
In1
U6-26t-67
Transcriptional unit

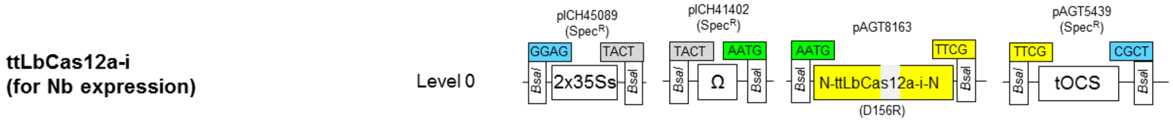
ttLbCas12a-i
(for Nb expression)
Level 0
pICH45089
(SpecR)
GGAG
TACT
BsaI
2x35Ss
pICH41402
(SpecR)
TACT
AATG
Ω
pAGT8163
AATG
TTCG
N-ttLbCas12a-i-N
(D156R)
pAGT5439
(SpecR)
TTCG
CGCT
tOCS

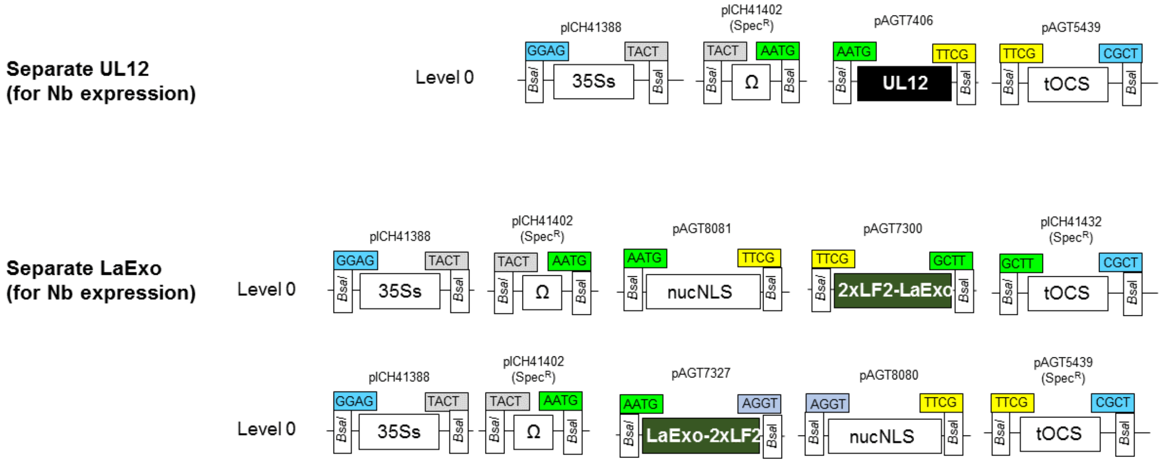
Separate UL12
(for Nb expression)
Level 0
pICH41388
GGAG
TACT
BsaI
35Ss
pICH41402
(SpecR)
TACT
AATG
Ω
pAGT7406
AATG
TTCG
UL12
pAGT5439
TTCG
CGCT
tOCS
Separate LaExo
(for Nb expression)
Level 0
pICH41388
GGAG
TACT
35Ss
pICH41402
(SpecR)
TACT
AATG
Ω
pAGT8081
AATG
TTCG
nucNLS
pAGT7300
TTCG
GCTT
2xLF2-LaExo
pICH41432
(SpecR)
GCTT
CGCT
tOCS
Level 0
pICH41388
GGAG
TACT
35Ss
pICH41402
(SpecR)
TACT
AATG
Ω
pAGT7327
AATG
AGGT
LaExo-2xLF2
pAGT8080
AGGT
TTCG
nucNLS
pAGT5439
(SpecR)
TTCG
CGCT
tOCS

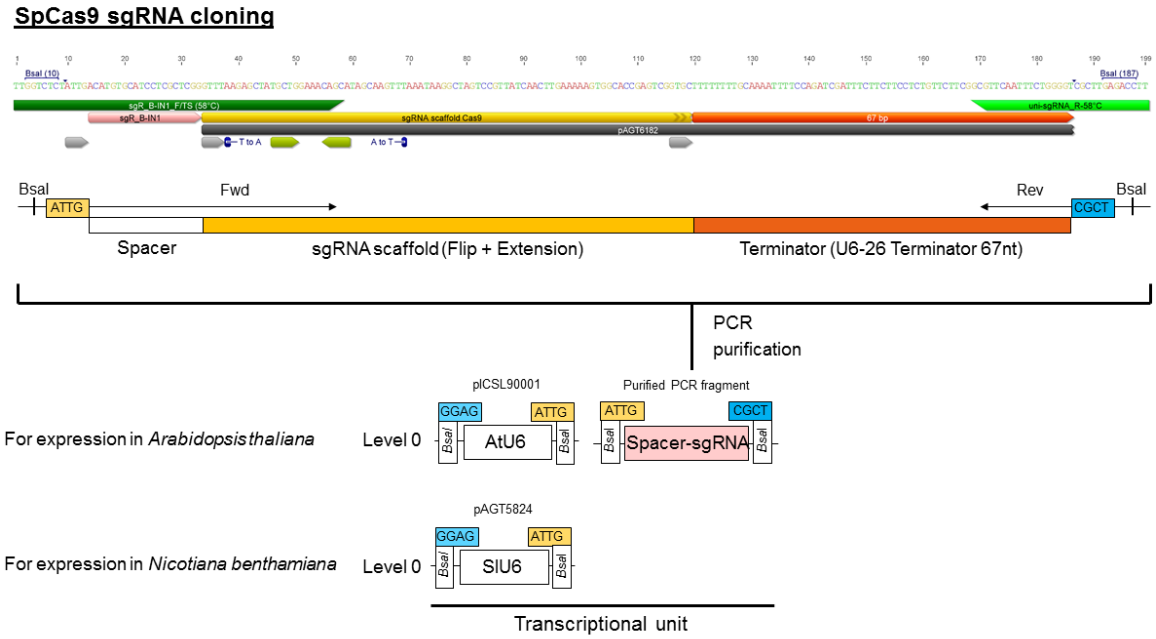
SpCas9 sgRNA cloning
BsaI
ATTG
Fwd
Rev
CGCT
BsaI
Spacer
sgRNA scaffold (Flip + Extension)
Terminator (U6-26 Terminator 67nt)
PCR
purification
pICSL90001
Purified PCR fragment
For expression in Arabidopsis thaliana
Level 0
GGAG
ATTG
AtU6
ATTG
CGCT
Spacer-sgRNA
pAGT5824
For expression in Nicotiana benthamiana
Level 0
GGAG
ATTG
SlU6
Transcriptional unit

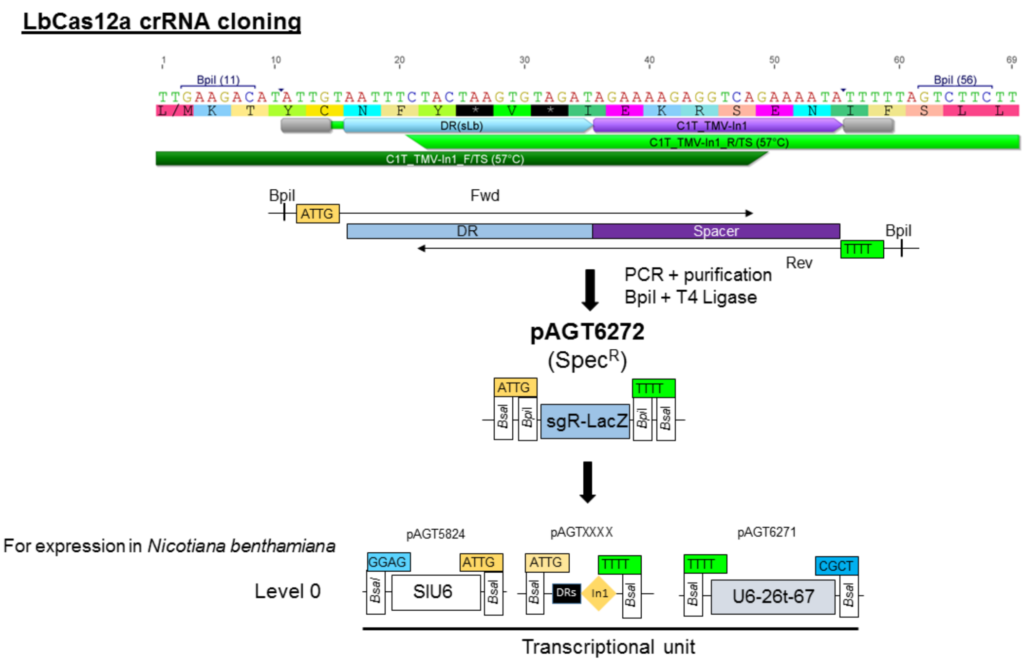
LbCas12a crRNA cloning
BpiI (11)
BpiI (56)
DR(sLb)
C1T_TMV-In1
C1T_TMV-In1_R/TS (57°C)
C1T_TMV-In1_F/TS (57°C)
BpiI
ATTG
Fwd
DR
Spacer
BpiI
TTTT
Rev
PCR + purification
BpiI + T4 Ligase
pAGT6272
(SpecR)
ATTG
TTTT
sgR-LacZ
For expression in Nicotiana benthamiana
Level 0
pAGT5824
GGAG
ATTG
SlU6
pAGTXXXX
ATTG
TTTT
DRs
In1
pAGT6271
TTTT
CGCT
U6-26t-67
Transcriptional unit

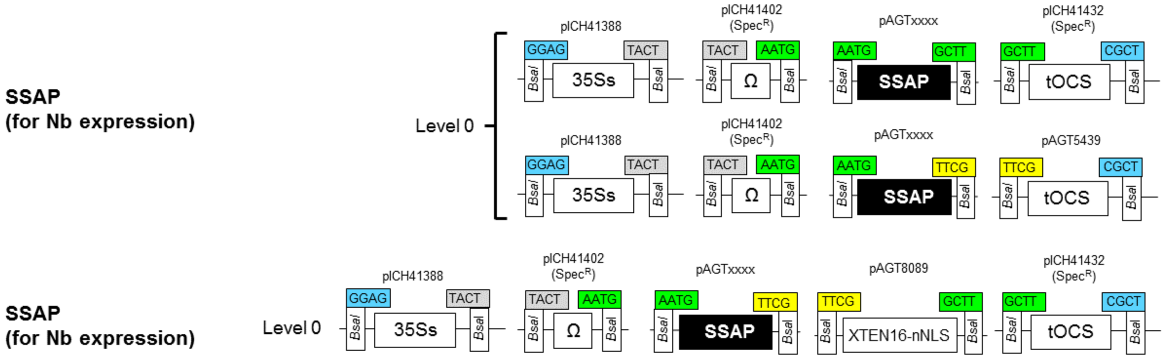
SSAP
(for Nb expression)
Level 0
pICH41388
GGAG
TACT
35Ss
pICH41402
(SpecR)
TACT
AATG
Ω
pAGTxxxx
AATG
GCTT
SSAP
pICH41432
(SpecR)
GCTT
CGCT
tOCS
pICH41388
35Ss
pICH41402
(SpecR)
Ω
pAGTxxxx
AATG
TTCG
SSAP
pAGT5439
TTCG
CGCT
tOCS
SSAP
(for Nb expression)
Level 0
pICH41388
GGAG
TACT
35Ss
pICH41402
(SpecR)
TACT
AATG
Ω
pAGTxxxx
AATG
TTCG
SSAP
pAGT8089
TTCG
GCTT
XTEN16-nNLS
pICH41432
(SpecR)
GCTT
CGCT
tOCS

**Cloning for E.coli expression**

***E.coli* codon optimized UL12 (pAGT8228)**
**(for UL12 protein purification after recombinant expression in *E.coli*)**

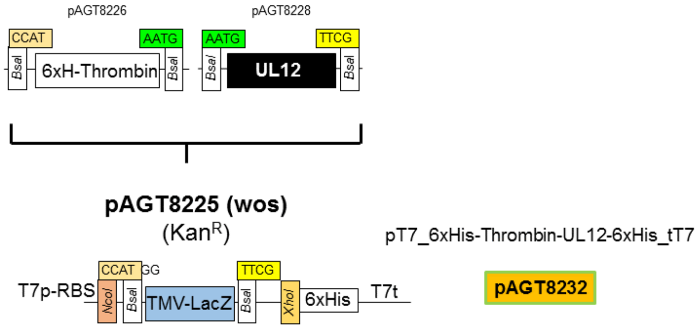
